# Emergence of zoonotic and multi-drug resistant *Streptococcus suis*

**DOI:** 10.1101/2025.06.30.662319

**Authors:** Jaime Brizuela, Gemma G. R. Murray, Parichart Boueroy, Andrew J. Balmer, Thidathip Wongsurawat, Piroon Jenjaroenpun, Peechanika Chopjitt, Rujirat Hatrongjit, Lucy A. Weinert, Anusak Kerdsin, Constance Schultsz

## Abstract

**Background:** *Streptococcus suis* is an emerging zoonotic porcine pathogen and a leading cause of adult bacterial meningitis in Southeast Asia, associated with raw pork consumption. Most zoonotic *S. suis* infections globally are by strains from lineage CC1 carrying a serotype 2 capsule. However, in Thailand, ∼40% of the reported zoonotic infections are caused by two endemic lineages, CC104 and CC233 which also have a serotype 2 capsule. In this study, we aimed to identify the drivers of the emergence and recent evolution of these two lineages.

**Methods:** We sequenced the whole genomes of 141 Thai *S. suis* zoonotic and porcine strains isolated over a 15-year period and combined them with a curated global dataset of 2761 published *S. suis* genomes. Using comparative genomics, Bayesian evolutionary models and multivariate analysis we investigated the emergence of zoonotic potential and multi-drug resistance in CC104 and CC233.

**Findings:** We estimated recent emergence dates for both CC104 (1990; 95% posterior: 1987-1992) and CC233 (2002; 95% posterior: 2000-2004). Both lineages acquired a serotype capsule 2 from CC1 through a capsule locus switching event, prior to their emergence. Both have also experienced multiple antimicrobial resistance (AMR) acquisition events, with some strains carrying 12 determinants encoding resistance against eight classes of antibiotics. Most importantly, CC104 and CC233 lineages are the first zoonotic lineages to have acquired increased resistance to penicillin and ceftriaxone, which form the standard therapy to treat *S. suis* infections in humans.

**Interpretation:** Horizontal transfer of multiple genomic regions can cause rapid emergence of novel multi-drug-resistant zoonotic *S. suis* lineages. As *S. suis* is mainly controlled and treated through the use of antibiotics in both pigs and humans, these findings highlight the urgent need for improved and enhanced surveillance, infection control, and treatments.

## INTRODUCTION

*Streptococcus suis* is an important porcine pathogen and is increasingly recognized as an emerging zoonotic pathogen. It causes invasive infections in both pigs and humans, with zoonotic infections typically resulting in meningitis, sepsis, high morbidity, and mortality.^1,2^ In Europe, zoonotic *S. suis* infections are mainly an occupational hazard for professions in contact with pigs or pork. In this setting, *S. suis* is typically transmitted through direct skin contact, with infection resulting from contact with skin lesions. However, in several Southeast Asian countries, *S. suis* is mainly transmitted through consumption of raw pork.^2,3^ In Thailand and Vietnam, human *S. suis* infections have become endemic and *S. suis* has caused multiple foodborne outbreaks with high mortality rates in China, Indonesia, and Thailand.^4–7^

*S. suis* can be classified into serotypes based on its capsular polysaccharides structure, and into sequence types (STs) which can be further grouped into clonal complexes (CC), based on the DNA sequences of seven housekeeping genes.^1,8^ The population structures and evolution of the ten *S. suis* CCs that are most commonly associated with disease in pigs globally were recently described and their estimated emergence dates range between mid-19^th^ century and mid-20^th^ century.^9^ The large majority of global zoonotic infections are caused by isolates in lineage CC1.^1^ Furthermore, while a diverse set of serotypes is associated with disease in pigs, ∼95% of the reported human infections are caused by strains with a serotype 2 capsule.^1^

Thailand has the highest estimated prevalence of zoonotic *S. suis* infections and has seen a recent emergence of two novel zoonotic lineages.^4^ In contrast to global human infections, infections in Thailand are caused by an unusually diverse set of lineages. Between 2006 and 2021 CC1 accounted for ∼55% of the reported zoonotic infections in Thailand, while 40% of the human infections were caused by two novel zoonotic lineages known as CC104 and CC233 which are now endemic to Thailand.^10^ In this study, we aimed to identify the drivers of the emergence and recent evolution of these two lineages. Using both short and long read whole genome sequencing of 141 Thai *S. suis* zoonotic and porcine strains isolated over a 15-year period, combined with a curated global dataset of 2761 published *S. suis* genomes, we show that independent horizontal transmission of genes encoding the polysaccharide capsule as well as genes encoding antibiotic resistance, has driven the recent evolution and emergence of these lineages.

## MATERIALS AND METHODS

### Thai strain collection and selection

The 141 *S. suis* isolates that were sequenced and analysed in this study were selected from 1120 isolates sampled from Thailand over 15 years (2006-2021) from three collections. Firstly, 873 zoonotic isolates were collected in a nation-wide active surveillance study between 2006 and 2012 (Supplementary Table 1.). Secondly, 141 zoonotic *S. suis* isolates were collected during a passive surveillance study in Northern and Northeast Thailand between 2013 and 2021 (Supplementary Table 2.) Finally, 106 isolates were collected from healthy pigs in Phayao province (Northern Thailand) between 2010 and 2011 (Supplementary Table 3.). Serotyping and multi-locus sequence typing (MLST) were performed previously and confirmed *in silico* for the sequenced isolates using the *S. suis* serotyping pipeline and mlst v2.19.0 (https://github.com/tseemann/mlst) with the PubMLST database (https://pubmlst.org/) and the *S. suis* MLST scheme.^8,11^

We performed stratified random sampling of the 1120 isolates to select 141 representative isolates for Illumina WGS and further analysis (Supplementary Figure 1; Supplementary Methods). We selected 121 zoonotic isolates and 20 isolates from healthy pigs, of which 10 were isolates with serotypes of interest (2, 7, and 14) and ten were randomly selected to give a total of 141 isolates (Supplementary Figure 1). Since there were no publicly available complete genomes from lineage CC104, we selected four isolates from the zoonotic lineage CC104 for additional long read sequencing using the Oxford Nanopore Technologies (ONT) platform. We combined these 141 isolates with 123 publicly available *S. suis* genomes from Thailand for genomic analyses (total n=264) (Supplementary Table 4, Supplementary Methods). Furthermore, we set the Thai isolates in the global context by combining them with a curated global collection of *S. suis* genomes (n = 2638) (Supplementary Methods).

### Modelling the evolution of lineages CC104 and CC233

To estimate the date of emergence of each lineage, we constructed two dated phylogenies for CC104 and CC233 respectively. We reconstructed two genome-wide SNP alignments by mapping the trimmed Illumina reads to the complete genomes of STC228 (CC104) and STC78 (CC233) using Snippy v4.6.0 (https://github.com/tseemann/snippy) and removed recombination from the alignment using Gubbins v3.3.1.^12^ We constructed dated phylogenies using BEAST v1.10 with a GTR+G model, a strict molecular clock and a constant population size.^13^ For isolates with an unknown year of isolation, we set a range spanning the dataset’s range. Additionally, to model the evolutionary history of both lineages we constructed a genome-wide SNP phylogeny as described above (using STC78 as a reference) including all CC104, CC233 as well as their closest related isolates from our global collection (n=130). Then, we used BEAST v1.10 to fit both a symmetric and an asymmetric discrete trait model to the posterior distribution of trees to model changes in serotype and country of isolation for both lineages (Supplementary Table 7).

**Analysis of Penicillin Binding Proteins (PBPs) alleles and β-lactam resistance**

We extracted the amino acid sequences of Pbp2B, Pbp2X, and MraY from the 264 *S. suis* Thai genomes to investigate whether the zoonotic lineages CC104 and CC233 had acquired PBP types conferring increased resistance against β-lactams. We selected these three proteins as a previous study identified key amino acid substitutions associated with increased β-lactam resistance in their amino acid sequences.^14^ We concatenated and aligned the amino acid sequences of the three genes using MAFFT v7.525,^15^ reconstructed an amino acid phylogeny using FastTree v2.1.11,^16^ and identified distinct PBP types. We made a representative selection of isolates based on their PBP types and tested their susceptibility against penicillin and ceftriaxone by determining the minimum inhibitory concentration (MIC) following CLSI 2023 guidelines (M100-33^rd^ edn) (Supplementary Methods).

Given the multidimensional nature of resistance against β-lactams in *S. suis* we used dimensionality reduction methods to systematically categorise isolates based on their PBP types (manuscript in preparation). We transformed the PBP amino acid alignment into a symmetrical matrix of the pairwise amino acid distances between isolates. Then, we used multidimensional scaling iterating with the SMACOF algorithm to reduce the data into a 2-dimensional map and used hierarchical clustering to classify isolates based on their PBP types. Finally, we used Spearman’s rank correlation to test for an association between diverging from the most common susceptible PBP type (PBP type “1”) and increased penicillin and ceftriaxone MICs.

For phenotypic susceptibility testing we used broth microdilution to determine the MIC of penicillin and ceftriaxone, following the CLSI guidelines (Supplementary Table 6) and the *S. suis* MIC breakpoints for penicillin following CLSI Veterinary medicine (VET01-7^th^ edn). *S. suis* does not have established ceftriaxone MIC breakpoints.

### Supplemental information

Additional methods on Illumina and Nanopore sequencing, assembly and annotation, blasts and phylogenies can be found in the Supplementary Methods.

### Role of the funding source

The funders did not play any role during the study design, collection of data, analysis, interpretation, writing of the report or decision to submit the study for publication.

## RESULTS

### The zoonotic Thai lineages CC104 and CC233 are part of the pathogenic *S. suis* population

The population structure of the Thai *S. suis* isolates resembles the global *S. suis* structure and includes isolates from most of the common global pathogenic clades (only CC17, CC20, and CC87 are absent) revealing the diversity of pathogenic and zoonotic *S. suis* in Thailand (Figure 1). We find that zoonotic infections in Thailand are caused by isolates from diverse globally distributed porcine pathogenic lineages in addition to two related Thai lineages CC104 and CC233 (Figure 1A). CC104 and CC233 predominantly carry a serotype 2 capsule and appear to be geographically restricted to Thailand, with the closest relatives being a set of double locus variants of serotype 7 recovered from diseased pigs in North America, Europe and Asia (Figure 1B). CC104 and CC233 both carry three recently identified pathogenicity-associated genomic islands that typify the main pathogenic *S. suis* lineages further supporting their placement within the diversity of global pathogenic lineages (Supplementary Figure 2).^9^ In addition, there are also two clades of isolates (CC221/234 and CC1688) associated with human infections that fall outside of the pathogenic population of *S. suis.* In particular, CC1688 has a serotype 2 capsule and may represent an emerging lineage.

**Figure 1.**
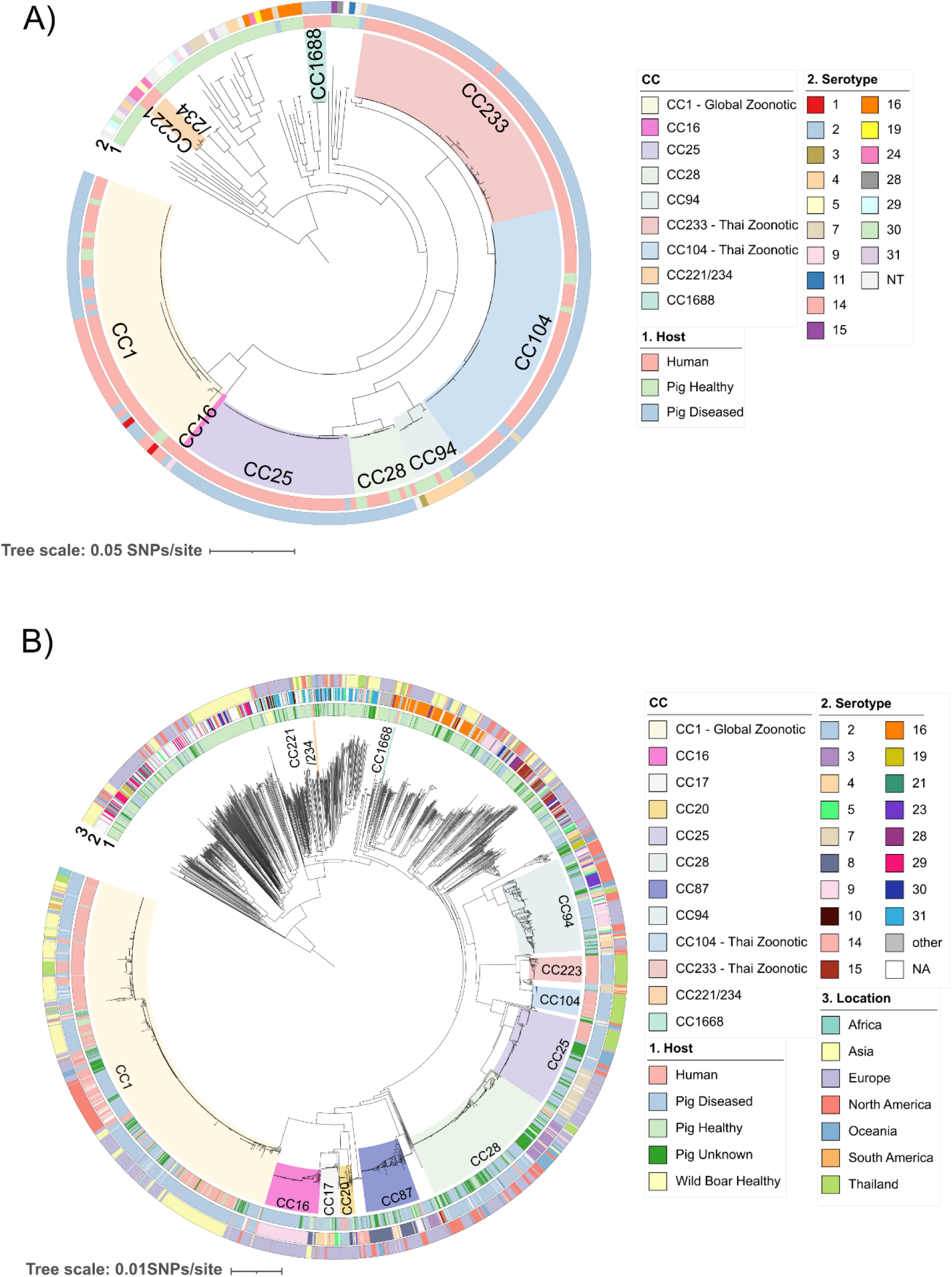
Population structure of Thai *S. suis*. A) Mid-point rooted maximum-likelihood (ML) core genome phylogeny of 264 Thai *S. suis* genomes constructed using Panaroo and IQ-TREE. Of the 264 genomes, 141 were sequenced during this study and 123 are publicly available genomes. Colours of highlighted clades indicate CC. The inner coloured ring (1) indicates host from which the sample was isolated and the outer coloured ring (2) indicates serotype. B) Global *S. suis* mid-point rooted core-genome phylogeny based on 2902 *S. suis* genomes constructed using Panaroo and FastTree. The colours of the highlighted clades indicate CC, the innermost coloured ring (1) indicates host from which the sample was isolated, the middle-coloured ring (2) indicates serotype, and the outer coloured ring (3) indicates continent/country of isolation. CC: Clonal complex, NT: Non-typable, NA: Data not available.

### The CC104 and CC233 lineages emerged as zoonotic in the last three decades following two independent serotype 2 acquisition events from CC1

We used the temporal distribution of our isolates to estimate the date of emergence and model the gain of zoonotic potential of CC104 and CC233. Both lineages have emerged recently: the dates of the most recent common ancestor (MRCA) of CC104 and CC233 were 1990 (95% Posterior 1987-1992) and 2002 (95% Posterior 2000-2004) (Figure 2A-C). These emergence dates are 7-8 years before the first reports of zoonotic infections caused by isolates belonging to both clades (1998 for CC104 and 2009 for CC233) (Figure 2A-C; Supplementary Table 4).^17^ Phylogeographic analysis suggests that the MRCA of CC104 and CC233 originated in Asia, however, uneven and sparse sampling across different geographic regions make these reconstructions difficult, with none of the ancestral state posterior probabilities being high (Supplementary Figure 3).

**Figure 2.**
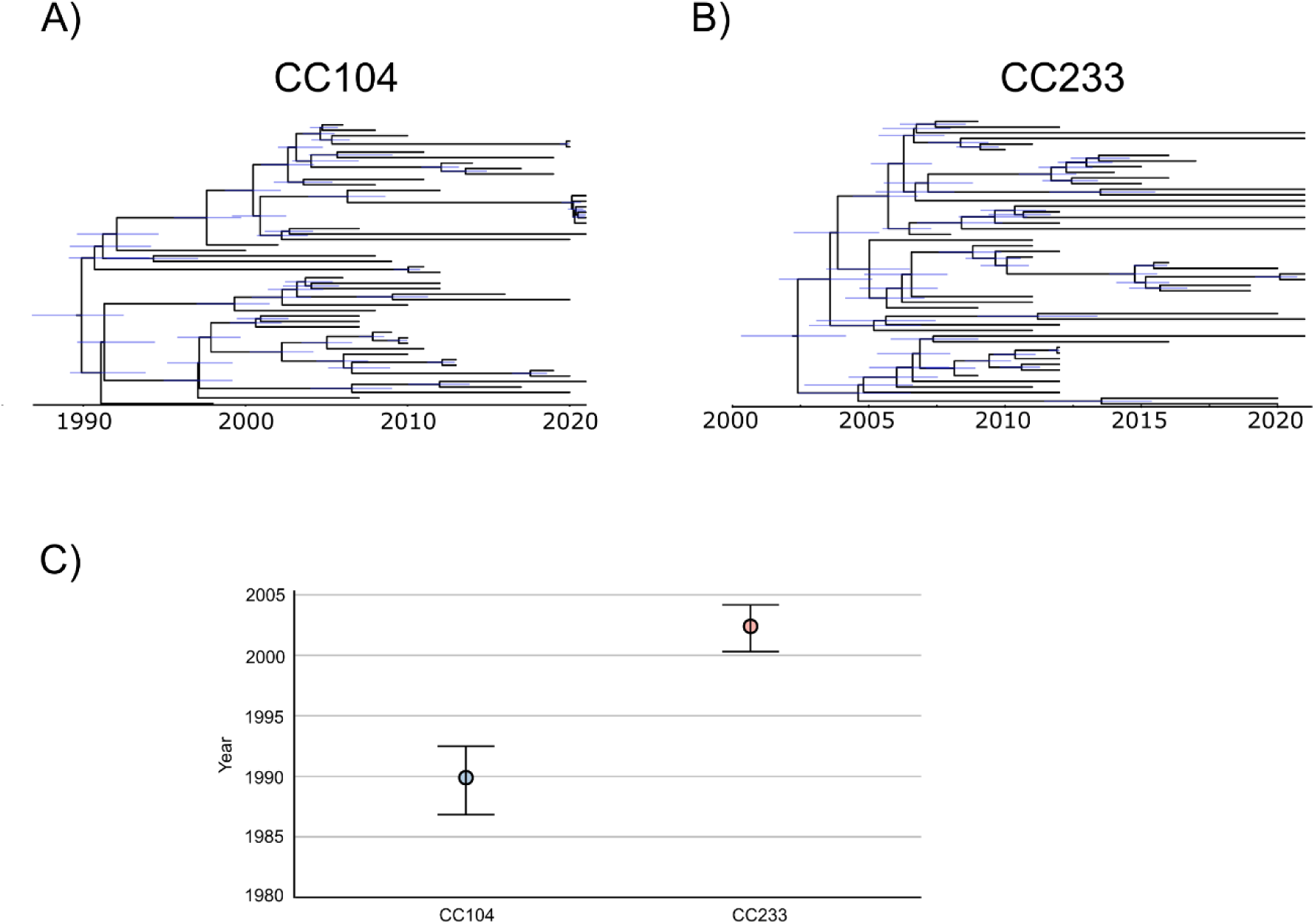
Lineages CC104 and CC233 have emerged recently. A-B) Dated phylogenies of the Thai zoonotic lineages CC104 and CC233 respectively. The dated phylogenies were reconstructed using BEAST with a strict molecular clock, GTR+G substitution model and constant population model. STC228 was used as a reference for the CC104 and STC78 for CC233. Node bars indicate 95 % confidence interval (CI) for inferred node ages. C) Age estimates for the emergence of the zoonotic lineages CC104 and CC233 based on their dated phylogenies. Error bars indicate the upper and lower bounds of the 95% CI.

Both CC104 and CC233 appear to have undergone capsular switching from a serotype 7 to a serotype 2 capsule, which is associated with gain of zoonotic potential (Figure 3A).^18^ We performed an ancestral state reconstruction of the capsule locus to model these capsular switching events. Our findings suggest that MRCA for both the CC104 and CC233 lineages was likely a serotype 7 strain and the expansion of both CC104 and CC233 was preceded by two capsule switching events where both lineages acquired a serotype 2 capsule (Supplementary Figure 4). Next, we investigated the origin of the serotype 2 capsule present in both CC104 and CC233. To do so, we reconstructed a serotype 2 capsular locus phylogeny including all serotype 2 isolates from Thailand, supplemented with serotype 2 isolates from our global collection (Figure 3B; Supplementary Methods). This showed that the serotype 2 capsule loci of CC104 and CC233 fall within the diversity of the serotype 2 capsule loci of CC1. Comparisons of mean pairwise nucleotide distance in core genes and those within the serotype 2 capsule locus revealed that despite being very divergent at a core genome level, genes within the serotype 2 capsule loci of CC1, CC104 and, CC233 were almost identical, with only 1 SNP difference between the most common serotype 2 locus sequence found in CC1 and the most common serotype 2 locus found in CC104 and CC233 (Figure 3C). Coupled with the fact that the published estimate of the date for CC1’s MRCA precedes our estimated MRCA dates for CC104 and CC233 (1850s, vs 1990 and 2002) suggests that CC104 and CC233 have acquired their serotype 2 locus from CC1.^9^ As many CC104 and CC233 isolates have identical serotype 2 loci, it is possible that it may have transferred from CC1 to one of these lineages and then from that lineage onto the other (Figure 3B). Our results also reveal other instances of the horizontal transfer of the serotype 2 capsule locus from CC1 to other CCs, revealing that these transfers are not isolated events (Figure 3B).

**Figure 3.**
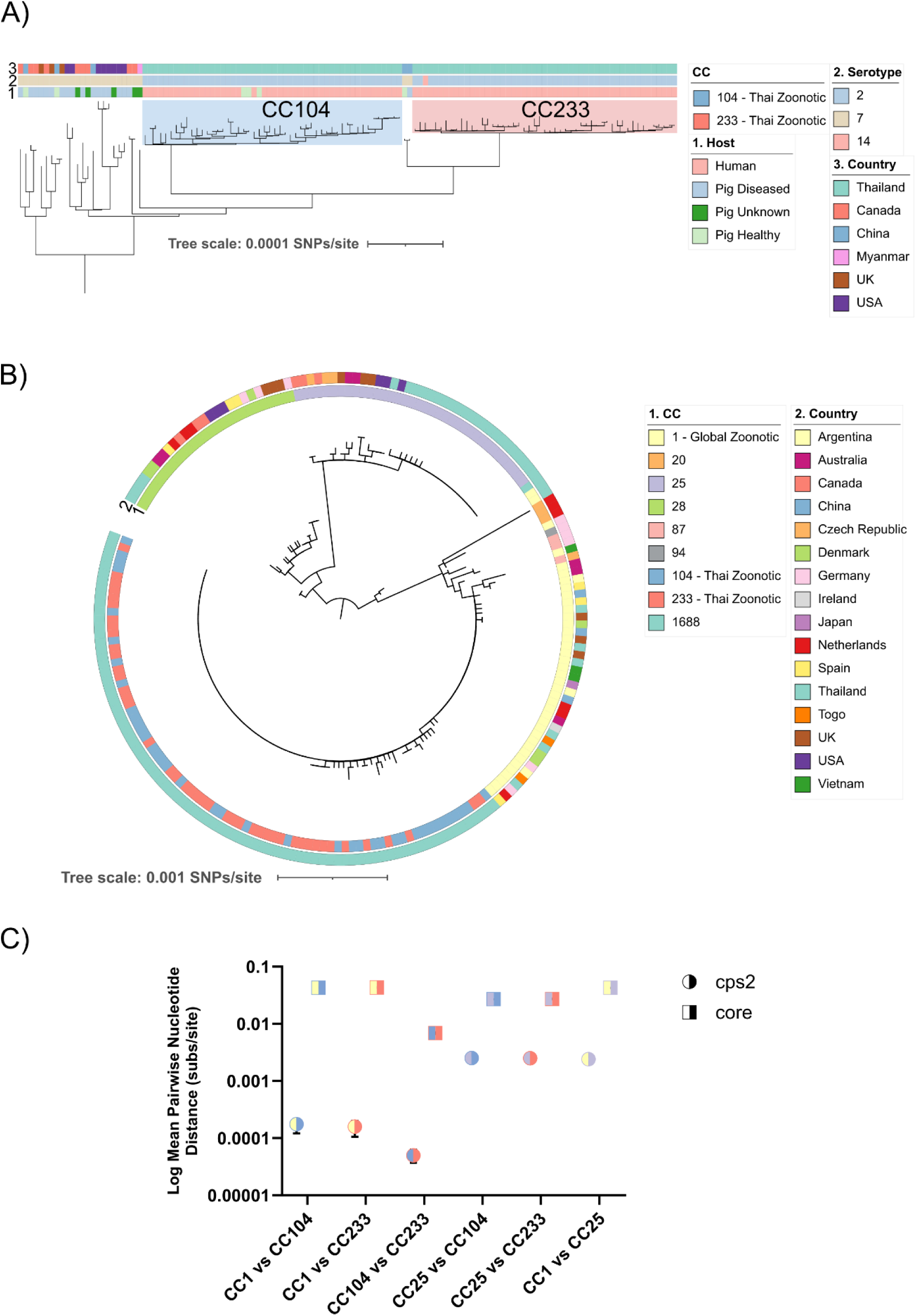
CC104 and CC233 acquired a serotype 2 capsular locus from CC1. A) SNP reference mapped phylogeny of CC104 and CC233 and their closest related strains. Colours of highlighted clades indicate CC. indicates host from which the sample was isolated, the middle-coloured ring (2) indicates serotype and the outer coloured ring (3) indicates the Country where the sample was isolated. B) Serotype 2 capsule locus phylogeny based on a concatenated nucleotide alignment of all capsule locus genes. Inner coloured ring (1) indicates CC and outer Coloured ring (2) indicates country of isolation. C) Mean pairwise nucleotide distance between different CCs based on the serotype 2 capsule locus genes and core genes. Error bars represent confidence intervals calculated by re-estimating the mean pairwise distance with 1000 bootstraps. Circles indicate mean pairwise distance between serotype 2 locus genes and squares represent mean pairwise distance between core genes. Symbols are coloured with the colour schemes of the CCs that are being compared.

### The CC104 lineage has acquired many antimicrobial resistance determinants through mobile genetic elements

Acquisition of antimicrobial resistance (AMR) genes through horizontal gene transfer can increase the fitness of novel bacterial lineages in settings of high antibiotic usage such as pig farms in Thailand.^19^ We therefore examined whether AMR could have promoted the emergence of CC104 and CC233 in Thailand. We found that these two lineages carry several antibiotic resistance determinants that are rarely observed in other pathogenic lineages of *S. suis*. For example, they carry a full-length copy of the chromosomal gene *vgaF*, which is associated with increased tiamulin resistance,^14^ whereas the this gene was truncated in the global pathogenic lineages CC1, CC16, CC25, CC28, and CC94 (Figure 4A).

**Figure 4.**
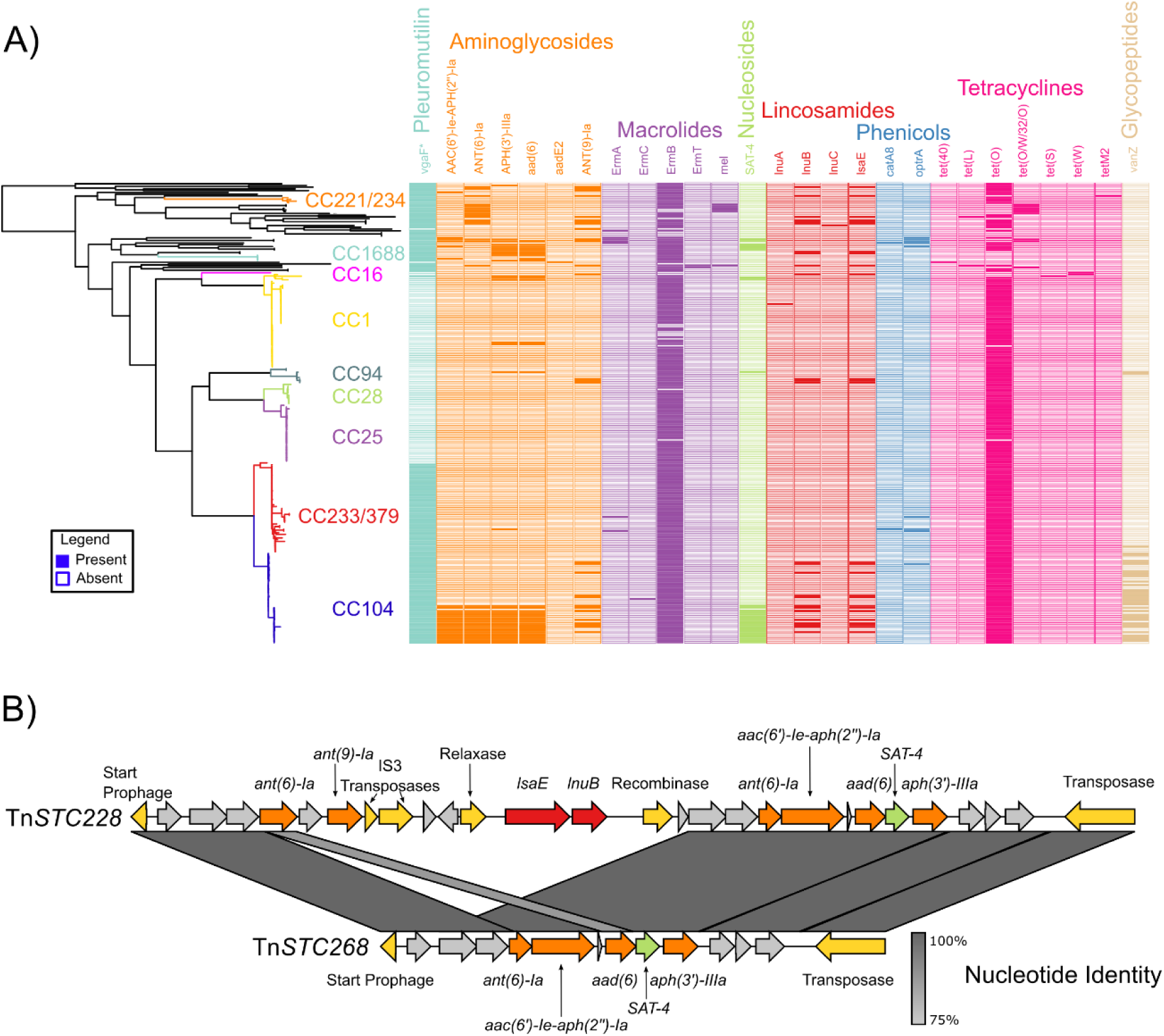
Horizontal Acquisition of AMR determinants by the zoonotic Thai *S. suis* lineages. A) Thai *S. suis* core genome phylogeny from Figure 1 visualized with a presence/absence matrix of AMR determinants. The nucleotide blast was performed using abricate v1.01 with a modified version of the CARD database and an 80% coverage and identity threshold. AMR determinants are grouped based on the antibiotic class they provide resistance against. B) Nucleotide alignment between the Tn*STC228* transposable element and the Tn*STC268* transposon found across the CC104-MDR subpopulation. AMR genes are coloured based on the antibiotic class they provide resistance against. Transposon genes are coloured yellow.

In addition, a subpopulation within the CC104 lineage has become multi-drug resistant (MDR) with some isolates carrying up to 11 AMR genes conferring resistance against 7 different antibiotic classes (Figure 4A). This sub-population (referred to as CC104-MDR throughout the rest of the manuscript) acquired a large transposable element (Tn*STC228)* carrying five aminoglycoside resistance genes, two lincosamide resistance genes and one nucleoside resistance gene (Figure 4B). However, some isolates have lost a large part of the element and only carry four of the aminoglycoside resistance genes and one nucleoside resistance gene (Figure 4B). Additionally, five CC104 isolates outside the main CC104-MDR subpopulation have acquired a prophage carrying an aminoglycoside resistance gene and two lincosamide resistance genes leading them also be resistant to 7 antibiotic classes (Supplementary Figure 5). Furthermore, a small plasmid carrying a homolog of the lipoglycopeptide resistance gene *vanZ* is carried by most of the CC104 isolates (Supplementary Figures 6-7).

### Thai zoonotic lineages CC104 and CC233 have increased β-lactam resistance through recombination in their penicillin binding proteins (PBP)

Thai zoonotic *S. suis* lineage CC233, along with Thai lineages CC221/234 and CC1688 had higher average MICs against penicillin and ceftriaxone than globally distributed lineages CC1, CC25, CC28, and CC94 (Figure 5 A-B). Furthermore, while the mean MIC of CC104 did not differ significantly from globally distributed CCs, isolates from the CC104-MDR subpopulation had a significantly higher MIC against penicillin and ceftriaxone than the rest of the CC104 isolates (Supplementary Figure 8). In streptococci, resistance against β-lactams is predominantly driven by amino acid changes in their penicillin binding proteins (PBPs). We combined a recently described multi-dimensional scaling (MDS) method with hierarchical clustering to categorise a representative selection of 78 Thai isolates into genetic clusters based on the amino acid variation in their PBPs, and compared the correlation between PBP types and β-lactam MICs (Supplementary Figure 9; Supplementary Figure 10).

**Figure 5.**
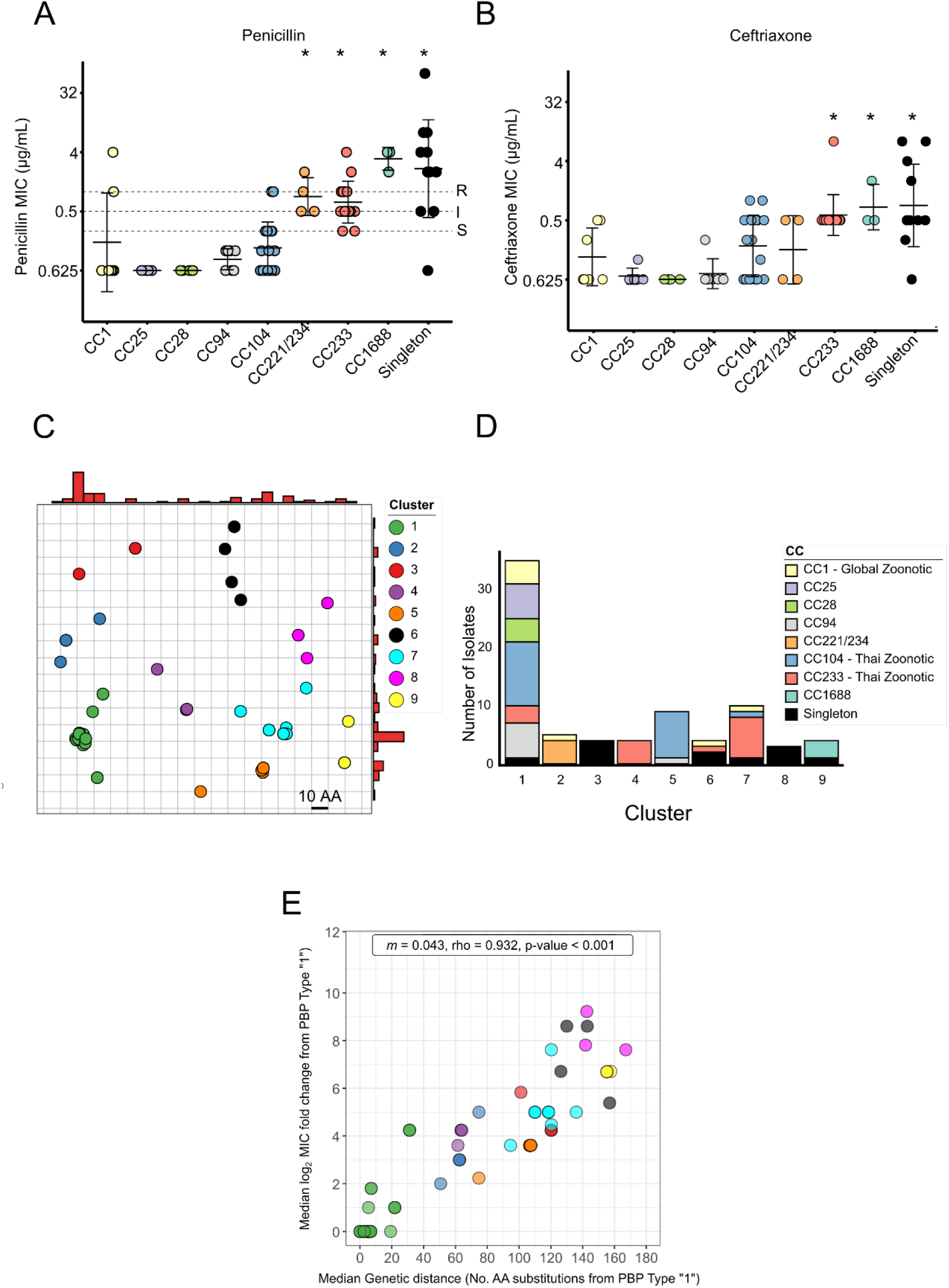
CC104 and CC233 have acquired resistant PBP types. A) and B) show the MICs for penicillin and ceftriaxone respectively. The y-axis was log_2_ transformed. Isolates are grouped by CC. Data were analysed using one-way ANOVA with Dunnett’s multiple comparisons, against CC1 (* indicates statistical significance: *p* <0.05). Horizontal lines on the penicillin plot indicate MIC breakpoints where S = susceptible, I= intermediate and R = resistant. For penicillin we used the recommended *S. suis* MIC breakpoints in CLSI Veterinary medicine (VET01-7^th^ edn). *S. suis* does not have established ceftriaxone MIC breakpoints. C) PBP genotype map of 78 representative Thai *S. suis* isolates. The alignment matrix was transformed into a two-dimensional space using multidimensional scaling (MDS). Gridlines represent 10 amino acid changes. Red histograms on the axis represent the relative distribution of isolates on the plot. Isolates were grouped into 9 clusters using hierarchical clustering algorithms based on their relative distances D) Clonal Complex distribution on each cluster from C. E) Spearman’s rank correlation between diverging from the susceptible PBP type “1” and increased β-lactam resistance. Rho = Spearman Correlation coefficient, *m =* slope. The median MIC value for both ceftriaxone and penicillin of each PBP type was used. PBP types were coloured based on their cluster. AA: Amino acid, CC: Clonal complex, rho: Spearman’s correlation coefficient, PBP: Penicillin binding protein.

We identified 9 clusters of PBP proteins carried by Thai *S. suis* isolates. Cluster 1 includes the majority of susceptible isolates and was composed of all the isolates from the zoonotic and pathogenic lineages CC1, CC25 CC28, CC94, and some isolates from CC104 and CC233 (Figure 5C-D; Supplementary Figure 11-13). There was a strong positive correlation (*m =* 0·043, rho = 0·932, and p<0·001) between divergence from PBP type 1 (the most common PBP type from Cluster 1) and increased MIC against penicillin and ceftriaxone (Figure 5E). All isolates from the main CC104-MDR subpopulation were in Cluster 5, which displayed increased MICs for both penicillin (mean Δlog2 = 1·5) and ceftriaxone (mean Δlog2 = 2·4) when compared to Cluster 1 (Figure 5D; Supplementary Figure 12-13). CC233 isolates had a diverse set of PBP types and were spread across clusters 1, 4, 6, and 7 (Figure 5D). Isolates from Clusters 4 (CC233), 6 (CC1, CC233, and Singletons) and 7 (CC1, CC104, CC233, and Singletons) had increased MICs against ceftriaxone (mean Δlog2 = 2·8) and penicillin (mean Δlog2 = 3·77) compared to Cluster 1 (Supplementary Figure 12-13). Acquisition of non-susceptible PBP types in the zoonotic lineages CC104 and CC233 appears to be recombination driven, with all three PBP genes displaying evidence of multiple recombination events (Supplementary Figure 14).

## Discussion

The emergence of the two novel zoonotic *S. suis* lineages CC104 and CC233 coincided with a switch from a serotype 7 capsule to the serotype 2 capsule acquired from the dominant global zoonotic clade. Their evolution is also characterised by the rapid acquisition of multiple AMR determinants, through horizontal gene transfer. Both lineages gained increased resistance against β-lactams through recombination within their Penicillin Binding Protein (PBPs) genes. Additionally, a recent CC104 subpopulation (CC104-MDR) acquired multiple mobile genetic elements carrying a large number of AMR determinants, resulting in multi-drug resistance. The recent emergence of these novel multi-drug resistant zoonotic lineages not only indicates that human infections are not restricted to isolates belonging to zoonotic lineage CC1 as previously suggested but also that novel zoonotic lineages will continue to emerge.^20^

While a recent study found that the main globally distributed pathogenic *S. suis* lineages emerged between the beginning of the 19^th^ century and the mid-20^th^ century,^9,21^ we estimate that CC104 and CC233 emerged much more recently in the late 20^th^ and early 21^st^ century. These dates follow the intensification of pig farming in Thailand which started in the 1960s after the import of commercial pig breeds from the United Kingdom and the United States.^22^ It is possible that these imports also led to the introduction of pathogenic lineages of *S. suis,* including CC1. As CC104 and CC233 have not been identified outside of Thailand it is likely that these lineages emerged within Thailand and have not yet spread to other countries. Nevertheless, as whole genome sequencing of *S. suis* isolates is uneven across different countries, it is possible that they have spread to other countries but have yet to be detected. In particular, countries importing live pigs from Thailand (Myanmar, Laos, Cambodia, and Vietnam) are at risk of acquiring CC104 and CC233.

The emergence of CC104 and CC233 as zoonotic agents is marked by the acquisition of a serotype 2 capsule, a common feature of zoonotic *S. suis* strains. For instance, the Dutch zoonotic lineage CC20 diverged from the European porcine pathogenic CC16 lineage following a switch in its capsule from serotype 9 to serotype 2.^21^ Additionally, the few sporadic zoonotic infections by CC87 and CC94 strains have been caused by divergent and rare serotype 2 strains.^3,9^ However, a serotype 2 capsule on its own is unlikely to be sufficient for *S. suis* to cause zoonotic infections and the genetic background is likely an additional determinant, since despite carrying a serotype 2 capsule, CC25 and CC28 isolates rarely cause zoonotic infections.^23^ The zoonotic lineage CC1 seems to act as a reservoir of serotype 2 capsular locus globally, transferring its capsule locus to other pathogenic lineages through horizontal gene transfer.^9^ The capsule switching events appear to be recent in the case of CC104, CC233, and CC1688, as their serotype 2 locus differ only by 1 SNP from the serotype 2 capsule in CC1 isolates. However, the precise mechanisms by which serotype 2 capsular polysaccharide facilitates disease in both pigs and humans remains unknown.

Thailand is amongst countries with the highest consumption of antibiotics per kilogram of meat in the world, with over 1000 tonnes of active pharmaceutical ingredients consumed in the pig farming industry per year.^24,25^ Amoxicillin is by far the most used antibiotic in porcine medicated feed in Thailand and shares a mechanism of action with penicillin and ceftriaxone (i.e. binding PBPs).^25^ Therefore, the high usage of amoxicillin is likely to have exerted a strong evolutionary selective pressure on Thai *S. suis* isolates, resulting in the co-selection of PBP types that also confer increased resistance against penicillin and ceftriaxone. The selection for resistant PBP types has led to a moderate increase in resistance to ceftriaxone, despite cephalosporins not being used in pig farming. Apart from amoxicillin, pleuromutilins, quinolones, macrolides, tetracyclines and aminoglycosides are also extensively used in pig farming in Thailand.^26^ The high usage of pleuromutilins may explain why lineages endemic to Thailand (CC104, CC233, and CC1688) carry the full-length version of *vgaF*, associated with increased pleuromutilin resistance. Additionally, the high antibiotic use will have facilitated the spread of the Tn*STC228* transposon carried by the CC104-MDR subpopulation, thus co-selecting for multiple AMR determinants. Taken together, high antimicrobial usage could have provided a favourable setting for lineages CC104 and CC233 to recently emerge as dominant pathogenic strains and potentially outcompete the CC1 lineage, especially considering that the CC1 isolates carry fewer AMR determinants, including susceptible PBP types. The evolution of AMR in CC104 and CC233 may have been facilitated by horizontal transfer from and recombination with commensal isolates of *S. suis* as they carry more AMR determinants on average than pathogenic isolates as well resistant PBP types (Supplementary Figure 15).

The evolution of CC104 and CC233 shows how horizontal gene transfers can lead to the emergence and spread of novel multi-drug resistant (MDR) zoonotic *S. suis* lineages. It is of special concern that these novel zoonotic *S. suis* lineages are not only MDR but also are accumulating resistance to critically important antibiotics such as ceftriaxone. For now, the human host remains an evolutionary dead end for *S. suis* as there have been no reports of human-to-human transmission and very little evidence for human carriage.^2^ Nonetheless, novel zoonotic and MDR strains may also be emerging in other countries that have recently undergone pig farming intensification and have a high level of antibiotic usage. This could result in not only an increased burden of disease in such countries but also in other regions via international trade of live animals. Therefore, while *S. suis* CC1 infections remain treatable with β-lactams, we predict that we will see an increasing burden of MDR human *S. suis* infections.

## Contributors

JB: Data Curation, Formal analysis, Investigation, Methodology, Validation, Visualization, Writing – original draft. GGRM: Methodology, Supervision, Validation, Writing -review and editing. PB: Data Curation, Investigation, Methodology, Writing – review and editing. AJB: Methodology, Supervision, Validation, Writing -review and editing. PW: Investigation, Methodology Resources, Writing – review and editing. PJ: Investigation, Methodology, Resources, Writing – review and editing. PC: Investigation, Methodology, Writing -review and editing RH: Investigation, Methodology, Writing – review and editing. LAW: Methodology, Software, Resources, Supervision, Validation, Writing – review and editing. AK: Conceptualization, Data Curation, Funding acquisition, Project Administration, Resources, Supervision, Writing – review and editing. CS: Conceptualization, Funding acquisition, Methodology, Project administration, Software, Resources, Supervision, Writing – review and editing.

## Ethical Statement

The medical records of 1015 human cases of *S. suis* infection were reviewed at local hospitals in Thailand by attending physicians as part of previous published studies. The clinical case record form was approved by the Ethics Committee of the Department of Medical Sciences, Ministry of Public Health, Thailand for all the previous studies from where this data derives.

## Conflicts of Interest

We declare no conflict of interests.

## Acknowledgements

We would like to thank Boas C. L. van der Putten from the Thomas J. Roodsant from the National Institute for Public Health and the Environment (RIVM) for their support and insights and valuable discussions.

This study was funded through the CANVAS research project (grant number: LSHM19137), a public-private partnership powered by Health∼Holland, Top Sector Life Sciences & Health, a research and innovation funding program of the Dutch government; by the ZoNMW through the NCOH Pandemic Preparedness Kickstarter (ZonMW Grant #10710022210003) project; and through Kasetsart University Research and Development Institute (KURDI), Bangkok, Thailand (grant number: FF(KU)26.67).

## Data Sharing

Raw Illumina and Nanopore FastQ reads have been deposited in the NCBI Short Read Archive under BioProject PRJNA1210814. The draft genome assemblies, annotated genomes and BioSample metadata can also be found under the same BioProject. The four complete genomes and their annotations generated in this study are deposited in BioProject PRJNA1221088.

## Notes

### Competing Interest Statement

The authors have declared no competing interest.

## Bibliography

1 Segura M, Aragon V, Brockmeier SL, et al. Update on *Streptococcus suis* Research and Prevention in the Era of Antimicrobial Restriction: 4th International Workshop on S. suis. Pathogens 2020; 9: 374.

2 Ho DTN, Le TPT, Wolbers M, et al. Risk Factors of *Streptococcus suis* Infection in Vietnam. A Case-Control Study. PLoS ONE 2011; 6: e17604.

3 Brizuela J, Roodsant TJ, Hasnoe Q, et al. Molecular Epidemiology of Underreported Emerging Zoonotic Pathogen *Streptococcus suis* in Europe. Emerg Infect Dis 2024; 30: 413–22.

4 Huong VTL, Ha N, Huy NT, et al. Epidemiology, Clinical Manifestations, and Outcomes of *Streptococcus suis* Infection in Humans. Emerg Infect Dis 2014; 20: 1105–14.

5 Yu H, Jing H, Chen Z, et al. Human *Streptococcus suis* Outbreak, Sichuan, China. Emerg Infect Dis 2006; 12: 914–20.

6 Brizuela J, Kajeekul R, Roodsant TJ, et al. *Streptococcus suis* outbreak caused by an emerging zoonotic strain with acquired multi-drug resistance in Thailand. Microb Genomics 2023; 9.

7 Tarini NMA, Susilawathi NM, Sudewi AAR, et al. A large cluster of human infections of *Streptococcus suis* in Bali, Indonesia. One Health 2022; 14: 100394.

8 King SJ, Leigh JA, Heath PJ, et al. Development of a multilocus sequence typing scheme for the pig pathogen *Streptococcus suis*: identification of virulent clones and potential capsular serotype exchange. J Clin Microbiol 2002; 40: 3671–80.

9 Murray GGR, Hossain ASMdM, Miller EL, et al. The emergence and diversification of a zoonotic pathogen from within the microbiota of intensively farmed pigs. Proc Natl Acad Sci 2023; 120: e2307773120.

10 Kerdsin A. Human *Streptococcus suis* Infections in Thailand: Epidemiology, Clinical Features, Genotypes, and Susceptibility. Trop Med Infect Dis 2022; 7: 359.

11 Athey TBT, Teatero S, Lacouture S, Takamatsu D, Gottschalk M, Fittipaldi N. Determining *Streptococcus suis* serotype from short-read whole-genome sequencing data. BMC Microbiol 2016; 16: 162.

12 Croucher NJ, Page AJ, Connor TR, et al. Rapid phylogenetic analysis of large samples of recombinant bacterial whole genome sequences using Gubbins. Nucleic Acids Res 2015; 43: e15.

13 Suchard MA, Lemey P, Baele G, Ayres DL, Drummond AJ, Rambaut A. Bayesian phylogenetic and phylodynamic data integration using BEAST 1.10. Virus Evol 2018; 4: vey016.

14 Hadjirin NF, Miller EL, Murray GGR, et al. Large-scale genomic analysis of antimicrobial resistance in the zoonotic pathogen *Streptococcus suis*. BMC Biol 2021; 19: 191.

15 Katoh K, Misawa K, Kuma K, Miyata T. MAFFT: a novel method for rapid multiple sequence alignment based on fast Fourier transform. Nucleic Acids Res 2002; 30: 3059.

16 Price MN, Dehal PS, Arkin AP. FastTree 2 – Approximately Maximum-Likelihood Trees for Large Alignments. PLoS ONE 2010; 5: e9490.

17 Takamatsu D, Wongsawan K, Osaki M, et al. *Streptococcus suis* in Humans, Thailand. Emerg Infect Dis 2008; 14: 181–3.

18 Roodsant TJ, Van Der Putten BCL, Tamminga SM, Schultsz C, Van Der Ark KCH. Identification of *Streptococcus suis* putative zoonotic virulence factors: A systematic review and genomic meta-analysis. Virulence 2021; 12: 2787–97.

19 Lekagul A, Tangcharoensathien V, Mills A, Rushton J, Yeung S. How antibiotics are used in pig farming: a mixed-methods study of pig farmers, feed mills and veterinarians in Thailand. BMJ Glob Health 2020; 5: e001918.

20 Dong X, Chao Y, Zhou Y, et al. The global emergence of a novel *Streptococcus suis* clade associated with human infections. EMBO Mol Med 2021; 13: e13810.

21 Willemse N, Howell KJ, Weinert LA, et al. An emerging zoonotic clone in the Netherlands provides clues to virulence and zoonotic potential of *Streptococcus suis*. Sci Rep 2016; 6: 28984.

22 Thanapongtharm W, Linard C, Chinson P, et al. Spatial analysis and characteristics of pig farming in Thailand. BMC Vet Res 2016; 12: 218.

23 Goyette-Desjardins G, Auger J-P, Xu J, Segura M, Gottschalk M. *Streptococcus suis*, an important pig pathogen and emerging zoonotic agent—an update on the worldwide distribution based on serotyping and sequence typing. Emerg Microbes Infect 2014; 3: 1–20.

24 Mulchandani R, Wang Y, Gilbert M, Boeckel TPV. Global trends in antimicrobial use in food-producing animals: 2020 to 2030. PLOS Glob Public Health 2023; 3: e0001305.

25 Health Policy and Systems Research on Antimicrobial Resistance (HPSR-AMR) Network. Highlights Thailand One Health Report 2021: Antimicrobial Consumption and Antimicrobial Resistance. 2024.

26 Coyne L, Arief R, Benigno C, et al. Characterizing Antimicrobial Use in the Livestock Sector in Three South East Asian Countries (Indonesia, Thailand, and Vietnam). Antibiotics 2019; 8: 33.

