## Supplementary Materials for "Emergence of zoonotic and multi-drug resistant *Streptococcus suis*"

### Supplementary Figures

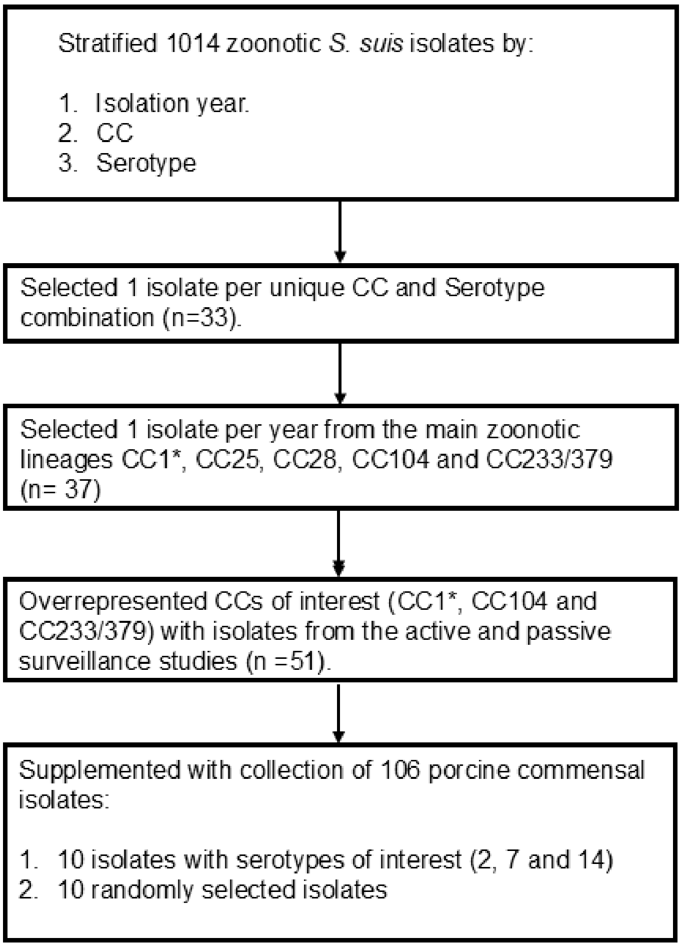

*An equal number of serotype2 and serotype 14 isolates were selected for CC1

**Supplementary Figure 1**. Isolate selection criteria for whole genome sequencing. We only included the index case from isolates belonging to single outbreaks to avoid sequencing duplicated strains.

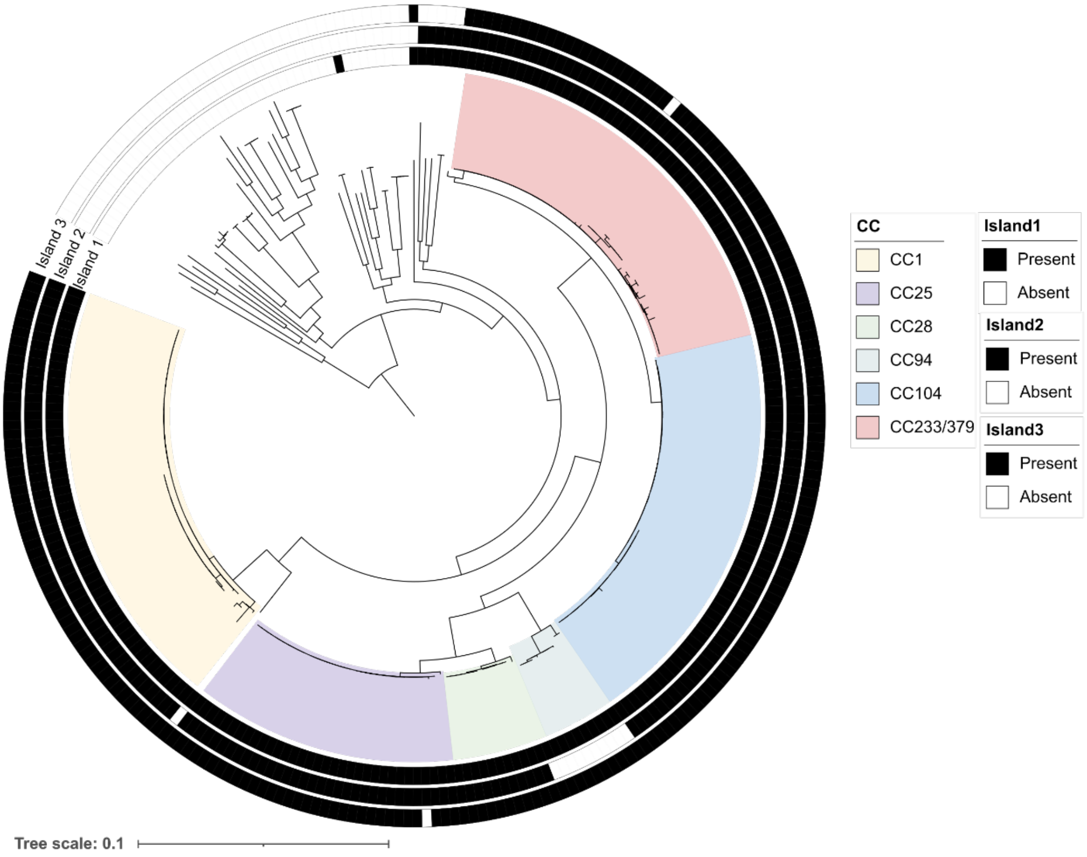

**Supplementary Figure 2**. Presence of the 3 genomic islands characteristic of *S. suis* pathogenic lineages on Thai *S. suis* isolates. Tree scale is in nucleotide substitutions per site. Coloured blocks indicate CC. Rings indicate presence/absence of each genomic island.

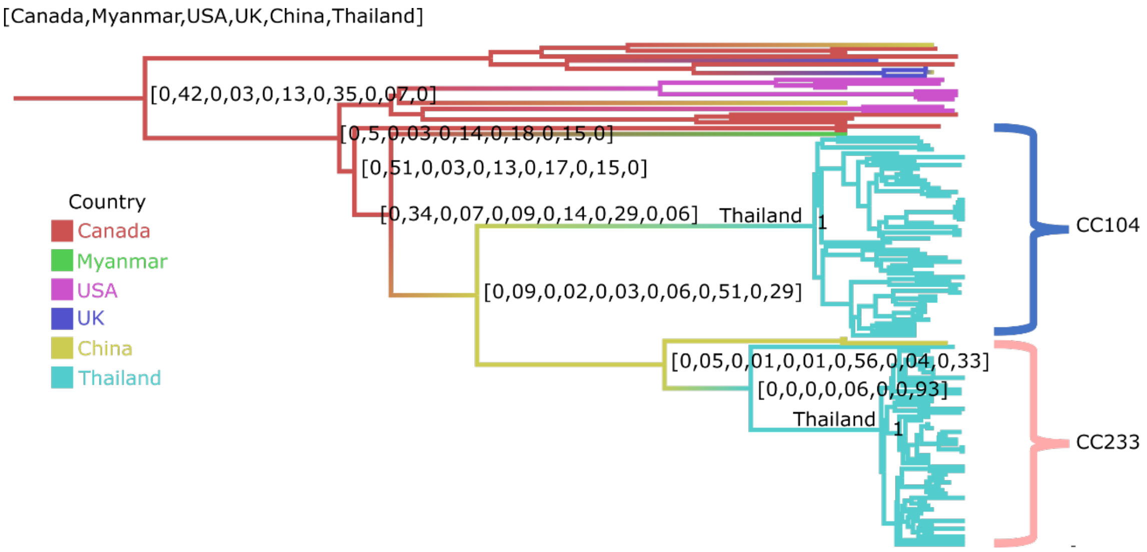

**Supplementary Figure 3**. Dated phylogeny of the Thai zoonotic lineages CC104 and CC233/379. The dated phylogeny was reconstructed using BEAST with a strict molecular clock, GTR+G substitution model and constant population model. The colour of the branches represents the most likely ancestral location derived from an asymmetric discrete trait model. The country-of-origin posterior probabilities for nodes from which CC104 and CC233 originate are indicated in brackets. The order of the posterior values associated with each country is shown at the top left.

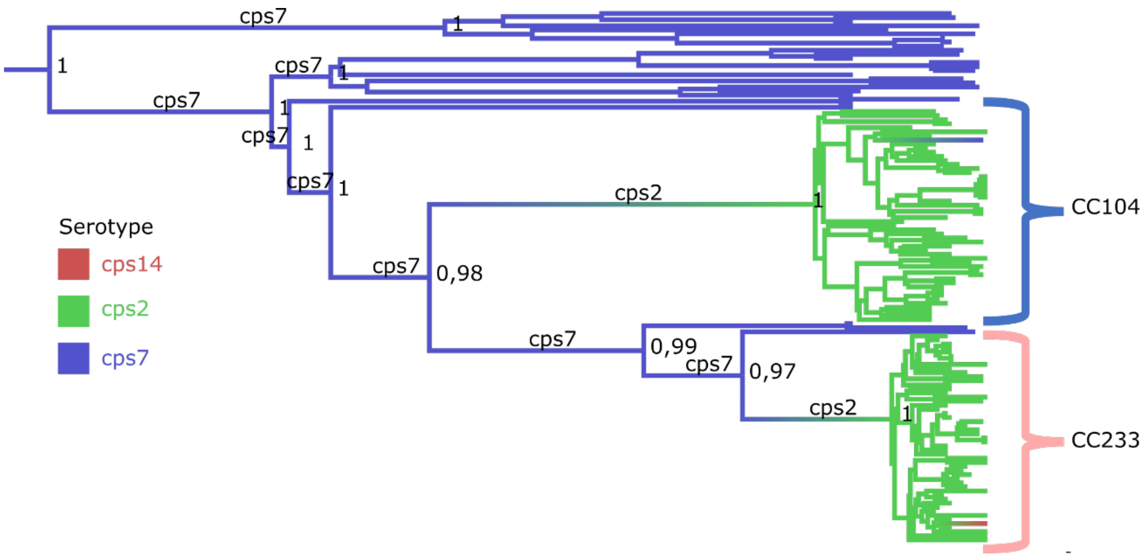

**Supplementary Figure 4**. Ancestral reconstruction of serotype using a symmetric discrete trait model. The colour of the branches represents the most likely ancestral serotype and the posterior probability for the ancestral state reconstruction on the main nodes are indicated.

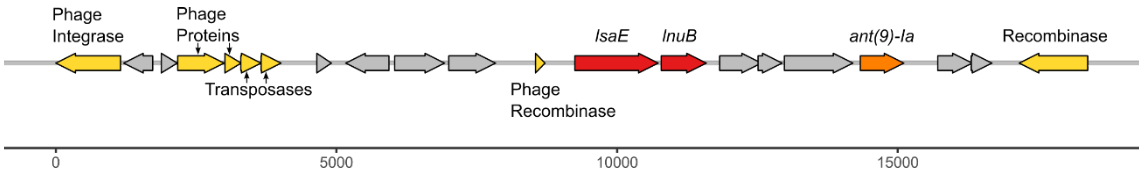

**Supplementary Figure 5**. Gene arrow map of prophage ΦSTC290 carrying AMR determinants. AMR genes are coloured based on the antibiotic class they confer resistance to as in Figure 4. Prophage genes are coloured in yellow.

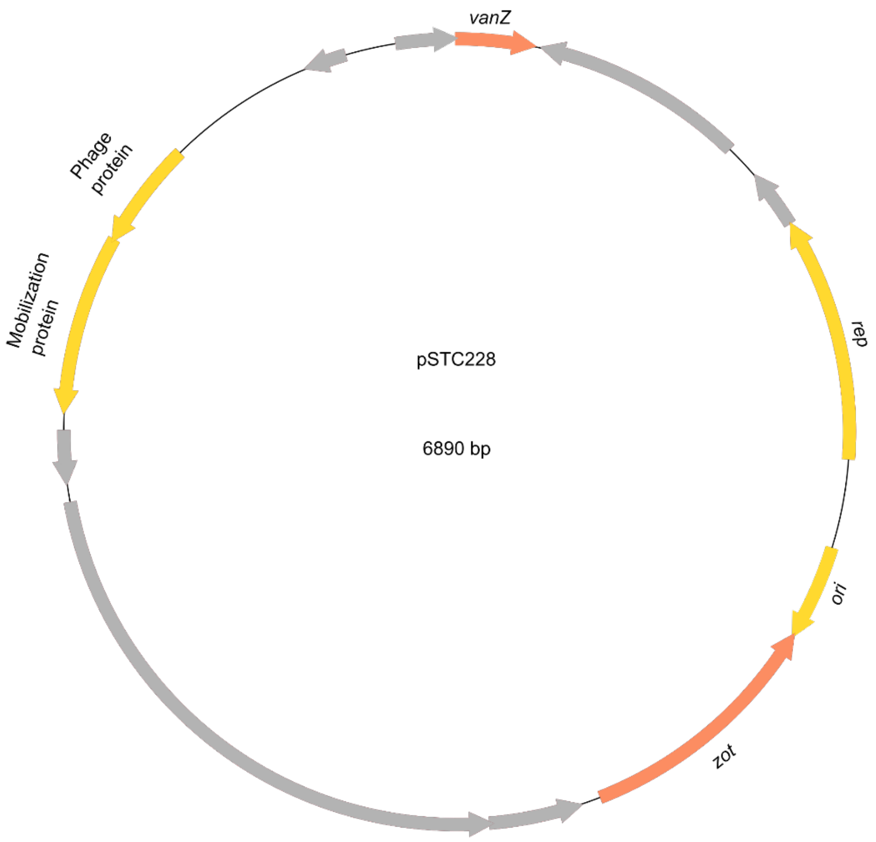

**Supplementary Figure 6**. Gene arrow map of the plasmid pSTC228 carrying vanZ which has spread across the zoonotic lineage CC104.

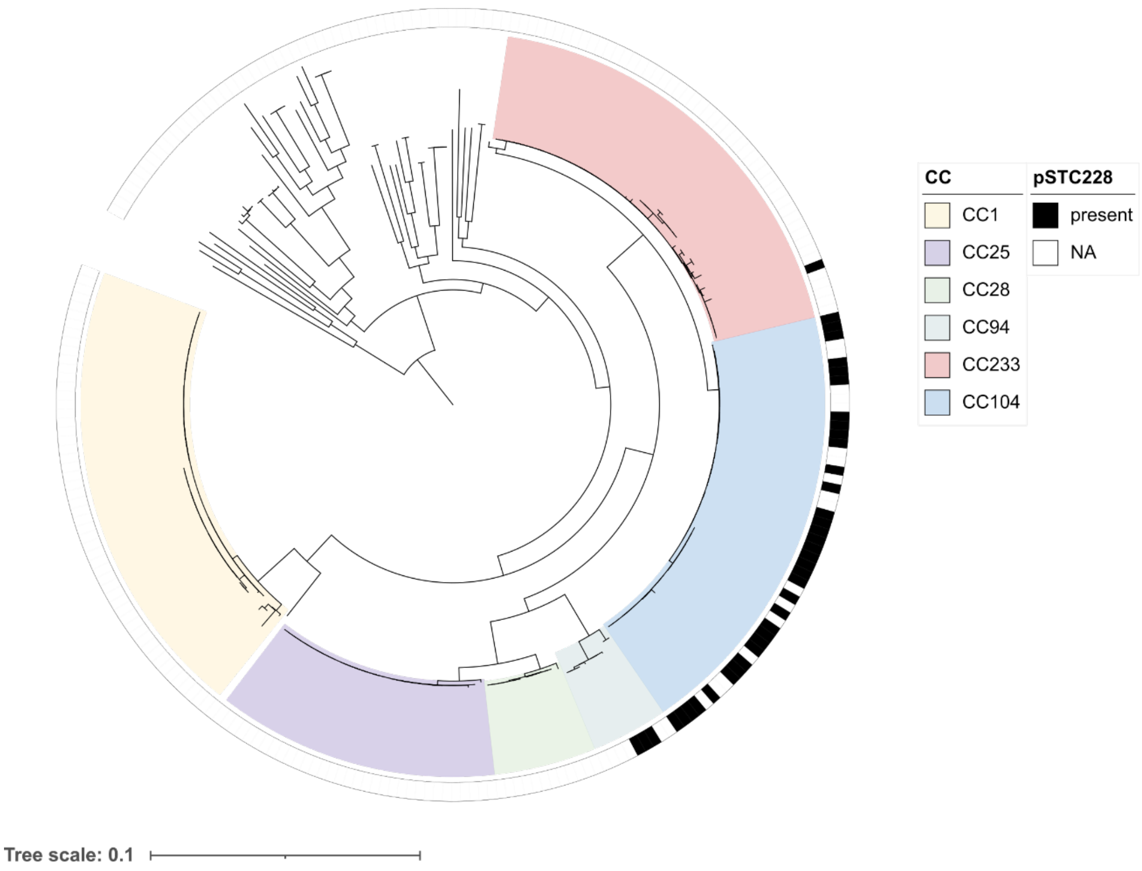

**Supplementary Figure 7**. Presence of the pSTC228 plasmid in the Thai *S. suis* genomes. The core genome phylogeny of the 264 Thai *S. suis* isolates shown in Figure 1A was used for this figure. Coloured blocks indicate CC. Presence of the pSTC228 plasmid is indicated in the outer coloured ring in black. Tree scale is in substitutions per site.

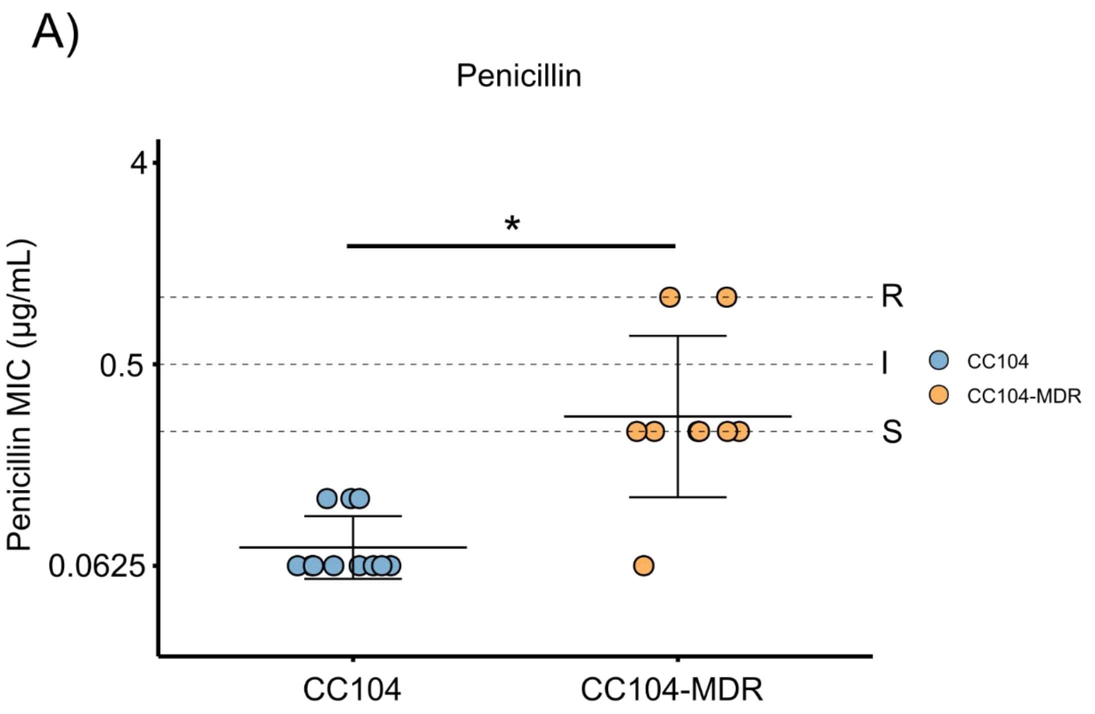

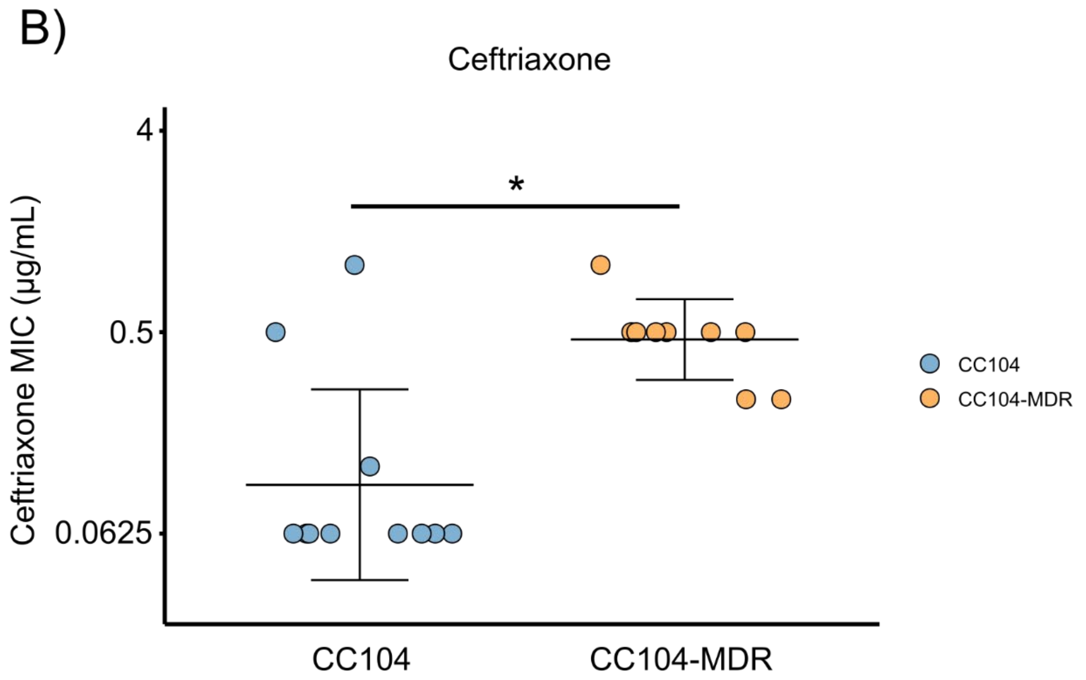

**Supplementary Figure 8**. Penicillin A) and Ceftriaxone B) MICs within CC104. The y-axis was log2 transformed. Samples are grouped as CC104 or CC104-MDR. A two-tailed t-test was performed (* = p<0.05). Horizontal lines on the Penicillin plot indicate MIC breakpoints where S = susceptible, I= intermediate and R = resistant. For penicillin we used the recommended S. suis MIC breakpoints in CLSI Veterinary medicine (VET01-7th edn). S. suis does not have established ceftriaxone MIC breakpoints.

A)

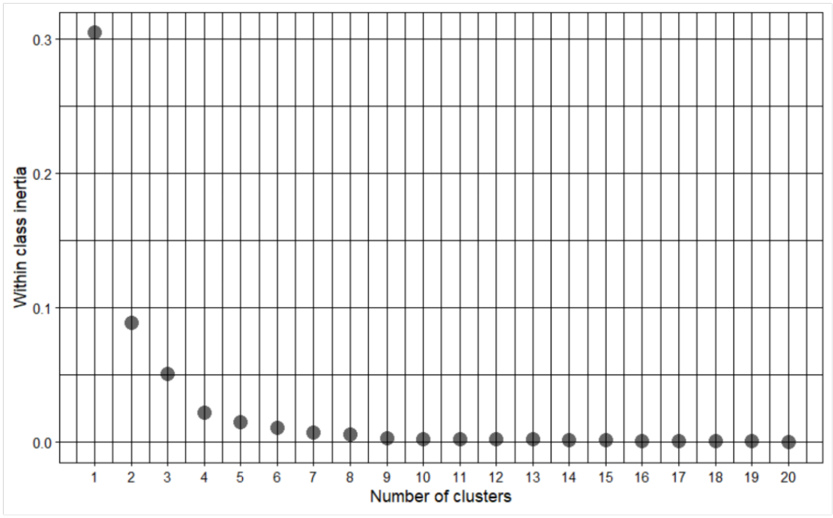

B)

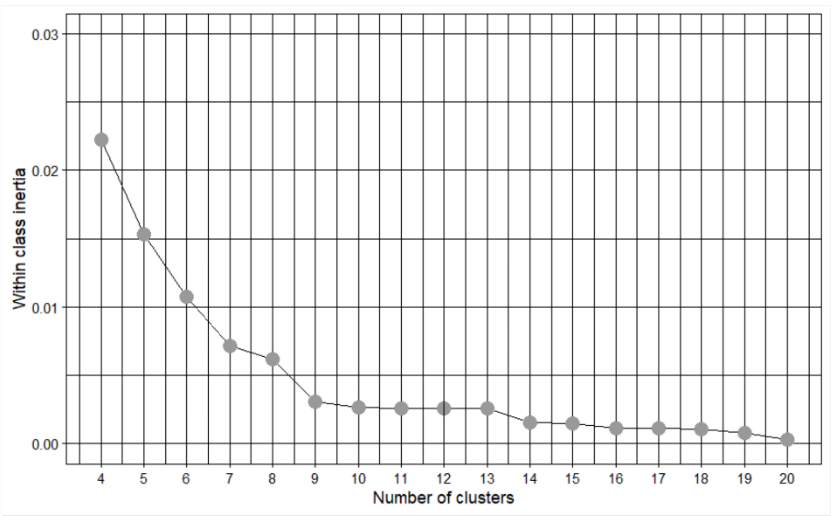

**Supplementary Figure 9**. Scree plot of the decrease in within cluster inertia as the number of clusters increases. A) 1-20 clusters. B) Zoomed in to 4-20 clusters.

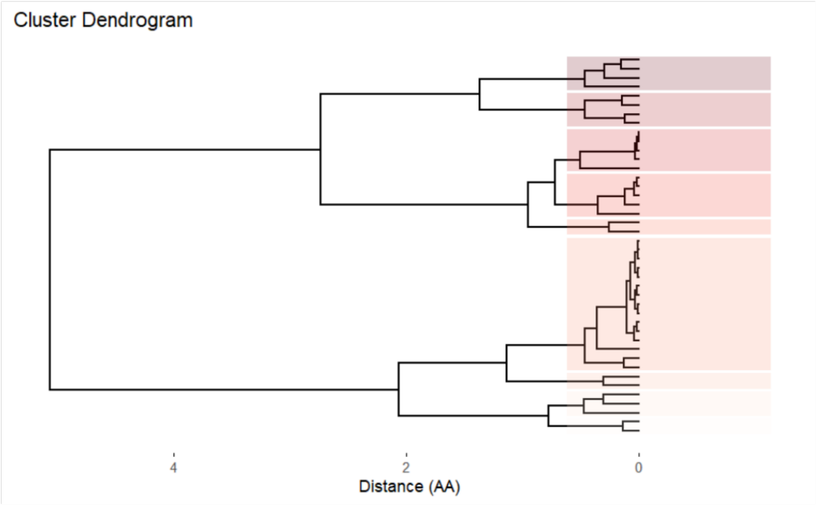

**Supplementary Figure 10**. Dendrogram of the different PBP alleles in Thai S. suis genomes and the distance between them. PBP alleles were clustered into 9 distinct clusters based on their pairwise distances using hierarchical clustering algorithms.

A)

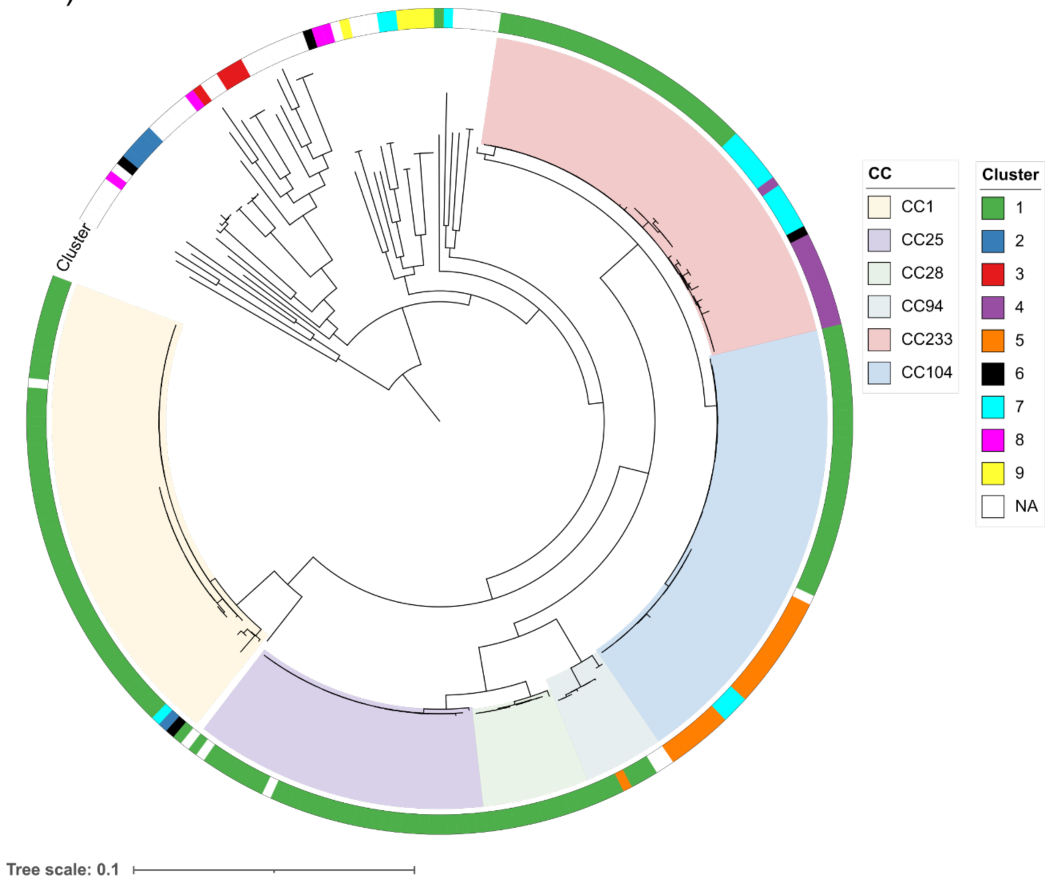

B)

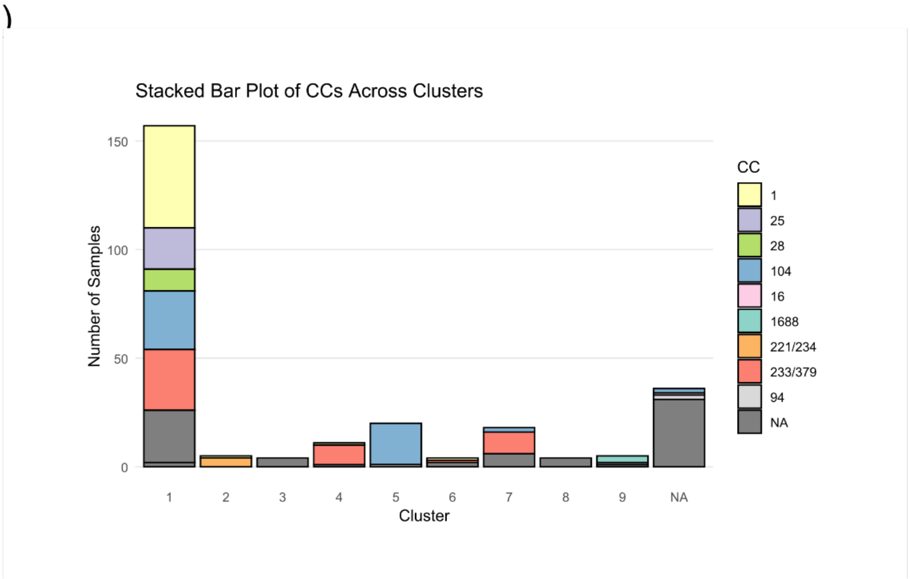

**Supplementary Figure 11**. A) Core genome S. suis phylogeny shown in Figure 1 overlayed with PBP allele cluster metadata from Figure 3. Coloured blocks indicate CC. Coloured ring indicates PBP Cluster. The tree scale is in substitutions per site. B) Distribution of CCs within each PBP cluster. NA = Not available.

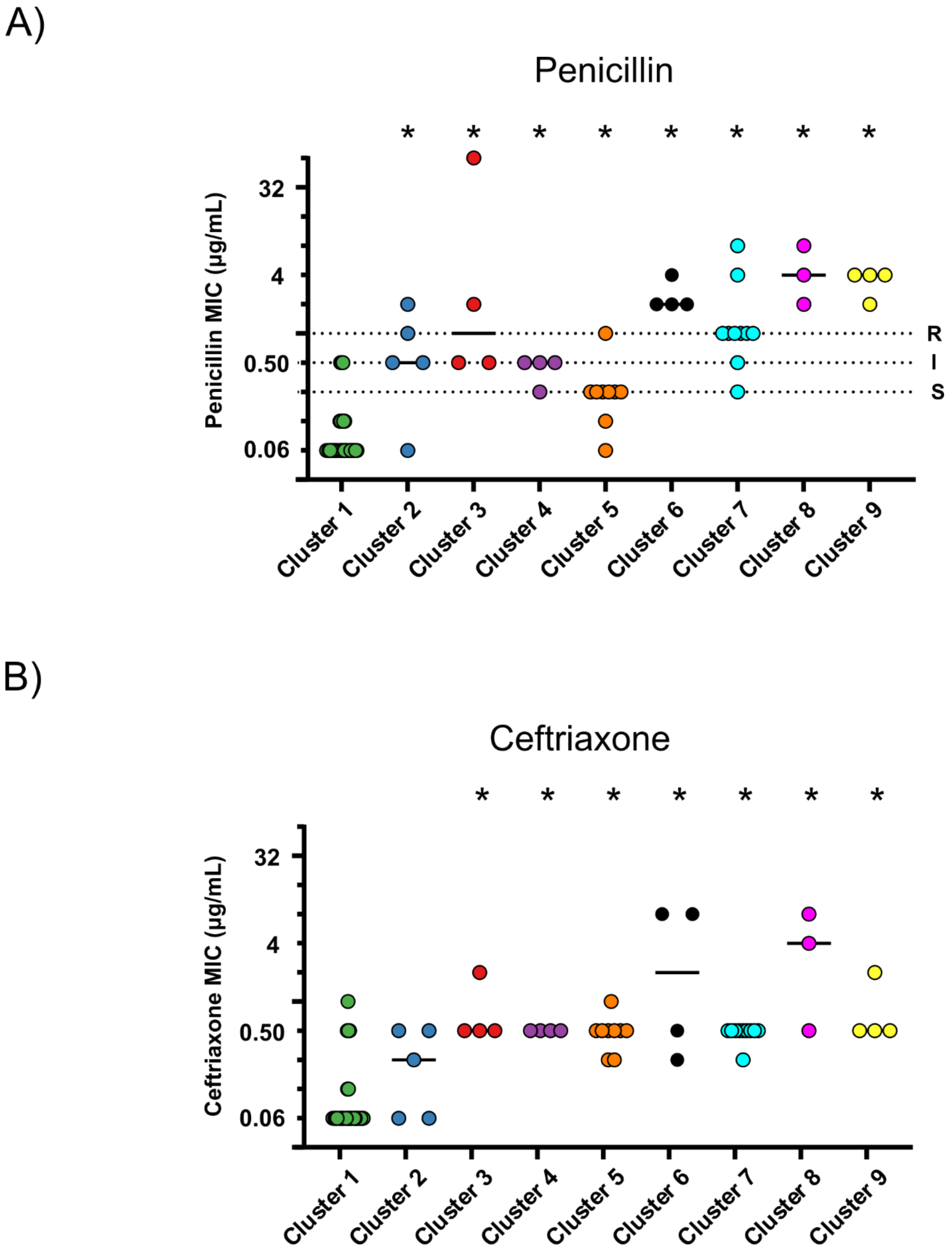

**Supplementary Figure 12**. A) and B) show the MICs for penicillin and ceftriaxone respectively. The y-axis was log2 transformed. Isolates are grouped by PBP Cluster. Data were analysed using one-way ANOVA with Dunnett's multiple comparisons, against CC1 (* indicates statistical significance: p <0.05). Horizontal lines on the penicillin plot indicate MIC breakpoints where S = susceptible, I= intermediate and R = resistant. For penicillin we used the recommended S. suis MIC breakpoints in CLSI Veterinary medicine (VET01-7th edn). S. suis does not have established ceftriaxone MIC breakpoints.

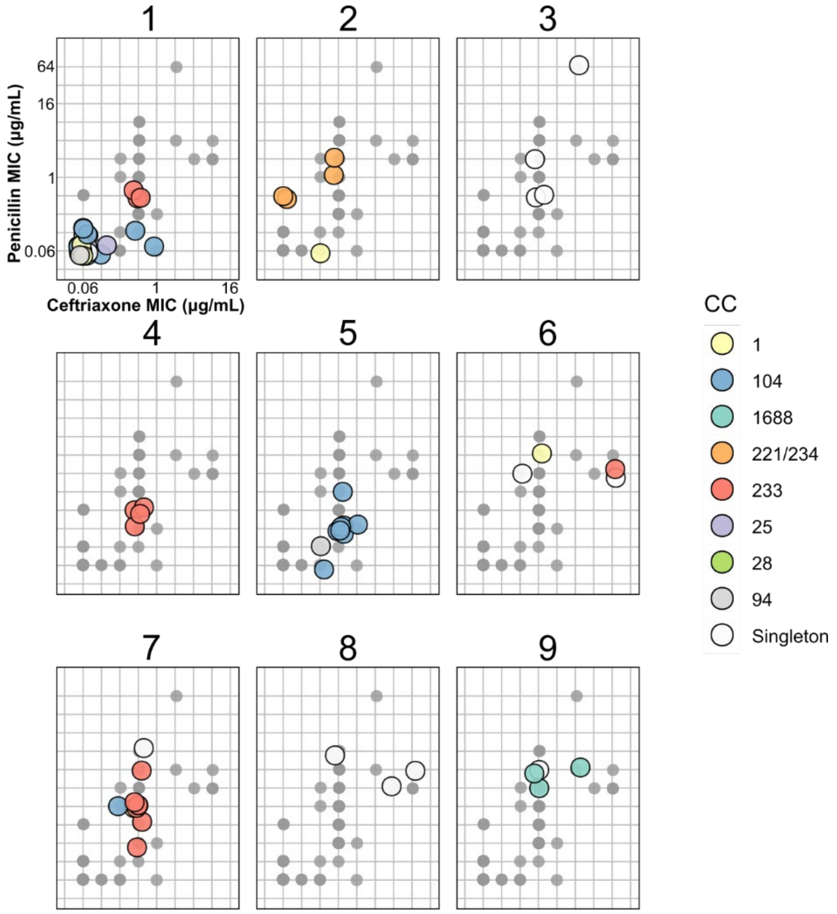

**Supplementary Figure 13**. Ceftriaxone and Penicillin MIC scatter plots for each of the 9 clusters. The ceftriaxone MIC values are displayed on the x-axis and the penicillin MIC values on the y-axis. The y-axis was log2 transformed. Clusters are labelled at the top of each panel and samples are coloured by CC. A light scattering effect was applied to each data point to visualize the density of isolates.

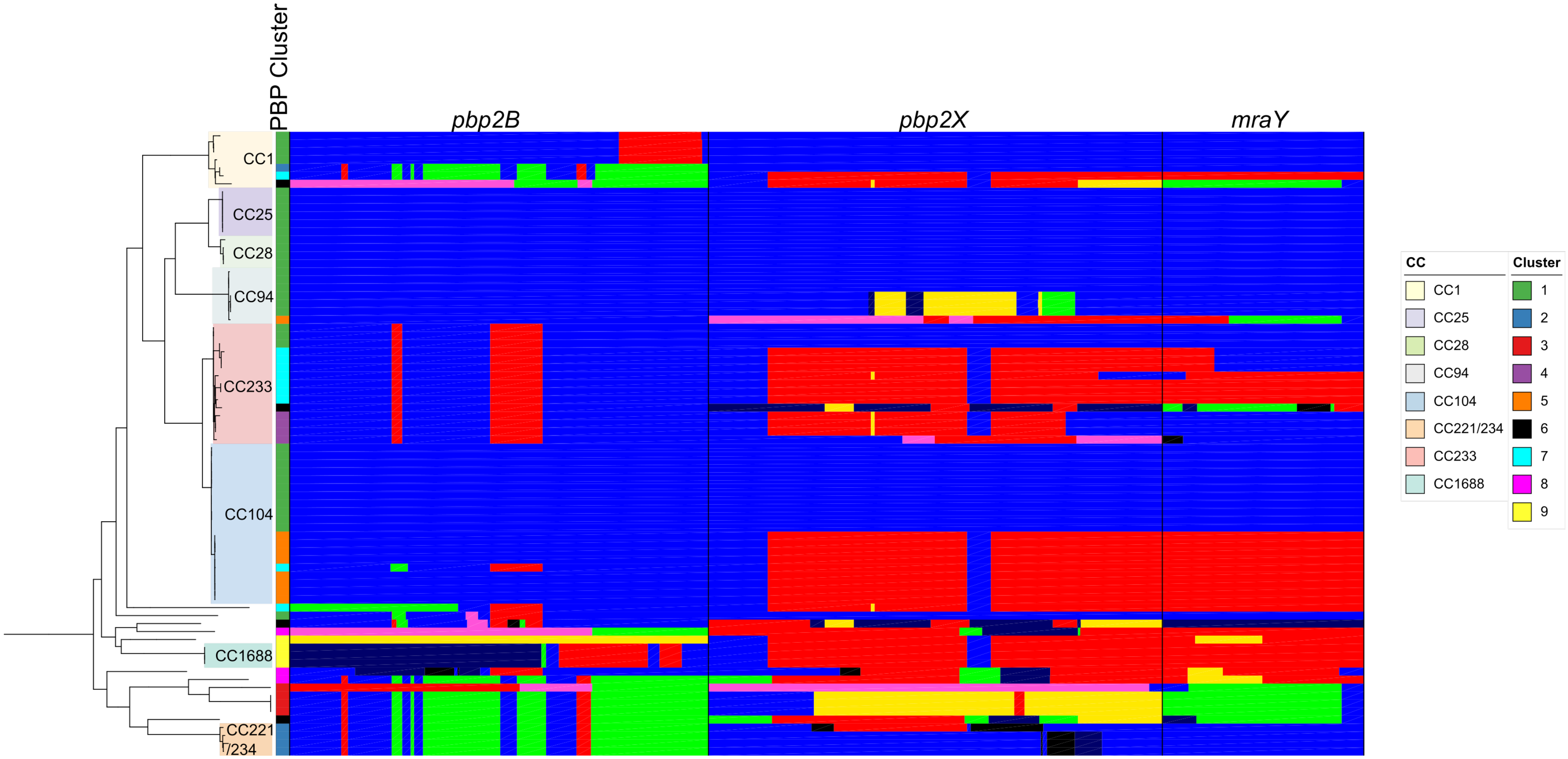

**Supplementary Figure 14**. Recombination of PBPs in 78 representative Thai S. suis isolates. The left side of the figure shows a core-genome phylogeny of the 78 Thai isolates. Coloured clades indicate CC. 1st Coloured Column indicates PBP type cluster. The coloured mosaic blocks next to the tree display the recombination results for each PBP performed with fastGEAR. Different colours indicate different PBP lineages as identified by fastGEAR. mraY is directly downstream of pbp2X in S. suis' genomes.

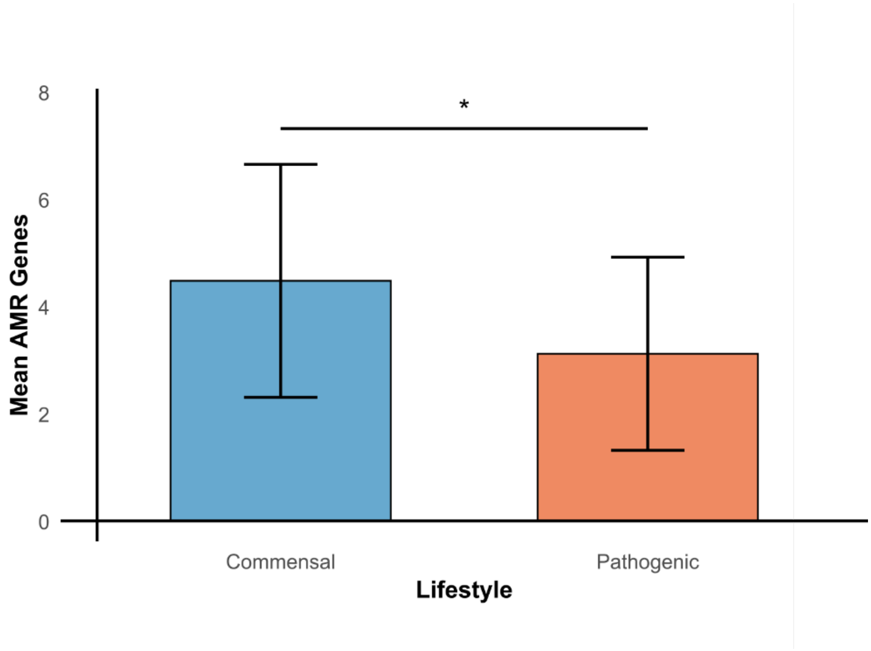

**Supplementary Figure 15**. Mean number of AMR genes carried by S. suis isolates by ecological niche. A two-tailed t-test was performed (* = p<0.05).

### Supplementary Methods

#### Stratified and randomized selection of *S. suis* isolates for whole genome sequencing

We stratified all zoonotic isolates (n= 1014) by isolation year, clonal complex (CC), and serotype. We first selected one isolate from each unique ST and serotype combination (n = 33). Subsequently, we selected one isolate per year from the main zoonotic lineages (CC1, CC25, CC28, CC104 and CC233/379; n=37). We supplemented CCs of interest (CC1, CC104 and CC233/379) with additional isolates, by selecting an equal number of isolates from both the active (Supplementary Table 1) and passive (Supplementary Table 2) surveillance studies (n = 51). In the case of CC1 we selected an equal number of serotype 2 and serotype 14 isolates.

#### Collection of public thai *S. suis* genomes

We scanned the literature for publications describing WGS on Thai *S. suis* isolates [1–7], NCBI's *S. suis* Biosamples collected in Thailand and with associated WGS data, as well as pubMLST's (https://pubmlst.org) genome database. Where possible, assemblies were extracted. When only short read sequencing data was available, we assembled short reads using Shovill (v1.09; https://github.com/tseemann/shovill) with SPAdes (v3.14.1) [8]. We manually removed duplicates cross-referencing sample ids and metadata.

#### Illumina whole genome sequencing and assembly

Pure cultures from the 141 selected isolates were grown in Todd-Hewitt Broth supplemented with 0.5% yeast extract (THY) at 37 $^{0}$C until sufficient bacterial mass was obtained. The cultures were pelleted and resuspended in DNA/RNA Shield buffer (Zymo Research, USA) and sent to MicrobesNG (https://microbesng.com/) for genomic DNA extraction and Illumina whole genome sequencing. The Nextera XT Library Prep Kit (Illumina, San Diego, USA) was used for preparing the genomic library and samples were sequenced with 250 bp pair-end reads using an Illumina NovaSeq 6000 (Illumina, San Diego, USA).

The raw Illumina reads were filtered and cleared of adapters at MicrobesNG (https://microbesng.com/) using a combination of Trimmomatic [9] and in-house scripts. We assembled the draft genomes Shovill (v1.09; https://github.com/tseemann/shovill) with SPAdes v3.14.1 [8] and assessed the quality of the draft assembly with QUAST v 4.6.3 [10]. We used Prokka v1.14 to annotate the draft assemblies [11].

#### Nanopore whole genome sequencing and assembly

Library preparation for Oxford Nanopore Technologies (ONT) sequencing followed the rapid barcoding DNA sequencing protocol with the SQK-RBK004 kit without DNA size selection (preserve the plasmid DNA) and the libraries were sequenced using a single R10.4.1 flow cell on a GridION sequencer. We base called and demultiplexed the raw data using Guppy v3.4.5 (ONT) specifying the high-accuracy model (-c dna_r9.4.1_450bps_hac.cfg). The ONT adapters were trimmed using Porechop v0.2.4 (https://github.com/rrwick/Porechop). Quality control of ONT reads was undertaken using Nanoplot v1.28.1. Hybrid assemblies with the ONT and Illumina data were performed using Unicycler v0.4.8 [12], and the genome sequences were checked for quality using QUAST v5.0.2 [10]. Draft and complete genomes were annotated using Prokka v1.14.6 [11].

#### Selection of isolates for minimum inhibitory concentration testing

We selected at least one sample from each available PBP allele (n=41) and a total of 3 samples from alleles that had more than two sample available (n=37) to perform minimum inhibitory concentration (MIC) testing with penicillin and ceftriaxone following the CLSI guidelines (Supplementary Table 6).

#### Core-genome phylogenies

To investigate the Thai *S. suis* population structure we reconstructed a core-genome single nucleotide polymorphism (SNP) phylogeny using all 264 Thai genomes. We used Panaroo v1.3.2 [13] to build the pangenome and align the core-genome. The number of constant sites in the core-genome alignment was calculated using SNP-sites v2.5.1 and the "-C flag" [14]. Then, we used IQ-TREE v2.0.3 [15] to reconstruct the Maximum Likelihood (ML) phylogeny with 1000 bootstraps, the GTR+Gamma model and the "-fconst" flag to account for the number of constant sites in the alignment.

To place the Thai *S. suis* population in a global context we combined the 264 Thai genomes with a publicly available and curated global collection of 2424 *S. suis* genomes [16]. We enhanced this global collection by supplementing it with all zoonotic (n = 206) isolates as well as isolates closely related to lineages CC104 and CC233 (double locus variants (DLV) from their ST) (n =15) from a previously curated collection [17]. We manually removed duplicates as described above. In total, our global collection contained 2902 *S. suis* genomes (Supplementary Table 5). We generated a core-genome alignment as described above and reconstructed a Neighbour-Joining tree using FastTree v2.1.11 [18].

**Serotype 2 capsule locus switching analysis**

To investigate serotype 2 capsule acquisition by lineages CC104 and CC233 we reconstructed a serotype 2 capsule locus nucleotide phylogeny. We included all serotype 2 *S. suis* genomes from Thailand and a stratified random selection of 75 serotype 2 genomes from our global collection. Briefly, we stratified the serotype 2 genomes by country and CC and randomly selected a maximum of 3 genomes from each CC in each country (Supplementary Table 8.) To extract the serotype 2 locus, we built a custom database in ABRicate v1.01 (https://github.com/tseemann/abricate) containing all the cps2 locus genes and blasted them against our genomes with an 90% coverage and identity threshold. We used an inhouse script (https://github.com/boasvdp/extract_genes_ABRicate) to extract the genes and concatenated and aligned them with MAFFT v7.525. The alignment was fed into FastTree v2.1.11 to reconstruct a Maximum-Likelihood phylogeny [18]. Finally, we compared the mean pairwise nucleotide distance across the different zoonotic clades, in both their core genes and cps2 locus genes, using bootstrapping to gauge error with *ape* v5.8 in a custom *R* script [19].

**Identification of antimicrobial resistance genes**

Antimicrobial resistance (AMR) genes were identified using ABRicate v1.01 (https://github.com/tseemann/abricate) and the CARD database supplemented with recently identified variants of antimicrobial resistance determinants found in *S. suis* [20]. We used a threshold of 80% coverage and nucleotide identity for the blast. Mobile genetic elements (MGEs) were identified by scanning complete genomes with multiple databases (ICEberg 2.0, PHASTEST, IslandViewer4 and TnCentral) [21–24].

### Supplementary Tables

**Supplementary Table 1**. Metadata regarding zoonotic S. suis isolates collected during the national wide active surviellance study performed between 2006-2012. NA stands for not available. CSF stands for Cerebrospinal fluid. NT stands for non typable. CC stands for clonal complex. ST stands for sequence type. VAG stands for virulence associated genes.

| Isolate | Isolation Year | Hospital/Province | Region | Sex | Age year | Specimen | Diagnosis | Serotype | CC | ST | VAG profile |
|---|---|---|---|---|---|---|---|---|---|---|---|
| 23165 | 2006 | Lampang | North | Male | 50 | Blood | Acute Meningitis | 2 | 1 | 1 | epf+/sly+/mrp+ |
| 22783 | 2006 | Phetchaboon | North | Male | 36 | CSF | Acute Meningitis, Sepsis | 2 | 1 | 1 | epf+/sly+/mrp+ |
| 22690 | 2006 | Phetchaboon | North | Female | 50 | Blood | Acute Meningitis | 2 | 1 | 1 | epf-/sly+/mrp+ |
| 22692 | 2006 | Lampang | North | Male | 48 | Blood | Meningitis | 14 | 1 | 105 | epf+/sly+/mrp+ |
| 22694 | 2006 | Nan | North | Male | 60 | Blood | Acute Meningitis, Sepsis, Septic Shock | 2 | 25 | 25 | epf-/sly-/mrp* |
| 14980 | 2006 | Mahasarakam | Northeast | NA | NA | CSF | Acute Meningitis | 2 | 1 | 1 | epf+/sly+/mrp+ |
| 22693 | 2006 | Phetchaboon | North | Male | 56 | Blood | Bacterial Cardiolitis | 2 | 104 | 104 | epf-/sly+/mrp- |
| 22714 | 2006 | Nan | North | Male | 75 | Blood | Sepsis | 2 | 25 | 25 | epf-/sly-/mrp* |
| 22715 | 2006 | Phetchaboon | North | Male | 53 | Blood | Acute Meningitis | 2 | 1 | 1 | epf+/sly+/mrp+ |
| 22695 | 2007 | Phetchabul | North | Female | 62 | Blood | Septic Arthritis | 14 | 1 | 105 | epf+/sly+/mrp+ |
| 22717 | 2006 | Nakhonratchasima | Northeast | Male | 53 | CSF | Acute Meningitis | 2 | 1 | 1 | epf+/sly+/mrp+ |
| 22716 | 2006 | Lampang | North | Male | 58 | Blood | Sepsis | 2 | 104 | 104 | epf-/sly+/mrp- |
| 22696 | 2006 | Utraradit | North | Female | 76 | Blood | Spondyonin | 2 | 104 | 104 | epf-/sly+/mrp- |
| 23211 | 2006 | Lampang | North | Male | 73 | CSF | Acute Meningitis | 2 | 1 | 1 | epf+/sly+/mrp+ |
| 23161 | 2006 | Phetchaboon | North | Male | 74 | CSF | Acute Meningitis | 2 | 28 | 28 | epf-/sly-/mrp+ |
| 23207 | 2006 | Phayao | North | Male | 41 | Blood | Acute Meningitis, Sepsis | 2 | 1 | 1 | epf+/sly+/mrp+ |
| 22697 | 2006 | Phetchaboon | North | Male | 80 | Blood | Sepsis | 2 | 28 | 28 | epf-/sly-/mrp+ |
| 23164 | 2006 | Lampang | North | Female | 68 | CSF | Acute Meningitis | 2 | 28 | 28 | epf-/sly-/mrp+ |
| 22698 | 2006 | Phetchaboon | North | Female | 48 | Blood | Sepsis | 2 | 25 | 25 | epf-/sly-/mrp* |
| 23163 | 2006 | Nan | North | Female | 55 | Blood | Acute Meningitis, Sepsis, Septic Shock | 2 | 104 | 104 | epf-/sly+/mrp- |

| | | | | | | | | | | | |
|---|---|---|---|---|---|---|---|---|---|---|---|
| 23168 | 2006 | Phayao | North | Male | 49 | Blood | Sepsis, Bronchitis | 2 | 104 | 104 | epf-/sly+/mrp- |
| 23167 | 2006 | Chonburi | East | Male | 47 | Blood | Acute Meningitis | 2 | 1 | 1 | epf+/sly+/mrp+ |
| 23169 | 2006 | Phayao | North | Female | 80 | Blood | Sepsis, Septic Shock | 2 | 104 | 104 | epf-/sly+/mrp- |
| 23208 | 2006 | Phetchaboon | North | Male | 63 | CSF | Acute Meningitis | 2 | 1 | 1 | epf+/sly+/mrp+ |
| 23170 | 2006 | Phetchaboon | North | Male | 50 | Blood | Acute Meningitis, Sepsis | 2 | 1 | 1 | epf+/sly+/mrps |
| 21508 | 2006 | Nakornsawan | Central | Male | 38 | Blood | Chronic Heart Failure | 2 | 104 | 104 | epf-/sly+/mrp- |
| 23171 | 2006 | Phetchaboon | North | Male | 45 | CSF | Acute Meningitis | 2 | 1 | 1 | epf+/sly+/mrps |
| 21507 | 2006 | Nakornsawan | Central | Male | 52 | Blood | Sepsis,Radial Artery Occlusion | 2 | 1 | 1 | epf+/sly+/mrp+ |
| 23172 | 2006 | Phetchaboon | North | Male | 68 | Blood | Sepsis | 2 | 1 | 1 | epf+/sly+/mrp+ |
| 23135 | 2006 | Phetchaboon | North | Male | 52 | Blood | Sepsis | 2 | 104 | 104 | epf-/sly+/mrp- |
| 21711 | 2006 | Lampang | North | Male | 73 | Blood | NA | 2 | 1 | 1 | epf+/sly+/mrp+ |
| 21927 | 2006 | Nakornpanom | Northeast | Male | 23 | Blood | Sepsis | 2 | 104 | 104 | epf-/sly+/mrp- |
| 23139 | 2006 | Nakhonsawan | North | Female | 48 | Blood | Sepsis | 2 | 104 | 104 | epf-/sly+/mrp- |
| 23140 | 2006 | Phetchaboon | North | Female | 88 | Blood | Sepsis | 2 | 104 | 104 | epf-/sly+/mrp- |
| 23142/23143 | 2006 | Nakhonsawan | North | Female | 67 | Blood | Sepsis | 2 | 1 | 1 | epf+/sly+/mrp+ |
| 23141 | 2006 | Lampang | North | Female | 57 | Blood | Acute Meningitis, Sepsis | 2 | 1 | 1 | epf+/sly+/mrp+ |
| 21876 | 2006 | Phetchaburi | Central | Male | 18 | Blood | Acute Meningitis | 2 | 1 | 1 | epf+/sly+/mrp+ |
| 23147 | 2006 | Nakhonsawan | North | Male | 70 | Blood | Acute Meningitis | 2 | 1 | 1 | epf+/sly+/mrp+ |
| 21980 | 2006 | Phetchaboon | North | Female | 38 | Blood | Sepsis | 2 | 1 | 1 | epf+/sly+/mrp+ |
| 23153 | 2006 | Phetchaboon | North | Male | 73 | CSF | Acute Meningitis | 2 | 1 | 1 | epf+/sly+/mrp+ |
| 23148 | 2006 | Nakhonsawan | North | Female | 50 | Blood | Sepsis | 2 | 1 | 1 | epf+/sly+/mrp+ |
| 22740 | 2006 | Phetchaboon | North | Female | 40 | CSF | Acute Meningitis | 2 | 1 | 1 | epf+/sly+/mrp+ |
| 24828 | 2007 | Lampang | North | Male | 50 | CSF | Acute Meningitis | 2 | 1 | 1 | epf+/sly+/mrp+ |
| 23410 | 2007 | Lampang | North | Male | 44 | Blood | Sepsis | 2 | 25 | 25 | epf-/sly-/mrp+ |
| 23711 | 2007 | Lampang | North | Female | 62 | Blood | Sepsis | 2 | 1 | 1 | epf+/sly+/mrp+ |
| 24002 | 2007 | Phetchaboon | North | Male | 71 | CSF | Acute Meningitis | 2 | 1 | 1 | epf+/sly+/mrp+ |

| | | | | | | | | | | | |
|---|---|---|---|---|---|---|---|---|---|---|---|
| 24004 | 2007 | Lampang | North | Female | 59 | Blood | Sepsis | 2 | 1 | 1 | epf+/sly+/mrp+ |
| 24005 | 2007 | Lampang | North | Male | 38 | CSF | Acute Meningitis | 2 | 1 | 1 | epf+/sly+/mrp+ |
| 24524 | 2007 | Phayao | North | Female | 78 | Blood | Sepsis | 14 | 1 | 105 | epf+/sly+/mrp+ |
| 24523 | 2007 | Phetchaboon | North | Male | 61 | Blood | Sepsis | 2 | 104 | 104 | epf-/sly+/mrp- |
| 24525 | 2007 | Phetchaboon | North | Female | 47 | Blood | Sepsis | 2 | 104 | 104 | epf-/sly+/mrp- |
| 24536 | 2007 | Nakhonsawan | North | Male | 41 | Blood | Sepsis | 2 | 1 | 1 | epf+/sly+/mrp+ |
| 24526 | 2007 | Lampang | North | Male | 75 | CSF | Acute Meningitis | 2 | 1 | 126 | epf+/sly+/mrp+ |
| 24009 | 2007 | Bangkok | North | Male | 43 | Blood | Acute Meningitis, Septicemia | 2 | 1 | 1 | epf+/sly+/mrp+ |
| 24527 | 2007 | Phetchaboon | North | Male | 41 | Blood | Acute Meningitis | 2 | 25 | 103 | epf-/sly-/mrp* |
| 24698 | 2007 | Nan | North | Male | 47 | Blood | Acute Meningitis, Sepsis, Septic Shock | 2 | 1 | 1 | epf+/sly+/mrp+ |
| 24124 | 2007 | Phayao | North | Male | 62 | Blood | Acute Meningitis | 2 | 1 | 1 | epf+/sly+/mrp+ |
| 24125 | 2007 | Phayao | North | Female | 49 | Blood | Sepsis | 2 | 1 | 1 | epf+/sly+/mrp+ |
| 24537 | 2007 | Nakhonpanom | Northeast | Male | 56 | CSF | Acute Meningitis | 2 | 1 | 1 | epf+/sly+/mrp+ |
| 24376 | 2007 | Sukhothai | North | Male | 48 | Blood | Acute Meningitis | 2 | 1 | 1 | epf+/sly+/mrp+ |
| 24205 | 2007 | Lampoon | North | Male | 40 | Blood | Sepsis | 2 | 25 | 25 | epf-/sly-/mrp* |
| 24206 | 2007 | Phayao | North | Male | 70 | Blood | Sepsis | 2 | 1 | 1 | epf+/sly+/mrp+ |
| 24213 | 2007 | Phayao | North | NA | NA | Blood | Sepsis | 2 | 1 | 1 | epf+/sly+/mrp+ |
| 24377 | 2007 | Sukhothai | North | Male | 68 | CSF | Acute Meningitis | 2 | 1 | 1 | epf+/sly+/mrp+ |
| 24378 | 2007 | Sukhothai | North | Male | 45 | CSF | Acute Meningitis | 2 | 1 | 1 | epf+/sly+/mrp+ |
| 24379 | 2007 | Sukhothai | North | Male | 44 | CSF | Acute Meningitis | 2 | 1 | 1 | epf+/sly+/mrp+ |
| 24451 | 2007 | Lampang | North | Male | 70 | CSF | Meningitis | 14 | 1 | 105 | epf+/sly+/mrp+ |
| 26152 | 2007 | Lopburi | Central | Male | 35 | CSF | Acute Meningitis | 2 | 1 | 1 | epf+/sly+/mrp+ |
| 26151 | 2007 | Nakhonsawan | Central | Male | 45 | Blood | Sepsis | 2 | 1 | 1 | epf+/sly+/mrp+ |
| 26147/26164 | 2007 | Nakhonsawan | Central | Male | 63 | Blood | Acute Meningitis | 2 | 1 | 1 | epf+/sly+/mrp+ |
| 26162/26163 | 2007 | Nakhonsawan | Central | Male | 56 | Blood | Sepsis | 2 | 104 | 104 | epf-/sly+/mrp- |
| 26153 | 2007 | Sukhothai | North | Male | 63 | Blood | Sepsis | 2 | 1 | 1 | epf+/sly+/mrp+ |

| | | | | | | | | | | | |
|---|---|---|---|---|---|---|---|---|---|---|---|
| 24529 | 2007 | Utraradit | North | Female | 32 | Blood | Acute Meningitis | 2 | 104 | 104 | epf-/sly+/mrp- |
| 24528 | 2007 | Phayao | North | Male | 59 | Blood | Septicemia | 2 | 104 | 104 | epf-/sly+/mrp- |
| 25437 | 2007 | Nan | North | Male | 72 | Blood | Sepsis | 2 | 25 | 25 | epf-/sly-/mrp* |
| 24579 | 2007 | Suphanburi | Central | Male | 60 | Blood | Sepsis, Acute Abdominal Pain | 2 | 104 | 104 | epf-/sly+/mrp- |
| 24651 | 2007 | Phayao | North | Male | 39 | CSF | Acute Meningitis | 2 | 1 | 1 | epf+/sly+/mrp+ |
| 24652 | 2007 | Nakhonsawan | North | Male | 64 | Blood | Sepsis | 2 | 104 | 104 | epf-/sly+/mrp- |
| 24883 | 2007 | Suphanburi | Central | Male | 61 | Blood | Acute Gastroenteritis | 2 | 104 | 104 | epf-/sly+/mrp- |
| 25436 | 2007 | Utraradit | North | Female | 74 | Blood | Sepsis | 2 | 25 | 25 | epf-/sly-/mrp* |
| 26154 | 2007 | Phetchaboon | North | Male | 29 | CSF | Acute Meningitis | 2 | 1 | 1 | epf+/sly+/mrp+ |
| 26155 | 2007 | Phetchaboon | North | Male | 44 | CSF | Acute Meningitis | 2 | 1 | 1 | epf+/sly+/mrp+ |
| 26156 | 2007 | Tak | North | Female | 43 | Blood | Acute Meningitis | 2 | 1 | 1 | epf+/sly+/mrp+ |
| 24826 | 2007 | Lampang | North | Male | 75 | Blood | Acute Meningitis, Sepsis | 2 | 1 | 1 | epf+/sly+/mrp+ |
| 22213 | 2007 | Lampang | North | Female | 65 | CSF | Acute Meningitis | 2 | 1 | 1 | epf+/sly+/mrp+ |
| 25438 | 2007 | Phayao | North | Male | 43 | CSF | Acute Meningitis | 2 | 1 | 1 | epf+/sly+/mrp+ |
| 23166 | 2007 | Nakhonsawan | North | Female | 53 | Blood | Sepsis | 2 | 104 | 104 | epf-/sly+/mrp- |
| 26161 | 2007 | Phayao | North | Female | 56 | Blood | Sepsis | 2 | 104 | 104 | epf-/sly+/mrp- |
| 24884 | 2007 | Phayao | North | Male | 56 | Blood | Sepsis | 2 | 104 | 104 | epf-/sly+/mrp- |
| 24827 | 2007 | Lampang | North | Female | 93 | Blood | Acute Meningitis | 2 | 1 | 1 | epf+/sly+/mrp+ |
| 26158 | 2007 | Phayao | North | Male | 77 | CSF | Acute Meningitis | 2 | 1 | 1 | epf+/sly+/mrp+ |
| 26160 | 2007 | Nakhonsawan | North | Male | 52 | CSF | Acute Meningitis, Sepsis | 2 | 1 | 1 | epf+/sly+/mrp+ |
| 25014 | 2007 | Phayao | North | Male | 47 | Blood | Sepsis | 2 | 104 | 104 | epf-/sly+/mrp- |
| 25015 | 2007 | Phayao | North | Female | 49 | Blood | Sepsis | 2 | 104 | 104 | epf-/sly+/mrp- |
| 24829 | 2007 | Sukhothai | North | Male | 51 | CSF | Acute Meningitis | 2 | 1 | 126 | epf+/sly+/mrp+ |
| 25378 | 2007 | Nan | North | Male | 53 | CSF | Acute Meningitis, Sepsis, Septic Shock | 2 | 1 | 1 | epf+/sly+/mrp+ |
| 24214 | 2007 | Lopburi | Central | Male | 50 | CSF | Acute Meningitis | 2 | 1 | 1 | epf+/sly+/mrp+ |
| 25016 | 2007 | Phayao | North | Male | 53 | Blood | Acute Meningitis, Sepsis | 2 | 104 | 104 | epf-/sly+/mrp- |

| | | | | | | | | | | | |
|---|---|---|---|---|---|---|---|---|---|---|---|
| 24886 | 2007 | Utraradit | North | Male | 51 | Blood | Sepsis | 2 | 104 | 104 | epf-/sly+/mrp- |
| 24887 | 2007 | Utraradit | North | Male | 69 | Blood | Sepsis | 2 | 104 | 104 | epf-/sly+/mrp- |
| 25018 | 2007 | Chiang Rai | North | Male | 48 | CSF | Bacterial Meningitis | 2 | 1 | 1 | epf+/sly+/mrp+ |
| 25042 | 2007 | Chiang Rai | North | Female | 74 | CSF | Acute Meningitis | 2 | 1 | 1 | epf+/sly+/mrp+ |
| 25377 | 2007 | Utraradit | North | Female | 71 | Blood | Sepsis, Septic Arthritis | 2 | 1 | 1 | epf+/sly+/mrp+ |
| 25686 | 2007 | Chonburi | East | Female | 80 | Blood | Septic Arthritis | 2 | 25 | 103 | epf-/sly-/mrp* |
| 25384 | 2007 | Phayao | North | Male | 58 | Blood | Sepsis | 2 | 1 | 1 | epf+/sly+/mrp+ |
| 23149 | 2007 | Nakhonsawan | North | NA | NA | Blood | Sepsis | 2 | 1 | 1 | epf+/sly+/mrp+ |
| 25382 | 2007 | Tak | North | Male | 68 | Blood | Acute Meningitis | 2 | 1 | 126 | epf+/sly+/mrp+ |
| 25443 | 2007 | Chaiyaphum | Northeast | Male | 53 | Blood | Sepsis | 2 | 1 | 1 | epf+/sly+/mrp+ |
| 25444 | 2007 | Tak | North | Male | 63 | Blood | Sepsis | 2 | 1 | 1 | epf+/sly+/mrp+ |
| 25687 | 2007 | Chaiyaphum | Northeast | Male | 63 | Blood | Acute Meningitis, Sepsis | 2 | 1 | 1 | epf+/sly+/mrp+ |
| 25688 | 2007 | Nakhonsawan | North | Male | 76 | Blood | Acute Meningitis | 2 | 1 | 1 | epf+/sly+/mrp+ |
| 25779/25780/25778/25840 | 2007 | Tak | North | NA | 48 | Blood | Sepsis | 14 | 1 | 105 | epf+/sly+/mrp+ |
| 25777 | 2007 | Nan | North | Male | 92 | Blood | Sepsis | 2 | 104 | 104 | epf-/sly+/mrp- |
| 25701 | 2007 | Phetchaboon | North | Female | 79 | CSF | Acute Meningitis | 2 | 1 | 1 | epf+/sly+/mrps |
| 25781 | 2007 | Saraburi | Central | Male | 53 | Blood | Acute Meningitis | 2 | 25 | 103 | epf-/sly-/mrp+ |
| 26012 | 2008 | Phetchabul | North | Female | 78 | Blood | Septic Arthritis | 14 | 1 | 105 | epf+/sly+/mrp+ |
| 26013 | 2007 | Sukhothai | North | Male | 47 | Blood | Sepsis, Severe MR | 2 | 1 | 1 | epf+/sly+/mrp+ |
| 27071 | 2008 | Lampang | North | Male | 68 | Blood | Meningitis | 14 | 1 | 105 | epf+/sly+/mrp+ |
| 26126 | 2008 | Lampang | North | Male | 55 | Blood | Sepsis | 2 | 1 | 1 | epf+/sly+/mrp+ |
| 26127 | 2008 | Lampang | North | Male | 48 | CSF | Acute Meningitis | 2 | 104 | 104 | epf-/sly+/mrp- |
| 26125 | 2008 | Lampang | North | Male | 64 | CSF | Acute Meningitis | 2 | 1 | 1 | epf+/sly+/mrp+ |
| 26128 | 2008 | Lampang | North | Male | 53 | Blood | Acute Meningitis | 2 | 25 | 25 | epf-/sly-/mrp+ |
| 26390 | 2008 | Bangkok | Central | Male | 71 | Blood | Meningitis | 14 | 1 | 105 | epf+/sly+/mrp+ |
| 26386 | 2008 | Nakhonsawan | North | Male | 47 | Blood | Acute Meningitis | 2 | 1 | 1 | epf+/sly+/mrp+ |

| | | | | | | | | | | | |
|---|---|---|---|---|---|---|---|---|---|---|---|
| 26349 | 2008 | Nakornpathom | Central | NA | NA | Blood | Sepsis | 2 | 104 | 104 | epf-/sly+/mrp- |
| 22712 | 2008 | Phetchaboon | North | Male | 82 | CSF | Acute Meningitis | 2 | 1 | 1 | epf+/sly+/mrp+ |
| 26484 | 2008 | Phetchaboon | North | Male | 53 | Blood | Acute Meningitis | 2 | 1 | 1 | epf+/sly+/mrp+ |
| 26648 | 2008 | Phitsanulok | North | Female | 57 | Blood | Septic Shock, Severe CAP, Act Renal Failure | 2 | 1 | 1 | epf+/sly+/mrp+ |
| 26431 | 2008 | Sakaew | East | Male | 44 | Blood | Sepsis | 2 | 104 | 104 | epf-/sly+/mrp- |
| 26645 | 2008 | Utraradit | North | Male | 69 | Blood | Sepsis | 2 | 104 | 104 | epf-/sly+/mrp- |
| 26647 | 2008 | Nakornratchasima | Northeast | Male | 76 | Blood | Acute Meningitis | 2 | 1 | 1 | epf+/sly+/mrp+ |
| 26646 | 2008 | Phayao | North | Female | 67 | Blood | Sepsis | 2 | 104 | 104 | epf-/sly+/mrp- |
| 27119 | 2008 | Sukhothai | North | Male | 69 | Blood | Sepsis | 2 | 104 | 104 | epf-/sly+/mrp- |
| 26717 | 2008 | Phitsanulok | North | Male | 64 | CSF | Acute Meningitis | 2 | 1 | 1 | epf+/sly+/mrp+ |
| 26108 | 2008 | Kon Kaen | Northeast | Male | 70 | Blood | Acute Meningitis, Sepsis | 2 | 1 | 1 | epf+/sly+/mrp+ |
| 26706 | 2008 | Utraradit | North | Male | 60 | Blood | Acute Meningitis, Septic Arthritis | 2 | 1 | 1 | epf+/sly+/mrp+ |
| 26718 | 2008 | Phitsanulok | North | Female | 55 | CSF | Acute Meningitis | 2 | 1 | 1 | epf+/sly+/mrp+ |
| 26719 | 2008 | Phitsanulok | North | Male | 69 | Blood | Sepsis | 2 | 104 | 104 | epf-/sly+/mrp- |
| 27024 | 2008 | Lampang | North | Male | 40 | CSF | Meningitis | 14 | 1 | 127 | epf+/sly+/mrp+ |
| 27125 | 2008 | Saraburi | Central | Male | 43 | Blood | Sepsis, Hypoglycemia, Group D Streptococcus Septicemia | 2 | 1 | 1 | epf+/sly+/mrp+ |
| 27100 | 2008 | Tak | North | Male | 43 | Blood | Bacterial Infection | 2 | 25 | 25 | epf-/sly-/mrp* |
| 27558 | 2008 | Tak | North | Male | 51 | CSF | Acute Meningitis | 2 | 1 | 1 | epf+/sly+/mrp+ |
| 27134 | 2008 | Chiang Rai | North | Female | 49 | Blood | Acute Pyelonephritis | 2 | 1 | 1 | epf+/sly+/mrp+ |
| 27133 | 2008 | Chiang Rai | North | Male | 42 | Blood | Oligoarthritis | 2 | 1 | 1 | epf+/sly+/mrp+ |
| 27103 | 2008 | Tak | North | Male | 40 | Blood | Sepsis | 2 | 25 | 25 | epf-/sly-/mrp* |
| 27573 | 2008 | Chiang Rai | North | Female | 57 | CSF | Acute Meningitis | 2 | 1 | 1 | epf+/sly+/mrp+ |
| 27571 | 2008 | Chiang Rai | North | Male | 59 | Blood | Febrial Illness, Meningitis | 2 | 1 | 1 | epf+/sly+/mrp+ |
| 27579 | 2008 | Lampang | North | Male | 69 | Blood | Sepsis | 2 | 104 | 104 | epf-/sly+/mrp- |
| 27554 | 2008 | Phetchaboon | North | Male | 43 | Blood | Acute Meningitis | 2 | 1 | 1 | epf+/sly+/mrp+ |

| | | | | | | | | | | | |
|---|---|---|---|---|---|---|---|---|---|---|---|
| 27339 | 2008 | Phetchaboon | North | Male | 38 | Blood | Acute Meningitis | 2 | 1 | 1 | epf+/sly+/mrp+ |
| 27714 | 2008 | Ratchaburi | Central | Male | 64 | CSF | Acute Meningitis | 2 | 1 | 1 | epf+/sly+/mrp+ |
| 27578 | 2008 | Phetchabul | North | Female | 63 | CSF | Meningitis | 14 | 1 | 105 | epf+/sly+/mrp+ |
| 27557 | 2008 | Nakhonsawan | North | Female | 50 | Blood | Acute Meningitis | 2 | 1 | 1 | epf+/sly+/mrp+ |
| 27560 | 2008 | Phetchaboon | North | Male | 59 | CSF | Acute Meningitis | 2 | 1 | 1 | epf+/sly+/mrp+ |
| 27567 | 2008 | Tak | North | Male | 64 | Blood | Sepsis | 2 | 25 | 25 | epf-/sly-/mrp+ |
| 27574 | 2008 | Phayao | North | Male | NA | Blood | Septicemia | 2 | 1 | 1 | epf+/sly+/mrp+ |
| 27562 | 2008 | Phitsanulok | North | Male | 62 | Blood | Acute Meningitis | 2 | 1 | 1 | epf+/sly+/mrp+ |
| 27807 | 2008 | Phetchaboon | North | Male | 43 | Blood | Congenitive Heart Failure (CHF) | 2 | 104 | 104 | epf-/sly+/mrp- |
| 27576 | 2008 | Utraradit | North | Male | 48 | Blood | Sepsis, Acute Pyelonephritis, Low Back Pain | 2 | 1 | 1 | epf+/sly+/mrp+ |
| 27577 | 2008 | Phetchaboon | North | Female | 42 | Blood | Acute Meningitis | 2 | 1 | 1 | epf+/sly+/mrp+ |
| 27518 | 2008 | Tak | North | Male | 53 | Blood | Acute Meningitis | 2 | 1 | 1 | epf+/sly+/mrp+ |
| NA | 2008 | Tak | North | Male | 61 | Blood | Acute Meningitis | 2 | 1 | 1 | epf+/sly+/mrp+ |
| 27937 | 2008 | Tak | North | Female | NA | Blood | Acute Meningitis | 2 | 1 | 1 | epf+/sly+/mrp+ |
| 27938 | 2008 | Nan | North | Female | 49 | Blood | Sepsis | 2 | 104 | 104 | epf- /sly+/mrp- |
| 27939 | 2008 | Nan | North | Male | 84 | Blood | Sepsis | 2 | 104 | 104 | epf- /sly+/mrp- |
| 27940 | 2008 | Nan | North | Female | 55 | Blood | Sepsis | 2 | 104 | 104 | epf- /sly+/mrp- |
| 27941 | 2008 | Nan | North | Male | 41 | Blood , CSF | Meningitis , Sepsis | 2 | 1 | 1 | epf +/sly+/mrp+ |
| 27942 | 2008 | Lampang | North | Male | 55 | Blood , CSF | Meningitis | 2 | 1 | 1 | epf+/sly+/mrp+ |
| 27943 | 2008 | Phitsanulok | North | Female | NA | NA | NA | 2 | 104 | 104 | epf- /sly+/mrp- |
| 27944 | 2008 | Phitsanulok | North | Male | 57 | CSF , Stool | Spondylodiscitis , Sepsis , Anemia | 2 | 1 | 1 | epf+ /sly+/mrp+ |
| 27968 | 2008 | Srakaew | East | Male | 86 | Blood | Pneumonia , Septicemia , Hypoglycemia , Acute on top ESRD | 2 | 233 | 233 | epf -/sly+/mrp- |
| 27996 | 2008 | Tak | North | Female | NA | Haemoculture | Sepsis | 2 | 1 | 1 | epf +/sly+/mrp+ |
| 28069 | 2008 | Chonburi | East | Male | 41 | Blood , CSF | Meningitis | 2 | 25 | 25 | epf- /sly- /mrp*** |

| | | | | | | | | | | | |
|---|---|---|---|---|---|---|---|---|---|---|---|
| NA | 2008 | Phitsanulok | North | Male | NA | Blood | Acute Meningitis | 2 | 1 | 1 | epf+/sly+/mrp+ |
| 27935 | 2008 | Tak | North | Female | 51 | Blood | Acute Meningitis | 2 | 1 | 1 | epf+/sly+/mrp+ |
| 27563 | 2008 | Bangkok | Central | NA | NA | Blood | NA | 2 | 104 | 104 | epf-/sly+/mrp- |
| 27564 | 2008 | Samutsongkram | Central | NA | NA | Blood | NA | 2 | 1 | 1 | epf+/sly+/mrp+ |
| 27569 | 2008 | Nakhonsawan | North | Male | 70 | Blood | Septicemia | 2 | 1 | 1 | epf+/sly+/mrp+ |
| 27936 | 2008 | Tak | North | Female | 48 | Blood | Sepsis | 2 | 1 | 1 | epf+/sly+/mrp+ |
| 26705 | 2008 | Utraradit | North | Female | 49 | CSF | Acute Meningitis, Sepsis | 2 | 1 | 1 | epf+/sly+/mrp+ |
| 27946 | 2008 | Suphanburi | Central | Female | 59 | Blood , CSF | Meningitis | 2 | 1 | 1 | epf+ /sly+/mrp+ |
| 27947 | 2008 | Chiang Mai | North | Male | NA | Haemoculture | Sepsis | 2 | 1 | 1 | epf +/sly+/mrp+ |
| 27948 | 2008 | Chiang Mai | North | Male | NA | Haemoculture | Sepsis | 2 | 1 | 1 | epf +/sly+/mrp+ |
| 27949 | 2008 | Chiang Mai | North | Male | NA | Haemoculture | Sepsis | 2 | 1 | 1 | epf+ /sly+/mrp+ |
| 27950 | 2008 | Chiang Mai | North | Male | NA | CSF | Meningitis | 2 | 1 | 1 | epf +/sly+/mrp+ |
| 27951 | 2008 | Chiang Mai | North | Male | NA | CSF | Meningitis | 2 | 1 | 1 | epf+ /sly+/mrp+ |
| 27952 | 2008 | Chiang Mai | North | Male | NA | Haemoculture | Sepsis | 2 | 1 | 1 | epf +/sly+/mrp+ |
| 27953 | 2008 | Chiang Mai | North | Male | NA | Haemoculture | Sepsis | 2 | 1 | 1 | epf+ /sly+/mrp+ |
| 27955 | 2008 | Chiang Mai | North | Male | NA | CSF | Meningitis | 2 | 1 | 1 | epf+ /sly+/mrp+ |
| 27956 | 2008 | Chiang Mai | North | Male | NA | Haemoculture | Sepsis | 2 | 1 | 1 | epf +/sly+/mrp+ |
| 27957 | 2008 | Chiang Mai | North | Male | NA | Haemoculture | Sepsis | 2 | 1 | 1 | epf+ /sly+/mrp+ |
| 27958 | 2008 | Chiang Mai | North | Male | NA | Haemoculture | Sepsis | 2 | 1 | 1 | epf +/sly+/mrp+ |
| 27960 | 2008 | Chiang Mai | North | Male | NA | Haemoculture | Sepsis | 2 | 1 | 1 | epf +/sly+/mrp+ |
| 27961 | 2008 | Sukhothai | North | Male | 67 | Blood , CSF | Sepsis | 2 | 1 | 1 | epf +/sly+/mrp+ |
| 27963 | 2008 | Sukhothai | North | Female | 76 | Blood | Severe AR | 2 | 28 | 28 | epf -/sly-/mrp+ |
| 27964 | 2008 | Sukhothai | North | Male | 79 | Blood | Septic Arthritis | 14 | 1 | 105 | epf+/sly+/mrp+ |

| | | | | | | | | | | | |
|---|---|---|---|---|---|---|---|---|---|---|---|
| 27964 | 2008 | Sukhothai | North | Male | NA | Haemoculture | Sepsis | 14 | 1 | 105 | epf +/sly+/mrp+ |
| 28074 | 2008 | Phitsanulok | North | Male | NA | Blood | Sepsis | 14 | 1 | 105 | epf +/sly+/mrp+ |
| 28075 | 2008 | Phitsanulok | North | Female | NA | Blood | Sepsis | 14 | 1 | 237 | epf-/sly+/mrpS |
| 28076 | 2008 | Phitsanulok | North | Male | NA | Blood | Sepsis | 14 | 1 | 105 | epf+ /sly+/mrp+ |
| 28077 | 2008 | Phitsanulok | North | Male | NA | Blood | Sepsis | 2 | 104 | 104 | epf -/sly+/mrp- |
| 28078 | 2008 | Phetchabul | North | Male | NA | CSF | Meningitis | 2 | 1 | 1 | epf +/sly+/mrp+ |
| 28080 | 2008 | Phetchabul | North | Male | NA | CSF | Meningitis | 2 | 1 | 1 | epf+ /sly+/mrp+ |
| 28082 | 2008 | Phitsanulok | North | Male | NA | Blood | NA | 2 | 1 | 1 | epf +/sly+/mrp+ |
| 28083 | 2008 | Lampang | North | Male | 54 | Blood , CSF | Meningitis | 2 | 1 | 1 | epf+ /sly+/mrp+ |
| 28085 | 2008 | Lampang | North | Male | 78 | Blood , CSF | Meningitis | 2 | 1 | 1 | epf +/sly/+mrp+ |
| 28088 | 2008 | Saraburi | Central | Male | 57 | Blood , CSF | Meningitis , Sepsis | 2 | 1 | 1 | epf +/sly/+mrp+ |
| 28089 | 2008 | Saraburi | Central | Male | 60 | Blood , CSF | Meningitis | 2 | 1 | 1 | epf +/sly+/mrp+ |
| 28729 | 2008 | Nakhornratchasima | Northeast | Male | NA | Blood | Sepsis | 2 | 1 | 1 | epf +/sly/+mrp+ |
| 28730 | 2008 | Sukhothai | North | Female | 77 | Blood | Pneumonia , Sepsis | 2 | 104 | 104 | epf -/sly+/mrp- |
| 27715 | 2008 | Ratchaburi | Central | Female | 72 | Blood | Chronic Renal Failure | 2 | 104 | 104 | epf-/sly+/mrp- |
| 28731 | 2008 | Chiang Rai | North | Female | 50 | CSF | Sepsis , ARF | 2 | 1 | 1 | epf +/sly+/mrp+ |
| 28732 | 2008 | Lampang | North | Male | 85 | Blood , CSF | Meningitis | 2 | 1 | 1 | epf +/sly+/mrp+ |
| 28733 | 2008 | Sukhothai | North | Male | NA | CSF | Meningitis | 2 | 1 | 1 | epf +/sly+/mrp+ |
| 28734 | 2008 | Chiang Rai | North | Male | 67 | Blood , CSF | Meningitis | 2 | 1 | 1 | epf +/sly+/mrp+ |
| 28736 | 2008 | Phetchabul | North | Male | NA | Blood | Sepsis | 2 | 1 | 1 | epf+ /sly+/mrp+ |
| 28737 | 2008 | Lampang | North | Male | 53 | Blood | Septic Shock | 2 | 104 | 104 | epf- /sly+/mrp- |
| 28739 | 2008 | Nan | North | Male | 51 | Blood | Arthritis , Sepsis | 2 | 28 | 28 | epf-/sly-/mrp+ |
| 28740 | 2008 | Nan | North | Male | 76 | Blood , CSF | Arthritis , Sepsis , Meningitis | 2 | 1 | 1 | epf +/sly+/mrp+ |

| | | | | | | | | | | | |
|---|---|---|---|---|---|---|---|---|---|---|---|
| 28741 | 2008 | Suphanburi | Central | Female | 55 | Blood | Pneumonia , Septic Shock | 2 | 104 | 104 | epf-/sly+/mrp- |
| 28744 | 2008 | Phitsanulok | North | Male | 49 | Blood | Septic Shock , Hypervolemic Shock | 2 | 104 | 104 | epf- /sly+/mrp- |
| 29276 | 2008 | Singburi | Central | Male | 31 | Blood , CSF | Meningitis | 2 | 104 | 104 | epf -/sly+/mrp- |
| 29279 | 2008 | Lampang | North | Female | 51 | Blood , CSF | Infective Endocarditis | 2 | 104 | 104 | epf -/sly+/mrp- |
| 29281 | 2008 | Phayao | North | Female | NA | Haemoculture | Sepsis | 2 | 104 | 104 | epf -/sly+/mrp- |
| 29282 | 2008 | Phayao | North | Male | NA | Haemoculture | Sepsis | 2 | 104 | 104 | epf -/sly+/mrp- |
| 29901 | 2008 | Phetchabul | North | Female | NA | CSF | Meningitis | 2 | 1 | 1 | epf +/sly+/mrp+ |
| 29903 | 2008 | Chaiyapum | Northeast | Male | 62 | Blood , CSF | Meningitis | 2 | 1 | 1 | epf +/sly+/mrp+ |
| 29904 | 2009 | Chiang Rai | North | Male | 70 | Blood | Arthritis | 2 | 1 | 1 | epf +/sly+/mrp+ |
| 29905 | 2009 | Nan | North | Male | 45 | Blood | Endocarditis , Sepsis | 2 | 104 | 104 | epf-/sly+/mrp- |
| 29906 | 2009 | Saraburi | Central | Male | 47 | CSF | Meningitis , Pneumonia | 2 | 1 | 1 | epf +/sly+/mrp+ |
| 29907 | 2009 | Uttraradit | North | Male | 67 | Blood , CSF | Meningo Encephalitis | 2 | 1 | 1 | epf +/sly+/mrp+ |
| 29908 | 2009 | Phetchabul | North | Male | NA | Blood | Sepsis | 2 | 1 | 1 | epf +/sly+/mrp+ |
| 29909 | 2009 | Bangkok | Central | Female | NA | Haemoculture | Sepsis | 2 | 104 | 104 | epf -/sly+/mrp- |
| 29911 | 2009 | Phetchabul | North | Male | NA | Blood | Sepsis | 2 | 1 | 1 | epf +/sly+/mrp+ |
| 29912 | 2009 | Lampang | North | Male | 52 | Blood , CSF | Meningitis | 2 | 104 | 104 | epf -/sly+/mrp- |
| 29915 | 2009 | Phitsanulok | North | Female | 49 | Blood | Septic Shock | 2 | 104 | 104 | epf -/sly+/mrp- |
| 30098 | 2009 | Kanchanaburi | Central | Female | 61 | Blood | Spondylodiscitis , Sepsis , UTI , DM , HT , DLP | 2 | 104 | 104 | epf -/sly+/mrp- |
| 30188 | 2009 | Phetchabul | North | Male | NA | Blood | Sepsis | 2 | 1 | 1 | epf +/sly+/mrp+ |
| 30189 | 2009 | Phetchabul | North | Male | NA | Blood | Sepsis | 2 | 104 | 104 | epf-/sly+/mrp- |
| 30190 | 2009 | Phitsanulok | North | Male | 69 | Blood , CSF | Meningitis , Act Renal Failure | 2 | 25 | 25 | epf-/sly-/mrp*** |
| 30192 | 2009 | Tak | North | Male | 34 | Blood | Meningitis , Schizophemia | 2 | 25 | 25 | epf -/sly-/mrp+ |
| 30194 | 2009 | Phitsanulok | North | Male | 71 | Blood | Septic Shock , AF C RVR , SVT , Respiratory Failure | 2 | 104 | 104 | epf- /sly+/mrp- |
| 30196 | 2009 | Saraburi | Central | Male | 65 | Blood , CSF | Meningitis | 2 | 25 | 25 | epf-/sly-/mrp*** |

| | | | | | | | | | | | |
|---|---|---|---|---|---|---|---|---|---|---|---|
| 30209 | 2009 | Lampang | North | Male | 41 | Blood | Septic Shock | 2 | 1 | 1 | epf +/sly+/mrp+ |
| 30198 | 2009 | Sukhothai | North | Male | 58 | Blood , CSF | Meningitis , Sepsis | 14 | 1 | 105 | epf +/sly+/mrp+ |
| 30214 | 2009 | Lampang | North | Male | 67 | Blood | Streptococcus viridan IE | 2 | 1 | 1 | epf +/sly+/mrp+ |
| 30215 | 2009 | Tak | North | Male | 49 | Blood | Mentaland Bahavioural disorders due to use of alcohols, other Septiccemia | 2 | 1 | 1 | epf +/sly+/mrp+ |
| 30331 | 2009 | Lampang | North | Male | 92 | CSF | Septic Shock | 2 | 1 | 1 | epf +/sly+/mrp+ |
| 30340 | 2009 | Phayao | North | Male | NA | CSF, Blood | Meningitis | 2 | 1 | 1 | epf +/sly+/mrp+ |
| 30341 | 2009 | Nan | North | Male | 69 | Blood | Pneumonia , Sepsis | 2 | 104 | 104 | epf -/sly+/mrp- |
| 30342 | 2009 | Phitsanulok | North | Male | NA | Blood | NA | 2 | 104 | 104 | epf -/sly+/mrp- |
| 30389 | 2009 | Srakaew | East | Male | NA | Haemoculture | Sepsis | 2 | 233 | 233 | epf -/sly+/mrp- |
| 30321 | 2009 | Yasothon | Northeast | Male | NA | CSF | Meningitis | 2 | 1 | 1 | epf +/sly+/mrp+ |
| 30327 | 2009 | Phetchabul | North | Male | NA | Blood | Sepsis | 2 | 104 | 104 | epf- /sly+/mrp- |
| 30333 | 2009 | Tak | North | Male | 46 | Blood | Inflammatory Spondylopathies | 14 | 1 | 105 | epf +/sly+/mrp+ |
| 30346 | 2009 | Phitsanulok | North | Male | 51 | Blood | Endocarditis , CHF , Severe MR | 2 | 233 | 233 | epf -/sly+/mrp- |
| 30348 | 2009 | Lampang | North | Male | 69 | Blood , CSF | B-StreptococcusGr. A Septicemia | 2 | 1 | 1 | epf +/sly+/mrp+ |
| 30349 | 2009 | Lampang | North | Female | 92 | Blood | Sepsis | 2 | 1 | 1 | epf +/sly+/mrp+ |
| 30350 | 2009 | Lampang | North | Male | 71 | Blood , CSF | Meningitis | 2 | 1 | 1 | epf +/sly+/mrp+ |
| 30351 | 2009 | Bangkok | Central | Male | NA | CSF | Meningitis | 2 | 1 | 1 | epf +/sly+/mrp+ |
| 30352 | 2009 | Uttraradit | North | Male | 45 | Blood | Endocarditis | 2 | 104 | 104 | epf -/sly+/mrp- |
| 30353 | 2009 | Sukhothai | North | Male | 54 | Blood , CSF | Meningitis | 2 | 1 | 1 | epf +/sly+/mrp+ |
| 30354 | 2009 | Sukhothai | North | Male | 62 | Blood , CSF | Meningitis | 2 | 1 | 1 | epf +/sly+/mrp+ |
| 30390 | 2009 | Srakaew | East | Male | NA | Haemoculture | Sepsis | 2 | 1 | 1 | epf +/sly+/mrp+ |
| 30356 | 2009 | Phetchabul | North | Male | NA | Blood | Sepsis | 2 | 104 | 104 | epf -/sly+/mrp- |
| 30528 | 2009 | Nakhonpanom | Northeast | Male | 59 | Blood | Sepsis | 2 | 1 | 1 | epf +/sly+/mrp+ |

| | | | | | | | | | | | |
|---|---|---|---|---|---|---|---|---|---|---|---|
| 30679 | 2009 | Chiang Rai | North | Female | 42 | Blood | Sepsis | 2 | 1 | 1 | epf +/sly+/mrp+ |
| 30680 | 2009 | Phitsanulok | North | Male | 55 | Blood , CSF | Meningitis , Hypolemic Shock | 2 | 1 | 1 | epf +/sly+/mrp+ |
| 30683 | 2009 | Sukhothai | North | Female | 41 | Blood , CSF | Meningitis | 2 | 1 | 1 | epf +/sly+/mrp+ |
| 30682 | 2009 | Uttraradit | North | Male | 45 | Blood | Endocarditis | 2 | 104 | 104 | epf -/sly+/mrp- |
| 30684 | 2009 | Nakhornratchasima | Northeast | Male | NA | CSF | Meningitis | 2 | 1 | 1 | epf +/sly+/mrp+ |
| 30685 | 2009 | Phitsanulok | North | Female | 50 | Blood | Acutet Ascending Cholangitis | 2 | 1 | 1 | epf +/sly+/mrp+ |
| 30689 | 2009 | Srakaew | East | Male | 42 | Blood | Septic Shock , Rlo Leptospirosis , Acute Gastroenteritis | 2 | 104 | 104 | epf -/sly+/mrp- |
| 30690 | 2009 | Nakhonpanom | Northeast | Male | 30 | Blood , CSF | Meningitis | 2 | 1 | 1 | epf +/sly+/mrp+ |
| 30700 | 2009 | Lampang | North | Male | 90 | Blood | Septic Shock | 2 | 1 | 1 | epf +/sly+/mrp+ |
| 30701 | 2009 | Uttraradit | North | Male | 45 | Blood , CSF | Meningitis | 14 | 1 | 105 | epf +/sly+/mrp+ |
| 30702 | 2009 | Uttraradit | North | Female | NA | Haemoculture | Sepsis | 2 | 104 | 104 | epf -/sly+/mrp- |
| 31087 | 2009 | Chanthaburi | East | Male | 58 | Blood | Acute Febrile Illness ,Viral Infection | 2 | 104 | 104 | epf -/sly+/mrp- |
| 30818 | 2009 | Srakaew | East | Female | NA | Haemoculture | Sepsis | 2 | 1 | 1 | epf +/sly+/mrp+ |
| 30820 | 2009 | Chiang Mai | North | Female | NA | CSF,Heamoculture | Meningitis | 2 | 1 | 1 | epf +/sly+/mrp+ |
| 30821 | 2009 | Chiang Mai | North | Male | NA | Haemoculture | Sepsis | 2 | 1 | 1 | epf +/sly+/mrp+ |
| 30822 | 2009 | Chiang Mai | North | Male | NA | Haemoculture | Sepsis | 2 | 1 | 1 | epf+/sly+/mrp+ |
| 30823 | 2009 | Chiang Mai | North | Male | NA | Haemoculture | Sepsis | 2 | 233 | 233 | epf -/sly+/mrp- |
| 30824 | 2009 | Chiang Mai | North | Male | NA | Haemoculture,CSF | Sepsis, Meningitis | 2 | 1 | 1 | epf +/sly+/mrp+ |
| 30825 | 2009 | Chiang Mai | North | Male | NA | Haemoculture | Sepsis | 2 | 1 | 1 | epf +/sly+/mrp+ |
| 30826 | 2009 | Phrae | North | Male | 59 | Blood , CSF | Meningitis , Acute Febril Illness Septic Shock | 2 | 25 | 103 | epf -/sly-/mrp*** |
| 30827 | 2009 | Phrae | North | Male | 53 | Blood | ARDS | 2 | 1 | 1 | epf +/sly+/mrp+ |
| 30828 | 2009 | Chiang Mai | North | Male | NA | CSF | Meningitis | 2 | 1 | 1 | epf +/sly+/mrp+ |
| 30829 | 2009 | Chiang Mai | North | Male | NA | CSF | Meningitis | 2 | 1 | 1 | epf +/sly+/mrp+ |

| | | | | | | | | | | | |
|---|---|---|---|---|---|---|---|---|---|---|---|
| 30830 | 2009 | Phrae | North | Male | 77 | Blood , CSF | Meningitis | 2 | 1 | 1 | epf +/sly+/mrp+ |
| 30831 | 2009 | Phrae | North | Female | 54 | Blood , CSF | Meningitis , Peripheral Vertigo | 2 | 1 | 1 | epf +/sly+/mrp+ |
| 30832 | 2009 | Chiang Mai | North | Male | NA | NA | NA | 2 | 1 | 1 | epf +/sly+/mrp+ |
| 30833 | 2009 | Phrae | North | Male | 51 | Blood , CSF | Ischemic Stroke | 2 | 1 | 1 | epf +/sly+/mrp+ |
| 30834 | 2009 | Chiang Mai | North | Male | NA | Haemoculture | Sepsis | 2 | 1 | 1 | epf +/sly+/mrp+ |
| 30835 | 2009 | Chiang Mai | North | Male | NA | Haemoculture | Sepsis | 2 | 1 | 1 | epf+/sly+/mrp+ |
| 30836 | 2009 | Phrae | North | Female | 49 | Blood , CSF | Sepsis | 2 | 1 | 1 | epf+ /sly+/mrp+ |
| 30842 | 2009 | Phrae | North | Male | 68 | Blood | Acute Myocardial Infection | 2 | 1 | 1 | epf+ /sly+/mrp+ |
| 30843 | 2009 | Phrae | North | Male | 51 | Blood , CSF | Meningitis , Leptospirosis | 2 | 1 | 1 | epf+ /sly+/mrp+ |
| 31089 | 2009 | Chiang Mai | North | Female | NA | NA | NA | 2 | 1 | 1 | epf+ /sly+/mrp+ |
| 31090 | 2009 | Chiang Mai | North | Female | NA | NA | NA | 2 | 1 | 1 | epf+ /sly+/mrp+ |
| 31091 | 2009 | Chiang Mai | North | Male | NA | NA | NA | 2 | 1 | 1 | epf+ /sly+/mrp+ |
| 31092 | 2009 | Chiang Mai | North | Male | NA | NA | NA | 2 | 1 | 1 | epf+ /sly+/mrp*** |
| 31093 | 2009 | Chiang Rai | North | Male | 56 | Blood | Meningitis | 2 | 1 | 1 | epf+ /sly+/mrp+ |
| 31094 | 2009 | Sukhothai | North | Male | 58 | Blood , CSF | Meningitis | 2 | 1 | 1 | epf+ /sly+/mrp+ |
| 31095 | 2009 | Phetchabul | North | Female | NA | Blood | Sepsis | 2 | 25 | 103 | epf- /sly- /mrp*** |
| 31096 | 2009 | Phetchabul | North | Female | NA | Blood | Sepsis | 2 | 1 | 1 | epf+ /sly+/mrp+ |
| 31098 | 2009 | Uttraradit | North | Male | 52 | Blood | Septic Shock , Acute Renal Failure (ATN) | 2 | 104 | 104 | epf -/sly+/mrp- |
| 31099 | 2009 | Uttraradit | North | Female | 50 | Blood , CSF | Meningitis | 2 | 1 | 1 | epf+ /sly+/mrp+ |
| 31100 | 2009 | Uttraradit | North | Male | 72 | Blood , CSF | Meningitis , Sepsis | 2 | 1 | 1 | epf+ /sly+/mrp+ |
| 31101 | 2009 | Uttraradit | North | Male | 62 | Blood | Septic Shock , Acute infective Diarrhea | 2 | 1 | 1 | epf+ /sly+/mrp+ |
| 31102 | 2009 | Uttraradit | North | Male | NA | Haemoculture | Sepsis | 2 | 1 | 1 | epf+ /sly+/mrp+ |

| | | | | | | | | | | | |
|---|---|---|---|---|---|---|---|---|---|---|---|
| 31106 | 2009 | Phetchabul | North | Male | NA | Blood | Sepsis | 2 | 104 | 104 | epf -/sly+/mrp- |
| 31107 | 2009 | Sukhothai | North | Male | 69 | Blood , CSF | Meningitis | 2 | 1 | 1 | epf+ /sly+/mrp+ |
| 30837 | 2009 | Nakhornratchasima | Northeast | Female | NA | Blood | Sepsis | 2 | 1 | 1 | epf+/sly+/mrp+ |
| 30838 | 2009 | Nakhornratchasima | Northeast | Male | NA | Blood | Sepsis | 2 | 1 | 1 | epf+ /sly+/mrp+ |
| 31278 | 2009 | Nan | North | Male | 53 | Blood | Sepsis , Viral Infection | 2 | 104 | 104 | epf -/sly+/mrp- |
| 31279 | 2009 | Chaiyaphum | Northeast | Male | 36 | Blood | Sepsis , Streptococcus Septicemia | 2 | 1 | 1 | epf+ /sly+/mrp+ |
| 31281 | 2009 | Phetchabul | North | Female | NA | CSF | Meningitis | 2 | 104 | 104 | epf -/sly+/mrp- |
| 34367 | 2010 | Sakon Nakhon | Northeast | Female | 79 | Blood | Sepsis | 2 | 1 | 1 | epf+/sly+/mrp+ |
| 32349 | 2010 | Lampang | North | NA | NA | Blood | Sepsis, Meningitis | 2 | 1 | 1 | epf+/sly+/mrp+ |
| 32490 | 2010 | Lampang | North | NA | NA | Blood | Sepsis | 2 | 1 | 1 | epf+/sly+/mrp+ |
| 32494 | 2010 | Phetchabul | North | NA | NA | Blood | Sepsis | 14 | 1 | 105 | epf+/sly+/mrp+ |
| 32495 | 2010 | Phetchabul | North | NA | NA | Blood | Sepsis, Meningitis | 14 | 1 | 105 | epf+/sly+/mrp+ |
| 32502 | 2010 | Udonthani | Northeast | NA | NA | Blood | Sepsis | 2 | 25 | 103 | epf-/sly-/mrp+ |
| 32514 | 2010 | Phetchabul | North | NA | NA | Blood | Sepsis | 14 | 1 | 105 | epf+/sly+/mrp+ |
| 32516 | 2010 | Phetchabul | North | NA | NA | CSF | Acute Meningitis | 14 | 1 | 105 | epf+/sly+/mrp+ |
| 32517 | 2010 | Phetchabul | North | NA | NA | CSF | Acute Meningitis | 14 | 1 | 105 | epf+/sly+/mrp+ |
| 32518 | 2010 | Phetchabul | North | NA | NA | CSF | Acute Meningitis | 2 | 104 | 104 | epf-/sly+/mrp- |
| 32563 | 2010 | Udonthani | Northeast | NA | NA | Blood | Sepsis, Meningitis | 5 | Singleton | 235 | epf+/sly+/mrp+ |
| 32578 | 2010 | Phetchabul | North | NA | NA | CSF | Acute Meningitis | 2 | 1 | 1 | epf+/sly+/mrp+ |
| 32580 | 2010 | Phetchabul | North | NA | NA | CSF | Acute Meningitis | 2 | 1 | 1 | epf+/sly+/mrp+ |
| 32583 | 2010 | Bhudhasothon | east | NA | NA | Blood | Sepsis, Meningitis | 2 | 104 | 104 | epf-/sly+/mrp- |
| 32588 | 2010 | Nan | North | NA | NA | Blood | Sepsis | 2 | 25 | 380 | epf-/sly-/mrp+ |
| 32599 | 2010 | Burirum | Northeast | NA | NA | Blood | Sepsis | 2 | 1 | 11 | epf-/sly+/mrp+ |
| 32681 | 2010 | Phayao | North | NA | NA | Blood | Sepsis | 14 | 1 | 105 | epf+/sly+/mrp+ |
| 32682 | 2010 | Phayao | North | NA | NA | Blood | Sepsis | 14 | 1 | 105 | epf+/sly+/mrp+ |

| | | | | | | | | | | | |
|---|---|---|---|---|---|---|---|---|---|---|---|
| 32695 | 2010 | Maharajnakhonrajasima | Northeast | NA | NA | Blood | Sepsis | 2 | 104 | 104 | epf-/sly+/mrp- |
| 32709 | 2010 | Nong Khai | Northeast | NA | NA | Blood | Sepsis, Meningitis | 2 | 1 | 1 | epf+/sly+/mrp+ |
| 32710 | 2010 | Nong Khai | Northeast | NA | NA | Blood | Sepsis | 2 | 1 | 1 | epf+/sly+/mrp+ |
| 32711 | 2010 | Maharajnakhonrajasima | Northeast | NA | NA | Blood | Sepsis | 2 | 1 | 1 | epf+/sly+/mrp+ |
| 32776 | 2010 | Taksin | Central | NA | NA | Blood | Sepsis | 2 | 1 | 1 | epf+/sly+/mrp+ |
| 32797 | 2010 | Phetchabul | North | NA | NA | Blood, CSF | Acute Meningitis | 2 | 1 | 1 | epf+/sly+/mrp+ |
| 32798 | 2010 | Phetchabul | North | NA | NA | Blood, CSF | Acute Meningitis | 2 | 104 | 104 | epf-/sly+/mrp- |
| 32801 | 2010 | Phetchabul | North | NA | NA | Blood | Sepsis | 2 | 104 | 104 | epf-/sly+/mrp- |
| 32803 | 2010 | Phetchabul | North | NA | NA | Blood | Sepsis, Meningitis | 2 | 104 | 104 | epf-/sly+/mrp- |
| 32815 | 2010 | Phayao | North | NA | NA | CSF | Acute Meningitis | 2 | 1 | 1 | epf+/sly+/mrp+ |
| 32840 | 2010 | Maharajnakhonrajasima | Northeast | NA | NA | Blood, CSF | Acute Meningitis | 2 | 1 | 1 | epf+/sly+/mrp+ |
| 32860 | 2010 | Phetchabul | North | NA | NA | Blood | Sepsis | 2 | 104 | 104 | epf-/sly+/mrp- |
| 32861 | 2010 | Phetchabul | North | NA | NA | Blood | Sepsis | 2 | 104 | 104 | epf-/sly+/mrp- |
| 32941 | 2010 | Lampang | North | NA | NA | Blood | Sepsis | 14 | 1 | 105 | epf+/sly+/mrp+ |
| 32942 | 2010 | Lampang | North | NA | NA | Blood | Sepsis | 2 | 104 | 104 | epf-/sly+/mrp- |
| 32963 | 2010 | Nakhon Nayok | Central | NA | NA | Blood | Sepsis | 2 | 1 | 1 | epf+/sly+/mrp+ |
| 32970 | 2010 | Phaholponpayuhasena | Central | NA | NA | Blood | Sepsis | 2 | 104 | 104 | epf-/sly+/mrp- |
| 32977 | 2010 | Nan | North | NA | NA | Blood | Sepsis | 2 | 25 | 25 | epf-/sly-/mrp+ |
| 32979 | 2010 | Nan | North | NA | NA | Blood | Sepsis | 2 | 104 | 104 | epf-/sly+/mrp- |
| 32999 | 2010 | Nan | North | NA | NA | Blood | Sepsis | 2 | 104 | 512 | epf-/sly+/mrp- |
| 33004 | 2010 | Lampang | North | NA | NA | Blood | Sepsis | 2 | 104 | 104 | epf-/sly+/mrp- |
| 33011 | 2010 | Phranungklao | Central | NA | NA | Blood | Sepsis | 2 | 1 | 1 | epf+/sly+/mrp+ |
| 33025 | 2010 | Chaophrayayommaraj | Central | NA | NA | Blood | Sepsis, Meningitis | 2 | 1 | 1 | epf+/sly+/mrp+ |
| 33028 | 2010 | Maesod | North | NA | NA | Blood | Sepsis | 2 | 1 | 1 | epf+/sly+/mrp+ |
| 33029 | 2010 | Phetchabul | North | NA | NA | Blood | Sepsis | 2 | 104 | 104 | epf-/sly+/mrp- |
| 33030 | 2010 | Phetchabul | North | NA | NA | Blood | Sepsis | 2 | 1 | 1 | epf+/sly+/mrp+ |

| | | | | | | | | | | | |
|---|---|---|---|---|---|---|---|---|---|---|---|
| 33031 | 2010 | Maesod | North | NA | NA | Blood | Sepsis | 2 | 1 | 1 | epf+/sly+/mrp+ |
| 33038 | 2010 | Phetchabul | North | NA | NA | CSF | Acute Meningitis | 2 | 1 | 1 | epf+/sly+/mrp+ |
| 33052 | 2010 | Phrapokklao | East | NA | NA | blood | Sepsis | 2 | 104 | 104 | epf-/sly+/mrp- |
| 33069 | 2010 | Phetchabul | North | NA | NA | CSF | Acute Meningitis | 2 | 104 | 104 | epf-/sly+/mrp- |
| 33075 | 2010 | Maesod | North | NA | NA | Blood | Sepsis | 2 | 1 | 1 | epf+/sly+/mrp+ |
| 33125 | 2010 | Phetchabul | North | NA | NA | Blood | Sepsis | 2 | 104 | 104 | epf-/sly+/mrp- |
| 33145 | 2010 | Hua Cheao | Central | NA | NA | Blood | Sepsis | 2 | 104 | 104 | epf-/sly+/mrp- |
| 33215 | 2010 | Lampang | North | NA | NA | Blood | Sepsis | 2 | 1 | 1 | epf+/sly+/mrp+ |
| 33243 | 2010 | Udonthani | Northeast | NA | NA | Blood | Sepsis | 2 | 25 | 381 | epf-/sly-/mrp+ |
| 33250 | 2010 | Phayao | North | NA | NA | CSF | Acute Meningitis | 2 | 1 | 1 | epf+/sly+/mrp+ |
| 33251 | 2010 | Phayao | North | NA | NA | Blood | Sepsis | 2 | 104 | 104 | epf-/sly+/mrp- |
| 33273 | 2010 | Phayao | North | NA | NA | Blood | Sepsis, Meningitis | 2 | 1 | 1 | epf+/sly+/mrp+ |
| 33277 | 2010 | Maharajnakhonrajasima | Northeast | NA | NA | Blood | Sepsis | 2 | 1 | 1 | epf+/sly+/mrp+ |
| 33288 | 2010 | Phayao | North | NA | NA | Blood | Sepsis | 2 | 1 | 1 | epf+/sly+/mrp+ |
| 33290 | 2010 | Phetchabul | North | NA | NA | Blood | Sepsis | 2 | 104 | 104 | epf-/sly+/mrp- |
| 33311 | 2010 | Maharajnakhonrajasima | Northeast | NA | NA | Blood | Sepsis | 2 | 1 | 144 | epf+/sly+/mrp+ |
| 33313 | 2010 | Maharajnakhonrajasima | Northeast | NA | NA | Blood | Sepsis | 2 | 1 | 1 | epf+/sly+/mrp+ |
| 33326 | 2010 | Lampang | North | NA | NA | Blood | Sepsis | 2 | 1 | 1 | epf+/sly+/mrp+ |
| 33327 | 2010 | Phetchabul | North | NA | NA | Blood | Sepsis | 2 | 104 | 104 | epf-/sly+/mrp- |
| 33329 | 2010 | Phetchabul | North | NA | NA | Blood | Sepsis | 24 | 221/234 | 221 | epf-/sly-/mrp- |
| 33382 | 2010 | Phetchabul | North | NA | NA | CSF | Acute Meningitis | 2 | 104 | 104 | epf-/sly+/mrp- |
| 33615 | 2010 | Taksin | Central | NA | NA | Blood | Sepsis | 2 | 104 | 104 | epf-/sly+/mrp- |
| 33659 | 2010 | Phetchabul | North | NA | NA | Blood | Sepsis | 2 | 104 | 104 | epf-/sly+/mrp- |
| 33663 | 2010 | Maharajnakhonrajasima | Northeast | NA | NA | CSF | Acute Meningitis | 2 | 1 | 1 | epf+/sly+/mrp+ |
| 33664 | 2010 | Maharajnakhonrajasima | Northeast | NA | NA | CSF | Acute Meningitis | 2 | 1 | 1 | epf+/sly+/mrp+ |
| 33703 | 2010 | Phetchabul | North | NA | NA | CSF | Acute Meningitis | 2 | 104 | 104 | epf-/sly+/mrp- |

| | | | | | | | | | | | |
|---|---|---|---|---|---|---|---|---|---|---|---|
| 33708 | 2010 | Lampang | North | NA | NA | Blood | Sepsis, Meningitis | 2 | 104 | 104 | epf-/sly+/mrp- |
| 33760 | 2010 | Singburi | Central | NA | NA | Blood | Sepsis | 2 | 104 | 104 | epf-/sly+/mrp- |
| 33764 | 2010 | Phetchabul | North | NA | NA | Blood | Sepsis | 2 | 1 | 1 | epf+/sly+/mrp+ |
| 33780 | 2010 | Rayong | East | NA | NA | Blood | Sepsis | 2 | 1 | 1 | epf+/sly+/mrp+ |
| 33802 | 2010 | Burirum | Northeast | Female | 81 | Blood | Sepsis | 2 | 1 | 1 | epf+/sly+/mrp+ |
| 33806 | 2010 | Bhudhachinaraj | North | Female | 45 | Blood | Sepsis | 2 | 1 | 1 | epf+/sly+/mrp+ |
| 33837 | 2010 | Phetchabul | North | Male | 49 | Blood | Sepsis | 2 | 104 | 104 | epf-/sly+/mrp- |
| 33838 | 2010 | Bhudhachinaraj | North | Male | 57 | Blood | Sepsis | 2 | 104 | 104 | epf-/sly+/mrp- |
| 33844 | 2010 | Bhudhachinaraj | North | Male | 60 | Blood | Sepsis | 2 | 1 | 1 | epf+/sly+/mrp+ |
| 33845 | 2010 | Maharajnakhonrajasima | Northeast | Male | 61 | Blood | Sepsis | 2 | 1 | 1 | epf+/sly+/mrp+ |
| 33872 | 2010 | Phayao | North | Male | 36 | Blood | Sepsis | 2 | 25 | 381 | epf-/sly-/mrp+ |
| 33899 | 2010 | Lampang | North | Male | 60 | Blood | Sepsis | 2 | 1 | 1 | epf+/sly+/mrp+ |
| 33900 | 2010 | Lampang | North | Male | 49 | Blood | Sepsis | 2 | 1 | 1 | epf+/sly+/mrp+ |
| 34003 | 2010 | Vajira | Central | Male | 53 | Blood | Sepsis | 2 | 1 | 1 | epf+/sly+/mrp+ |
| 34015 | 2010 | Hua Cheao | Central | Female | 70 | Blood | Sepsis | 2 | 104 | 392 | epf-/sly+/mrp- |
| 34026 | 2010 | Phramonkutklao | Central | Male | NA | Blood | Sepsis | 2 | 1 | 1 | epf+/sly+/mrp+ |
| 34104 | 2010 | Lampang | North | Male | 88 | Blood | Sepsis | 2 | 104 | 104 | epf-/sly+/mrp- |
| 34109 | 2010 | Lampang | North | Male | 49 | Blood | Sepsis | 14 | 1 | 11 | epf-/sly+/mrp+ |
| 34125 | 2010 | Singburi | Central | Male | 61 | Blood | Sepsis | 2 | 1 | 1 | epf+/sly+/mrp+ |
| 34185 | 2010 | Phetchabul | North | Male | 43 | Blood | Sepsis | 2 | 1 | 1 | epf+/sly+/mrp+ |
| 34252 | 2010 | Phayao | North | Male | 41 | Blood | Sepsis | 2 | 1 | 1 | epf+/sly+/mrp+ |
| 34253 | 2010 | Phayao | North | NA | 47 | Blood | Sepsis | 2 | 1 | 1 | epf+/sly+/mrp+ |
| 34803 | 2011 | Lampang | North | Male | 47 | Blood | Sepsis | 2 | 1 | 1 | epf+/sly+/mrp+ |
| 38942 | 2012 | Nakhonping | North | Male | NA | Blood, CSF | Meningitis | 2 | 1 | 1 | epf+/sly+/mrp+ |
| 35074 | 2011 | Bhudhachinaraj | North | Male | 41 | Blood | Sepsis | 2 | 104 | 104 | epf-/sly+/mrp- |
| 35161 | 2011 | Phetchabul | North | Male | 43 | Blood | Sepsis | 2 | 1 | 1 | epf+/sly+/mrp+ |

| | | | | | | | | | | | |
|---|---|---|---|---|---|---|---|---|---|---|---|
| 35357 | 2011 | Sa Kaeo | East | NA | NA | Blood | Sepsis | 2 | 1 | 1 | epf+/sly+/mrp+ |
| 35765 | 2011 | Rayong | East | Female | 37 | Blood | Sepsis | 2 | 104 | 104 | epf-/sly+/mrp- |
| 35767 | 2011 | Burirum | Northeast | Female | 56 | Blood | Sepsis, Meningitis | 2 | 1 | 1 | epf+/sly+/mrp+ |
| 35769 | 2011 | Phetchabul | North | Male | 70 | Blood | Sepsis | 2 | 104 | 104 | epf-/sly+/mrp- |
| 35816 | 2011 | Rayong | East | Male | 36 | Blood | Sepsis | 2 | 104 | 391 | epf-/sly+/mrp- |
| 35823 | 2011 | Pichit | Central | Male | 37 | Blood | Sepsis | 2 | 104 | 104 | epf-/sly+/mrp- |
| 35872 | 2011 | Sa Kaeo | East | Male | 65 | Blood | Sepsis | 2 | 1 | 1 | epf+/sly+/mrp+ |
| 35888 | 2011 | Maharajnakhonrajasima | Northeast | Male | NA | Blood | Sepsis | 2 | 1 | 1 | epf+/sly+/mrp+ |
| 35895 | 2011 | Sa Kaeo | East | Male | 56 | Blood | Sepsis | 2 | 25 | 381 | epf-/sly-/mrp+ |
| 35896 | 2011 | Phetchabul | North | Female | 40 | CSF | Acute Meningitis | 2 | 1 | 105 | epf+/sly+/mrp+ |
| 35897 | 2011 | Phetchabul | North | Male | 47 | Blood | Sepsis | 2 | 1 | 1 | epf+/sly+/mrp+ |
| 35900 | 2011 | Rayong | East | Male | 43 | Blood | Sepsis | 2 | 104 | 104 | epf-/sly+/mrp- |
| 35907 | 2011 | Maesod | North | Female | 47 | Blood | Sepsis | 2 | 1 | 1 | epf+/sly+/mrp+ |
| 35912 | 2011 | Phetchabul | North | Male | 51 | Blood | Sepsis, Meningitis | 2 | 1 | 1 | epf+/sly+/mrp+ |
| 35924 | 2011 | Maharajnakhonrajasima | Northeast | Male | 56 | Blood | Sepsis | 2 | 104 | 104 | epf-/sly+/mrp- |
| 35931 | 2011 | Maharajnakhonrajasima | Northeast | Male | 67 | Blood | Sepsis | 2 | 1 | 1 | epf+/sly+/mrp+ |
| 35940 | 2011 | Phetchabul | North | Male | 21 | Blood | Sepsis | 2 | 233 | 233 | epf-/sly+/mrp- |
| 35986 | 2011 | Phaholponpayuhasena | Central | Male | NA | Blood | Sepsis | 2 | 104 | 104 | epf-/sly+/mrp- |
| 36046 | 2011 | Maharajnakhonrajasima | Northeast | Male | 66 | Blood | Sepsis | 2 | 1 | 1 | epf+/sly+/mrp+ |
| 35928 | 2011 | Singburi | Central | Female | 66 | Blood | Sepsis | 2 | 104 | 104 | epf-/sly+/mrp- |
| 35943 | 2011 | Nakhonping | North | Male | 45 | Blood | Sepsis | 2 | 104 | 104 | epf-/sly+/mrp- |
| 35949 | 2011 | Phrajomklao | Central | Male | 34 | Blood | Sepsis | 2 | 233 | 233 | epf-/sly+/mrp- |
| 35952 | 2011 | Loei | Northeast | Male | 54 | CSF | Acute Meningitis | 2 | 1 | 1 | epf+/sly+/mrp+ |
| 35955 | 2011 | Maharajnakhonrajasima | Northeast | Male | NA | Blood | Sepsis | 2 | 104 | 104 | epf-/sly+/mrp- |
| 35965 | 2011 | Maharajnakhonrajasima | Northeast | Female | 69 | Blood | Sepsis | 2 | 104 | 104 | epf-/sly+/mrp- |
| 35967 | 2011 | Somdejphrachaotaksin | North | Male | 52 | Blood | Sepsis | 2 | 233 | 233 | epf-/sly+/mrp- |

| | | | | | | | | | | | |
|---|---|---|---|---|---|---|---|---|---|---|---|
| 35997 | 2011 | Chiang Rai | North | Male | 34 | Blood | Sepsis | 2 | 1 | 144 | epf+/sly+/mrp+ |
| 35998 | 2011 | Chiang Rai | North | Male | 34 | Blood | Sepsis | 2 | 1 | 1 | epf+/sly+/mrp+ |
| 35999 | 2011 | Chiang Rai | North | Male | 34 | Blood | Sepsis | 2 | 1 | 1 | epf+/sly+/mrp+ |
| 36000 | 2011 | Phetchabul | North | Female | 56 | Blood | Sepsis | 2 | 1 | 144 | epf+/sly+/mrp+ |
| 36004 | 2011 | Pichit | Central | Male | 75 | Blood | Sepsis, Meningitis | 2 | 1 | 1 | epf+/sly+/mrp+ |
| 36005 | 2011 | Pichit | Central | Male | 51 | Blood | Sepsis | 2 | 1 | 1 | epf+/sly+/mrp+ |
| 36031 | 2011 | Maharajnakhonrajasima | Northeast | Male | 61 | Blood | Sepsis | 2 | 1 | 1 | epf+/sly+/mrp+ |
| 36032 | 2011 | Phetchabul | North | Female | 50 | Blood | Sepsis | 2 | 104 | 104 | epf-/sly+/mrp- |
| 36037 | 2011 | Sakon Nakhon | Northeast | Male | 58 | Blood | Sepsis | 2 | 1 | 1 | epf+/sly+/mrp+ |
| 36048 | 2011 | Phetchabul | North | Female | 49 | Blood | Sepsis | 2 | 1 | 1 | epf+/sly+/mrp+ |
| 36053 | 2011 | Lampang | North | Male | 69 | Blood | Sepsis | 2 | 104 | 104 | epf-/sly+/mrp- |
| 36054 | 2011 | Lampang | North | Male | 61 | Blood | Sepsis, Meningitis | 4 | 94 | 94 | epf-/sly+/mrp+ |
| 36067 | 2011 | Maharajnakhonrajasima | Northeast | Male | NA | Blood | Sepsis | 2 | 104 | 104 | epf-/sly+/mrp- |
| 36173 | 2011 | Burirum | Northeast | Male | 73 | Blood | Sepsis | 2 | 104 | 104 | epf-/sly+/mrp- |
| 36178 | 2011 | Phayao | North | Male | NA | CSF | Acute Meningitis | 2 | 1 | 1 | epf+/sly+/mrp+ |
| 36179 | 2011 | Phayao | North | NA | 39 | CSF | Acute Meningitis | 2 | 1 | 1 | epf+/sly+/mrp+ |
| 36193 | 2011 | St. Louis | Central | Male | NA | Blood | Sepsis | 2 | 104 | 104 | epf-/sly+/mrp- |
| 36219 | 2011 | Udonthani | Northeast | Male | 47 | CSF | Acute Meningitis | 2 | 1 | 1 | epf+/sly+/mrp+ |
| 36083 | 2011 | Nong Khai | Northeast | Female | 66 | Blood | Sepsis | 2 | 104 | 104 | epf-/sly+/mrp- |
| 36298 | 2011 | Nakhon Phanom | Northeast | Female | 54 | Blood | Sepsis, Meningitis | 2 | 1 | 1 | epf+/sly+/mrp+ |
| 36303 | 2011 | Nakhon Phanom | Northeast | Male | NA | Blood | Sepsis, Meningitis | 2 | 1 | 1 | epf+/sly+/mrp+ |
| 36315 | 2011 | Nakhon Phanom | Northeast | Female | 64 | Blood | Sepsis | 2 | 1 | 144 | epf+/sly+/mrp+ |
| 36355 | 2011 | Pichit | Central | NA | NA | Blood | Sepsis | 2 | 1 | 144 | epf+/sly+/mrp+ |
| 36363 | 2011 | Nakhon Phanom | Northeast | Male | NA | Blood | Sepsis | 2 | 1 | 144 | epf+/sly+/mrp+ |
| 36367 | 2011 | Chaophrayayommaraj | Central | Male | NA | Blood, CSF | Acute Meningitis | 2 | 1 | 1 | epf+/sly+/mrp+ |
| 36369 | 2011 | Lampang | North | Male | 57 | Blood | Sepsis | 2 | 104 | 512 | epf-/sly+/mrp- |

| | | | | | | | | | | | |
|---|---|---|---|---|---|---|---|---|---|---|---|
| 36378 | 2011 | Phranungklao | Central | Male | NA | Blood | Sepsis | 2 | 1 | 1 | epf+/sly+/mrp+ |
| 36430 | 2011 | Bhudhachinaraj | North | Male | 77 | Blood | Sepsis | 2 | 104 | 104 | epf-/sly+/mrp- |
| 36435 | 2011 | Phetchabul | North | Male | 88 | NA | NA | 2 | 1 | 1 | epf+/sly+/mrp+ |
| 36436 | 2011 | Phetchabul | North | Male | 50 | Blood | Sepsis | 2 | 104 | 104 | epf-/sly+/mrp- |
| 36439 | 2011 | Nakhonping | North | Male | 55 | Blood | Sepsis | 2 | 1 | 1 | epf+/sly+/mrp+ |
| 36440 | 2011 | Lampang | North | Female | 67 | Blood | Sepsis, Meningitis | 2 | 1 | 1 | epf+/sly+/mrp+ |
| 36535 | 2011 | Maharajnakhonrajasima | Northeast | Male | 48 | Blood | Sepsis | 2 | 104 | 104 | epf-/sly+/mrp- |
| 36536 | 2011 | Maharajnakhonrajasima | Northeast | Male | NA | Blood | Sepsis | 2 | 104 | 104 | epf-/sly+/mrp- |
| 36545 | 2011 | Phramonkutklao | Central | Female | NA | Blood | Sepsis | 2 | 104 | 104 | epf-/sly+/mrp- |
| 36563 | 2011 | Phetchabul | North | Male | 65 | NA | NA | 2 | 104 | 104 | epf-/sly+/mrp- |
| 36564 | 2011 | Phetchabul | North | Male | 47 | Blood | Sepsis | 2 | 104 | 104 | epf-/sly+/mrp- |
| 36565 | 2011 | Phetchabul | North | Male | 47 | Blood | Sepsis | 2 | 1 | 1 | epf+/sly+/mrp+ |
| 36566 | 2011 | Phetchabul | North | Female | 92 | Synovial Fluid | Septic Arthritis | 2 | 1 | 1 | epf+/sly+/mrp+ |
| 36569 | 2011 | Sakon Nakhon | Northeast | Male | 49 | CSF | Acute Meningitis | 2 | 1 | 1 | epf+/sly+/mrp+ |
| 36570 | 2011 | Nakhonping | North | Male | 59 | Blood | Sepsis | 2 | 104 | 104 | epf-/sly+/mrp- |
| 36580 | 2011 | Maharajnakhonrajasima | Northeast | Male | 85 | Blood | Sepsis | 2 | 104 | 104 | epf-/sly+/mrp- |
| 36581 | 2011 | Maharajnakhonrajasima | Northeast | Male | 32 | Blood | Sepsis | 2 | 104 | 104 | epf-/sly+/mrp- |
| 36602 | 2011 | Lampang | North | Male | 51 | Body Fluid | NA | 2 | 233 | 379 | epf-/sly+/mrp- |
| 36610 | 2011 | Nakhonping | North | Male | 48 | Blood | Sepsis | 2 | 104 | 391 | epf-/sly+/mrp- |
| 36620 | 2011 | Nakhon Phanom | Northeast | Male | 42 | Blood | Sepsis | 2 | 1 | 1 | epf+/sly+/mrp+ |
| 36639 | 2011 | Sakon Nakhon | Northeast | Male | 70 | Blood | Sepsis | 2 | 1 | 1 | epf+/sly+/mrp+ |
| 36716 | 2011 | Lampang | North | Male | 47 | CSF | Acute Meningitis | 2 | 1 | 1 | epf+/sly+/mrp+ |
| 36717 | 2011 | Phetchabul | North | Male | 54 | Blood | Sepsis | 2 | 1 | 1 | epf+/sly+/mrp+ |
| 36323 | 2011 | Udonthani | Northeast | Male | 67 | Blood | Sepsis | 2 | 104 | 391 | epf-/sly+/mrp- |
| 37438 | 2011 | Lampang | North | Male | 51 | Blood | Sepsis | 2 | 104 | 104 | epf-/sly+/mrp- |
| 36375 | 2011 | Nong Khai | Northeast | Male | 46 | Blood | Sepsis | 2 | 25 | 395 | epf-/sly-/mrp+ |

| | | | | | | | | | | | |
|---|---|---|---|---|---|---|---|---|---|---|---|
| 38825 | 2011 | Pichit | Central | Male | 44 | Blood | Sepsis | 2 | 104 | 104 | epf-/sly+/mrp- |
| 36738 | 2011 | Udonthani | Northeast | Male | 61 | Blood | Sepsis | 2 | 25 | 380 | epf-/sly-/mrp+ |
| 36762 | 2011 | Uttaradit | North | Male | 44 | Blood | Sepsis | 2 | 1 | 1 | epf+/sly+/mrp+ |
| 36763 | 2011 | Uttaradit | North | Male | NA | Blood | Sepsis | NT | 28 | 28 | epf-/sly-/mrp+ |
| 36829 | 2011 | Taksin | Central | Male | 62 | Blood | Sepsis | 2 | 104 | 104 | epf-/sly+/mrp- |
| 36906 | 2011 | Chaophrayayommaraj | Central | Male | NA | Blood | Sepsis, Meningitis | 2 | 104 | 104 | epf-/sly+/mrp- |
| 36921 | 2011 | Chaophrayayommaraj | Central | Male | NA | Blood | Sepsis | 2 | 104 | 104 | epf-/sly+/mrp- |
| 37656 | 2011 | Maharajnakhonrajasima | Northeast | Male | 64 | Blood | Sepsis, Meningitis | 2 | 1 | 1 | epf+/sly+/mrp+ |
| 38291 | 2011 | Pichit | Central | Male | 55 | Blood | Sepsis | 2 | 104 | 104 | epf-/sly+/mrp- |
| 37083 | 2011 | Udonthani | Northeast | Male | 36 | Blood | Sepsis | 2 | 104 | 104 | epf-/sly+/mrp- |
| 37646 | 2011 | Lampang | North | Female | 47 | Blood | Sepsis | 2 | 1 | 1 | epf+/sly+/mrp+ |
| 38782 | 2011 | Lampang | North | Male | 56 | Blood | Sepsis | 2 | 104 | 104 | epf-/sly+/mrp- |
| 37084 | 2011 | Uttaradit | North | Male | 59 | Blood | Sepsis | 2 | 104 | 104 | epf-/sly+/mrp- |
| 37549 | 2011 | Phetchabul | North | Female | 62 | CSF | Acute Meningitis | 2 | 1 | 1 | epf+/sly+/mrp+ |
| 38055 | 2011 | Uttaradit | North | Male | 62 | Blood | Sepsis | 2 | 1 | 1 | epf+/sly+/mrp+ |
| 37647 | 2011 | Lampang | North | Female | 46 | Blood | Sepsis | 2 | 233 | 233 | epf-/sly+/mrp- |
| 37548 | 2011 | Nakhonping | North | Male | 46 | Blood | Sepsis | 2 | 1 | 1 | epf+/sly+/mrp+ |
| 37554 | 2011 | Nakhonping | North | Female | 80 | Blood | Sepsis | 2 | 233 | 233 | epf-/sly+/mrp- |
| 37622 | 2011 | Nakhonping | North | Male | 55 | Blood | Sepsis | 2 | 104 | 104 | epf-/sly+/mrp- |
| 37879 | 2011 | Nan | North | Male | 55 | Blood | Sepsis | 2 | 104 | 104 | epf-/sly+/mrp- |
| 37945 | 2011 | Maharajnakhonrajasima | Northeast | Male | 79 | Blood | Sepsis | 2 | 1 | 1 | epf+/sly+/mrp+ |
| 37675 | 2011 | Uttaradit | North | Male | 47 | Blood | Sepsis | 2 | 1 | 1 | epf+/sly+/mrp+ |
| 37833 | 2011 | Bhudhachinaraj | North | Male | 77 | Blood | Sepsis | 2 | 104 | 104 | epf-/sly+/mrp- |
| 37902 | 2011 | Phetchabul | North | Male | 51 | Synovial Fluid | Septic Arthritis | 2 | 1 | 1 | epf+/sly+/mrp+ |
| 38172 | 2011 | Maharajnakhonrajasima | Northeast | Female | 75 | CSF | Acute Meningitis | 2 | 1 | 1 | epf+/sly+/mrp+ |
| 38512 | 2011 | Phetchabul | North | Male | 49 | Blood | Sepsis | 2 | 104 | 104 | epf-/sly+/mrp- |

| | | | | | | | | | | | |
|---|---|---|---|---|---|---|---|---|---|---|---|
| 37880 | 2011 | Lampang | North | Male | 38 | Blood | Sepsis | 2 | 233 | 379 | epf-/sly+/mrp- |
| 38433 | 2011 | Burirum | Northeast | Female | 52 | Blood | Sepsis | 2 | 25 | 380 | epf-/sly-/mrp+ |
| 38824 | 2011 | Phetchabul | North | Male | 35 | Blood | Sepsis | 14 | 1 | 105 | epf+/sly+/mrp+ |
| 38702 | 2011 | Udonthani | Northeast | Male | 43 | Blood | Sepsis | 2 | 1 | 1 | epf+/sly+/mrp+ |
| 38704 | 2011 | Phetchabul | North | Male | 48 | Blood | Sepsis | 2 | 104 | 104 | epf-/sly+/mrp- |
| 38245 | 2011 | Maharajnakhonrajasima | Northeast | Female | 69 | Blood | Sepsis | 2 | 104 | 104 | epf-/sly+/mrp- |
| 38727 | 2011 | Lampang | North | Male | 46 | Blood | Sepsis | 2 | 233 | 233 | epf-/sly+/mrp- |
| 38721 | 2011 | Nan | North | Male | 50 | Blood | Sepsis | 2 | 104 | 104 | epf-/sly+/mrp- |
| 38732 | 2011 | Nan | North | Male | 85 | Blood | Sepsis | 2 | 104 | 514 | epf-/sly+/mrp- |
| 38175 | 2011 | Saraburi | Central | Female | 35 | Blood | Sepsis | 2 | 104 | 104 | epf-/sly+/mrp- |
| 38826 | 2011 | Phetchabul | North | Male | 72 | Blood | Sepsis | 14 | 1 | 105 | epf+/sly+/mrp+ |
| 38827 | 2011 | Nakhon Phanom | Northeast | Male | 48 | Blood | Sepsis | 2 | 1 | 1 | epf+/sly+/mrp+ |
| 38828 | 2011 | Phrapokklao | East | Male | 63 | Blood | Sepsis | 14 | 1 | 105 | epf+/sly+/mrp+ |
| 38521 | 2011 | Lampang | North | Male | 47 | Blood | Sepsis | 14 | 1 | 105 | epf+/sly+/mrp+ |
| 41442 | 2012 | Nakhon Phanom | Northeast | Male | 48 | Blood | Sepsis | 2 | 1 | 1 | epf+/sly+/mrp+ |
| 38728 | 2012 | Sakon Nakhon | Northeast | Male | 43 | Blood | Sepsis, Meningitis | 2 | 1 | 1 | epf+/sly+/mrp+ |
| 39619 | 2012 | Uttaradit | North | Male | 66 | Blood | Sepsis | 2 | 1 | 1 | epf+/sly+/mrp+ |
| 41440 | 2012 | Sa Kaeo | East | Male | 54 | Blood | Sepsis | 2 | 1 | 1 | epf+/sly+/mrp+ |
| 38730 | 2012 | Maharajnakhonrajasima | Northeast | Male | 60 | Blood | Sepsis | 2 | 104 | 391 | epf-/sly+/mrp- |
| 38985 | 2012 | Maharajnakhonrajasima | Northeast | Male | 77 | Blood | Sepsis | 2 | 1 | 1 | epf+/sly+/mrp+ |
| 38778 | 2012 | Saraburi | Central | Male | 46 | Blood | Sepsis | 2 | 1 | 1 | epf+/sly+/mrp+ |
| 38939 | 2012 | Hat Yai | South | Male | 53 | Blood | Sepsis | 2 | 104 | 104 | epf-/sly+/mrp- |
| 39497 | 2012 | Lampang | North | Female | 47 | Blood | Sepsis | 2 | 1 | 1 | epf+/sly+/mrp+ |
| 38790 | 2012 | Saraburi | Central | Female | 45 | Blood | Sepsis | 2 | 1 | 1 | epf+/sly+/mrp+ |
| 38937 | 2012 | Saraburi | Central | Male | 44 | Blood | Sepsis, Meningitis | 2 | 1 | 1 | epf+/sly+/mrp+ |
| 39558 | 2012 | Uttaradit | North | Female | 68 | Blood | Sepsis | 2 | 1 | 1 | epf+/sly+/mrp+ |

| | | | | | | | | | | | |
|---|---|---|---|---|---|---|---|---|---|---|---|
| 39014 | 2012 | Phetchabul | North | Male | 58 | CSF | Meningitis | 2 | 1 | 1 | epf+/sly+/mrp+ |
| 39265 | 2012 | Hat Yai | South | Male | 66 | Blood | Sepsis | 2 | 1 | 1 | epf+/sly+/mrp+ |
| 39104 | 2012 | Khon Kaen | Northeast | Male | 56 | Blood | Sepsis | 2 | 25 | 380 | epf-/sly-/mrp+ |
| 39559 | 2012 | Uttaradit | North | Male | 48 | Blood | Sepsis | 2 | 1 | 1 | epf+/sly+/mrp+ |
| 39587 | 2012 | Uttaradit | North | Male | 43 | Blood | Sepsis | 2 | 1 | 1 | epf+/sly+/mrp+ |
| 39151 | 2012 | Sakon Nakhon | Northeast | Male | 55 | Blood | Sepsis | 2 | 1 | 1 | epf+/sly+/mrp+ |
| 39152 | 2012 | Khon Kaen | Northeast | Male | 44 | Blood | Sepsis | 2 | 1 | 1 | epf+/sly+/mrp+ |
| 39232 | 2012 | Bhudhachinaraj | North | Female | NA | Blood | Sepsis | 2 | 1 | 1 | epf+/sly+/mrp+ |
| 39234 | 2012 | Bhudhachinaraj | North | Female | NA | Blood | Sepsis, Meningitis | 2 | 1 | 1 | epf+/sly+/mrp+ |
| 39481 | 2012 | Nakhon Phanom | Northeast | Female | 39 | Blood | Sepsis | 2 | 1 | 1 | epf+/sly+/mrp+ |
| 39162 | 2012 | Maharajnakhonrajasima | Northeast | Male | 72 | Blood | Sepsis | 2 | 104 | 104 | epf-/sly+/mrp- |
| 39209 | 2012 | Phetchabul | North | Male | 27 | CSF | Meningitis | 2 | 1 | 1 | epf+/sly+/mrp+ |
| 41470 | 2012 | Maharajnakhonrajasima | Northeast | Male | 74 | Blood | Sepsis, Meningitis | 2 | 1 | 1 | epf+/sly+/mrp+ |
| 39492 | 2012 | Nakhon Phanom | Northeast | Male | 41 | Blood | Sepsis | 2 | 1 | 1 | epf+/sly+/mrp+ |
| 39544 | 2012 | Lampang | North | Male | 74 | CSF | Meningitis | 2 | 1 | 1 | epf+/sly+/mrp+ |
| 39249 | 2012 | Maharajnakhonrajasima | Northeast | Male | 78 | Blood | Sepsis | 2 | 1 | 1 | epf+/sly+/mrp+ |
| 39479 | 2012 | Maharajnakhonrajasima | Northeast | Male | 34 | Blood | Sepsis | 2 | 1 | 105 | epf+/sly+/mrp+ |
| 39480 | 2012 | Maharajnakhonrajasima | Northeast | Male | 53 | Blood | Sepsis | 2 | 25 | 380 | epf-/sly-/mrp+ |
| 39236 | 2012 | Bhudhachinaraj | North | Male | NA | Blood | Sepsis | 2 | 233 | 233 | epf-/sly+/mrp- |
| 39235 | 2012 | Bhudhachinaraj | North | Male | NA | Blood | Sepsis | 2 | 1 | 1 | epf+/sly+/mrp+ |
| 39233 | 2012 | Bhudhachinaraj | North | Male | NA | Blood | Sepsis | 2 | 104 | 104 | epf-/sly+/mrp- |
| 39258 | 2012 | Lampang | North | Male | 32 | Blood | Sepsis | 2 | 1 | 1 | epf+/sly+/mrp+ |
| 40010 | 2012 | Hua Cheao | Central | Female | 41 | Blood | Sepsis, Meningitis | 2 | 104 | 104 | epf-/sly+/mrp- |
| 41471 | 2012 | Lampang | North | Male | 74 | Blood | Sepsis, Meningitis | 2 | 1 | 1 | epfs/sly+/mrp+ |
| 39509 | 2012 | Phetchabul | North | Male | 53 | Blood | Sepsis | 2 | 1 | 1 | epf+/sly+/mrp+ |
| 39499 | 2012 | Lampang | North | Male | 68 | Blood | Sepsis | 2 | 1 | 1 | epf+/sly+/mrp+ |

| | | | | | | | | | | | |
|---|---|---|---|---|---|---|---|---|---|---|---|
| 39547 | 2012 | Bhudhachinaraj | North | Male | 72 | Blood | Sepsis | 2 | 104 | 104 | epf-/sly+/mrp- |
| 39502 | 2012 | Nakhon Phanom | Northeast | Male | 50 | Blood | Sepsis | 2 | 1 | 337 | epf+/sly+/mrp+ |
| 39542 | 2012 | Saraburi | Central | Male | 80 | Blood | Sepsis | 2 | 1 | 1 | epf+/sly+/mrp+ |
| 39548 | 2012 | Bhudhachinaraj | North | Male | 29 | Blood | Sepsis | 2 | 104 | 104 | epf-/sly+/mrp- |
| 39557 | 2012 | Uttaradit | North | Female | 55 | Blood | Sepsis | 2 | 104 | 104 | epf-/sly+/mrp- |
| 39504 | 2012 | Saraburi | Central | Male | 61 | Blood | Sepsis | 2 | 1 | 1 | epf+/sly+/mrp+ |
| 39957 | 2012 | Uttaradit | North | Male | 40 | Blood | Sepsis | 14 | 1 | 105 | epf+/sly+/mrp+ |
| 39554 | 2012 | Saraburi | Central | Female | 79 | Blood | Sepsis, Meningitis | 2 | 1 | 1 | epf+/sly+/mrp+ |
| 40082 | 2012 | Taksin | Central | Male | 61 | Blood | Sepsis | 2 | 104 | 104 | epf-/sly+/mrp- |
| 39603 | 2012 | Saraburi | Central | Male | 59 | Blood | Sepsis | 2 | 1 | 1 | epf+/sly+/mrp+ |
| 39778 | 2012 | Nakhon Phanom | Northeast | Male | 28 | Blood | Sepsis | 2 | 1 | 1 | epf+/sly+/mrp+ |
| 39969 | 2012 | Nan | North | Male | 70 | Blood | Sepsis, Meningitis | 2 | 104 | 104 | epf-/sly+/mrp- |
| 39764 | 2012 | Bhudhachinaraj | North | Male | 51 | Blood | Sepsis | 2 | 1 | 1 | epf+/sly+/mrp+ |
| 39950 | 2012 | Phetchabul | North | Male | 69 | Blood | Sepsis | 2 | 1 | 1 | epf+/sly+/mrp+ |
| 39424 | 2012 | Maharajnakhonrajasima | Northeast | Male | 42 | Blood | Sepsis | 2 | 1 | 1 | epf+/sly+/mrp+ |
| 39729 | 2012 | Saraburi | Central | NA | NA | Blood | Sepsis | 2 | 104 | 104 | epf-/sly+/mrp- |
| 39891 | 2012 | Phetchabul | North | Male | 39 | Blood | Sepsis | 2 | 104 | 104 | epf-/sly+/mrp- |
| 39970 | 2012 | Nan | North | Male | 40 | Blood | Sepsis, Meningitis | 2 | 104 | 104 | epf-/sly+/mrp- |
| 39824 | 2012 | Phetchabul | North | Male | 47 | Blood | Sepsis | 14 | 1 | 105 | epf+/sly+/mrp+ |
| 40150 | 2012 | Phayao | North | Male | 51 | Blood | Sepsis, Meningitis | 2 | 1 | 1 | epf+/sly+/mrp+ |
| 39720 | 2012 | Bhudhachinaraj | North | Male | 33 | Blood | Sepsis, Meningitis | 2 | 1 | 1 | epf+/sly+/mrp+ |
| 39825 | 2012 | Phetchabul | North | Female | 74 | CSF | Meningitis | 14 | 1 | 105 | epf+/sly+/mrp+ |
| 39830 | 2012 | Nakhon Phanom | Northeast | Male | 46 | Blood | Sepsis | 2 | 1 | 1 | epf+/sly+/mrp+ |
| 39971 | 2012 | Nan | North | Male | 34 | Blood | Sepsis | 2 | 104 | 512 | epf-/sly+/mrp- |
| 40011 | 2012 | Phetchabul | North | Male | 91 | Blood | Sepsis | 2 | 1 | 1 | epf+/sly+/mrp+ |
| 40133 | 2012 | Bhudhachinaraj | North | Male | 57 | Blood | Sepsis | 2 | 104 | 513 | epf-/sly+/mrp- |

| | | | | | | | | | | | |
|---|---|---|---|---|---|---|---|---|---|---|---|
| 39845 | 2012 | Saraburi | Central | Male | 48 | Blood | Sepsis | 2 | 104 | 104 | epf-/sly+/mrp- |
| 39846 | 2012 | Saraburi | Central | Male | 37 | Blood | Sepsis | 2 | 25 | 25 | epf-/sly-/mrp+ |
| 39827 | 2012 | Phetchabul | North | Male | 71 | CSF | Meningitis | 14 | 1 | 105 | epf+/sly+/mrp+ |
| 39843 | 2012 | Saraburi | Central | Male | 65 | Blood | Sepsis | 2 | 25 | 25 | epf-/sly-/mrp+ |
| 39931 | 2012 | Nakhonping | North | Male | NA | Blood | Sepsis | 2 | 1 | 1 | epf+/sly+/mrp+ |
| 39838 | 2012 | Nakhonping | North | Male | NA | Blood | Sepsis | 2 | 104 | 104 | epf-/sly+/mrp- |
| 39835 | 2012 | Chaophrayayommaraj | Central | Male | NA | Blood | Sepsis | 2 | 1 | 1 | epf+/sly+/mrp+ |
| 39895 | 2012 | Maharajnakhonrajasima | Northeast | Female | 71 | Blood | Sepsis | 2 | 1 | 1 | epf+/sly+/mrp+ |
| 39958 | 2012 | Uttaradit | North | Male | 62 | Blood | Sepsis | 2 | 1 | 1 | epf+/sly+/mrp+ |
| 39966 | 2012 | Nakhon Phanom | Northeast | Female | 48 | Blood | Sepsis | 2 | 1 | 1 | epf+/sly+/mrp+ |
| 39973 | 2012 | Nan | North | Male | 66 | Blood | Sepsis | 2 | 104 | 104 | epf-/sly+/mrp- |
| 40163 | 2012 | Sakon Nakhon | Northeast | Male | 40 | CSF | Meningitis | 2 | 1 | 1 | epf+/sly+/mrp+ |
| 40424 | 2012 | Chaophrayayommaraj | Central | Male | NA | Blood | Sepsis | 2 | 25 | 380 | epf-/sly-/mrp+ |
| 40483 | 2012 | Chaophrayayommaraj | Central | Male | NA | Blood | Sepsis | 2 | 104 | 104 | epf-/sly+/mrp- |
| 40753 | 2012 | Maharajnakhonrajasima | Northeast | Male | 30 | Synovial Fluid | Septic Arthritis | 2 | 1 | 1 | epf+/sly+/mrp+ |
| 39949 | 2012 | Taksin | Central | Male | 69 | Blood | Sepsis | 2 | 25 | 25 | epf-/sly-/mrp+ |
| 39959 | 2012 | Uttaradit | North | Male | 50 | Blood | Sepsis | 2 | 104 | 391 | epf-/sly+/mrp- |
| 39990 | 2012 | Phetchabul | North | Female | 38 | CSF | Meningitis | 2 | 1 | 1 | epf+/sly+/mrp+ |
| 39848 | 2012 | Saraburi | Central | Male | 50 | Blood | Sepsis | 2 | 1 | 1 | epf+/sly+/mrp+ |
| 39960 | 2012 | Uttaradit | North | Male | 56 | Blood | Sepsis | 2 | 104 | 104 | epf-/sly+/mrp- |
| 39847 | 2012 | Saraburi | Central | Male | 72 | Blood | Sepsis, Meningitis | 2 | 1 | 1 | epf+/sly+/mrp+ |
| 39855 | 2012 | Maharajnakhonrajasima | Northeast | Male | NA | Blood | Sepsis | 2 | 1 | 1 | epf+/sly+/mrp+ |
| 39974 | 2012 | Nan | North | Male | 46 | Blood | Sepsis | 2 | 104 | 392 | epf-/sly+/mrp- |
| 40014 | 2012 | Uttaradit | North | Male | 70 | Blood | Sepsis | 2 | 104 | 391 | epf-/sly+/mrp- |
| 40517 | 2012 | Lampang | North | Male | NA | Blood | Sepsis | 2 | 1 | 1 | epf+/sly+/mrp+ |
| 39980 | 2012 | Saraburi | Central | Female | 37 | Blood | Sepsis, Meningitis | 2 | 1 | 1 | epf+/sly+/mrp+ |

| | | | | | | | | | | | |
|---|---|---|---|---|---|---|---|---|---|---|---|
| 40016 | 2012 | Phetchabul | North | Male | 29 | CSF | Meningitis | 2 | 1 | 1 | epf+/sly+/mrp+ |
| 39982 | 2012 | Saraburi | Central | Male | 48 | Blood | Sepsis | 2 | 1 | 1 | epf+/sly+/mrp+ |
| 39983 | 2012 | Saraburi | Central | Female | 42 | Blood | Sepsis | 2 | 1 | 1 | epf+/sly+/mrp+ |
| 40128 | 2012 | Nakhon Phanom | Northeast | Male | 23 | Blood | Sepsis | 2 | 1 | 337 | epf+/sly+/mrp+ |
| 40137 | 2012 | Bhudhachinaraj | North | Male | 68 | Blood | Sepsis | 2 | 104 | 104 | epf-/sly+/mrp- |
| 40135 | 2012 | Bhudhachinaraj | North | Male | 64 | Blood | Sepsis | 2 | 104 | 104 | epf-/sly+/mrp- |
| 40142 | 2012 | Bhudhachinaraj | North | Male | 43 | Blood | Sepsis | 2 | 104 | 104 | epf-/sly+/mrp- |
| 40146 | 2012 | Saraburi | Central | NA | 89 | Blood, CSF | Meningitis | 2 | 1 | 1 | epf+/sly+/mrp+ |
| 40414 | 2012 | Bhudhachinaraj | North | Male | NA | Blood | Sepsis | 2 | 233 | 233 | epf-/sly+/mrp- |
| 40154 | 2012 | Sakon Nakhon | Northeast | Female | 51 | CSF | Meningitis | 2 | 1 | 1 | epf+/sly+/mrp+ |
| 40465 | 2012 | St. Louis | Central | Male | 41 | Blood, CSF | Meningitis | 2 | 1 | 1 | epfs/sly+/mrp+ |
| 40240 | 2012 | Saraburi | Central | Female | 77 | Blood | Sepsis | 2 | 25 | 103 | epf-/sly-/mrp+ |
| 40305 | 2012 | Saraburi | Central | Male | 75 | Blood | Sepsis | 2 | 104 | 104 | epf-/sly+/mrp- |
| 40339 | 2012 | Nakhon Phanom | Northeast | Male | 48 | Blood | Sepsis, Meningitis | 2 | 1 | 1 | epf+/sly+/mrp+ |
| 40679 | 2012 | Phayao | North | Male | NA | Blood | Sepsis | 2 | 1 | 1 | epf+/sly+/mrp+ |
| 40363 | 2012 | Maharajnakhonrajasima | Northeast | Male | 59 | Blood | Sepsis | 2 | 104 | 104 | epf-/sly+/mrp- |
| 40678 | 2012 | Phayao | North | Male | NA | Blood | Sepsis | 2 | 104 | 104 | epf-/sly+/mrp- |
| 40411 | 2012 | Uttaradit | North | Male | 62 | Blood | Sepsis | 2 | 104 | 104 | epf-/sly+/mrp- |
| 40414 | 2012 | Saraburi | Central | Male | 50 | Blood | Sepsis | 2 | 104 | 514 | epf-/sly+/mrp- |
| 40415 | 2012 | Saraburi | Central | Female | 75 | CSF | Meningitis | 2 | 1 | 1 | epf+/sly+/mrp+ |
| 40417 | 2012 | Saraburi | Central | Male | 34 | Blood | Sepsis | 2 | 25 | 25 | epf-/sly-/mrp+ |
| 40236 | 2012 | Bhudhachinaraj | North | Male | 42 | Blood | Sepsis | 2 | 104 | 104 | epf-/sly+/mrp- |
| 40237 | 2012 | Bhudhachinaraj | North | Male | 79 | Blood | Sepsis | 2 | 233 | 233 | epf-/sly+/mrp- |
| 40300 | 2012 | Maesod | North | Male | 41 | Blood | Sepsis | 2 | 1 | 1 | epf+/sly+/mrp+ |
| 40301 | 2012 | Bhudhachinaraj | North | Male | 60 | Blood | Sepsis | 2 | 233 | 233 | epf-/sly+/mrp- |
| 40302 | 2012 | Bhudhachinaraj | North | Male | 68 | Blood | Sepsis | 2 | 104 | 393 | epf-/sly+/mrp- |

| | | | | | | | | | | | |
|---|---|---|---|---|---|---|---|---|---|---|---|
| 40410 | 2012 | Nakhonping | North | Male | NA | Blood | Sepsis | 2 | 104 | 392 | epf-/sly+/mrp- |
| 40484 | 2012 | Bhudhachinaraj | North | Male | 36 | CSF | Meningitis | 14 | 1 | 105 | epf+/sly+/mrp+ |
| 40707 | 2012 | Phetchabul | North | Female | 54 | Blood | Sepsis | 2 | 1 | 1 | epf+/sly+/mrp+ |
| 40749 | 2012 | Bhudhachinaraj | North | Female | 68 | Blood | Sepsis | 2 | 1 | 337 | epf+/sly+/mrp+ |
| 40750 | 2012 | Bhudhachinaraj | North | Female | 65 | Blood | Sepsis | 2 | 233 | 233 | epf-/sly+/mrp- |
| 40803 | 2012 | Phranungklao | Central | Female | 56 | Blood | Sepsis | 2 | 233 | 233 | epf-/sly+/mrp- |
| 40804 | 2012 | Phranungklao | Central | Female | 79 | Blood | Sepsis | 2 | 233 | 233 | epf-/sly+/mrp- |
| 40486 | 2012 | Uttaradit | North | Male | 69 | Blood | Sepsis | 2 | 104 | 104 | epf-/sly+/mrp- |
| 40364 | 2012 | Maharajnakhonrajasima | Northeast | Male | NA | Blood | Sepsis | 2 | 233 | 233 | epf-/sly+/mrp- |
| 40412 | 2012 | Uttaradit | North | Male | 53 | Blood | Sepsis | 2 | 104 | 104 | epf-/sly+/mrp- |
| 40586 | 2012 | Nakhon Phanom | Northeast | Male | 64 | Blood | Sepsis | 2 | 1 | 1 | epf+/sly+/mrp+ |
| 40453 | 2012 | Maharajnakhonrajasima | Northeast | Male | 45 | Blood | Sepsis | 2 | 104 | 104 | epf-/sly+/mrp- |
| 40511 | 2012 | Sakon Nakhon | Northeast | Male | 64 | Blood | Sepsis, Meningitis | 2 | 1 | 1 | epf+/sly+/mrp+ |
| 40672 | 2012 | Nakhon Phanom | Northeast | Male | 47 | Blood | Sepsis | 2 | 1 | 1 | epf+/sly+/mrp+ |
| 40744 | 2012 | Bhudhachinaraj | North | Male | 71 | Blood | Sepsis | 2 | 1 | 1 | epf+/sly+/mrp+ |
| 40614 | 2012 | Burirum | Northeast | Male | 64 | Blood | Sepsis | 2 | 1 | 1 | epf+/sly+/mrp+ |
| 40621 | 2012 | Bhudhasothon | East | Male | 56 | Blood | Sepsis | 2 | 104 | 392 | epf-/sly+/mrp- |
| 40585 | 2012 | Sakon Nakhon | Northeast | Male | 43 | Blood | Sepsis | 2 | 1 | 1 | epf+/sly+/mrp+ |
| 40625 | 2012 | Maharajnakhonrajasima | Northeast | Male | 75 | Blood | Sepsis | 2 | 1 | 1 | epf+/sly+/mrp+ |
| 40623 | 2012 | Phetchabul | North | Male | 53 | Blood | Sepsis, Meningitis | 2 | 1 | 1 | epf+/sly+/mrp+ |
| 40746 | 2012 | Bhudhachinaraj | North | Male | 36 | Blood | Sepsis | 2 | 104 | 104 | epf-/sly+/mrp- |
| 40747 | 2012 | Bhudhachinaraj | North | Male | 27 | Blood | Sepsis | 2 | 233 | 233 | epf-/sly+/mrp- |
| 40752 | 2012 | Maharajnakhonrajasima | Northeast | Male | 46 | Blood | Sepsis | 2 | 1 | 1 | epf+/sly+/mrp+ |
| 41112 | 2012 | Phramonkutklao | Central | Male | 39 | Blood | Sepsis | 2 | 104 | 391 | epf-/sly+/mrp- |
| 41121 | 2012 | Udonthani | Northeast | Male | 40 | CSF | Meningitis | 2 | 233 | 233 | epf-/sly+/mrp- |
| 41218 | 2012 | Lampang | North | Female | 47 | Blood | Sepsis | 2 | 233 | 233 | epf-/sly+/mrp- |

| | | | | | | | | | | | |
|---|---|---|---|---|---|---|---|---|---|---|---|
| 41234 | 2012 | Nakhon Phanom | Northeast | Female | 62 | Blood | Sepsis | 2 | 1 | 337 | epf+/sly+/mrp+ |
| 41117 | 2012 | Nan | North | NA | NA | Blood | Sepsis | 2 | 1 | 1 | epf+/sly+/mrp+ |
| 40833 | 2012 | Phetchabul | North | Male | 41 | Blood | Sepsis | 2 | 104 | 104 | epf-/sly+/mrp- |
| 41087 | 2012 | Phayao | North | Male | 63 | Blood | Sepsis | 2 | 1 | 1 | epf+/sly+/mrp+ |
| 41122 | 2012 | Phetchabul | North | Male | 88 | Blood | Sepsis | 2 | 1 | 1 | epf+/sly+/mrp+ |
| 41123 | 2012 | Phetchabul | North | Male | 59 | Blood | Sepsis | 2 | 104 | 104 | epf-/sly+/mrp- |
| 41115 | 2012 | Saraburi | Central | Male | 77 | Blood | Sepsis | 2 | 104 | 104 | epf-/sly+/mrp- |
| 41126 | 2012 | Maharajnakhonrajasima | Northeast | Male | NA | CSF | Meningitis | 2 | 233 | 233 | epf-/sly+/mrp- |
| 41106 | 2012 | Roi Et | Northeast | Male | 56 | Blood | Sepsis | 2 | 104 | 104 | epf-/sly+/mrp- |
| 41120 | 2012 | Nan | North | Male | 50 | Blood | Sepsis | 2 | 104 | 104 | epf-/sly+/mrp- |
| 41594 | 2012 | Phetchabul | North | Male | 66 | Blood | Sepsis | 2 | Singleton | 236 | epf-/sly+/mrp+ |
| 41219 | 2012 | Sa Kaeo | East | Male | 48 | Blood | Sepsis | 2 | 233 | 233 | epf-/sly+/mrp- |
| 41477 | 2012 | Nan | North | Male | 73 | Blood | Sepsis | 2 | 233 | 233 | epf-/sly+/mrp- |
| 41236 | 2012 | Nakhon Phanom | Northeast | Male | 60 | Blood | Sepsis, Meningitis | 2 | 1 | 1 | epf+/sly+/mrp+ |
| 41570 | 2012 | Phayao | North | Male | 61 | Blood | Sepsis | 5 | 221/234 | 221 | epf+/sly+/mrp+ |
| 41476 | 2012 | Nan | North | Male | 59 | Blood | Sepsis, Meningitis | 2 | 104 | 104 | epf-/sly+/mrp- |
| 41478 | 2012 | Nan | North | Male | 70 | Blood | Sepsis | 2 | 104 | 104 | epf-/sly+/mrp- |
| 41306 | 2012 | Saraburi | Central | Male | 54 | Blood | Sepsis | 2 | 1 | 1 | epf+/sly+/mrp+ |
| 41315 | 2012 | Phetchabul | North | Male | 42 | Blood | Sepsis | 2 | 104 | 391 | epf-/sly+/mrp- |
| 41307 | 2012 | Saraburi | Central | Female | 83 | Blood | Sepsis, Meningitis | 2 | 1 | 1 | epf+/sly+/mrp+ |
| 41614 | 2012 | Sakon Nakhon | Northeast | Male | 60 | Blood | Sepsis | 2 | 1 | 1 | epf+/sly+/mrp+ |
| 41493 | 2012 | Phetchabul | North | Male | 61 | Blood | Sepsis | 2 | 1 | 1 | epf+/sly+/mrp+ |
| 41737 | 2012 | Uttaradit | North | Female | 66 | Blood | Sepsis | 2 | 104 | 104 | epf-/sly+/mrp- |
| 41707 | 2012 | Uttaradit | North | Female | 55 | Blood | Sepsis | 2 | 1 | 1 | epf+/sly+/mrp+ |
| 41743 | 2012 | Saraburi | Central | Female | 70 | Blood | Sepsis | 2 | 1 | 1 | epf+/sly+/mrp+ |
| 41948 | 2012 | Lampang | North | Female | 70 | Synovial Fluid | Septic Arthritis | 2 | 1 | 1 | epf+/sly+/mrp+ |

| | | | | | | | | | | | |
|---|---|---|---|---|---|---|---|---|---|---|---|
| 42039 | 2012 | Saraburi | Central | Female | 54 | Blood | Sepsis | 2 | 1 | 1 | epf+/sly+/mrp+ |
| 41766 | 2012 | Singburi | Central | Male | 75 | Blood | Sepsis | 2 | 104 | 391 | epf-/sly+/mrp- |
| 41840 | 2012 | Phetchabul | North | Male | 63 | Blood | Sepsis, Meningitis | 2 | 1 | 1 | epf+/sly+/mrp+ |
| 41745 | 2012 | Maesod | North | Male | 85 | Blood | Sepsis, Meningitis | 2 | 1 | 1 | epf+/sly+/mrp+ |
| 42073 | 2012 | Lampang | North | Male | 47 | Blood | Sepsis | 2 | 28 | 382 | epf-/sly-/mrp+ |
| 41940 | 2012 | Nakhon Phanom | Northeast | Male | 36 | Blood | Sepsis | 2 | 1 | 337 | epf+/sly+/mrp+ |
| 41962 | 2012 | Uttaradit | North | Male | 57 | Blood | Sepsis | 2 | 104 | 104 | epf-/sly+/mrp- |
| 41870 | 2012 | Bhudhachinaraj | North | Male | 73 | Blood | Sepsis | 2 | 1 | 1 | epf+/sly+/mrp+ |
| 42324 | 2012 | Bhudhachinaraj | North | Male | NA | Blood | Sepsis | 2 | 104 | 104 | epf-/sly+/mrp- |
| 42052 | 2012 | Uttaradit | North | Male | 55 | Blood | Sepsis | 2 | 104 | 104 | epf-/sly+/mrp- |
| 52053 | 2012 | Uttaradit | North | Male | 61 | Blood | Sepsis, Meningitis | 2 | 1 | 1 | epf+/sly+/mrp+ |
| 42044 | 2012 | Phetchabul | North | Male | 69 | Blood | Sepsis | 2 | 1 | 1 | epf+/sly+/mrp+ |
| 42102 | 2012 | Nakhon Pathom | Central | Male | 88 | Blood | Sepsis | 2 | 1 | 1 | epf+/sly+/mrp+ |
| 42113 | 2012 | Maharajnakhonrajasima | Northeast | Male | 66 | Blood | Sepsis | 2 | 1 | 1 | epf+/sly+/mrp+ |
| 42317 | 2012 | Bhudhachinaraj | North | Female | NA | Blood | Sepsis | 2 | 104 | 104 | epf-/sly+/mrp- |
| 42411 | 2012 | Nakhon Phanom | Northeast | Female | 55 | Blood | Sepsis | 2 | 1 | 1 | epf+/sly+/mrp+ |
| 42428 | 2012 | Nakhon Phanom | Northeast | Female | 61 | Blood | Sepsis | 2 | 1 | 1 | epf+/sly+/mrp+ |
| 42437 | 2012 | Phetchabul | North | Female | 68 | Blood | Sepsis | 2 | 104 | 104 | epf-/sly+/mrp- |
| 42054 | 2012 | Uttaradit | North | Male | 33 | Blood | Sepsis | 2 | 1 | 1 | epf+/sly+/mrp+ |
| 42050 | 2012 | Taksin | Central | Male | 68 | Blood | Sepsis | 2 | 1 | 1 | epf+/sly+/mrp+ |
| 42527 | 2012 | Bhudhachinaraj | North | Male | NA | Blood | Sepsis | 2 | 233 | 233 | epf-/sly+/mrp- |
| 42323 | 2012 | Bhudhachinaraj | North | Male | NA | Blood | Sepsis | 2 | 104 | 104 | epf-/sly+/mrp- |
| 42213 | 2012 | Phetchabul | North | Male | 34 | Blood | Sepsis | 2 | 104 | 104 | epf-/sly+/mrp- |
| 42181 | 2012 | Pichit | Central | Male | 32 | Blood | Sepsis | 2 | 104 | 104 | epf-/sly+/mrp- |
| 42080 | 2012 | Pichit | Central | NA | 55 | Blood | Sepsis | 2 | 104 | 104 | epf-/sly+/mrp- |
| 42319 | 2012 | Bhudhachinaraj | North | Male | NA | Blood | Sepsis | 2 | 28 | 382 | epf-/sly-/mrp+ |

| | | | | | | | | | | | |
|---|---|---|---|---|---|---|---|---|---|---|---|
| 42216 | 2012 | Phetchabul | North | Male | 33 | Blood | Sepsis | 2 | 104 | 104 | epf-/sly+/mrp- |
| 42219 | 2012 | Phetchabul | North | Male | 40 | CSF | Meningitis | 2 | 1 | 1 | epf+/sly+/mrp+ |
| 42310 | 2012 | Saraburi | Central | Male | 47 | Blood | Sepsis, Meningitis | 2 | 1 | 1 | epf+/sly+/mrp+ |
| 42316 | 2012 | Uttaradit | North | Male | 69 | Blood | Sepsis | 2 | 1 | 1 | epf+/sly+/mrp+ |
| 42506 | 2012 | Pichit | Central | Male | 56 | Blood | Sepsis | 2 | 233 | 233 | epf-/sly+/mrp- |
| 42312 | 2012 | Maharajnakhonrajasima | Northeast | Male | 59 | Blood | Sepsis | 2 | 1 | 1 | epf+/sly+/mrp+ |
| 42337 | 2012 | Phetchabul | North | Male | 47 | CSF | Meningitis | 2 | 104 | 104 | epf-/sly+/mrp- |
| 42495 | 2012 | Saraburi | Central | Male | 42 | Blood | Sepsis | 2 | 1 | 1 | epf+/sly+/mrp+ |
| 42578 | 2012 | Nakhon Phanom | Northeast | Male | 54 | Blood | Sepsis | 2 | 1 | 1 | epf+/sly+/mrp+ |
| 42494 | 2012 | Saraburi | Central | Male | 65 | Blood | Sepsis | 2 | 1 | 1 | epf+/sly+/mrp+ |
| 42425 | 2012 | Rayong | East | Male | 32 | Blood | Sepsis | 2 | 104 | 104 | epf-/sly+/mrp- |
| 42445 | 2012 | Maesod | North | Female | NA | Blood | Sepsis | 2 | 104 | 104 | epf-/sly+/mrp- |
| 42610 | 2012 | Phayao | North | Female | 80 | Blood | Sepsis | 2 | 104 | 104 | epf-/sly+/mrp- |
| 42711 | 2012 | Hat Yai | South | Male | 51 | CSF | Meningitis | 2 | 1 | 1 | epf+/sly+/mrp+ |
| 42507 | 2012 | Singburi | Central | Male | 67 | Blood | Sepsis | 2 | 104 | 104 | epf-/sly+/mrp- |
| 42457 | 2012 | Phrajomklao | Central | Male | 63 | Blood | Sepsis | 2 | 104 | 104 | epf-/sly+/mrp- |
| 42617 | 2012 | Phayao | North | Male | 80 | Blood | Sepsis, Meningitis | 2 | 1 | 1 | epf+/sly+/mrp+ |
| 42570 | 2012 | Uttaradit | North | Male | 79 | Blood | Sepsis | 2 | 1 | 1 | epf+/sly+/mrp+ |
| 42587 | 2012 | Rayong | East | Male | 68 | Blood | Sepsis | 2 | 104 | 104 | epf-/sly+/mrp- |
| 42563 | 2012 | Phramonkutklao | Central | Male | 80 | Blood | Sepsis | 2 | 28 | 28 | epf-/sly-/mrp+ |
| 42780 | 2012 | Phrapokklao | East | Male | 65 | Blood | Sepsis | 2 | 1 | 105 | epf+/sly+/mrp+ |
| 42713 | 2012 | Uttaradit | North | Male | 60 | Blood | Sepsis | 2 | 104 | 104 | epf-/sly+/mrp- |
| 42725 | 2012 | Chaophrayayommaraj | Central | Male | NA | Blood | Sepsis | 2 | 104 | 104 | epf-/sly+/mrp- |
| 42994 | 2012 | Maharajnakhonrajasima | Northeast | Male | 73 | CSF | Meningitis | 2 | 1 | 1 | epf+/sly+/mrp+ |
| 42611 | 2012 | Sakon Nakhon | Northeast | Male | 38 | Blood | Sepsis | 2 | 1 | 1 | epf+/sly+/mrp+ |
| 42699 | 2012 | Uttaradit | North | Male | 52 | Blood | Sepsis | 2 | 104 | 104 | epf-/sly+/mrp- |

| | | | | | | | | | | | |
|---|---|---|---|---|---|---|---|---|---|---|---|
| 42714 | 2012 | Uttaradit | North | Male | 45 | Blood | Sepsis | 2 | 104 | 514 | epf-/sly+/mrp- |
| 42847 | 2012 | Maharajnakhonrajasima | Northeast | Female | 76 | CSF | Meningitis | 2 | 1 | 1 | epf+/sly+/mrp+ |
| 42772 | 2012 | Udonthani | Northeast | Male | NA | Blood | Sepsis | 2 | 1 | 1 | epf+/sly+/mrp+ |
| 42771 | 2012 | Phranungklao | Central | Female | NA | CSF | Meningitis | 2 | 1 | 1 | epf+/sly+/mrp+ |
| 42841 | 2012 | Uttaradit | North | Male | 66 | Blood | Sepsis, Meningitis | 14 | 1 | 105 | epf+/sly+/mrp+ |
| 42870 | 2012 | Saraburi | Central | Male | 52 | Blood | Sepsis, Meningitis | 2 | 1 | 1 | epf+/sly+/mrp+ |
| 43013 | 2012 | Bhudhachinaraj | North | Male | NA | Blood | Sepsis | 2 | 104 | 104 | epf-/sly+/mrp- |
| 42885 | 2012 | Saraburi | Central | Male | 89 | Blood | Sepsis | 2 | 28 | 28 | epf-/sly-/mrp+ |
| 43021 | 2012 | Phetchabul | North | Male | 64 | Blood | Sepsis | 2 | 1 | 1 | epf+/sly+/mrp+ |
| 43024 | 2012 | Saraburi | Central | Female | 53 | Blood | Sepsis | 2 | 1 | 1 | epf+/sly+/mrp+ |
| 21386 | 2006 | Nakornsawan | Central | Male | 54 | Blood | Acute Meningitis | 2 | 1 | 1 | epf+/sly+/mrp+ |
| 22687 | 2006 | Nan | North | Male | 76 | Blood | Sepsis, Septic Shock | 2 | 104 | 104 | epf-/sly+/mrp- |
| 22689 | 2006 | Nan | North | Male | 45 | Blood | Sepsis, Septic Shock, Septic Arthritis | 2 | 104 | 104 | epf-/sly+/mrp- |
| 22713 | 2006 | Nan | North | Male | 65 | Blood | Sepsis, Sepsis Shock, Septic Arthritis | 2 | 104 | 104 | epf-/sly+/mrp- |
| 31995 | 2010 | Phetchabul | North | Male | NA | Blood | Sepsis | 2 | 1 | 1 | epf+/sly+/mrp+ |
| 31998 | 2010 | Phetchabul | North | NA | NA | CSF | Acute Meningitis | 14 | 1 | 105 | epf+/sly+/mrp+ |
| 32006 | 2010 | Phetchabul | North | NA | NA | Blood | Sepsis | 14 | 1 | 105 | epf+/sly+/mrp+ |
| 32007 | 2010 | Phetchabul | North | NA | NA | Blood | Sepsis | 2 | 104 | 104 | epf-/sly+/mrp- |
| 32008 | 2010 | Phetchabul | North | NA | NA | Blood | Sepsis | 2 | 104 | 104 | epf-/sly+/mrp- |
| 32011 | 2010 | Phetchabul | North | NA | NA | Blood | Sepsis | 2 | 104 | 104 | epf-/sly+/mrp- |
| 32024 | 2010 | Surathani | South | NA | NA | CSF | Acute Meningitis | 2 | 1 | 1 | epf+/sly+/mrp+ |
| 32027 | 2010 | Maharajnakhonrajasima | Northeast | NA | NA | Blood | Sepsis | 2 | 104 | 104 | epf-/sly+/mrp- |
| 32098 | 2010 | Lampang | North | NA | NA | Blood | Sepsis | 24 | 221/234 | 234 | epf-/sly-/mrp- |
| 32101 | 2010 | Phetchabul | North | NA | NA | CSF | Acute Meningitis | 2 | 1 | 1 | epf+/sly+/mrp+ |
| 32105 | 2010 | Maharajnakhonrajasima | Northeast | NA | NA | Blood | Sepsis | 2 | 1 | 1 | epf+/sly+/mrp+ |
| 32334 | 2010 | Phetchabul | North | NA | NA | Blood | Sepsis | 2 | 104 | 104 | epf-/sly+/mrp- |

| | | | | | | | | | | | |
|---|---|---|---|---|---|---|---|---|---|---|---|
| 32336 | 2010 | Phetchabul | North | NA | NA | Blood | Sepsis, Meningitis | 14 | 1 | 105 | epf+/sly+/mrp+ |
| 32340 | 2010 | Phetchabul | North | NA | NA | CSF | Acute Meningitis | 14 | 1 | 105 | epf+/sly+/mrp+ |
| 32341 | 2010 | Phayao | North | NA | NA | Blood | Sepsis, Meningitis | 2 | 1 | 1 | epf+/sly+/mrp+ |
| 32342 | 2010 | Phayao | North | NA | NA | Blood | Sepsis | 2 | 1 | 1 | epf+/sly+/mrp+ |
| 32471 | 2010 | Phetchabul | North | NA | NA | Blood | Sepsis | 2 | 1 | 1 | epf+/sly+/mrp+ |
| 32472 | 2010 | Phetchabul | North | NA | NA | blood | Sepsis | 2 | 104 | 104 | epf-/sly+/mrp- |
| 32478 | 2010 | Phetchabul | North | NA | NA | Blood | Sepsis | 2 | 104 | 104 | epf-/sly+/mrp- |
| 32479 | 2010 | Phetchabul | North | NA | NA | CSF | Acute Meningitis | 14 | 1 | 105 | epf+/sly+/mrp+ |
| 32481 | 2010 | Phetchabul | North | NA | NA | Blood | Sepsis | 14 | 1 | 105 | epf+/sly+/mrp+ |
| 32482 | 2010 | Phetchabul | North | NA | NA | Blood | Sepsis | 2 | 104 | 104 | epf-/sly+/mrp- |
| 32487 | 2010 | Phayao | North | NA | NA | Blood | Sepsis | 2 | 1 | 1 | epf+/sly+/mrp+ |
| 32510 | 2010 | Hua Cheao | Central | NA | NA | Blood | Sepsis | 2 | 104 | 104 | epf-/sly+/mrp- |
| 32511 | 2010 | Maharajnakhonrajasima | Northeast | NA | NA | Blood | Sepsis | 2 | 1 | 1 | epf+/sly+/mrp+ |
| 32512 | 2010 | Phetchabul | North | NA | NA | CSF | Acute Meningitis | 14 | 1 | 105 | epf+/sly+/mrp+ |
| 32546 | 2010 | Phetchabul | North | NA | NA | Blood | Sepsis | 2 | 104 | 104 | epf-/sly+/mrp- |
| 35592 | 2011 | Phramonkutklao | Central | Male | NA | Blood | Sepsis, Meningitis | 2 | 1 | 1 | epf+/sly+/mrp+ |
| 35611 | 2011 | Phrapokklao | East | Male | 45 | Blood | Sepsis | 2 | 233 | 233 | epf-/sly+/mrp- |
| 35680 | 2011 | Chaophrayayommaraj | Central | Male | NA | Blood | Sepsis | 2 | 104 | 104 | epf-/sly+/mrp- |
| 35688 | 2011 | Rayong | East | Female | 63 | Blood | Sepsis | 2 | 233 | 233 | epf-/sly+/mrp- |
| 35690 | 2011 | Nakhon Phanom | Northeast | Male | NA | Blood | Sepsis | 2 | 1 | 1 | epf+/sly+/mrp+ |
| 35728 | 2011 | Maharajnakhonrajasima | Northeast | Male | NA | Blood | Sepsis | 14 | 1 | 105 | epf+/sly+/mrp+ |
| 35736 | 2011 | Phayao | North | Male | 72 | Blood | Sepsis | 2 | 1 | 298 | epf+/sly+/mrp+ |
| 35737 | 2011 | Phayao | North | Male | 56 | Blood | Sepsis | 2 | 1 | 1 | epf+/sly+/mrp+ |
| 35738 | 2011 | Phayao | North | Male | 52 | Blood | Sepsis | 2 | 104 | 104 | epf-/sly+/mrp- |
| 35739 | 2011 | Phayao | North | Female | 47 | Blood | Sepsis | 2 | 1 | 1 | epf+/sly+/mrp+ |
| 35755 | 2011 | Maharajnakhonrajasima | Northeast | Male | 32 | CSF | Acute Meningitis | 2 | 1 | 1 | epf+/sly+/mrp+ |

| | | | | | | | | | | | |
|---|---|---|---|---|---|---|---|---|---|---|---|
| 39564 | 2012 | Phetchabul | North | Female | 75 | Blood | Sepsis | 2 | 1 | 1 | epf+/sly+/mrp+ |
| 39565 | 2012 | Taksin | Central | Female | 56 | Blood | Sepsis | 24 | 221/234 | 221 | epf-/sly-/mrp- |
| 39598 | 2012 | Sakon Nakhon | Northeast | Male | 61 | CSF | Meningitis | 2 | 1 | 1 | epf+/sly+/mrp+ |
| 39716 | 2012 | Loei | Northeast | Male | 49 | CSF | Meningitis | 2 | 1 | 1 | epf+/sly+/mrp+ |
| 39728 | 2012 | Sakon Nakhon | Northeast | Female | 47 | Blood | Sepsis, Meningitis | 2 | 1 | 1 | epf+/sly+/mrp+ |
| 40085 | 2012 | Lampang | North | Female | 91 | Blood | Sepsis | 2 | 233 | 379 | epf-/sly+/mrp- |
| 33377 | 2010 | Pichit | Central | NA | NA | Blood | Sepsis | 2 | 104 | 104 | epf-/sly+/mrp- |
| 33378 | 2010 | Pichit | Central | NA | NA | Blood | Sepsis | 2 | 1 | 1 | epf+/sly+/mrp+ |
| 33394 | 2010 | Mongkutwattana | Central | NA | NA | Blood | Sepsis | 2 | 233 | 233 | epf-/sly+/mrp- |
| 33434 | 2010 | Nakhon Pathom | Central | NA | NA | Blood | Sepsis | 2 | 1 | 1 | epf+/sly+/mrp+ |
| 33436 | 2010 | Maharajnakhonrajasima | Northeast | NA | NA | Blood | Sepsis | 2 | 1 | 1 | epf+/sly+/mrp+ |
| 33514 | 2010 | Maharajnakhonrajasima | Northeast | NA | NA | Blood, CSF | Acute Meningitis | 2 | 1 | 1 | epf+/sly+/mrp+ |
| 33522 | 2010 | Phetchabul | North | NA | NA | CSF | Acute Meningitis | 2 | 1 | 1 | epf+/sly+/mrp+ |
| 33523 | 2010 | Maharajnakhonrajasima | Northeast | NA | NA | Blood | Sepsis | 2 | 104 | 104 | epf-/sly+/mrp- |
| 33590 | 2010 | Maharajnakhonrajasima | Northeast | NA | NA | Blood | Sepsis | 2 | 1 | 1 | epf+/sly+/mrp+ |
| 33591 | 2010 | Maharajnakhonrajasima | Northeast | NA | NA | Blood | Sepsis | 2 | 1 | 1 | epf+/sly+/mrp+ |
| 36642 | 2011 | Sa Kaeo | East | Male | NA | Blood | Sepsis | 2 | 1 | 1 | epf+/sly+/mrp+ |
| 36643 | 2011 | Sa Kaeo | East | Male | NA | Blood | Sepsis | 2 | 1 | 1 | epf+/sly+/mrp+ |
| 36644 | 2011 | Sa Kaeo | East | Male | NA | Blood | Sepsis | 2 | 1 | 144 | epf+/sly+/mrp+ |
| 36730 | 2011 | Maharajnakhonrajasima | Northeast | Female | 32 | Blood | Sepsis | 2 | 104 | 104 | epf-/sly+/mrp- |
| 36733 | 2011 | Burirum | Northeast | Male | 45 | Blood | Sepsis | 2 | 1 | 1 | epf+/sly+/mrp+ |
| 36743 | 2011 | Sakon Nakhon | Northeast | Male | 42 | CSF | Acute Meningitis | 2 | 1 | 1 | epf+/sly+/mrp+ |
| 36745 | 2011 | Phrapokklao | East | Female | 48 | Blood | Sepsis | 2 | 233 | 233 | epf-/sly+/mrp- |
| 36767 | 2011 | Sa Kaeo | East | Male | NA | Blood | Sepsis | 2 | 1 | 1 | epf+/sly+/mrp+ |
| 36817 | 2011 | Sakon Nakhon | Northeast | Male | 36 | Blood | Sepsis | 2 | 1 | 1 | epf+/sly+/mrp+ |
| 36821 | 2011 | Uttaradit | North | Female | 50 | Blood | Sepsis | 2 | 104 | 104 | epf-/sly+/mrp- |

| | | | | | | | | | | | |
|---|---|---|---|---|---|---|---|---|---|---|---|
| 36830 | 2011 | Nakhonping | North | Male | 77 | Blood | Sepsis | 2 | 25 | 380 | epf-/sly-/mrp+ |
| 36871 | 2011 | Maharajnakhonrajasima | Northeast | Male | 42 | Blood | Sepsis | 2 | 104 | 104 | epf-/sly+/mrp- |
| 36900 | 2011 | Nan | North | Male | 49 | Blood | Sepsis | 2 | 29 | 516 | epf-/sly-/mrp+ |
| 36901 | 2011 | Nan | North | Male | 45 | Blood | Sepsis | 2 | 104 | 512 | epf-/sly+/mrp- |
| 36903 | 2011 | Nan | North | Male | 62 | Blood | Sepsis | 2 | 104 | 512 | epf-/sly+/mrp- |
| 36905 | 2011 | Chaophrayayommaraj | Central | Female | NA | Blood | Sepsis | 2 | 233 | 233 | epf-/sly+/mrp- |
| 36917 | 2011 | Nan | North | Female | 49 | Blood | Sepsis, Meningitis | 2 | 104 | 512 | epf-/sly+/mrp- |
| 36918 | 2011 | Nan | North | Male | 44 | Blood | Sepsis | 2 | 233 | 233 | epf-/sly+/mrp- |
| 36919 | 2011 | Nan | North | Male | 80 | Blood | Sepsis | 2 | 104 | 512 | epf-/sly+/mrp- |
| 36920 | 2011 | Nan | North | Male | 78 | Blood | Sepsis | 2 | 104 | 512 | epf-/sly+/mrp- |
| 36928 | 2011 | Lampang | North | Male | 61 | Blood | Sepsis | 2 | 1 | 1 | epf+/sly+/mrp+ |
| 36976 | 2011 | Phetchabul | North | Male | 57 | Blood | Sepsis, Meningitis | 2 | 1 | 1 | epf+/sly+/mrp+ |
| 36979 | 2011 | Roi Et | Northeast | Male | 59 | Blood | Sepsis | 2 | 1 | 1 | epf+/sly+/mrp+ |
| 36994 | 2011 | Nakhon Phanom | Northeast | Male | NA | Blood | Sepsis | 2 | 1 | 1 | epf+/sly+/mrp+ |
| 36999 | 2011 | Maharajnakhonrajasima | Northeast | Female | 61 | Blood | Sepsis | 2 | 25 | 380 | epf-/sly-/mrp+ |
| 37037 | 2011 | Nakhonping | North | Male | 50 | Blood | Sepsis | 2 | 104 | 104 | epf-/sly+/mrp- |
| 37045 | 2011 | Nakhonping | North | Male | 58 | Blood | Sepsis | 2 | 104 | 104 | epf-/sly+/mrp- |
| 37193 | 2011 | Phetchabul | North | Male | 70 | Blood | Sepsis | 2 | 1 | 1 | epf+/sly+/mrp+ |
| 37655 | 2011 | Maharajnakhonrajasima | Northeast | Male | 50 | Blood | Sepsis | 2 | 104 | 104 | epf-/sly+/mrp- |
| 38290 | 2011 | Lampang | North | Male | NA | Blood | Sepsis | 2 | 1 | 1 | epf+/sly+/mrp+ |
| 41375 | 2012 | Bhudhachinaraj | North | Male | 54 | Blood | Sepsis, Meningitis | 2 | 1 | 1 | epf+/sly+/mrp+ |
| 41386 | 2012 | Singburi | Central | Male | 56 | Blood, CSF | Meningitis | 2 | 1 | 1 | epf+/sly+/mrp+ |
| 41656 | 2012 | Phetchabul | North | Female | 74 | Blood | Sepsis | 2 | 1 | 1 | epf+/sly+/mrp+ |
| 34365 | 2010 | Lampang | North | Male | 47 | Blood | Sepsis, Meningitis | 2 | 1 | 1 | epf+/sly+/mrp+ |
| 34436 | 2010 | Lampang | North | Male | 52 | Blood | Sepsis | 2 | 1 | 1 | epf+/sly+/mrp+ |
| 34472 | 2010 | Vajira | Central | Female | NA | NA | NA | 2 | 104 | 104 | epf-/sly+/mrp- |

| | | | | | | | | | | | |
|---|---|---|---|---|---|---|---|---|---|---|---|
| 34479 | 2010 | Mongkutwattana | Central | Male | 51 | Blood | Sepsis | 2 | 25 | 25 | epf-/sly-/mrp+ |
| 38436 | 2011 | Nakhon Phanom | Northeast | NA | NA | NA | NA | 2 | 1 | 1 | epf+/sly+/mrp+ |
| 38439 | 2011 | Hat Yai | South | Female | 46 | Blood | Sepsis | 2 | 1 | 1 | epf+/sly+/mrp+ |
| 43011 | 2012 | Maesod | North | Female | 59 | Blood | Sepsis, Meningitis | 2 | 1 | 1 | epf+/sly+/mrp+ |
| 43022 | 2012 | Phetchabul | North | Female | 46 | Blood | Sepsis | 2 | 104 | 104 | epf-/sly+/mrp- |
| 21928 | 2006 | Lampang | North | Female | 48 | CSF | Meningitis | 14 | 1 | 105 | epf+/sly+/mrp+ |
| 34506 | 2011 | Phetchabul | North | Male | 80 | Blood | Sepsis | 2 | 104 | 104 | epf-/sly+/mrp- |
| 34517 | 2011 | Phetchabul | North | Male | 76 | Blood, CSF | Acute Meningitis | 2 | 1 | 1 | epf+/sly+/mrp+ |
| 34579 | 2011 | Bhudhachinaraj | North | Male | 74 | Blood | Sepsis | 2 | 25 | 381 | epf-/sly-/mrp+ |
| 34584 | 2011 | Rayong | East | Male | 73 | Blood | Sepsis | 2 | 29 | 515 | epf-/sly-/mrp+ |
| 34725 | 2011 | Roi Et | Northeast | Male | 42 | CSF | Acute Meningitis | 2 | 1 | 1 | epf+/sly+/mrp+ |
| 34758 | 2011 | Burirum | Northeast | Male | 72 | Blood | Sepsis | 2 | 25 | 380 | epf-/sly-/mrp+ |
| 34853 | 2011 | Phayao | North | Male | 59 | CSF | Acute Meningitis | 2 | 1 | 1 | epf+/sly+/mrp+ |
| 34854 | 2011 | Phayao | North | NA | 64 | Blood | Sepsis | 2 | 1 | 1 | epf+/sly+/mrp+ |
| 38544 | 2012 | Phetchabul | North | Female | 57 | Blood | Sepsis | 2 | 104 | 104 | epf-/sly+/mrp- |
| 41445 | 2012 | Maesod | North | Male | NA | Blood | Sepsis | 2 | 1 | 105 | epf+/sly+/mrp+ |
| 43434 | 2006 | Phetchabul | North | Male | 63 | Blood | Sepsis | 2 | 233 | 233 | epf-/sly+/mrp- |

**Supplementary Table 2**. Metadata regarding zoonotic S. suis isolates collected in the passive surveillance study carried out in North and Northeast Thailand between 2013 and 2021. NA stands for not available. CSF stands for cerebrospinal fluid. CC stands for clonal complex. Serotype and CC were determined by PCR.

| Isolate | Hospital | Region | Age | Sex | Specimen | Isolation year | Serotype | CC |
|---|---|---|---|---|---|---|---|---|
| STC217 | Chiang Mai | North | NA | NA | Blood | 2013 | 2 | 1 |
| STC218 | Chiang Mai | North | NA | NA | CSF | 2013 | 2 | 1 |
| STC219 | Chiang Mai | North | NA | NA | Blood | 2013 | 2 | 1 |
| STC220 | Chiang Mai | North | NA | NA | CSF | 2014 | 2 | 1 |
| STC221 | Phrae | North | NA | NA | Blood | 2014 | 2 | 1 |
| STC222 | Phrae | North | NA | NA | Blood | 2014 | 2 | 1 |
| STC223 | Phrae | North | NA | NA | Blood | 2014 | 2 | 233 |
| STC224 | Phrae | North | NA | NA | Blood | 2014 | 2 | 1 |
| STC225 | Phrae | North | NA | NA | Blood | 2014 | 2 | 1 |
| STC226 | Phrae | North | NA | NA | Blood | 2014 | 2 | 1 |
| STC227 | Phrae | North | NA | NA | Blood | 2014 | 2 | 1 |
| STC228 | Phrae | North | NA | NA | Blood | 2014 | 2 | 104 |
| STC229 | Phrae | North | NA | NA | Blood | 2014 | 2 | 1 |
| STC230 | Phrae | North | NA | NA | Blood | 2014 | 2 | 1 |
| STC231 | Chiang Mai | North | NA | NA | Blood | 2015 | 2 | 1 |
| STC232 | Phrae | North | NA | NA | Blood | 2015 | 2 | 1 |
| STC233 | Chiang Mai | North | NA | NA | Blood | 2015 | 2 | 233 |
| STC234 | Chiang Mai | North | NA | NA | Blood | 2015 | 2 | 1 |
| STC235 | Chiang Mai | North | NA | NA | Blood | 2015 | 2 | 233 |
| STC236 | Chiang Mai | North | NA | NA | Blood | 2015 | 2 | 233 |
| STC237 | Chiang Mai | North | NA | NA | Blood | 2015 | 2 | 233 |
| STC238 | Chiang Mai | North | NA | NA | CSF | 2016 | 2 | 1 |
| STC272 | Chiang Mai | North | NA | NA | Blood | 2016 | 2 | 233 |

| | | | | | | | | |
|---|---|---|---|---|---|---|---|---|
| STC239 | Chiang Mai | North | NA | NA | Blood | 2016 | 2 | 233 |
| STC240 | Chiang Mai | North | NA | NA | Blood | 2016 | 2 | 233 |
| STC241 | Chiang Mai | North | NA | NA | Blood | 2016 | 2 | 233 |
| STC242 | Lamphoon | North | NA | NA | Blood | 2016 | 2 | 104 |
| STC243 | Lamphoon | North | NA | NA | Blood | 2016 | 2 | 233 |
| STC244 | Phrae | North | NA | NA | Blood | 2017 | 2 | 104 |
| STC245 | Phrae | North | NA | NA | Blood | 2017 | 2 | 104 |
| STC246 | Phrae | North | NA | NA | Blood | 2017 | 2 | 104 |
| STC247 | Chiang Mai | North | NA | NA | Blood | 2017 | 2 | 1 |
| STC248 | Chiang Mai | North | NA | NA | Blood | 2017 | 2 | 1 |
| STC249 | Nan | North | NA | NA | Blood | 2017 | 2 | 104 |
| STC250 | Chiang Mai | North | NA | NA | Blood | 2017 | 2 | 233 |
| STC7 | Phetchaboon | North | 84 | Female | Blood | 2019 | 14 | 1 |
| STC251 | Chiang Mai | North | NA | NA | Blood | 2019 | 2 | 104 |
| STC252 | Chiang Mai | North | NA | NA | Blood | 2019 | 2 | 233 |
| STC253 | Chiang Mai | North | NA | NA | Blood | 2019 | 2 | 104 |
| STC254 | Phrae | North | NA | NA | Blood | 2019 | 2 | 104 |
| STC255 | Chiang Mai | North | NA | NA | Blood | 2019 | 2 | 233 |
| STC256 | Chiang Mai | North | NA | NA | Blood | 2019 | 2 | 104 |
| STC257 | Chiang Mai | North | NA | NA | Blood | 2019 | 2 | 233 |
| STC273 | Chiang Mai | North | NA | NA | Blood | 2019 | 2 | 1 |
| STC274 | Chiang Mai | North | NA | NA | Blood | 2019 | 2 | 233 |
| STC275 | Chiang Mai | North | NA | NA | Blood | 2019 | 2 | 1 |
| STC276 | Chiang Mai | North | NA | NA | CSF | 2019 | 2 | 233 |
| STC277 | Chiang Mai | North | NA | NA | Blood | 2020 | 2 | 1 |
| STC278 | Chiang Mai | North | NA | NA | Blood | 2020 | 2 | 233 |

| | | | | | | | | |
|---|---|---|---|---|---|---|---|---|
| STC279 | Lampang | North | NA | NA | Blood | 2020 | 2 | 233 |
| STC280 | Lampang | North | NA | NA | Blood | 2020 | 2 | 233 |
| STC281 | Lampang | North | NA | NA | Blood | 2020 | 2 | 233 |
| STC282 | Lampang | North | NA | NA | Blood | 2020 | 2 | 104 |
| STC283 | Lampang | North | NA | NA | Blood | 2020 | 2 | 104 |
| STC284 | Lampang | North | NA | NA | Blood | 2020 | 2 | 104 |
| STC285 | Lampang | North | NA | NA | Blood | 2020 | 2 | 104 |
| STC286 | Lampang | North | NA | NA | Blood | 2020 | 2 | 233 |
| STC287 | Lampang | North | NA | NA | Blood | 2020 | 2 | 104 |
| STC288 | Lampang | North | NA | NA | Blood | 2020 | 2 | 233 |
| STC289 | Lampang | North | NA | NA | Blood | 2020 | 2 | 1 |
| STC290 | Lampang | North | NA | NA | Blood | 2020 | 2 | 104 |
| STC291 | Lampang | North | NA | NA | Blood | 2020 | 2 | 104 |
| STC292 | Lampang | North | NA | NA | Blood | 2020 | 2 | 104 |
| STC293 | Chiang Mai | North | NA | NA | Blood | 2020 | 2 | 1 |
| STC294 | Chiang Mai | North | NA | NA | Blood | 2020 | 2 | 233 |
| STC19 | Roi-Et | Northeast | 52 | Male | Blood | 2021 | 2 | 1 |
| STC31 | Phetchabun | North | 61 | Male | Blood | 2021 | 2 | 233 |
| STC70 | Roi-Et | Northeast | 80 | Male | Blood | 2021 | 2 | 233 |
| STC72 | Roi-Et | Northeast | NA | Male | Blood | 2021 | 2 | 104 |
| STC95 | Phetchabun | North | 62 | Female | CSF | 2021 | 2 | 1 |
| STC96 | Phetchabun | North | 30 | Male | CSF | 2021 | 2 | 1 |
| STC97 | Phetchabun | North | 75 | Female | Blood | 2021 | 2 | 1 |
| STC103 | Phetchabun | North | 56 | Female | Blood | 2021 | 2 | 1 |
| STC106 | Roi-Et | Northeast | 53 | Female | Blood | 2021 | 2 | 1 |
| STC108 | Roi-Et | Northeast | 85 | Male | Blood | 2021 | 2 | 1 |

| | | | | | | | | |
|---|---|---|---|---|---|---|---|---|
| STC111 | Roi-Et | Northeast | 37 | Male | Blood | 2021 | 2 | 1 |
| STC115 | Surin | Northeast | 69 | Male | Blood | 2021 | 2 | 1 |
| STC116 | Surin | Northeast | 43 | Male | Blood | 2021 | 2 | 1 |
| STC123 | Phetchabun | North | 62 | Female | Blood | 2021 | 2 | 1 |
| STC125 | Phetchabun | North | 56 | Female | CSF | 2021 | 2 | 1 |
| STC127 | Nakhon Ratchasima | Northeast | NA | NA | Blood | 2021 | 2 | 1 |
| STC129 | Nakhon Ratchasima | Northeast | NA | NA | Blood | 2021 | 2 | 1 |
| STC130 | Nakhon Ratchasima | Northeast | NA | NA | Blood | 2021 | 2 | 233 |
| STC131 | Nakhon Ratchasima | Northeast | NA | NA | Blood | 2021 | 2 | 233 |
| STC132 | Nakhon Ratchasima | Northeast | NA | NA | Blood | 2021 | 2 | 233 |
| STC133 | Nakhon Ratchasima | Northeast | NA | NA | Blood | 2021 | 2 | 1 |
| STC134 | Nakhon Ratchasima | Northeast | NA | NA | Blood | 2021 | 2 | 1 |
| STC135 | Nakhon Ratchasima | Northeast | NA | NA | Blood | 2021 | 2 | 233 |
| STC137 | Phetchabun | North | 56 | Male | Blood | 2021 | 2 | 233 |
| STC141 | Phetchabun | North | 57 | Female | Blood | 2021 | 2 | 233 |
| STC143 | Surin | Northeast | NA | Male | NA | 2021 | 2 | 1 |
| STC144 | Surin | Northeast | NA | Female | NA | 2021 | 2 | 1 |
| STC145 | Surin | Northeast | NA | Female | NA | 2021 | 2 | 1 |
| STC146 | Surin | Northeast | NA | Male | NA | 2021 | 2 | 1 |
| STC147 | Chiang Mai | North | 63 | Male | Blood | 2021 | 2 | 25 |
| STC149 | Phetchabun | North | 43 | Female | Blood | 2021 | 14 | 233 |
| STC152 | Nakhon Ratchasima | Northeast | NA | NA | Blood | 2021 | 2 | 104 |
| STC153 | Nakhon Ratchasima | Northeast | NA | NA | Blood | 2021 | 2 | 233 |
| STC154 | Nakhon Ratchasima | Northeast | NA | NA | Blood | 2021 | 2 | 104 |
| STC155 | Nakhon Ratchasima | Northeast | NA | NA | Blood | 2021 | 2 | 1 |
| STC156 | Nakhon Ratchasima | Northeast | NA | NA | Blood | 2021 | 2 | 1 |

| | | | | | | | | |
|---|---|---|---|---|---|---|---|---|
| STC157 | Nakhon Ratchasima | Northeast | NA | NA | Mitral valve | 2021 | 2 | 1 |
| STC172 | Nakhon Ratchasima | Northeast | NA | NA | Blood | 2021 | 2 | 1 |
| STC173 | Nakhon Ratchasima | Northeast | NA | NA | Blood | 2021 | 2 | 233 |
| STC174 | Nakhon Ratchasima | Northeast | NA | NA | Blood | 2021 | 2 | 233 |
| STC175 | Nakhon Ratchasima | Northeast | NA | NA | Blood | 2021 | 2 | 1 |
| STC176 | Nakhon Ratchasima | Northeast | NA | NA | Blood | 2021 | 2 | 1 |
| STC177 | Nakhon Ratchasima | Northeast | NA | NA | Blood | 2021 | 2 | 1 |
| STC178 | Nakhon Ratchasima | Northeast | NA | NA | Blood | 2021 | 2 | 1 |
| STC182 | Roi-Et | Northeast | 74 | Male | Blood | 2021 | 2 | 1 |
| STC191 | Nakhon Ratchasima | Northeast | NA | NA | Blood | 2021 | 2 | 233 |
| STC192 | Nakhon Ratchasima | Northeast | NA | NA | Blood | 2021 | 2 | 1 |
| STC193 | Nakhon Ratchasima | Northeast | NA | NA | Blood | 2021 | 2 | 1 |
| STC194 | Nakhon Ratchasima | Northeast | NA | NA | Blood | 2021 | 2 | 1 |
| STC195 | Nakhon Ratchasima | Northeast | NA | NA | Blood | 2021 | 2 | 1 |
| STC196 | Nakhon Ratchasima | Northeast | NA | NA | Blood | 2021 | 2 | 1 |
| STC197 | Nakhon Ratchasima | Northeast | NA | NA | Blood | 2021 | 2 | 1 |
| STC201 | Phetchabun | North | 61 | Male | Blood | 2021 | 2 | 1 |
| STC202 | Phetchabun | North | 69 | Female | Blood | 2021 | 2 | 25 |
| STC203 | Phetchabun | North | 58 | Male | CSF | 2021 | 2 | 1 |
| STC209 | Nakhon Ratchasima | Northeast | NA | NA | Blood | 2021 | 2 | NA |
| STC210 | Nakhon Ratchasima | Northeast | NA | NA | Blood | 2021 | 2 | 1 |
| STC211 | Nakhon Ratchasima | Northeast | NA | NA | Blood | 2021 | 2 | 1 |
| STC212 | Nakhon Ratchasima | Northeast | NA | NA | Blood | 2021 | 2 | 233 |
| STC213 | Nakhon Ratchasima | Northeast | NA | NA | Blood | 2021 | 2 | 233 |
| STC214 | Nakhon Ratchasima | Northeast | NA | NA | Blood | 2021 | 2 | 1 |
| STC215 | Nakhon Ratchasima | Northeast | NA | NA | Blood | 2021 | 2/14 | 1 |

| | | | | | | | | |
|---|---|---|---|---|---|---|---|---|
| STC216 | Nakhon Ratchasima | Northeast | NA | NA | Blood | 2021 | 2 | 1 |
| STC258 | Nakhon Ratchasima | Northeast | NA | NA | Blood | 2021 | 14 | NA |
| STC259 | Nakhon Ratchasima | Northeast | NA | NA | Blood | 2021 | 2 | 1 |
| STC260 | Nakhon Ratchasima | Northeast | NA | NA | Blood | 2021 | 2 | 233 |
| STC261 | Nakhon Ratchasima | Northeast | NA | NA | Blood | 2021 | 2 | 104 |
| STC262 | Nakhon Ratchasima | Northeast | NA | NA | Blood | 2021 | 2 | 104 |
| STC263 | Nakhon Ratchasima | Northeast | NA | NA | Blood | 2021 | 2 | 1 |
| STC264 | Nakhon Ratchasima | Northeast | NA | NA | Blood | 2021 | 2 | 1 |
| STC265 | Nakhon Ratchasima | Northeast | NA | NA | Blood | 2021 | 2 | 1 |
| STC266 | Nakhon Ratchasima | Northeast | NA | NA | Blood | 2021 | 2 | 104 |
| STC267 | Nakhon Ratchasima | Northeast | NA | NA | Blood | 2021 | 2 | 104 |
| STC268 | Nakhon Ratchasima | Northeast | NA | NA | Blood | 2021 | 2 | 104 |
| STC269 | Nakhon Ratchasima | Northeast | NA | NA | Blood | 2021 | 2 | 104 |
| STC270 | Nakhon Ratchasima | Northeast | NA | NA | Blood | 2021 | 2 | 104 |

**Supplementary Table 3**. Metadata regarding S. suis isolates collected from healthy pigs in Phayao province between 2010 and 2011. NA stands for not available. ST stands for sequence type.

| Isolate | Serotype | ST | Specimen | Isolation Year | Province | Region |
|---|---|---|---|---|---|---|
| 34376 | 9 | NA | Pig tonsil | 2010-2011 | Phayao | North |
| 34383 | NA | NA | Pig tonsil | 2010-2011 | Phayao | North |
| 34384 | 2 | 2 | Pig tonsil | 2010-2011 | Phayao | North |
| 34389 | 3 | NA | Pig tonsil | 2010-2011 | Phayao | North |
| 34390 | 31 | NA | Pig tonsil | 2010-2011 | Phayao | North |
| 34391 | 31 | NA | Pig tonsil | 2010-2011 | Phayao | North |
| 34394 | 3 | NA | Pig tonsil | 2010-2011 | Phayao | North |
| 34397 | 3 | NA | Pig tonsil | 2010-2011 | Phayao | North |
| 34399 | 3 | NA | Pig tonsil | 2010-2011 | Phayao | North |
| 34401 | 16 | NA | Pig tonsil | 2010-2011 | Phayao | North |
| 34403 | 16 | NA | Pig tonsil | 2010-2011 | Phayao | North |
| 34404 | 16 | NA | Pig tonsil | 2010-2011 | Phayao | North |
| 34406 | 2 | 1 | Pig tonsil | 2010-2011 | Phayao | North |
| 34408 | 15 | NA | Pig tonsil | 2010-2011 | Phayao | North |
| 34409 | 16 | NA | Pig tonsil | 2010-2011 | Phayao | North |
| 34410 | 31 | NA | Pig tonsil | 2010-2011 | Phayao | North |
| 34426 | 9 | NA | Pig tonsil | 2010-2011 | Phayao | North |
| 34428 | 3 | NA | Pig tonsil | 2010-2011 | Phayao | North |
| 34429 | 9 | NA | Pig tonsil | 2010-2011 | Phayao | North |
| 34430 | 3 | NA | Pig tonsil | 2010-2011 | Phayao | North |
| 34509 | 5 | NA | Pig tonsil | 2010-2011 | Phayao | North |
| 34522 | 16 | NA | Pig tonsil | 2010-2011 | Phayao | North |
| 34523 | 16 | NA | Pig tonsil | 2010-2011 | Phayao | North |

| | | | | | | |
|---|---|---|---|---|---|---|
| 34525 | 16 | NA | Pig tonsil | 2010-2011 | Phayao | North |
| 34526 | NA | NA | Pig tonsil | 2010-2011 | Phayao | North |
| 34527 | 16 | NA | Pig tonsil | 2010-2011 | Phayao | North |
| 34528 | 9 | NA | Pig tonsil | 2010-2011 | Phayao | North |
| 34529 | 18 | NA | Pig tonsil | 2010-2011 | Phayao | North |
| 34530 | 16 | NA | Pig tonsil | 2010-2011 | Phayao | North |
| 34531 | 16 | NA | Pig tonsil | 2010-2011 | Phayao | North |
| 34532 | 16 | NA | Pig tonsil | 2010-2011 | Phayao | North |
| 34533 | 31 | NA | Pig tonsil | 2010-2011 | Phayao | North |
| 34534 | 16 | NA | Pig tonsil | 2010-2011 | Phayao | North |
| 34535 | 15 | NA | Pig tonsil | 2010-2011 | Phayao | North |
| 34536 | 2 | 299 | Pig tonsil | 2010-2011 | Phayao | North |
| 34537 | 2 | 104 | Pig tonsil | 2010-2011 | Phayao | North |
| 34538 | 16 | NA | Pig tonsil | 2010-2011 | Phayao | North |
| 34539 | 24 | NA | Pig tonsil | 2010-2011 | Phayao | North |
| 34540 | 16 | NA | Pig tonsil | 2010-2011 | Phayao | North |
| 34541 | 24 | NA | Pig tonsil | 2010-2011 | Phayao | North |
| 34542 | NA | NA | Pig tonsil | 2010-2011 | Phayao | North |
| 34544 | NA | NA | Pig tonsil | 2010-2011 | Phayao | North |
| 34545 | 4 | NA | Pig tonsil | 2010-2011 | Phayao | North |
| 34547 | 16 | NA | Pig tonsil | 2010-2011 | Phayao | North |
| 34548 | 16 | NA | Pig tonsil | 2010-2011 | Phayao | North |
| 34549 | NA | NA | Pig tonsil | 2010-2011 | Phayao | North |
| 34550 | 31 | NA | Pig tonsil | 2010-2011 | Phayao | North |
| 34551 | NA | NA | Pig tonsil | 2010-2011 | Phayao | North |
| 34552 | 19 | NA | Pig tonsil | 2010-2011 | Phayao | North |

| | | | | | | |
|---|---|---|---|---|---|---|
| 34553 | 4 | NA | Pig tonsil | 2010-2011 | Phayao | North |
| 34554 | 3 | NA | Pig tonsil | 2010-2011 | Phayao | North |
| 34555 | 16 | NA | Pig tonsil | 2010-2011 | Phayao | North |
| 34556 | 16 | NA | Pig tonsil | 2010-2011 | Phayao | North |
| 34557 | NA | NA | Pig tonsil | 2010-2011 | Phayao | North |
| 34558 | 31 | NA | Pig tonsil | 2010-2011 | Phayao | North |
| 34559 | 9 | NA | Pig tonsil | 2010-2011 | Phayao | North |
| 34560 | 24 | NA | Pig tonsil | 2010-2011 | Phayao | North |
| 34561 | 16 | NA | Pig tonsil | 2010-2011 | Phayao | North |
| 34562 | 31 | NA | Pig tonsil | 2010-2011 | Phayao | North |
| 34563 | 16 | NA | Pig tonsil | 2010-2011 | Phayao | North |
| 34564 | 16 | NA | Pig tonsil | 2010-2011 | Phayao | North |
| 34565 | 16 | NA | Pig tonsil | 2010-2011 | Phayao | North |
| 34566 | 31 | NA | Pig tonsil | 2010-2011 | Phayao | North |
| 34567 | 5 | NA | Pig tonsil | 2010-2011 | Phayao | North |
| 34568 | 16 | NA | Pig tonsil | 2010-2011 | Phayao | North |
| 34569 | 16 | NA | Pig tonsil | 2010-2011 | Phayao | North |
| 34572 | 4 | NA | Pig tonsil | 2010-2011 | Phayao | North |
| 34573 | 28 | NA | Pig tonsil | 2010-2011 | Phayao | North |
| 34574 | 29 | NA | Pig tonsil | 2010-2011 | Phayao | North |
| 34575 | 3 | NA | Pig tonsil | 2010-2011 | Phayao | North |
| 34576 | NA | NA | Pig tonsil | 2010-2011 | Phayao | North |
| 34677 | 7 | NA | Pig tonsil | 2010-2011 | Phayao | North |
| 34678 | 29 | NA | Pig tonsil | 2010-2011 | Phayao | North |
| 34679 | 29 | NA | Pig tonsil | 2010-2011 | Phayao | North |
| 34680 | 9 | NA | Pig tonsil | 2010-2011 | Phayao | North |

| | | | | | | |
|---|---|---|---|---|---|---|
| 34682 | 9 | NA | Pig tonsil | 2010-2011 | Phayao | North |
| 34684 | 31 | NA | Pig tonsil | 2010-2011 | Phayao | North |
| 34685 | 16 | NA | Pig tonsil | 2010-2011 | Phayao | North |
| 34687 | 3 | NA | Pig tonsil | 2010-2011 | Phayao | North |
| 34688 | 7 | NA | Pig tonsil | 2010-2011 | Phayao | North |
| 34691 | 3 | NA | Pig tonsil | 2010-2011 | Phayao | North |
| 34692 | 31 | NA | Pig tonsil | 2010-2011 | Phayao | North |
| 34693 | 4 | NA | Pig tonsil | 2010-2011 | Phayao | North |
| 34697 | 28 | NA | Pig tonsil | 2010-2011 | Phayao | North |
| 34703 | 7 | NA | Pig tonsil | 2010-2011 | Phayao | North |
| 34704 | 4 | NA | Pig tonsil | 2010-2011 | Phayao | North |
| 34705 | 3 | NA | Pig tonsil | 2010-2011 | Phayao | North |
| 35125 | 2 | NA | Pig tonsil | 2010-2011 | Phayao | North |
| 35131 | 29 | NA | Pig tonsil | 2010-2011 | Phayao | North |
| 35150 | 2 | 1 | Pig tonsil | 2010-2011 | Phayao | North |
| 35156 | 15 | NA | Pig tonsil | 2010-2011 | Phayao | North |
| 35407 | 29 | NA | Pig tonsil | 2010-2011 | Phayao | North |
| 35409 | 29 | NA | Pig tonsil | 2010-2011 | Phayao | North |
| 35410 | 29 | NA | Pig tonsil | 2010-2011 | Phayao | North |
| 35412 | 29 | NA | Pig tonsil | 2010-2011 | Phayao | North |
| 35415 | 2 | 28 | Pig tonsil | 2010-2011 | Phayao | North |
| 35538 | 31 | NA | Pig tonsil | 2010-2011 | Phayao | North |
| 35539 | 31 | NA | Pig tonsil | 2010-2011 | Phayao | North |
| 35540 | 31 | NA | Pig tonsil | 2010-2011 | Phayao | North |
| 35541 | 1 | 237 | Pig tonsil | 2010-2011 | Phayao | North |
| 35542 | 31 | NA | Pig tonsil | 2010-2011 | Phayao | North |

| | | | | | | |
|---|---|---|---|---|---|---|
| 35543 | 31 | NA | Pig tonsil | 2010-2011 | Phayao | North |
| 35544 | 31 | NA | Pig tonsil | 2010-2011 | Phayao | North |
| 35545 | 31 | NA | Pig tonsil | 2010-2011 | Phayao | North |
| 35546 | 11 | NA | Pig tonsil | 2010-2011 | Phayao | North |
| 35781 | 14 | 1 | Pig tonsil | 2010-2011 | Phayao | North |

**Supplementary Table 4**. Metadata regarding the 264 Thai S. suis isolates used for genomic analysis during this study. The dataset contains 141 new genomes generated during this study and 123 publicly available Thai S. suis genomes. Na stands for not available. CC stands for clonal complex. ST stands for sequence type.

| **Isolate** | **Year** | **Province** | **Region** | **Serotype** | **CC** | **ST** | **Host** | **Clinical Manifestation** | **Biosample** | **Reference** |
|---|---|---|---|---|---|---|---|---|---|---|
| 21928 | 2006 | Lampang | North | 14 | 1 | 105 | Human | Meningitis | NA | [25] |
| 25779 | 2007 | Tak | North | 14 | 1 | 105 | Human | Sepsis | NA | [25] |
| 26390 | 2008 | Bangkok | Central | 14 | 1 | 105 | Human | Meningitis | NA | [25] |
| 27024 | 2008 | Lampang | North | 14 | 1 | 127 | Human | Meningitis | NA | [25] |
| 28075 | 2008 | Phitsanulok | North | 14 | 1 | 237 | Human | Sepsis | NA | [25] |
| 30333 | 2009 | Tak | North | 14 | 1 | 105 | Human | Spondylodiscitis | NA | [25] |
| 31998 | 2010 | Phetchabun | North | 14 | 1 | 105 | Human | Meningitis | NA | [25] |
| 32481 | 2010 | Phetchabun | North | 14 | 1 | 105 | Human | Sepsis | NA | [25] |
| 32516 | 2010 | Phetchabun | North | 14 | 1 | 105 | Human | Meningitis | NA | [25] |
| 35728 | 2011 | Nakhon Ratchasima | Northeast | 14 | 1 | 105 | Human | Sepsis | NA | [25] |
| 42841 | 2012 | Uttaradit | North | 14 | 1 | 105 | Human | Sepsis & Meningitis | NA | [25] |
| SS-194 | 2018 | Ubon Ratchathani | Northeast | 7 | 104 | 1726 | Pig Diseased | Respiratory | NA | [26] |
| SS-230 | 2018 | Prachin Buri | Central | 7 | 94 | 373 | Pig Diseased | Respiratory | NA | [26] |
| SS-236 | 2018 | Chachoengsao | Central | 7 | 94 | 373 | Pig Diseased | Respiratory | NA | [26] |
| SS-500 | 2020 | Nakhon Pathom | Central | 7 | 104 | 104 | Pig Diseased | Respiratory | NA | [26] |
| SS-560 | 2020 | Ratchaburi | Central | 7 | NA | 1809 | Pig Diseased | Respiratory | NA | [26] |
| STC407 | 2021 | Nakhon Ratchasima | | 2 | 233 | 1713 | Human | Sepsis | NA | [27] |
| MNCM21 | 1998 | Chiang Mai | North | 2 | 104 | 101 | Human | Meningitis | SAMD00009282 | NA |
| MNCM50 | 2002 | Chiang Mai | North | 2 | 104 | 104 | Human | Pulmonary edema | SAMD00009289 | NA |
| MNCM01 | 2000 | Chiang Mai | North | 2 | 1 | 1 | Human | Endocarditis | SAMD00039895 | NA |
| MNCM02 | 2000 | Chiang Mai | North | 2 | 25 | 25 | Human | NA | SAMD00039896 | NA |
| MNCM03 | 2000 | Chiang Mai | North | 2 | 25 | 25 | Human | NA | SAMD00039897 | NA |

| | | | | | | | | | | |
|---|---|---|---|---|---|---|---|---|---|---|
| MNCM06 | 2000 | Chiang Mai | North | 2 | 1 | 1 | Human | Meningitis | SAMD00039898 | NA |
| MNCM11 | 2001 | Chiang Mai | North | 2 | 25 | 25 | Human | NA | SAMD00039899 | NA |
| MNCM13 | 2001 | Chiang Mai | North | 2 | 25 | 25 | Human | NA | SAMD00039900 | NA |
| MNCM16 | 2000 | Chiang Mai | North | 2 | 1 | 1 | Human | Meningitis | SAMD00039901 | NA |
| MNCM33 | 2002 | Chiang Mai | North | 2 | 25 | 103 | Human | Meningitis | SAMD00039902 | NA |
| MNCM36 | 2002 | Chiang Mai | North | 2 | 25 | 103 | Human | NA | SAMD00039903 | NA |
| MNCM38 | 2002 | Chiang Mai | North | 2 | 25 | 3040 | Human | NA | SAMD00039904 | NA |
| MNCM52 | 2002 | Chiang Mai | North | 2 | 25 | NA | Human | NA | SAMD00039905 | NA |
| MNCM54 | 2002 | Chiang Mai | North | 2 | 25 | 102 | Human | Meningitis | SAMD00039906 | NA |
| LPH03 | 2001 | Lamphun | North | 2 | 25 | 103 | Human | Meningitis | SAMD00039907 | NA |
| LPH05 | 2001 | Lamphun | North | 2 | 25 | 103 | Human | Sepsis | SAMD00039908 | NA |
| MNCM17 | 2000 | Chiang Mai | North | 2 | 104 | 104 | Human | NA | SAMD00039909 | NA |
| MNCM34 | 2002 | Chiang Mai | North | 2 | 25 | 25 | Human | NA | SAMD00039910 | NA |
| MNCM39 | 2002 | Chiang Mai | North | 2 | 28 | 28 | Human | NA | SAMD00039911 | NA |
| DTU_2020_1000565_1_SI_THA_SAR_056 | 2020 | Saraburi | Central | 2 | 233 | 3041 | Human | NA | SAMEA111506260 | [28] |
| NSUI036 | 2002 | NA | NA | 2 | 28 | 28 | Human | NA | SAMN03652189 | [6] |
| NSUI033 | 2000 | NA | NA | 2 | 25 | 25 | Human | Meningitis | SAMN04231436 | [5] |
| NSUI034 | 2000 | NA | NA | 2 | 25 | 25 | Human | Sepsis | SAMN04231437 | [5] |
| NSUI035 | 2001 | NA | NA | 2 | 25 | 25 | Human | Meningitis | SAMN04231438 | [5] |
| NSUI037 | 2002 | NA | NA | 2 | 25 | 25 | Human | Sepsis | SAMN04231439 | [5] |
| NSUI038 | 2002 | NA | NA | 2 | 25 | 25 | Human | Sepsis | SAMN04231440 | [5] |
| NSUI039 | 2001 | NA | NA | 2 | 25 | 25 | Human | Sepsis | SAMN04231441 | [5] |
| NSUI040 | 2002 | NA | NA | 2 | 25 | 25 | Human | Sepsis | SAMN04231442 | [5] |
| NSUI092 | 2001 | NA | NA | 2 | 25 | 25 | Human | Endocarditis | SAMN04231474 | [5] |
| 1584695 | 2013 | NA | NA | 9 | 16 | 16 | Human | Sepsis | SAMN05919918 | [7] |
| HU_SS02 | 2007 | Phayao | North | 2 | 1 | 1 | Human | NA | SAMN12911088 | [2] |

| | | | | | | | | | | |
|---|---|---|---|---|---|---|---|---|---|---|
| DP_SS20 | 2007 | Nakhon Si Thammarat | South | 2 | 1 | 1 | Pig Diseased | Sepsis | SAMN12911089 | [2] |
| AP_SS21 | 2012 | Phayao | North | NT | NA | NA | Pig Healthy | Healthy | SAMN12911090 | [2] |
| AP_SS24 | 2007 | Phayao | North | 2 | 1 | 1 | Pig Healthy | Healthy | SAMN12911092 | [2] |
| AP_SS25 | 2007 | Phayao | North | 2 | 1 | 1 | Pig Healthy | Healthy | SAMN12911093 | [2] |
| DP_SS26 | 2007 | Nakhon Pathom | Central | 2 | 1 | 1 | Pig Diseased | Sepsis | SAMN12911094 | [2] |
| DP_SS28 | 2007 | Nakhon Pathom | Central | 2 | 104 | 104 | Pig Diseased | Sepsis | SAMN12911095 | [2] |
| DP_SS29 | 2012 | Nakhon Pathom | Central | 2 | 25 | 25 | Pig Diseased | Sepsis | SAMN12911096 | [2] |
| HU_SS30 | 2007 | Phayao | North | 2 | 104 | 104 | Human | Sepsis | SAMN13172488 | [2] |
| 33329 | 2010 | Phetchabun | North | 24 | 221/234 | 221 | Human | Sepsis | SAMN17277757 | [29] |
| 32098 | 2010 | Lampang | North | 24 | 221/234 | 234 | Human | Sepsis | SAMN17277758 | [29] |
| 32494 | 2010 | Phetchabun | North | 14 | 1 | 105 | Human | Sepsis | SAMN17277759 | [30] |
| 30190 | 2009 | Phitsanulok | North | 2 | 25 | 25 | Human | Meningitis | SAMN17277760 | [30] |
| 27715 | 2008 | Ratchaburi | Central | 2 | 104 | 104 | Human | Sepsis | SAMN17277761 | [30] |
| 40747 | 2012 | Phitsanulok | North | 2 | 233 | 233 | Human | Sepsis | SAMN17277762 | [30] |
| 41126 | 2012 | Nakhon Ratchasima | Northeas t | 2 | 233 | 233 | Human | Meningitis | SAMN17277763 | [30] |
| 40085 | 2012 | Lampang | North | 2 | 233 | 379 | Human | Sepsis | SAMN17277764 | [30] |
| 40484 | 2012 | Phitsanulok | North | 2 | 233 | 233 | Human | Meningitis | SAMN17277765 | [30] |
| 37647 | 2011 | Lampang | North | 2 | 233 | 233 | Human | Sepsis | SAMN17277766 | [30] |
| 24525 | 2007 | Phetchabun | North | 2 | 104 | 104 | Human | Sepsis | SAMN18585446 | [30] |
| 26154 | 2007 | Phetchabun | North | 2 | 1 | 1 | Human | Meningitis | SAMN18585447 | [30] |
| 39565 | 2012 | Bangkok | Central | 24 | 221/234 | 221 | Human | Sepsis | SAMN19600291 | [29] |
| 41570 | 2020 | Phayao | North | 5 | 221/234 | 221 | Human | Sepsis | SAMN20090113 | [29] |
| STC78 | 2021 | Nakhon Ratchasima | Northeas t | 2 | 233 | 1656 | Human | Sepsis | SAMN29401318 | [1] |
| STC127 | 2021 | Nakhon Ratchasima | Northeas t | 2 | 1688 | 1688 | Human | Sepsis | SAMN29401333 | [1] |
| STC129 | 2021 | Nakhon Ratchasima | Northeas t | 2 | 1688 | 1688 | Human | Sepsis & Meningitis | SAMN29401334 | [1] |

| | | | | | | | | | | |
|---|---|---|---|---|---|---|---|---|---|---|
| STC130 | 2021 | Nakhon Ratchasima | Northeas t | 2 | 233 | 379 | Human | Sepsis | SAMN29401335 | 1 |
| STC131 | 2021 | Nakhon Ratchasima | Northeas t | 2 | 233 | 233 | Human | Sepsis | SAMN29401336 | 1 |
| STC152 | 2021 | Nakhon Ratchasima | Northeas t | 2 | 104 | 104 | Human | Meningitis | SAMN29401337 | 1 |
| STC153 | 2021 | Nakhon Ratchasima | Northeas t | 2 | 233 | 233 | Human | NA | SAMN29401338 | 1 |
| STC154 | 2021 | Nakhon Ratchasima | Northeas t | 2 | 1688 | 1688 | Human | Spondylodiscitis | SAMN29401339 | 1 |
| 8789_SUI_1 | 2015 | Chiang Mai | North | 16 | NA | 1923 | Pig Healthy | Healthy | SAMN29502508 | 3 |
| 8790_SUI_2 | 2015 | Chiang Mai | North | 16 | NA | 1923 | Pig Healthy | Healthy | SAMN29502509 | 3 |
| 8797_SUI_4_1 | 2015 | Chiang Mai | North | NT | NA | 1924 | Pig Healthy | Healthy | SAMN29502515 | 3 |
| 8798_SUI_4_2 | 2015 | Chiang Mai | North | 4 | NA | 1924 | Pig Healthy | Healthy | SAMN29502516 | 3 |
| 8799_SUI_4_3 | 2015 | Chiang Mai | North | 4 | NA | 1924 | Pig Healthy | Healthy | SAMN29502517 | 3 |
| 8811_SUI_9_1 | 2015 | Chiang Mai | North | 16 | NA | 1925 | Pig Healthy | Healthy | SAMN29502527 | 3 |
| 8813_SUI_10 | 2015 | Chiang Mai | North | 16 | NA | 1925 | Pig Healthy | Healthy | SAMN29502528 | 3 |
| 8814_SUI_11 | 2015 | Chiang Mai | North | NT | NA | 1926 | Pig Healthy | Healthy | SAMN29502529 | 3 |
| 8815_SUI_12 | 2015 | Chiang Mai | North | 11 | NA | 1926 | Pig Healthy | Healthy | SAMN29502530 | 3 |
| 8816_SUI_13 | 2015 | Chiang Mai | North | NA | NA | 1927 | Pig Healthy | Healthy | SAMN29502531 | 3 |
| 8818_SUI_15 | 2015 | Chiang Mai | North | NA | NA | 1928 | Pig Healthy | Healthy | SAMN29502533 | 3 |
| 8819_SUI_16 | 2015 | Chiang Mai | North | NA | NA | 1929 | Pig Healthy | Healthy | SAMN29502534 | 3 |
| 8820_SUI_17 | 2015 | Chiang Mai | North | NA | NA | 1929 | Pig Healthy | Healthy | SAMN29502535 | 3 |
| 8822_SUI_19 | 2015 | Chiang Mai | North | NA | NA | 1929 | Pig Healthy | Healthy | SAMN29502537 | 3 |
| 8823_SUI_20 | 2015 | Chiang Mai | North | NA | NA | 1930 | Pig Healthy | Healthy | SAMN29502538 | 3 |
| 8826_SUI_24 | 2015 | Chiang Mai | North | NA | NA | 1931 | Pig Healthy | Healthy | SAMN29502541 | 3 |
| 8827_SUI_25 | 2015 | Chiang Mai | North | NT | NA | 1932 | Pig Healthy | Healthy | SAMN29502542 | 3 |
| 8835_SUI_33 | 2015 | Chiang Mai | North | 16 | NA | 1933 | Pig Healthy | Healthy | SAMN29502550 | 3 |
| 8845_SUI_42 | 2015 | Chiang Mai | North | NT | NA | 1934 | Pig Healthy | Healthy | SAMN29502560 | 3 |
| 8846_SUI_43 | 2015 | Chiang Mai | North | NT | NA | 1934 | Pig Healthy | Healthy | SAMN29502561 | 3 |

| | | | | | | | | | | |
|---|---|---|---|---|---|---|---|---|---|---|
| 8852_SUI_50 | 2015 | Chiang Mai | North | 29 | NA | NA | Pig Healthy | Healthy | SAMN29502567 | [3] |
| 8853_SUI_51 | 2015 | Chiang Mai | North | 30 | NA | 1935 | Pig Healthy | Healthy | SAMN29502568 | [3] |
| 8854_SUI_52 | 2015 | Chiang Mai | North | 4 | NA | 1936 | Pig Healthy | Healthy | SAMN29502569 | [3] |
| 8858_SUI_56 | 2015 | Chiang Mai | North | 31 | NA | 1937 | Pig Healthy | Healthy | SAMN29502573 | [3] |
| 8859_SUI_57 | 2015 | Chiang Mai | North | NT | NA | 3042 | Pig Healthy | Healthy | SAMN29502574 | [3] |
| 8860_SUI_H132_53 | 2010 | Chiang Mai | North | 2 | 104 | 104 | Human | Meningitis | SAMN29502575 | [3] |
| 8861_SUI_H219_53 | 2010 | Chiang Mai | North | 2 | 1 | 105 | Human | Meningitis | SAMN29502576 | [3] |
| 8862_SUI_T2NE | 2013 | Chiang Mai | North | 2 | 1 | 1 | Pig Diseased | Sepsis | SAMN29502577 | [3] |
| 8863_SUI_T2N | 2013 | Chiang Mai | North | 2 | 1 | 1 | Pig Diseased | Sepsis | SAMN29502578 | [3] |
| 8864_SUI_THD10_6 | 2013 | Chiang Mai | North | 31 | NA | 1938 | Pig Healthy | Healthy | SAMN29502579 | [3] |
| 8865_SUI_TSK11_2 | 2013 | Chiang Mai | North | 2 | 28 | 28 | Pig Healthy | Healthy | SAMN29502580 | [3] |
| 8866_SUI_TSK16_2 | 2013 | Chiang Mai | North | 2 | 25 | 1939 | Pig Healthy | Healthy | SAMN29502581 | [3] |
| 8867_SUI_TP12_2 | 2013 | Chiang Mai | North | 2 | 104 | 104 | Pig Healthy | Healthy | SAMN29502582 | [3] |
| 8868_SUI_TP12_3 | 2013 | Chiang Mai | North | 2 | 104 | 104 | Pig Healthy | Healthy | SAMN29502583 | [3] |
| 8869_SUI_T13_1N | 2013 | Chiang Mai | North | 28 | NA | 1940 | Pig Healthy | Healthy | SAMN29502584 | [3] |
| 8871_SUI_TJ21_2S | 2013 | Chiang Mai | North | NT | NA | 3031 | Pig Healthy | Healthy | SAMN29502585 | [3] |
| 35541 | 2011 | Phayao | North | 1 | 1 | 237 | Pig Healthy | Healthy | SAMN31277171 | [31] |
| 38828 | 2011 | Chanthaburi | East | 1 | 1 | 105 | Human | Sepsis | SAMN31277172 | [31] |
| 34572 | 2011 | Phayao | North | 4 | 94 | 1689 | Pig Healthy | Healthy | SAMN31277173 | [32] |
| 36054 | 2011 | Lampang | North | 4 | 94 | 94 | Human | Sepsis | SAMN31277174 | [32] |
| TRG6 | 2010 | Ratchaburi | Central | 4 | 94 | 94 | Pig Diseased | Pneumonia | SAMN31277175 | [32] |
| 34545 | 2011 | Phayao | North | 4 | 4 | 1689 | Pig Healthy | Healthy | SAMN31277176 | [32] |
| 34553 | 2011 | Phayao | North | 4 | 94 | 1689 | Pig Healthy | Healthy | SAMN31277177 | [32] |
| 34693 | 2011 | Phayao | North | 4 | 94 | 94 | Pig Healthy | Healthy | SAMN31277178 | [32] |
| 34704 | 2011 | Phayao | North | 4 | 94 | 94 | Pig Healthy | Healthy | SAMN31277179 | [32] |
| 22696 | 2006 | Uttaradit | North | 2 | 104 | 104 | Human | Spondylodiscitis | SAMN46258697 | This study |

| | | | | | | | | | | |
|---|---|---|---|---|---|---|---|---|---|---|
| 22714 | 2006 | Nan | North | 2 | 25 | 25 | Human | Sepsis | SAMN46258698 | This study |
| 23163 | 2006 | Nan | North | 2 | 104 | 104 | Human | Meningitis | SAMN46258699 | This study |
| 23164 | 2006 | Lampang | North | 2 | 28 | 28 | Human | Meningitis | SAMN46258700 | This study |
| 23167 | 2006 | Chon Buri | East | 2 | 1 | 1 | Human | Meningitis | SAMN46258701 | This study |
| 24205 | 2007 | Lampoon | North | 2 | 25 | 25 | Human | Sepsis | SAMN46258702 | This study |
| 24526 | 2007 | Lampang | North | 2 | 1 | 126 | Human | Meningitis | SAMN46258703 | This study |
| 24529 | 2007 | Uttaradit | North | 2 | 104 | 104 | Human | Meningitis | SAMN46258704 | This study |
| 25777 | 2007 | Nan | North | 2 | 104 | 104 | Human | Sepsis | SAMN46258705 | This study |
| 25781 | 2007 | Saraburi | Central | 2 | 25 | 103 | Human | Meningitis | SAMN46258706 | This study |
| 26127 | 2008 | Lampang | North | 2 | 104 | 104 | Human | Meningitis | SAMN46258707 | This study |
| 26647 | 2008 | Nakhon Ratchasima | Northeast | 2 | 104 | 104 | Human | Meningitis | SAMN46258708 | This study |
| 27103 | 2008 | Tak | North | 2 | 25 | 25 | Human | Sepsis | SAMN46258709 | This study |
| 27964 | 2008 | Sukhothai | North | 14 | 1 | 105 | Human | Sepsis | SAMN46258710 | This study |
| 27968 | 2008 | Sa Kaeo | East | 2 | 233 | 233 | Human | Sepsis | SAMN46258711 | This study |
| 28074 | 2008 | Phitsanulok | North | 14 | 1 | 105 | Human | Sepsis | SAMN46258712 | This study |
| 28739 | 2008 | Nan | North | 2 | 28 | 28 | Human | Sepsis | SAMN46258713 | This study |
| 29281 | 2008 | Phayao | North | 2 | 104 | 104 | Human | Sepsis | SAMN46258714 | This study |
| 30198 | 2009 | Sukhothai | North | 14 | 1 | 105 | Human | Sepsis & Meningitis | SAMN46258715 | This study |
| 30341 | 2009 | Nan | North | 2 | 104 | 104 | Human | Sepsis | SAMN46258716 | This study |
| 30346 | 2009 | Phitsanulok | North | 2 | 233 | 233 | Human | Endocarditis | SAMN46258717 | This study |
| 30389 | 2009 | Sa Kaeo | East | 2 | 233 | 233 | Human | Sepsis | SAMN46258718 | This study |
| 30823 | 2009 | Chiang Mai | North | 2 | 233 | 233 | Human | Sepsis | SAMN46258719 | This study |
| 30827 | 2009 | Phrae | North | 2 | 1 | 1 | Human | ARDS | SAMN46258720 | This study |
| 31095 | 2009 | Phetchabun | North | 2 | 25 | 103 | Human | Sepsis | SAMN46258721 | This study |
| 31106 | 2009 | Phetchabun | North | 2 | 104 | 104 | Human | Sepsis | SAMN46258722 | This study |

| | | | | | | | | | | |
|---|---|---|---|---|---|---|---|---|---|---|
| 32599 | 2010 | Buri Ram | Northeast | 2 | 1 | 11 | Human | Sepsis | SAMN46258723 | This study |
| 32861 | 2010 | Phetchabun | North | 2 | 104 | 104 | Human | Sepsis | SAMN46258724 | This study |
| 32941 | 2010 | Lampang | North | 14 | 1 | 105 | Human | Sepsis | SAMN46258725 | This study |
| 33004 | 2010 | Lampang | North | 2 | 104 | 104 | Human | Sepsis | SAMN46258726 | This study |
| 33029 | 2010 | Phetchabun | North | 2 | 104 | 104 | Human | Sepsis | SAMN46258727 | This study |
| 33243 | 2010 | Udon Thani | Northeast | 2 | 25 | 381 | Human | Sepsis | SAMN46258728 | This study |
| 33394 | 2010 | Bangkok | Central | 2 | 233 | 233 | Human | Sepsis | SAMN46258729 | This study |
| 34384 | 2010 | Phayao | North | 2 | 1 | 2 | Pig Healthy | Healthy | SAMN46258730 | This study |
| 34391 | 2010 | Phayao | North | 31 | NA | 2759 | Pig Healthy | Healthy | SAMN46258731 | This study |
| 34408 | 2010 | Phayao | North | 15 | NA | 3020 | Pig Healthy | Healthy | SAMN46258732 | This study |
| 34428 | 2010 | Phayao | North | 3 | 28 | 117 | Pig Healthy | Healthy | SAMN46258733 | This study |
| 34429 | 2010 | Phayao | North | 9 | NA | 301 | Pig Healthy | Healthy | SAMN46258734 | This study |
| 34533 | 2010 | Phayao | North | 31 | NA | 2763 | Pig Healthy | Healthy | SAMN46258735 | This study |
| 34536 | 2010 | Phayao | North | 2 | 28 | 299 | Pig Healthy | Healthy | SAMN46258736 | This study |
| 34537 | 2010 | Phayao | North | 2 | 104 | 104 | Pig Healthy | Healthy | SAMN46258737 | This study |
| 34538 | 2010 | Phayao | North | 16 | NA | 3021 | Pig Healthy | Healthy | SAMN46258738 | This study |
| 34552 | 2010 | Phayao | North | 19 | NA | 3034 | Pig Healthy | Healthy | SAMN46258739 | This study |
| 34555 | 2010 | Phayao | North | 16 | NA | 3026 | Pig Healthy | Healthy | SAMN46258740 | This study |
| 34560 | 2010 | Phayao | North | 24 | NA | 3038 | Pig Healthy | Healthy | SAMN46258741 | This study |
| 34584 | 2011 | Rayong | East | 2 | 29 | 515 | Human | Sepsis | SAMN46258742 | This study |
| 34677 | 2010 | Phayao | North | 7 | NA | 302 | Pig Healthy | Healthy | SAMN46258743 | This study |
| 34679 | 2010 | Phayao | North | 29 | NA | 3028 | Pig Healthy | Healthy | SAMN46258744 | This study |
| 34688 | 2010 | Phayao | North | 7 | NA | 302 | Pig Healthy | Healthy | SAMN46258745 | This study |
| 34703 | 2010 | Phayao | North | 7 | NA | 302 | Pig Healthy | Healthy | SAMN46258746 | This study |
| 34758 | 2011 | Buri Ram | Northeast | 2 | 25 | 380 | Human | Sepsis | SAMN46258747 | This study |

| | | | | | | | | | | |
|---|---|---|---|---|---|---|---|---|---|---|
| 35125 | 2010 | Phayao | North | 2 | 28 | 28 | Pig Healthy | Healthy | SAMN46258748 | This study |
| 35150 | 2010 | Phayao | North | 2 | 1 | 1 | Pig Healthy | Healthy | SAMN46258749 | This study |
| 35415 | 2010 | Phayao | North | 2 | 28 | 28 | Pig Healthy | Healthy | SAMN46258750 | This study |
| 35688 | 2011 | Rayong | East | 2 | 233 | 233 | Human | Sepsis | SAMN46258751 | This study |
| 35736 | 2011 | Phayao | North | 2 | 1 | 298 | Human | Sepsis | SAMN46258752 | This study |
| 35781 | 2010 | Phayao | North | 14 | 1 | 1 | Pig Healthy | Healthy | SAMN46258753 | This study |
| 35896 | 2011 | Phetchabun | North | 2 | 1 | 105 | Human | Meningitis | SAMN46258754 | This study |
| 35940 | 2011 | Phetchabun | North | 2 | 233 | 233 | Human | Sepsis | SAMN46258755 | This study |
| 35949 | 2011 | Phetchaburi | Central | 2 | 233 | 233 | Human | Sepsis | SAMN46258756 | This study |
| 36363 | 2011 | Nakhon Phanom | Northeast | 2 | 1 | 144 | Human | Sepsis | SAMN46258757 | This study |
| 36369 | 2011 | Lampang | North | 2 | 104 | 512 | Human | Sepsis | SAMN46258758 | This study |
| 36375 | 2011 | Nong Khai | Northeast | 2 | 25 | 395 | Human | Sepsis | SAMN46258759 | This study |
| 36602 | 2011 | Lampang | North | 2 | 233 | 379 | Human | NA | SAMN46258760 | This study |
| 36610 | 2011 | Chiang Mai | North | 2 | 104 | 391 | Human | Sepsis | SAMN46258761 | This study |
| 36763 | 2011 | Uttaradit | North | NT | 28 | 28 | Human | Sepsis | SAMN46258762 | This study |
| 36900 | 2011 | Nan | North | 2 | 29 | 516 | Human | Sepsis | SAMN46258763 | This study |
| 36921 | 2011 | Suphanburi | Central | 2 | 104 | 104 | Human | Sepsis | SAMN46258764 | This study |
| 37554 | 2011 | Chiang Mai | North | 2 | 233 | 233 | Human | Sepsis | SAMN46258765 | This study |
| 37880 | 2011 | Lampang | North | 2 | 233 | 379 | Human | Sepsis | SAMN46258766 | This study |
| 38727 | 2011 | Lampang | North | 2 | 233 | 233 | Human | Sepsis | SAMN46258767 | This study |
| 38826 | 2011 | Phetchabun | North | 14 | 1 | 105 | Human | Sepsis | SAMN46258768 | This study |
| 39236 | 2012 | Phitsanulok | North | 2 | 233 | 233 | Human | Sepsis | SAMN46258769 | This study |
| 39825 | 2012 | Phetchabun | North | 14 | 1 | 105 | Human | Meningitis | SAMN46258770 | This study |
| 39957 | 2012 | Uttaradit | North | 14 | 1 | 105 | Human | Sepsis | SAMN46258771 | This study |
| 39971 | 2012 | Nan | North | 2 | 104 | 512 | Human | Sepsis | SAMN46258772 | This study |

| | | | | | | | | | | |
|---|---|---|---|---|---|---|---|---|---|---|
| 40128 | 2012 | Nakhon Phanom | Northeas t | 2 | 1 | 337 | Human | Sepsis | SAMN46258773 | This study |
| 40133 | 2012 | Phitsanulok | North | 2 | 104 | 513 | Human | Sepsis | SAMN46258774 | This study |
| 40301 | 2012 | Phitsanulok | North | 2 | 233 | 233 | Human | Sepsis | SAMN46258775 | This study |
| 40302 | 2012 | Phitsanulok | North | 2 | 233 | 393 | Human | Sepsis | SAMN46258776 | This study |
| 40414 | 2012 | Saraburi | Central | 2 | 104 | 514 | Human | Sepsis | SAMN46258777 | This study |
| 40417 | 2012 | Saraburi | Central | 2 | 25 | 25 | Human | Sepsis | SAMN46258778 | This study |
| 40614 | 2012 | Buri Ram | Northeas t | 2 | 1 | 1 | Human | Sepsis | SAMN46258779 | This study |
| 40621 | 2012 | Cachoengsao | East | 2 | 104 | 392 | Human | Sepsis | SAMN46258780 | This study |
| 40750 | 2012 | Phitsanulok | North | 2 | 233 | 233 | Human | Sepsis | SAMN46258781 | This study |
| 40804 | 2012 | Nonthaburi | Central | 2 | 233 | 233 | Human | Sepsis | SAMN46258782 | This study |
| 41219 | 2012 | Sa Kaeo | East | 2 | 233 | 233 | Human | Sepsis | SAMN46258783 | This study |
| 41445 | 2012 | Tak | North | 2 | 1 | 105 | Human | Sepsis | SAMN46258784 | This study |
| 41477 | 2012 | Nan | North | 2 | 233 | 233 | Human | Sepsis | SAMN46258785 | This study |
| 41594 | 2012 | Phetchabun | North | 2 | NA | 236 | Human | Sepsis | SAMN46258786 | This study |
| 42319 | 2012 | Phitsanulok | North | 2 | 28 | 382 | Human | Sepsis | SAMN46258787 | This study |
| 42527 | 2012 | Phitsanulok | North | 2 | 233 | 233 | Human | Sepsis | SAMN46258788 | This study |
| STC123 | 2021 | Phetchabun | North | 2 | 1 | 105 | Human | NA | SAMN46258789 | This study |
| STC135 | 2021 | Nakhon Ratchasima | Northeas t | 2 | 233 | 379 | Human | NA | SAMN46258790 | This study |
| STC141 | 2021 | Phetchabun | North | 2 | 233 | 233 | Human | NA | SAMN46258791 | This study |
| STC147 | 2021 | Chiang Mai | North | 2 | 25 | 25 | Human | NA | SAMN46258792 | This study |
| STC149 | 2021 | Phetchabun | North | 14 | 233 | 3032 | Human | NA | SAMN46258793 | This study |
| STC173 | 2021 | Nakhon Ratchasima | Northeas t | 2 | 233 | 233 | Human | NA | SAMN46258794 | This study |
| STC19 | 2021 | Roi Et | Northeas t | 2 | 1688 | 1688 | Human | NA | SAMN46258795 | This study |
| STC212 | 2021 | Nakhon Ratchasima | Northeas t | 2 | 233 | 233 | Human | NA | SAMN46258796 | This study |
| STC217 | 2013 | Chiang Mai | North | 2 | 1 | 1 | Human | NA | SAMN46258797 | This study |

| | | | | | | | | | | |
|---|---|---|---|---|---|---|---|---|---|---|
| STC223 | 2014 | Phrae | North | 2 | 233 | 233 | Human | NA | SAMN46258798 | This study |
| STC224 | 2014 | Phrae | North | 2 | 1 | 1 | Human | NA | SAMN46258799 | This study |
| STC228 | 2014 | Phrae | North | 2 | 104 | 104 | Human | NA | SAMN46258800 | This study |
| STC231 | 2015 | Chiang Mai | North | 2 | 1 | 1 | Human | NA | SAMN46258801 | This study |
| STC233 | 2015 | Chiang Mai | North | 2 | 233 | 233 | Human | NA | SAMN46258802 | This study |
| STC235 | 2015 | Chiang Mai | North | 2 | 233 | 233 | Human | NA | SAMN46258803 | This study |
| STC238 | 2016 | Chiang Mai | North | 2 | 233 | 233 | Human | NA | SAMN46258804 | This study |
| STC239 | 2016 | Chiang Mai | North | 2 | 233 | 233 | Human | NA | SAMN46258805 | This study |
| STC240 | 2016 | Chiang Mai | North | 2 | 233 | 379 | Human | NA | SAMN46258806 | This study |
| STC242 | 2016 | Lamphun | North | 2 | 104 | 104 | Human | NA | SAMN46258807 | This study |
| STC243 | 2016 | Lamphun | North | 2 | 233 | 233 | Human | NA | SAMN46258808 | This study |
| STC244 | 2017 | Phrae | North | 2 | 104 | 104 | Human | NA | SAMN46258809 | This study |
| STC246 | 2017 | Phrae | North | 2 | 104 | 104 | Human | NA | SAMN46258810 | This study |
| STC247 | 2017 | Chiang Mai | North | 2 | 1 | 1 | Human | NA | SAMN46258811 | This study |
| STC250 | 2017 | Chiang Mai | North | 2 | 233 | 233 | Human | NA | SAMN46258812 | This study |
| STC251 | 2019 | Chiang Mai | North | 2 | 104 | 104 | Human | NA | SAMN46258813 | This study |
| STC253 | 2019 | Chiang Mai | North | 2 | 104 | 104 | Human | NA | SAMN46258814 | This study |
| STC256 | 2019 | Chiang Mai | North | 2 | 104 | 104 | Human | NA | SAMN46258815 | This study |
| STC257 | 2019 | Chiang Mai | North | 2 | 233 | 379 | Human | NA | SAMN46258816 | This study |
| STC260 | 2021 | Nakhon Ratchasima | Northeast | 2 | 233 | 1656 | Human | NA | SAMN46258817 | This study |
| STC262 | 2021 | Nakhon Ratchasima | Northeast | 2 | 104 | 104 | Human | NA | SAMN46258818 | This study |
| STC266 | 2021 | Nakhon Ratchasima | Northeast | 2 | 104 | 104 | Human | NA | SAMN46258819 | This study |
| STC267 | 2021 | Nakhon Ratchasima | Northeast | 2 | 104 | 104 | Human | NA | SAMN46258820 | This study |
| STC268 | 2021 | Nakhon Ratchasima | Northeast | 2 | 104 | 104 | Human | NA | SAMN46258821 | This study |
| STC269 | 2021 | Nakhon Ratchasima | Northeast | 2 | 104 | 104 | Human | NA | SAMN46258822 | This study |

| | | | | | | | | | | |
|---|---|---|---|---|---|---|---|---|---|---|
| STC270 | 2021 | Nakhon Ratchasima | Northeas t | 2 | 104 | 104 | Human | NA | SAMN46258823 | This study |
| STC274 | 2019 | Chiang Mai | North | 2 | 233 | 379 | Human | NA | SAMN46258824 | This study |
| STC275 | 2019 | Chiang Mai | North | 2 | 1 | 1 | Human | NA | SAMN46258825 | This study |
| STC279 | 2020 | Lampang | North | 2 | 233 | 233 | Human | NA | SAMN46258826 | This study |
| STC280 | 2020 | Lampang | North | 2 | 233 | 379 | Human | NA | SAMN46258827 | This study |
| STC282 | 2020 | Lampang | North | 2 | 104 | 104 | Human | NA | SAMN46258828 | This study |
| STC283 | 2020 | Lampang | North | 2 | 104 | 104 | Human | NA | SAMN46258829 | This study |
| STC286 | 2020 | Lampang | North | 2 | 233 | 233 | Human | NA | SAMN46258830 | This study |
| STC290 | 2020 | Lampang | North | 2 | 104 | 104 | Human | NA | SAMN46258831 | This study |
| STC291 | 2020 | Lampang | North | 2 | 104 | 104 | Human | NA | SAMN46258832 | This study |
| STC292 | 2020 | Lampang | North | 2 | 104 | 104 | Human | NA | SAMN46258833 | This study |
| STC293 | 2020 | Chiang Mai | North | 2 | 1 | 105 | Human | NA | SAMN46258834 | This study |
| STC31 | 2021 | Phetchabun | North | 2 | 233 | 3041 | Human | NA | SAMN46258835 | This study |
| STC7 | 2019 | Phetchabun | North | 14 | 1 | 11 | Human | NA | SAMN46258836 | This study |
| STC72 | 2021 | Roi Et | Northeas t | 2 | 104 | 104 | Human | NA | SAMN46258837 | This study |

**Supplementary Table 5**. Metadata of 2909 genomes from our Global collection. NA stands for not Available. UK stands for United Kingdom. USA stands for United States of America. ST states for sequence type.

| Isolate name | Year | Country | Host | Disease association | ST | Serotype | Reference | Bioproject |
|---|---|---|---|---|---|---|---|---|
| 0435-10_S10_L001 | 2019 | USA | Pig Diseased | Unknown | 225 | 7 | [16] | PRJNA972671 |
| 0435-11_S11_L001 | 2019 | USA | Pig Diseased | Unknown | 1 | NA | [16] | PRJNA972671 |
| 0435-12_S12_L001 | 2019 | USA | Pig Diseased | Unknown | 1 | 14 | [16] | PRJNA972671 |
| 0435-13_S13_L001 | 2019 | USA | Pig Diseased | Unknown | NA | 9 | [16] | PRJNA972671 |
| 0435-14_S14_L001 | 2019 | USA | Pig Diseased | Unknown | 1 | 14 | [16] | PRJNA972671 |
| 0435-15_S15_L001 | 2019 | USA | Pig Diseased | Unknown | 1 | NA | [16] | PRJNA972671 |
| 0435-16_S16_L001 | 2019 | USA | Pig Diseased | Unknown | NA | 3 | [16] | PRJNA972671 |
| 0435-17_S17_L001 | 2019 | USA | Pig Diseased | Unknown | 1 | 14 | [16] | PRJNA972671 |
| 0435-18_S18_L001 | 2019 | USA | Pig Diseased | Unknown | NA | 23 | [16] | PRJNA972671 |
| 0435-19_S19_L001 | 2019 | USA | Pig Diseased | Unknown | 17 | 4 | [16] | PRJNA972671 |
| 0435-2_S2_L001 | 2018 | USA | Pig Diseased | Unknown | 28 | 2 | [16] | PRJNA972671 |
| 0435-20_S20_L001 | 2019 | USA | Pig Diseased | Unknown | NA | 18 | [16] | PRJNA972671 |
| 0435-21_S21_L001 | 2019 | USA | Pig Diseased | Unknown | NA | 18 | [16] | PRJNA972671 |
| 0435-22_S22_L001 | 2019 | USA | Pig Diseased | Unknown | 1 | 14 | [16] | PRJNA972671 |
| 0435-23_S23_L001 | 2019 | USA | Pig Diseased | Unknown | 94 | 4 | [16] | PRJNA972671 |
| 0435-24_S24_L001 | 2019 | USA | Pig Diseased | Unknown | NA | 2 | [16] | PRJNA972671 |
| 0435-25_S25_L001 | 2019 | USA | Pig Diseased | Unknown | NA | NA | [16] | PRJNA972671 |
| 0435-26_S26_L001 | 2019 | USA | Pig Diseased | Unknown | 1 | 14 | [16] | PRJNA972671 |
| 0435-27_S27_L001 | 2019 | USA | Pig Diseased | Unknown | 1 | NA | [16] | PRJNA972671 |
| 0435-28_S28_L001 | 2020 | USA | Pig Diseased | Unknown | 1 | 14 | [16] | PRJNA972671 |
| 0435-29_S29_L001 | 2019 | USA | Pig Diseased | Unknown | 1 | 14 | [16] | PRJNA972671 |
| 0435-3_S3_L001 | 2019 | USA | Pig Diseased | Unknown | NA | 2 | [16] | PRJNA972671 |
| 0435-30_S30_L001 | 2019 | USA | Pig Diseased | Unknown | 1 | 14 | [16] | PRJNA972671 |

| | | | | | | | | |
|---|---|---|---|---|---|---|---|---|
| 0435-31_S31_L001 | 2018 | USA | Pig Diseased | Unknown | 1 | 14 | [16] | PRJNA972671 |
| 0435-32_S32_L001 | 2019 | USA | Pig Diseased | Unknown | 28 | 2 | [16] | PRJNA972671 |
| 0435-33_S33_L001 | 2019 | USA | Pig Diseased | Unknown | 108 | 23 | [16] | PRJNA972671 |
| 0435-34_S34_L001 | 2019 | USA | Pig Diseased | Unknown | NA | 3 | [16] | PRJNA972671 |
| 0435-4_S4_L001 | 2019 | USA | Pig Diseased | Unknown | NA | 28 | [16] | PRJNA972671 |
| 0435-5_S5_L001 | 2019 | USA | Pig Diseased | Unknown | 87 | 8 | [16] | PRJNA972671 |
| 0435-6_S6_L001 | 2019 | USA | Pig Diseased | Unknown | 373 | 5 | [16] | PRJNA972671 |
| 0435-7_S7_L001 | 2019 | USA | Pig Diseased | Unknown | 87 | 8 | [16] | PRJNA972671 |
| 0435-9_S9_L001 | 2019 | USA | Pig Diseased | Unknown | 225 | 7 | [16] | PRJNA972671 |
| 108-100D_S95_L001 | 2018 | USA | Pig Diseased | Unknown | 1 | 14 | [16] | PRJNA972671 |
| 108-101D_S96_L001 | 2019 | USA | Pig Diseased | Unknown | 1 | 14 | [16] | PRJNA972671 |
| 108-19D_S19_L001 | 2018 | USA | Pig Diseased | Unknown | 1 | 14 | [16] | PRJNA972671 |
| 108-20D_S20_L001 | 2018 | USA | Pig Diseased | Unknown | 1 | 14 | [16] | PRJNA972671 |
| 108-22D_S21_L001 | 2018 | USA | Pig Diseased | Unknown | 94 | 7 | [16] | PRJNA972671 |
| 108-23D_S22_L001 | 2018 | USA | Pig Diseased | Unknown | NA | 2 | [16] | PRJNA972671 |
| 108-24D_S23_L001 | 2018 | USA | Pig Diseased | Unknown | 225 | 7 | [16] | PRJNA972671 |
| 108-26D_S25_L001 | 2018 | USA | Pig Diseased | Unknown | NA | NA | [16] | PRJNA972671 |
| 108-28D_S27_L001 | 2018 | USA | Pig Diseased | Unknown | NA | 28 | [16] | PRJNA972671 |
| 108-29D_S28_L001 | 2018 | USA | Pig Diseased | Unknown | NA | 30 | [16] | PRJNA972671 |
| 108-30D_S29_L001 | 2018 | USA | Pig Diseased | Unknown | 28 | NA | [16] | PRJNA972671 |
| 108-31D_S30_L001 | 2018 | USA | Pig Diseased | Unknown | 1 | NA | [16] | PRJNA972671 |
| 108-32D_S31_L001 | 2018 | USA | Pig Diseased | Unknown | 1 | 14 | [16] | PRJNA972671 |
| 108-34D_S33_L001 | 2018 | USA | Pig Diseased | Unknown | 1 | 14 | [16] | PRJNA972671 |
| 108-35D_S34_L001 | 2018 | USA | Pig Diseased | Unknown | 1 | 14 | [16] | PRJNA972671 |
| 108-36D_S35_L001 | 2018 | USA | Pig Diseased | Unknown | 1 | 14 | [16] | PRJNA972671 |
| 108-37D_S36_L001 | 2018 | USA | Pig Diseased | Unknown | 1 | NA | [16] | PRJNA972671 |

| | | | | | | | | |
|---|---|---|---|---|---|---|---|---|
| 108-38D_S37_L001 | 2018 | USA | Pig Diseased | Unknown | 1 | 14 | [16] | PRJNA972671 |
| 108-39D_S38_L001 | 2018 | USA | Pig Diseased | Unknown | 1 | 14 | [16] | PRJNA972671 |
| 108-40D_S39_L001 | 2018 | USA | Pig Diseased | Unknown | 1 | 14 | [16] | PRJNA972671 |
| 108-41D_S40_L001 | 2018 | USA | Pig Diseased | Unknown | NA | NA | [16] | PRJNA972671 |
| 108-42D_S41_L001 | 2018 | USA | Pig Diseased | Unknown | 1 | 14 | [16] | PRJNA972671 |
| 108-43D_S42_L001 | 2018 | USA | Pig Diseased | Unknown | 1 | 14 | [16] | PRJNA972671 |
| 108-44D_S43_L001 | 2018 | USA | Pig Diseased | Unknown | NA | 8 | [16] | PRJNA972671 |
| 108-45D_S44_L001 | 2018 | USA | Pig Diseased | Unknown | NA | 15 | [16] | PRJNA972671 |
| 108-46D_S45_L001 | 2018 | USA | Pig Diseased | Unknown | NA | NA | [16] | PRJNA972671 |
| 108-47D_S46_L001 | 2018 | USA | Pig Diseased | Unknown | 1 | 14 | [16] | PRJNA972671 |
| 108-48D_S47_L001 | 2018 | USA | Pig Diseased | Unknown | 1 | 14 | [16] | PRJNA972671 |
| 108-49D_S48_L001 | 2018 | USA | Pig Diseased | Unknown | 1 | 14 | [16] | PRJNA972671 |
| 108-51D_S49_L001 | 2018 | USA | Pig Diseased | Unknown | NA | 5 | [16] | PRJNA972671 |
| 108-52D_S50_L001 | 2018 | USA | Pig Diseased | Unknown | 28 | 2 | [16] | PRJNA972671 |
| 108-54D_S51_L001 | 2018 | USA | Pig Diseased | Unknown | 28 | 2 | [16] | PRJNA972671 |
| 108-55D_S52_L001 | 2018 | USA | Pig Diseased | Unknown | NA | NA | [16] | PRJNA972671 |
| 108-56D_S53_L001 | 2018 | USA | Pig Diseased | Unknown | 1 | 14 | [16] | PRJNA972671 |
| 108-57D_S54_L001 | 2018 | USA | Pig Diseased | Unknown | 1 | 14 | [16] | PRJNA972671 |
| 108-58D_S55_L001 | 2018 | USA | Pig Diseased | Unknown | 1 | 14 | [16] | PRJNA972671 |
| 108-59D_S56_L001 | 2018 | USA | Pig Diseased | Unknown | 1 | 14 | [16] | PRJNA972671 |
| 108-60D_S57_L001 | 2018 | USA | Pig Diseased | Unknown | 94 | 5 | [16] | PRJNA972671 |
| 108-61D_S58_L001 | 2018 | USA | Pig Diseased | Unknown | 27 | 3 | [16] | PRJNA972671 |
| 108-62D_S59_L001 | 2018 | USA | Pig Diseased | Unknown | 28 | 2 | [16] | PRJNA972671 |
| 108-63D_S60_L001 | 2018 | USA | Pig Diseased | Unknown | NA | 29 | [16] | PRJNA972671 |
| 108-64D_S61_L001 | 2018 | USA | Pig Diseased | Unknown | NA | 4 | [16] | PRJNA972671 |
| 108-74D_S69_L001 | 2018 | USA | Pig Diseased | Unknown | NA | 31 | [16] | PRJNA972671 |

| | | | | | | | | |
|---|---|---|---|---|---|---|---|---|
| 108-75D_S70_L001 | 2018 | USA | Pig Diseased | Unknown | 1 | 14 | [16] | PRJNA972671 |
| 108-76D_S71_L001 | 2018 | USA | Pig Diseased | Unknown | NA | 9 | [16] | PRJNA972671 |
| 108-77D_S72_L001 | 2018 | USA | Pig Diseased | Unknown | 1 | 14 | [16] | PRJNA972671 |
| 108-78D_S73_L001 | 2018 | USA | Pig Diseased | Unknown | 1 | 14 | [16] | PRJNA972671 |
| 108-79D_S74_L001 | 2018 | USA | Pig Diseased | Unknown | NA | 4 | [16] | PRJNA972671 |
| 108-80D_S75_L001 | 2018 | USA | Pig Diseased | Unknown | 1 | 14 | [16] | PRJNA972671 |
| 108-81D_S76_L001 | 2018 | USA | Pig Diseased | Unknown | 1 | 14 | [16] | PRJNA972671 |
| 108-82D_S77_L001 | 2018 | USA | Pig Diseased | Unknown | NA | 9 | [16] | PRJNA972671 |
| 108-83D_S78_L001 | 2018 | USA | Pig Diseased | Unknown | 28 | 2 | [16] | PRJNA972671 |
| 108-84D_S79_L001 | 2018 | USA | Pig Diseased | Unknown | NA | 10 | [16] | PRJNA972671 |
| 108-85D_S80_L001 | 2018 | USA | Pig Diseased | Unknown | 1 | 14 | [16] | PRJNA972671 |
| 108-89D_S84_L001 | 2018 | USA | Pig Diseased | Unknown | 28 | 2 | [16] | PRJNA972671 |
| 108-90D_S85_L001 | 2018 | USA | Pig Diseased | Unknown | NA | 31 | [16] | PRJNA972671 |
| 108-91D_S86_L001 | 2018 | USA | Pig Diseased | Unknown | NA | 9 | [16] | PRJNA972671 |
| 108-94D_S89_L001 | 2018 | USA | Pig Diseased | Unknown | NA | NA | [16] | PRJNA972671 |
| 108-95D_S90_L001 | 2018 | USA | Pig Diseased | Unknown | NA | 23 | [16] | PRJNA972671 |
| 108-96D_S91_L001 | 2018 | USA | Pig Diseased | Unknown | NA | NA | [16] | PRJNA972671 |
| 108-97D_S92_L001 | 2018 | USA | Pig Diseased | Unknown | NA | NA | [16] | PRJNA972671 |
| 108-98D_S93_L001 | 2018 | USA | Pig Diseased | Unknown | 28 | 2 | [16] | PRJNA972671 |
| 108-99D_S94_L001 | 2018 | USA | Pig Diseased | Unknown | 17 | 4 | [16] | PRJNA972671 |
| 1236-82_S82_L001 | 2019 | USA | Pig Diseased | Unknown | 94 | 3 | [16] | PRJNA972671 |
| 1236-83_S83_L001 | 2017 | USA | Pig Diseased | Unknown | 28 | 2 | [16] | PRJNA972671 |
| 1236-85_S85_L001 | 2019 | USA | Pig Diseased | Unknown | NA | NA | [16] | PRJNA972671 |
| 1237-10_S10_L001 | 2019 | USA | Pig Diseased | Unknown | 28 | 2 | [16] | PRJNA972671 |
| 1237-11_S11_L001 | 2019 | USA | Pig Diseased | Unknown | NA | NA | [16] | PRJNA972671 |
| 1237-14_S14_L001 | 2019 | USA | Pig Diseased | Unknown | NA | 3 | [16] | PRJNA972671 |

| | | | | | | | | |
|---|---|---|---|---|---|---|---|---|
| 1237-15_S15_L001 | 2019 | USA | Pig Diseased | Unknown | 1 | NA | [16] | PRJNA972671 |
| 1237-17_S17_L001 | 2019 | USA | Pig Diseased | Unknown | 1 | 14 | [16] | PRJNA972671 |
| 1237-19_S19_L001 | 2019 | USA | Pig Diseased | Unknown | 1 | NA | [16] | PRJNA972671 |
| 1237-20_S20_L001 | 2019 | USA | Pig Diseased | Unknown | NA | 9 | [16] | PRJNA972671 |
| 1237-21_S21_L001 | 2019 | USA | Pig Diseased | Unknown | NA | 9 | [16] | PRJNA972671 |
| 1237-22_S22_L001 | 2019 | USA | Pig Diseased | Unknown | 1 | NA | [16] | PRJNA972671 |
| 1237-24_S24_L001 | 2019 | USA | Pig Diseased | Unknown | NA | NA | [16] | PRJNA972671 |
| 1237-25_S25_L001 | 2019 | USA | Pig Diseased | Unknown | NA | 9 | [16] | PRJNA972671 |
| 1237-26_S26_L001 | 2019 | USA | Pig Diseased | Unknown | NA | 9 | [16] | PRJNA972671 |
| 1237-27_S27_L001 | 2019 | USA | Pig Diseased | Unknown | NA | NA | [16] | PRJNA972671 |
| 1237-28_S28_L001 | 2019 | USA | Pig Diseased | Unknown | NA | 9 | [16] | PRJNA972671 |
| 1237-29_S29_L001 | 2019 | USA | Pig Diseased | Unknown | NA | 9 | [16] | PRJNA972671 |
| 1237-30_S30_L001 | 2019 | USA | Pig Diseased | Unknown | NA | 23 | [16] | PRJNA972671 |
| 1237-31_S31_L001 | 2019 | USA | Pig Diseased | Unknown | 1 | 14 | [16] | PRJNA972671 |
| 1237-32_S32_L001 | 2019 | USA | Pig Diseased | Unknown | 1 | 14 | [16] | PRJNA972671 |
| 1237-33_S33_L001 | 2019 | USA | Pig Diseased | Unknown | 1 | 14 | [16] | PRJNA972671 |
| 1237-34_S34_L001 | 2019 | USA | Pig Diseased | Unknown | 1 | 14 | [16] | PRJNA972671 |
| 1237-35_S35_L001 | 2019 | USA | Pig Diseased | Unknown | 1 | NA | [16] | PRJNA972671 |
| 1237-36_S36_L001 | 2019 | USA | Pig Diseased | Unknown | 1 | 14 | [16] | PRJNA972671 |
| 1237-37_S37_L001 | 2019 | USA | Pig Diseased | Unknown | 1 | 14 | [16] | PRJNA972671 |
| 1237-38_S38_L001 | 2019 | USA | Pig Diseased | Unknown | 1 | NA | [16] | PRJNA972671 |
| 1237-39_S39_L001 | 2019 | USA | Pig Diseased | Unknown | NA | NA | [16] | PRJNA972671 |
| 1237-40_S40_L001 | 2019 | USA | Pig Diseased | Unknown | NA | NA | [16] | PRJNA972671 |
| 1237-41_S41_L001 | 2019 | USA | Pig Diseased | Unknown | 108 | 23 | [16] | PRJNA972671 |
| 1237-43_S43_L001 | 2019 | USA | Pig Diseased | Unknown | 1 | 14 | [16] | PRJNA972671 |
| 1237-6_S6_L001 | 2019 | USA | Pig Diseased | Unknown | 1 | 2 | [16] | PRJNA972671 |

| | | | | | | | | |
|---|---|---|---|---|---|---|---|---|
| 1237-7_S7_L001 | 2019 | USA | Pig Diseased | Unknown | 1 | 14 | [16] | PRJNA972671 |
| 1237-8_S8_L001 | 2019 | USA | Pig Diseased | Unknown | 1 | NA | [16] | PRJNA972671 |
| 19591_PH2016-001 | 2016 | Germany | Pig Diseased | Systemic | 15 | 3 | [16] | PRJNA1009400 |
| 19592_PH2016-002 | 2016 | Germany | Pig Diseased | Systemic | 1 | 2 | [16] | PRJNA1009400 |
| 19593_PH2016-003 | 2016 | Germany | Pig Diseased | Systemic | 94 | 4 | [16] | PRJNA1009400 |
| 19594_PH2016-004 | 2016 | Germany | Pig Diseased | Systemic | NA | 3 | [16] | PRJNA1009400 |
| 19595_PH2016-006 | 2016 | Germany | Pig Diseased | Systemic | 29 | 7 | [16] | PRJNA1009400 |
| 19596_PH2016-007 | 2016 | Germany | Pig Diseased | Systemic | 198 | 8 | [16] | PRJNA1009400 |
| 19597_PH2016-008 | 2016 | Germany | Pig Diseased | Systemic | 16 | 9 | [16] | PRJNA1009400 |
| 19598_PH2016-009 | 2016 | Germany | Pig Diseased | Systemic | 28 | 2 | [16] | PRJNA1009400 |
| 19599_PH2016-010 | 2016 | Germany | Pig Diseased | Systemic | 16 | 9 | [16] | PRJNA1009400 |
| 19600_PH2016-011 | 2016 | Germany | Pig Diseased | Systemic | NA | 9 | [16] | PRJNA1009400 |
| 19601_PH2016-012 | 2016 | Germany | Pig Diseased | Systemic | 16 | 9 | [16] | PRJNA1009400 |
| 19602_PH2016-014 | 2016 | Germany | Pig Diseased | Systemic | NA | NA | [16] | PRJNA1009400 |
| 19603_PH2016-016 | 2016 | Germany | Pig Diseased | Systemic | 1 | 2 | [16] | PRJNA1009400 |
| 19604_PH2016-017 | 2016 | Germany | Pig Diseased | Systemic | 16 | 9 | [16] | PRJNA1009400 |
| 19605_PH2016-018 | 2016 | Germany | Pig Diseased | Systemic | 16 | 9 | [16] | PRJNA1009400 |
| 19606_PH2016-019 | 2016 | Germany | Pig Diseased | Systemic | 28 | 2 | [16] | PRJNA1009400 |
| 19607_PH2016-020 | 2016 | Germany | Pig Diseased | Systemic | 29 | 7 | [16] | PRJNA1009400 |
| 19608_PH2016-021 | 2016 | Germany | Pig Diseased | Systemic | 28 | 2 | [16] | PRJNA1009400 |
| 19609_PH2016-022 | 2016 | Germany | Pig Diseased | Systemic | 16 | 9 | [16] | PRJNA1009400 |
| 19610_PH2016-024 | 2016 | Germany | Pig Diseased | Systemic | 89 | 7 | [16] | PRJNA1009400 |
| 19611_PH2016-025 | 2016 | Germany | Pig Diseased | Systemic | 308 | 8 | [16] | PRJNA1009400 |
| 19612_PH2016-026 | 2016 | Germany | Pig Diseased | Systemic | 94 | 4 | [16] | PRJNA1009400 |
| 19613_PH2016-027 | 2016 | Germany | Pig Diseased | Systemic | 94 | 4 | [16] | PRJNA1009400 |
| 19614_PH2016-028 | 2016 | Germany | Pig Diseased | Systemic | 98 | 5 | [16] | PRJNA1009400 |

| | | | | | | | | |
|---|---|---|---|---|---|---|---|---|
| 19615_PH2016-029 | 2016 | Germany | Pig Diseased | Systemic | 16 | 9 | [16] | PRJNA1009400 |
| 19616_PH2016-030 | 2016 | Germany | Pig Diseased | Systemic | NA | 9 | [16] | PRJNA1009400 |
| 19618_PH2016-033 | 2016 | Germany | Pig Diseased | Systemic | NA | 16 | [16] | PRJNA1009400 |
| 19619_PH2016-034 | 2016 | Germany | Pig Diseased | Systemic | 29 | 7 | [16] | PRJNA1009400 |
| 19620_PH2016-035 | 2016 | Germany | Pig Diseased | Systemic | 17 | 4 | [16] | PRJNA1009400 |
| 19621_PH2016-037 | 2016 | Germany | Pig Diseased | Systemic | 16 | 9 | [16] | PRJNA1009400 |
| 19622_PH2016-038 | 2016 | Germany | Pig Diseased | Systemic | 124 | 14 | [16] | PRJNA1009400 |
| 19623_PH2016-039 | 2016 | Germany | Pig Diseased | Systemic | 29 | 7 | [16] | PRJNA1009400 |
| 19624_PH2016-040 | 2016 | Germany | Pig Diseased | Systemic | 16 | 9 | [16] | PRJNA1009400 |
| 19625_PH2016-041 | 2016 | Germany | Pig Diseased | Systemic | NA | 18 | [16] | PRJNA1009400 |
| 19626_PH2016-042 | 2016 | Germany | Pig Diseased | Systemic | 28 | 2 | [16] | PRJNA1009400 |
| 19627_PH2016-043 | 2016 | Germany | Pig Diseased | Systemic | 16 | 9 | [16] | PRJNA1009400 |
| 19628_PH2016-044 | 2016 | Germany | Pig Diseased | Systemic | 1 | 14 | [16] | PRJNA1009400 |
| 19629_PH2016-045 | 2016 | Germany | Pig Diseased | Systemic | 16 | 9 | [16] | PRJNA1009400 |
| 19631_PH2016-047 | 2016 | Germany | Pig Diseased | Systemic | 1 | 2 | [16] | PRJNA1009400 |
| 19632_PH2016-048 | 2016 | Germany | Pig Diseased | Systemic | 1 | 2 | [16] | PRJNA1009400 |
| 19633_PH2016-049 | 2016 | Germany | Pig Diseased | Systemic | 29 | 7 | [16] | PRJNA1009400 |
| 19634_PH2016-051 | 2016 | Germany | Pig Diseased | Respiratory | NA | NA | [16] | PRJNA1009400 |
| 19635_PH2016-052 | 2016 | Germany | Pig Diseased | Respiratory | 28 | 2 | [16] | PRJNA1009400 |
| 19636_PH2016-053 | 2016 | Germany | Pig Diseased | Respiratory | 15 | 3 | [16] | PRJNA1009400 |
| 19638_PH2016-055 | 2016 | Germany | Pig Diseased | Respiratory | 28 | 2 | [16] | PRJNA1009400 |
| 19639_PH2016-056 | 2016 | Germany | Pig Diseased | Respiratory | 87 | 8 | [16] | PRJNA1009400 |
| 19640_PH2016-057 | 2016 | Germany | Pig Diseased | Respiratory | 17 | 4 | [16] | PRJNA1009400 |
| 19641_PH2016-059 | 2016 | Germany | Pig Diseased | Respiratory | NA | 29 | [16] | PRJNA1009400 |
| 19644_PH2016-062 | 2016 | Germany | Pig Diseased | Respiratory | 29 | 7 | [16] | PRJNA1009400 |
| 19645_PH2016-063 | 2016 | Germany | Pig Diseased | Respiratory | 17 | 4 | [16] | PRJNA1009400 |

| | | | | | | | | |
|---|---|---|---|---|---|---|---|---|
| 19646_PH2016-064 | 2016 | Germany | Pig Diseased | Respiratory | NA | 21 | 16 | PRJNA1009400 |
| 19647_PH2016-065 | 2016 | Germany | Pig Diseased | Respiratory | NA | NA | 16 | PRJNA1009400 |
| 19648_PH2016-066 | 2016 | Germany | Pig Diseased | Respiratory | 94 | 4 | 16 | PRJNA1009400 |
| 19649_PH2016-067 | 2016 | Germany | Pig Diseased | Respiratory | 17 | 3 | 16 | PRJNA1009400 |
| 19650_PH2016-068 | 2016 | Germany | Pig Diseased | Respiratory | NA | 18 | 16 | PRJNA1009400 |
| 19651_PH2016-069 | 2016 | Germany | Pig Diseased | Respiratory | 28 | 2 | 16 | PRJNA1009400 |
| 19652_PH2016-070 | 2016 | Germany | Pig Diseased | Respiratory | NA | 15 | 16 | PRJNA1009400 |
| 19653_PH2016-071 | 2016 | Germany | Pig Diseased | Respiratory | NA | 16 | 16 | PRJNA1009400 |
| 19654_PH2016-072 | 2016 | Germany | Pig Diseased | Respiratory | NA | 18 | 16 | PRJNA1009400 |
| 19655_PH2016-073 | 2016 | Germany | Pig Diseased | Respiratory | 15 | NA | 16 | PRJNA1009400 |
| 19656_PH2016-074 | 2016 | Germany | Pig Diseased | Respiratory | NA | 31 | 16 | PRJNA1009400 |
| 19657_PH2016-075 | 2016 | Germany | Pig Diseased | Respiratory | NA | 19 | 16 | PRJNA1009400 |
| 19658_PH2016-076 | 2016 | Germany | Pig Diseased | Respiratory | 17 | 4 | 16 | PRJNA1009400 |
| 19659_PH2016-077 | 2016 | Germany | Pig Diseased | Respiratory | 28 | 2 | 16 | PRJNA1009400 |
| 19660_PH2016-078 | 2016 | Germany | Pig Diseased | Respiratory | 28 | 2 | 16 | PRJNA1009400 |
| 19661_PH2016-079 | 2016 | Germany | Pig Diseased | Respiratory | 308 | 8 | 16 | PRJNA1009400 |
| 19662_PH2016-080 | 2016 | Germany | Pig Diseased | Respiratory | 28 | 2 | 16 | PRJNA1009400 |
| 19664_PH2016-082 | 2016 | Germany | Pig Diseased | Respiratory | 87 | 8 | 16 | PRJNA1009400 |
| 19665_PH2016-083 | 2016 | Germany | Pig Diseased | Respiratory | 28 | 2 | 16 | PRJNA1009400 |
| 19666_PH2016-084 | 2016 | Germany | Pig Diseased | Respiratory | 29 | 7 | 16 | PRJNA1009400 |
| 19667_PH2016-086 | 2016 | Germany | Pig Diseased | Respiratory | 87 | 8 | 16 | PRJNA1009400 |
| 19668_PH2016-087 | 2016 | Germany | Pig Diseased | Respiratory | 29 | 7 | 16 | PRJNA1009400 |
| 19669_PH2016-088 | 2016 | Germany | Pig Diseased | Respiratory | 16 | 9 | 16 | PRJNA1009400 |
| 19670_PH2016-089 | 2016 | Germany | Pig Diseased | Respiratory | 17 | 4 | 16 | PRJNA1009400 |
| 19671_PH2016-090 | 2016 | Germany | Pig Diseased | Respiratory | 15 | 3 | 16 | PRJNA1009400 |
| 19672_PH2016-091 | 2016 | Germany | Pig Diseased | Respiratory | NA | 16 | 16 | PRJNA1009400 |

| | | | | | | | | |
|---|---|---|---|---|---|---|---|---|
| 19673_PH2016-092 | 2016 | Germany | Pig Diseased | Respiratory | NA | 18 | [16] | PRJNA1009400 |
| 19674_PH2016-093 | 2016 | Germany | Pig Diseased | Respiratory | 89 | 7 | [16] | PRJNA1009400 |
| 19675_PH2016-094 | 2016 | Germany | Pig Diseased | Respiratory | 308 | 8 | [16] | PRJNA1009400 |
| 19676_PH2016-095 | 2016 | Germany | Pig Diseased | Respiratory | NA | NA | [16] | PRJNA1009400 |
| 19677_PH2016-096 | 2016 | Germany | Pig Diseased | Respiratory | 54 | 4 | [16] | PRJNA1009400 |
| 19678_PH2016-097 | 2016 | Germany | Pig Diseased | Respiratory | NA | 3 | [16] | PRJNA1009400 |
| 19680_PH2016-099 | 2016 | Germany | Pig Diseased | Respiratory | 87 | 8 | [16] | PRJNA1009400 |
| 19681_PH2016-100 | 2016 | Germany | Pig Diseased | Respiratory | 28 | 2 | [16] | PRJNA1009400 |
| 19682_PH2016-101 | 2016 | Germany | Pig Healthy | Non-Clinical | 17 | 4 | [16] | PRJNA1009400 |
| 19683_PH2016-102 | 2016 | Germany | Pig Healthy | Non-Clinical | NA | 28 | [16] | PRJNA1009400 |
| 19684_PH2016-104 | 2016 | Germany | Pig Healthy | Non-Clinical | 17 | 4 | [16] | PRJNA1009400 |
| 19685_PH2016-105 | 2016 | Germany | Pig Healthy | Non-Clinical | NA | 9 | [16] | PRJNA1009400 |
| 19687_PH2016-107 | 2016 | Germany | Pig Healthy | Non-Clinical | 87 | 8 | [16] | PRJNA1009400 |
| 19688_PH2016-108 | 2016 | Germany | Pig Healthy | Non-Clinical | NA | 29 | [16] | PRJNA1009400 |
| 19689_PH2016-109 | 2016 | Germany | Pig Healthy | Non-Clinical | NA | 16 | [16] | PRJNA1009400 |
| 19690_PH2016-110 | 2016 | Germany | Pig Healthy | Non-Clinical | NA | 28 | [16] | PRJNA1009400 |
| 19691_PH2016-111 | 2016 | Germany | Pig Healthy | Non-Clinical | NA | 30 | [16] | PRJNA1009400 |
| 19692_PH2016-112 | 2016 | Germany | Pig Healthy | Non-Clinical | NA | 31 | [16] | PRJNA1009400 |
| 19693_PH2016-113 | 2016 | Germany | Pig Healthy | Non-Clinical | NA | NA | [16] | PRJNA1009400 |
| 19694_PH2016-114 | 2016 | Germany | Pig Healthy | Non-Clinical | 56 | 2 | [16] | PRJNA1009400 |
| 19695_PH2016-115 | 2016 | Germany | Pig Healthy | Non-Clinical | NA | 15 | [16] | PRJNA1009400 |
| 19696_PH2016-116 | 2016 | Germany | Pig Healthy | Non-Clinical | NA | 5 | [16] | PRJNA1009400 |
| 19697_PH2016-117 | 2016 | Germany | Pig Healthy | Non-Clinical | NA | 15 | [16] | PRJNA1009400 |
| 19698_PH2016-118 | 2016 | Germany | Pig Healthy | Non-Clinical | NA | 29 | [16] | PRJNA1009400 |
| 19699_PH2016-119 | 2016 | Germany | Pig Healthy | Non-Clinical | NA | 29 | [16] | PRJNA1009400 |
| 19700_PH2016-120 | 2016 | Germany | Pig Healthy | Non-Clinical | NA | NA | [16] | PRJNA1009400 |

| | | | | | | | | |
|---|---|---|---|---|---|---|---|---|
| 19701_PH2016-121 | 2016 | Germany | Pig Healthy | Non-Clinical | NA | 5 | 16 | PRJNA1009400 |
| 19702_PH2016-123 | 2016 | Germany | Pig Healthy | Non-Clinical | NA | 19 | 16 | PRJNA1009400 |
| 19703_PH2016-124 | 2016 | Germany | Pig Healthy | Non-Clinical | NA | 28 | 16 | PRJNA1009400 |
| 19704_PH2016-125 | 2016 | Germany | Pig Healthy | Non-Clinical | 17 | 4 | 16 | PRJNA1009400 |
| 19705_PH2016-126 | 2016 | Germany | Pig Healthy | Non-Clinical | 87 | 8 | 16 | PRJNA1009400 |
| 19706_PH2016-127 | 2016 | Germany | Pig Healthy | Non-Clinical | NA | 31 | 16 | PRJNA1009400 |
| 19707_PH2016-128 | 2016 | Germany | Pig Healthy | Non-Clinical | 87 | NA | 16 | PRJNA1009400 |
| 19708_PH2016-129 | 2016 | Germany | Pig Healthy | Non-Clinical | NA | NA | 16 | PRJNA1009400 |
| 19709_PH2016-130 | 2016 | Germany | Pig Healthy | Non-Clinical | 17 | 4 | 16 | PRJNA1009400 |
| 19710_PH2016-131 | 2016 | Germany | Pig Healthy | Non-Clinical | NA | 2 | 16 | PRJNA1009400 |
| 19711_PH2016-132 | 2016 | Germany | Pig Healthy | Non-Clinical | 17 | 4 | 16 | PRJNA1009400 |
| 19712_PH2016-133 | 2016 | Germany | Pig Healthy | Non-Clinical | 87 | 8 | 16 | PRJNA1009400 |
| 19713_PH2016-134 | 2016 | Germany | Pig Healthy | Non-Clinical | NA | 29 | 16 | PRJNA1009400 |
| 19714_PH2016-135 | 2016 | Germany | Pig Healthy | Non-Clinical | NA | 16 | 16 | PRJNA1009400 |
| 19716_PH2016-137 | 2016 | Germany | Pig Healthy | Non-Clinical | NA | 29 | 16 | PRJNA1009400 |
| 19718_PH2016-139 | 2016 | Germany | Pig Healthy | Non-Clinical | 28 | 2 | 16 | PRJNA1009400 |
| 19719_PH2016-140 | 2016 | Germany | Pig Healthy | Non-Clinical | NA | 8 | 16 | PRJNA1009400 |
| 19721_PH2016-142 | 2016 | Germany | Pig Healthy | Non-Clinical | NA | 21 | 16 | PRJNA1009400 |
| 19722_PH2016-143 | 2016 | Germany | Pig Healthy | Non-Clinical | NA | NA | 16 | PRJNA1009400 |
| 19723_PH2016-144 | 2016 | Germany | Pig Healthy | Non-Clinical | 87 | 8 | 16 | PRJNA1009400 |
| 19724_PH2016-145 | 2016 | Germany | Pig Healthy | Non-Clinical | NA | 13 | 16 | PRJNA1009400 |
| 19725_PH2016-146 | 2016 | Germany | Pig Healthy | Non-Clinical | NA | 15 | 16 | PRJNA1009400 |
| 19726_PH2016-147 | 2016 | Germany | Pig Healthy | Non-Clinical | NA | 29 | 16 | PRJNA1009400 |
| 19727_PH2016-148 | 2017 | Germany | Pig Healthy | Non-Clinical | NA | NA | 16 | PRJNA1009400 |
| 19728_PH2016-149 | 2017 | Germany | Pig Healthy | Non-Clinical | NA | 29 | 16 | PRJNA1009400 |
| 19729_PH2016-150 | 2017 | Germany | Pig Healthy | Non-Clinical | NA | 29 | 16 | PRJNA1009400 |

| | | | | | | | | |
|---|---|---|---|---|---|---|---|---|
| 19779_M101513_S1 | 2017 | Spain | Pig Diseased | Systemic | 1 | 2 | [16] | PRJNA1009400 |
| 19780_M101565_S2 | 2017 | Spain | Pig Diseased | Systemic | 308 | 8 | [16] | PRJNA1009400 |
| 19781_M101830_S3 | 2017 | Spain | Pig Diseased | Systemic | 123 | 9 | [16] | PRJNA1009400 |
| 19782_M101866_S4 | 2018 | Spain | Pig Diseased | Systemic | 123 | 9 | [16] | PRJNA1009400 |
| 19783_M101866_S5 | 2018 | Spain | Pig Diseased | Systemic | 123 | 9 | [16] | PRJNA1009400 |
| 19784_M101866_S6 | 2017 | Spain | Pig Diseased | Systemic | 123 | 9 | [16] | PRJNA1009400 |
| 19785_M102095_S7 | 2017 | Spain | Pig Diseased | Systemic | 1 | 2 | [16] | PRJNA1009400 |
| 19786_M102327_S8 | 2017 | Spain | Pig Diseased | Systemic | NA | 9 | [16] | PRJNA1009400 |
| 19787_M102942_S9 | 2017 | Spain | Pig Diseased | Systemic | 123 | 9 | [16] | PRJNA1009400 |
| 19788_M102942_S10 | 2017 | Spain | Pig Diseased | Systemic | 123 | 9 | [16] | PRJNA1009400 |
| 19789_M102942_S11 | 2017 | Spain | Pig Diseased | Systemic | 123 | 9 | [16] | PRJNA1009400 |
| 19790_M103642_S12 | 2017 | Spain | Pig Diseased | Systemic | 123 | 9 | [16] | PRJNA1009400 |
| 19791_M103815_S13 | 2017 | Spain | Pig Diseased | Systemic | 123 | 9 | [16] | PRJNA1009400 |
| 19792_M103866_S14 | 2017 | Spain | Pig Diseased | Systemic | 123 | 9 | [16] | PRJNA1009400 |
| 19793_M103947_S15 | 2017 | Spain | Pig Diseased | Systemic | 123 | 9 | [16] | PRJNA1009400 |
| 19794_M103947_S16 | 2017 | Spain | Pig Diseased | Systemic | 123 | 9 | [16] | PRJNA1009400 |
| 19795_M104104_S17 | 2017 | Spain | Pig Diseased | Systemic | 123 | 9 | [16] | PRJNA1009400 |
| 19796_M104104_S18 | 2017 | Spain | Pig Diseased | Systemic | 123 | 9 | [16] | PRJNA1009400 |
| 19797_M104300_S19 | 2017 | Spain | Pig Diseased | Systemic | 1 | 2 | [16] | PRJNA1009400 |
| 19798_M104300_S20 | 2017 | Spain | Pig Diseased | Systemic | 1 | 2 | [16] | PRJNA1009400 |
| 19799_M104421_S21 | 2017 | Spain | Pig Diseased | Systemic | 123 | 9 | [16] | PRJNA1009400 |
| 19800_M104576_S22 | 2017 | Spain | Pig Diseased | Systemic | 123 | 9 | [16] | PRJNA1009400 |
| 19801_M104878_S23 | 2017 | Spain | Pig Diseased | Systemic | 123 | 9 | [16] | PRJNA1009400 |
| 19802_M105040_S24 | 2017 | Spain | Pig Diseased | Systemic | 1 | 2 | [16] | PRJNA1009400 |
| 19803_M105040_S25 | 2017 | Spain | Pig Diseased | Systemic | 1 | 2 | [16] | PRJNA1009400 |
| 19804_M105052_S26 | 2017 | Spain | Pig Diseased | Systemic | NA | 19 | [16] | PRJNA1009400 |

| | | | | | | | | |
|---|---|---|---|---|---|---|---|---|
| 19805_M105237_S27 | 2017 | Spain | Pig Diseased | Systemic | 123 | 9 | [16] | PRJNA1009400 |
| 19806_M105244_S28 | 2017 | Spain | Pig Diseased | Systemic | 1 | 2 | [16] | PRJNA1009400 |
| 19807_M105244_S29 | 2017 | Spain | Pig Diseased | Systemic | 1 | 2 | [16] | PRJNA1009400 |
| 19808_M105552_S30 | 2017 | Spain | Pig Diseased | Systemic | NA | 8 | [16] | PRJNA1009400 |
| 19809_M105623_S31 | 2017 | Spain | Pig Diseased | Systemic | 123 | 9 | [16] | PRJNA1009400 |
| 19810_M105628_S32 | 2017 | Spain | Pig Diseased | Systemic | 1 | 14 | [16] | PRJNA1009400 |
| 19811_M105690_S33 | 2017 | Spain | Pig Diseased | Systemic | 123 | 9 | [16] | PRJNA1009400 |
| 19812_M105966_S34 | 2017 | Spain | Pig Diseased | Systemic | 16 | 9 | [16] | PRJNA1009400 |
| 19813_M106115_S35 | 2017 | Spain | Pig Diseased | Systemic | NA | 29 | [16] | PRJNA1009400 |
| 19814_M106199_S36 | 2017 | Spain | Pig Diseased | Systemic | 123 | 9 | [16] | PRJNA1009400 |
| 19815_M106199_S37 | 2017 | Spain | Pig Diseased | Systemic | 123 | 9 | [16] | PRJNA1009400 |
| 19816_M106252_S38 | 2018 | Spain | Pig Diseased | Systemic | 54 | 3 | [16] | PRJNA1009400 |
| 19817_M106252_S39 | 2018 | Spain | Pig Diseased | Systemic | NA | 31 | [16] | PRJNA1009400 |
| 19818_M106471_S40 | 2018 | Spain | Pig Diseased | Systemic | NA | 30 | [16] | PRJNA1009400 |
| 19819_M106728_S41 | 2018 | Spain | Pig Diseased | Systemic | 15 | 3 | [16] | PRJNA1009400 |
| 19820_M106729_S42 | 2018 | Spain | Pig Diseased | Systemic | 123 | 9 | [16] | PRJNA1009400 |
| 19821_M106729_S43 | 2018 | Spain | Pig Diseased | Systemic | 123 | 9 | [16] | PRJNA1009400 |
| 19822_M106729_S44 | 2018 | Spain | Pig Diseased | Systemic | 123 | 9 | [16] | PRJNA1009400 |
| 19823_M106819_S45 | 2018 | Spain | Pig Diseased | Systemic | 123 | 9 | [16] | PRJNA1009400 |
| 19824_M106898_S46 | 2018 | Spain | Pig Diseased | Systemic | 123 | NA | [16] | PRJNA1009400 |
| 19825_M107022_S47 | 2018 | Spain | Pig Diseased | Systemic | 123 | 9 | [16] | PRJNA1009400 |
| 19826_M107147_S48 | 2018 | Spain | Pig Diseased | Systemic | 123 | 9 | [16] | PRJNA1009400 |
| 19827_M107167_S49 | 2018 | Spain | Pig Diseased | Systemic | 125 | 9 | [16] | PRJNA1009400 |
| 19828_M107306_S50 | 2018 | Spain | Pig Diseased | Systemic | 123 | 9 | [16] | PRJNA1009400 |
| 19829_M101487_R1 | 2017 | Spain | Pig Diseased | Respiratory | NA | 17 | [16] | PRJNA1009400 |
| 19830_M101969_R2 | 2017 | Spain | Pig Diseased | Respiratory | NA | 3 | [16] | PRJNA1009400 |

| | | | | | | | | |
|---|---|---|---|---|---|---|---|---|
| 19832_M102541_R4 | 2018 | Spain | Pig Diseased | Respiratory | NA | NA | [16] | PRJNA1009400 |
| 19833_M103045_R5 | 2018 | Spain | Pig Diseased | Respiratory | NA | 19 | [16] | PRJNA1009400 |
| 19834_M103246_R6 | 2017 | Spain | Pig Diseased | Respiratory | 123 | 9 | [16] | PRJNA1009400 |
| 19835_M103466_R7 | 2017 | Spain | Pig Diseased | Respiratory | 17 | 23 | [16] | PRJNA1009400 |
| 19836_M103466_R8 | 2017 | Spain | Pig Diseased | Respiratory | NA | 24 | [16] | PRJNA1009400 |
| 19837_M103466_R9 | 2017 | Spain | Pig Diseased | Respiratory | NA | 29 | [16] | PRJNA1009400 |
| 19838_M103466_R10 | 2017 | Spain | Pig Diseased | Respiratory | 29 | 7 | [16] | PRJNA1009400 |
| 19839_M103565_R11 | 2017 | Spain | Pig Diseased | Respiratory | NA | 19 | [16] | PRJNA1009400 |
| 19840_M103644_R12 | 2017 | Spain | Pig Diseased | Respiratory | 15 | 3 | [16] | PRJNA1009400 |
| 19841_M103904_R13 | 2017 | Spain | Pig Diseased | Respiratory | NA | 17 | [16] | PRJNA1009400 |
| 19842_M103905_R14 | 2017 | Spain | Pig Diseased | Respiratory | NA | 21 | [16] | PRJNA1009400 |
| 19843_M103975_R15 | 2017 | Spain | Pig Diseased | Respiratory | NA | 29 | [16] | PRJNA1009400 |
| 19845_M103975_R17 | 2017 | Spain | Pig Diseased | Respiratory | NA | 21 | [16] | PRJNA1009400 |
| 19846_M104396_R18 | 2017 | Spain | Pig Diseased | Respiratory | NA | 21 | [16] | PRJNA1009400 |
| 19847_M104396_R19 | 2017 | Spain | Pig Diseased | Respiratory | NA | 21 | [16] | PRJNA1009400 |
| 19848_M107687_R20 | 2018 | Spain | Pig Diseased | Respiratory | 17 | 4 | [16] | PRJNA1009400 |
| 19849_M104767_R21 | 2017 | Spain | Pig Diseased | Respiratory | NA | 17 | [16] | PRJNA1009400 |
| 19850_M104827_R22 | 2017 | Spain | Pig Diseased | Respiratory | NA | 10 | [16] | PRJNA1009400 |
| 19851_M107695_R23 | 2018 | Spain | Pig Diseased | Respiratory | NA | 9 | [16] | PRJNA1009400 |
| 19852_M104890_R24 | 2017 | Spain | Pig Diseased | Respiratory | 15 | 3 | [16] | PRJNA1009400 |
| 19853_M104928_R25 | 2017 | Spain | Pig Diseased | Respiratory | 54 | 4 | [16] | PRJNA1009400 |
| 19854_M105082_R26 | 2017 | Spain | Pig Diseased | Respiratory | NA | 29 | [16] | PRJNA1009400 |
| 19855_M105082_R27 | 2017 | Spain | Pig Diseased | Respiratory | NA | NA | [16] | PRJNA1009400 |
| 19856_M105082_R28 | 2017 | Spain | Pig Diseased | Respiratory | NA | 21 | [16] | PRJNA1009400 |
| 19857_M105235_R29 | 2017 | Spain | Pig Diseased | Respiratory | NA | 16 | [16] | PRJNA1009400 |
| 19858_M105281_R30 | 2017 | Spain | Pig Diseased | Respiratory | 24 | 7 | [16] | PRJNA1009400 |

| | | | | | | | | |
|---|---|---|---|---|---|---|---|---|
| 19859_M105435_R31 | 2017 | Spain | Pig Diseased | Respiratory | 28 | 2 | 16 | PRJNA1009400 |
| 19860_M105455_R32 | 2017 | Spain | Pig Diseased | Respiratory | 28 | 2 | 16 | PRJNA1009400 |
| 19861_M105548_R33 | 2017 | Spain | Pig Diseased | Respiratory | NA | 15 | 16 | PRJNA1009400 |
| 19862_M106003_R34 | 2017 | Spain | Pig Diseased | Respiratory | 15 | 3 | 16 | PRJNA1009400 |
| 19863_M106107_R35 | 2017 | Spain | Pig Diseased | Respiratory | NA | 4 | 16 | PRJNA1009400 |
| 19864_M106316_R36 | 2018 | Spain | Pig Diseased | Respiratory | NA | NA | 16 | PRJNA1009400 |
| 19865_M106387_R37 | 2018 | Spain | Pig Diseased | Respiratory | 15 | 3 | 16 | PRJNA1009400 |
| 19866_M106456_R38 | 2018 | Spain | Pig Diseased | Respiratory | NA | 29 | 16 | PRJNA1009400 |
| 19867_M106485_R39 | 2018 | Spain | Pig Diseased | Respiratory | 1 | 2 | 16 | PRJNA1009400 |
| 19868_M106715_R40 | 2018 | Spain | Pig Diseased | Respiratory | NA | 29 | 16 | PRJNA1009400 |
| 19869_M106931_R41 | 2018 | Spain | Pig Diseased | Respiratory | 123 | 9 | 16 | PRJNA1009400 |
| 19870_M106961_R42 | 2018 | Spain | Pig Diseased | Respiratory | NA | 8 | 16 | PRJNA1009400 |
| 19871_M107169_R43 | 2018 | Spain | Pig Diseased | Respiratory | 123 | 9 | 16 | PRJNA1009400 |
| 19872_M107170_R44 | 2018 | Spain | Pig Diseased | Respiratory | 123 | 9 | 16 | PRJNA1009400 |
| 19873_M107170_R45 | 2018 | Spain | Pig Diseased | Respiratory | 123 | 9 | 16 | PRJNA1009400 |
| 19874_M107244_R46 | 2018 | Spain | Pig Diseased | Respiratory | 94 | 4 | 16 | PRJNA1009400 |
| 19875_M107440_R47 | 2018 | Spain | Pig Diseased | Respiratory | 16 | 9 | 16 | PRJNA1009400 |
| 19876_M107545_R48 | 2018 | Spain | Pig Diseased | Respiratory | NA | 12 | 16 | PRJNA1009400 |
| 19877_M108095_R49 | 2018 | Spain | Pig Diseased | Respiratory | NA | 9 | 16 | PRJNA1009400 |
| 19878_M107642_R50 | 2018 | Spain | Pig Diseased | Respiratory | 17 | 4 | 16 | PRJNA1009400 |
| 19879_M101999_C1 | 2017 | Spain | Pig Healthy | Non-Clinical | NA | 8 | 16 | PRJNA1009400 |
| 19881_AR_1_C3 | 2014 | Spain | Pig Healthy | Non-Clinical | NA | NA | 16 | PRJNA1009400 |
| 19882_CR1_C4 | 2014 | Spain | Pig Healthy | Non-Clinical | NA | 29 | 16 | PRJNA1009400 |
| 19883_CR8_C5 | 2014 | Spain | Pig Healthy | Non-Clinical | 29 | 7 | 16 | PRJNA1009400 |
| 19884_CR9_C6 | 2014 | Spain | Pig Healthy | Non-Clinical | NA | 29 | 16 | PRJNA1009400 |
| 19885_GM2_2_C7 | 2014 | Spain | Pig Healthy | Non-Clinical | NA | 29 | 16 | PRJNA1009400 |

| | | | | | | | | |
|---|---|---|---|---|---|---|---|---|
| 19886_GM8_2_C8 | 2014 | Spain | Pig Healthy | Non-Clinical | NA | 21 | [16] | PRJNA1009400 |
| 19887_VL1_2_C9 | 2014 | Spain | Pig Healthy | Non-Clinical | NA | 21 | [16] | PRJNA1009400 |
| 19889_VL5_4_C11 | 2014 | Spain | Pig Healthy | Non-Clinical | NA | 29 | [16] | PRJNA1009400 |
| 19890_VL8_7_C12 | 2014 | Spain | Pig Healthy | Non-Clinical | NA | 16 | [16] | PRJNA1009400 |
| 19891_WB2_3_C13 | 2015 | Spain | Wild Boar Healthy | Non-Clinical | NA | NA | [16] | PRJNA1009400 |
| 19892_WB4_2_C14 | 2015 | Spain | Wild Boar Healthy | Non-Clinical | NA | 29 | [16] | PRJNA1009400 |
| 19893_SS15055_N2_C15 | 2015 | Spain | Wild Boar Healthy | Non-Clinical | 1 | 2 | [16] | PRJNA1009400 |
| 19894_SS15144_T1_C16 | 2015 | Spain | Wild Boar Healthy | Non-Clinical | 1 | 2 | [16] | PRJNA1009400 |
| 2020Jan-S.suis-1_S1_L001 | 2019 | USA | Pig Diseased | Unknown | NA | 10 | [16] | PRJNA972671 |
| 2020Jan-S.suis-10_S10_L001 | 2019 | USA | Pig Diseased | Unknown | 28 | 2 | [16] | PRJNA972671 |
| 2020Jan-S.suis-11_S11_L001 | 2019 | USA | Pig Diseased | Unknown | NA | 2 | [16] | PRJNA972671 |
| 2020Jan-S.suis-12_S12_L001 | 2019 | USA | Pig Diseased | Unknown | 87 | 8 | [16] | PRJNA972671 |
| 2020Jan-S.suis-13_S13_L001 | 2019 | USA | Pig Diseased | Unknown | 108 | 3 | [16] | PRJNA972671 |
| 2020Jan-S.suis-14_S14_L001 | 2019 | USA | Pig Diseased | Unknown | 54 | 4 | [16] | PRJNA972671 |
| 2020Jan-S.suis-15_S15_L001 | 2019 | USA | Pig Diseased | Unknown | NA | 16 | [16] | PRJNA972671 |
| 2020Jan-S.suis-16_S16_L001 | 2019 | USA | Pig Diseased | Unknown | NA | NA | [16] | PRJNA972671 |
| 2020Jan-S.suis-17_S17_L001 | 2019 | USA | Pig Diseased | Unknown | NA | 15 | [16] | PRJNA972671 |
| 2020Jan-S.suis-18_S18_L001 | 2019 | USA | Pig Diseased | Unknown | 16 | 9 | [16] | PRJNA972671 |
| 2020Jan-S.suis-24_S24_L001 | 2019 | USA | Pig Diseased | Unknown | NA | 3 | [16] | PRJNA972671 |
| 2020Jan-S.suis-25_S25_L001 | 2019 | USA | Pig Diseased | Unknown | 1 | 14 | [16] | PRJNA972671 |
| 2020Jan-S.suis-26_S26_L001 | 2019 | USA | Pig Diseased | Unknown | 28 | 2 | [16] | PRJNA972671 |
| 2020Jan-S.suis-27_S27_L001 | 2019 | USA | Pig Diseased | Unknown | 1 | NA | [16] | PRJNA972671 |
| 2020Jan-S.suis-28_S28_L001 | 2019 | USA | Pig Diseased | Unknown | 28 | 2 | [16] | PRJNA972671 |
| 2020Jan-S.suis-30_S30_L001 | 2019 | USA | Pig Diseased | Unknown | NA | 9 | [16] | PRJNA972671 |
| 2020Jan-S.suis-31_S31_L001 | 2019 | USA | Pig Diseased | Unknown | 28 | 2 | [16] | PRJNA972671 |
| 2020Jan-S.suis-32_S32_L001 | 2019 | USA | Pig Diseased | Unknown | 108 | 4 | [16] | PRJNA972671 |

| | | | | | | | | |
|---|---|---|---|---|---|---|---|---|
| 2020Jan-S.suis-33_S33_L001 | 2019 | USA | Pig Diseased | Unknown | 28 | 2 | [16] | PRJNA972671 |
| 2020Jan-S.suis-34_S34_L001 | 2019 | USA | Pig Diseased | Unknown | 108 | 23 | [16] | PRJNA972671 |
| 2020Jan-S.suis-35_S35_L001 | 2019 | USA | Pig Diseased | Unknown | 28 | 2 | [16] | PRJNA972671 |
| 2020Jan-S.suis-36_S36_L001 | 2019 | USA | Pig Diseased | Unknown | 94 | 7 | [16] | PRJNA972671 |
| 2020Jan-S.suis-37_S37_L001 | 2019 | USA | Pig Diseased | Unknown | 108 | 5 | [16] | PRJNA972671 |
| 2020Jan-S.suis-4_S4_L001 | 2019 | USA | Pig Diseased | Unknown | NA | 10 | [16] | PRJNA972671 |
| 2020Jan-S.suis-5_S5_L001 | 2019 | USA | Pig Diseased | Unknown | NA | NA | [16] | PRJNA972671 |
| 2020Jan-S.suis-68_S67_L001 | 2019 | USA | Pig Diseased | Unknown | 1 | 14 | [16] | PRJNA972671 |
| 2020Jan-S.suis-69_S68_L001 | 2019 | USA | Pig Diseased | Unknown | 1 | NA | [16] | PRJNA972671 |
| 2020Jan-S.suis-7_S7_L001 | 2019 | USA | Pig Diseased | Unknown | 28 | 2 | [16] | PRJNA972671 |
| 2020Jan-S.suis-70_S69_L001 | 2019 | USA | Pig Diseased | Unknown | 1 | 14 | [16] | PRJNA972671 |
| 2020Jan-S.suis-71_S70_L001 | 2019 | USA | Pig Diseased | Unknown | 1 | 14 | [16] | PRJNA972671 |
| 2020Jan-S.suis-74_S73_L001 | 2019 | USA | Pig Diseased | Unknown | 108 | 5 | [16] | PRJNA972671 |
| 2020Jan-S.suis-8_S8_L001 | 2019 | USA | Pig Diseased | Unknown | 1 | NA | [16] | PRJNA972671 |
| 2020Jan-S.suis-9_S9_L001 | 2019 | USA | Pig Diseased | Unknown | 1 | 14 | [16] | PRJNA972671 |
| 21134_DNR1 | 2018 | Denmark | Pig Diseased | Respiratory | 87 | 8 | [16] | PRJNA1009400 |
| 21135_DNR2 | 2018 | Denmark | Pig Diseased | Respiratory | 28 | 2 | [16] | PRJNA1009400 |
| 21136_DNR3 | 2018 | Denmark | Pig Diseased | Respiratory | 28 | 2 | [16] | PRJNA1009400 |
| 21137_DNR4 | 2018 | Denmark | Pig Diseased | Respiratory | 28 | 2 | [16] | PRJNA1009400 |
| 21138_DNR5 | 2018 | Denmark | Pig Diseased | Respiratory | 28 | 2 | [16] | PRJNA1009400 |
| 21139_DNR6 | 2018 | Denmark | Pig Diseased | Respiratory | NA | 8 | [16] | PRJNA1009400 |
| 21140_DNR7 | 2018 | Denmark | Pig Diseased | Respiratory | 17 | 4 | [16] | PRJNA1009400 |
| 21141_DNR8 | 2018 | Denmark | Pig Diseased | Respiratory | 28 | 2 | [16] | PRJNA1009400 |
| 21142_DNR9 | 2018 | Denmark | Pig Diseased | Respiratory | 27 | 3 | [16] | PRJNA1009400 |
| 21143_DNR10 | 2018 | Denmark | Pig Diseased | Respiratory | NA | 3 | [16] | PRJNA1009400 |
| 21144_DNR11 | 2018 | Denmark | Pig Diseased | Respiratory | NA | NA | [16] | PRJNA1009400 |

| | | | | | | | | |
|---|---|---|---|---|---|---|---|---|
| 21145_DNR12 | 2018 | Denmark | Pig Diseased | Respiratory | 29 | 7 | [16] | PRJNA1009400 |
| 21146_DNR13 | 2018 | Denmark | Pig Diseased | Respiratory | 29 | 7 | [16] | PRJNA1009400 |
| 21147_DNR14 | 2018 | Denmark | Pig Diseased | Respiratory | 28 | 2 | [16] | PRJNA1009400 |
| 21148_DNR15 | 2018 | Denmark | Pig Diseased | Respiratory | 29 | 7 | [16] | PRJNA1009400 |
| 21149_DNR16 | 2018 | Denmark | Pig Diseased | Respiratory | NA | 28 | [16] | PRJNA1009400 |
| 21150_DNR17 | 2018 | Denmark | Pig Diseased | Respiratory | 29 | 7 | [16] | PRJNA1009400 |
| 21151_DNR18 | 2018 | Denmark | Pig Diseased | Respiratory | NA | 30 | [16] | PRJNA1009400 |
| 21152_DNR19 | 2018 | Denmark | Pig Diseased | Respiratory | 29 | 7 | [16] | PRJNA1009400 |
| 21153_DNR20 | 2018 | Denmark | Pig Diseased | Respiratory | 28 | 2 | [16] | PRJNA1009400 |
| 21154_DNR21 | 2018 | Denmark | Pig Diseased | Respiratory | 28 | 2 | [16] | PRJNA1009400 |
| 21155_DNR22 | 2018 | Denmark | Pig Diseased | Respiratory | 29 | 7 | [16] | PRJNA1009400 |
| 21156_DNR23 | 2018 | Denmark | Pig Diseased | Respiratory | NA | 5 | [16] | PRJNA1009400 |
| 21157_DNR24 | 2018 | Denmark | Pig Diseased | Respiratory | 94 | 4 | [16] | PRJNA1009400 |
| 21158_DNR25 | 2018 | Denmark | Pig Diseased | Respiratory | NA | 8 | [16] | PRJNA1009400 |
| 21159_DNR26 | 2018 | Denmark | Pig Diseased | Respiratory | 29 | 7 | [16] | PRJNA1009400 |
| 21160_DNR27 | 2018 | Denmark | Pig Diseased | Respiratory | 28 | 2 | [16] | PRJNA1009400 |
| 21161_DNR28 | 2018 | Denmark | Pig Diseased | Respiratory | 28 | 2 | [16] | PRJNA1009400 |
| 21162_DNR29 | 2018 | Denmark | Pig Diseased | Respiratory | NA | 14 | [16] | PRJNA1009400 |
| 21163_DNR30 | 2018 | Denmark | Pig Diseased | Respiratory | NA | 14 | [16] | PRJNA1009400 |
| 21164_DNR31 | 2018 | Denmark | Pig Diseased | Respiratory | NA | 14 | [16] | PRJNA1009400 |
| 21165_DNR32 | 2018 | Denmark | Pig Diseased | Respiratory | 29 | 7 | [16] | PRJNA1009400 |
| 21166_DNR33 | 2018 | Denmark | Pig Diseased | Respiratory | 28 | 2 | [16] | PRJNA1009400 |
| 21167_DNR34 | 2018 | Denmark | Pig Diseased | Respiratory | NA | 8 | [16] | PRJNA1009400 |
| 21168_DNR35 | 2018 | Denmark | Pig Diseased | Respiratory | 87 | 8 | [16] | PRJNA1009400 |
| 21169_DNR36 | 2018 | Denmark | Pig Diseased | Respiratory | 1 | 2 | [16] | PRJNA1009400 |
| 21170_DNR37 | 2018 | Denmark | Pig Diseased | Respiratory | 1 | 2 | [16] | PRJNA1009400 |

| | | | | | | | | |
|---|---|---|---|---|---|---|---|---|
| 21171_DNR38 | 2018 | Denmark | Pig Diseased | Respiratory | 1 | 2 | [16] | PRJNA1009400 |
| 21172_DNR39 | 2018 | Denmark | Pig Diseased | Respiratory | 1 | 2 | [16] | PRJNA1009400 |
| 21173_DNR40 | 2018 | Denmark | Pig Diseased | Respiratory | 28 | 2 | [16] | PRJNA1009400 |
| 21174_DNR41 | 2018 | Denmark | Pig Diseased | Respiratory | NA | 3 | [16] | PRJNA1009400 |
| 21175_DNR42 | 2018 | Denmark | Pig Diseased | Respiratory | 28 | 2 | [16] | PRJNA1009400 |
| 21176_DNR43 | 2018 | Denmark | Pig Diseased | Respiratory | 28 | 2 | [16] | PRJNA1009400 |
| 21177_DNR44 | 2018 | Denmark | Pig Diseased | Respiratory | 54 | 4 | [16] | PRJNA1009400 |
| 21178_DNR45 | 2018 | Denmark | Pig Diseased | Respiratory | 54 | 4 | [16] | PRJNA1009400 |
| 21179_DNR46 | 2018 | Denmark | Pig Diseased | Respiratory | 29 | 7 | [16] | PRJNA1009400 |
| 21180_DNR47 | 2018 | Denmark | Pig Diseased | Respiratory | 87 | 8 | [16] | PRJNA1009400 |
| 21181_DNR48 | 2018 | Denmark | Pig Diseased | Respiratory | NA | 8 | [16] | PRJNA1009400 |
| 21182_DNR49 | 2018 | Denmark | Pig Diseased | Respiratory | 54 | 5 | [16] | PRJNA1009400 |
| 21183_DNR50 | 2018 | Denmark | Pig Diseased | Respiratory | NA | 31 | [16] | PRJNA1009400 |
| 21184_DNS1 | 2018 | Denmark | Pig Diseased | Systemic | 1 | 2 | [16] | PRJNA1009400 |
| 21185_DNS2 | 2018 | Denmark | Pig Diseased | Systemic | 94 | 4 | [16] | PRJNA1009400 |
| 21186_DNS3 | 2018 | Denmark | Pig Diseased | Systemic | 16 | 9 | [16] | PRJNA1009400 |
| 21187_DNS4 | 2018 | Denmark | Pig Diseased | Systemic | 16 | 9 | [16] | PRJNA1009400 |
| 21188_DNS5 | 2018 | Denmark | Pig Diseased | Systemic | 17 | 4 | [16] | PRJNA1009400 |
| 21189_DNS6 | 2018 | Denmark | Pig Diseased | Systemic | 1 | 2 | [16] | PRJNA1009400 |
| 21190_DNS7 | 2018 | Denmark | Pig Diseased | Systemic | 1 | 2 | [16] | PRJNA1009400 |
| 21191_DNS8 | 2018 | Denmark | Pig Diseased | Systemic | 54 | 5 | [16] | PRJNA1009400 |
| 21192_DNS9 | 2018 | Denmark | Pig Diseased | Systemic | 16 | 9 | [16] | PRJNA1009400 |
| 21193_DNS10 | 2018 | Denmark | Pig Diseased | Systemic | 16 | 9 | [16] | PRJNA1009400 |
| 21194_DNS11 | 2018 | Denmark | Pig Diseased | Systemic | 16 | 9 | [16] | PRJNA1009400 |
| 21195_DNS12 | 2018 | Denmark | Pig Diseased | Systemic | 28 | 2 | [16] | PRJNA1009400 |
| 21196_DNS13 | 2018 | Denmark | Pig Diseased | Systemic | 29 | 7 | [16] | PRJNA1009400 |

| | | | | | | | | |
|---|---|---|---|---|---|---|---|---|
| 21197_DNS14 | 2018 | Denmark | Pig Diseased | Systemic | 1 | 2 | [16] | PRJNA1009400 |
| 21198_DNS15 | 2018 | Denmark | Pig Diseased | Systemic | 1 | 2 | [16] | PRJNA1009400 |
| 21199_DNS16 | 2018 | Denmark | Pig Diseased | Systemic | 1 | 2 | [16] | PRJNA1009400 |
| 21200_DNS17 | 2018 | Denmark | Pig Diseased | Systemic | 1 | 2 | [16] | PRJNA1009400 |
| 21201_DNS18 | 2018 | Denmark | Pig Diseased | Systemic | 1 | 2 | [16] | PRJNA1009400 |
| 21202_DNS19 | 2018 | Denmark | Pig Diseased | Systemic | 1 | 2 | [16] | PRJNA1009400 |
| 21204_DNS21 | 2018 | Denmark | Pig Diseased | Systemic | 28 | 2 | [16] | PRJNA1009400 |
| 21205_DNS22 | 2018 | Denmark | Pig Diseased | Systemic | 28 | 2 | [16] | PRJNA1009400 |
| 21206_DNS23 | 2018 | Denmark | Pig Diseased | Systemic | 28 | 2 | [16] | PRJNA1009400 |
| 21207_DNS24 | 2018 | Denmark | Pig Diseased | Systemic | 29 | 7 | [16] | PRJNA1009400 |
| 21208_DNS25 | 2018 | Denmark | Pig Diseased | Systemic | 1 | 2 | [16] | PRJNA1009400 |
| 21209_DNS26 | 2018 | Denmark | Pig Diseased | Systemic | 1 | 2 | [16] | PRJNA1009400 |
| 21210_DNS27 | 2018 | Denmark | Pig Diseased | Systemic | 1 | 2 | [16] | PRJNA1009400 |
| 21211_DNS28 | 2018 | Denmark | Pig Diseased | Systemic | 28 | 2 | [16] | PRJNA1009400 |
| 21212_DNS29 | 2018 | Denmark | Pig Diseased | Systemic | 28 | 2 | [16] | PRJNA1009400 |
| 21213_DNS30 | 2018 | Denmark | Pig Diseased | Systemic | 28 | 2 | [16] | PRJNA1009400 |
| 21215_DNS32 | 2018 | Denmark | Pig Diseased | Systemic | 29 | 7 | [16] | PRJNA1009400 |
| 21216_DNS33 | 2018 | Denmark | Pig Diseased | Systemic | 29 | 7 | [16] | PRJNA1009400 |
| 21217_DNS34 | 2018 | Denmark | Pig Diseased | Systemic | NA | 9 | [16] | PRJNA1009400 |
| 21218_DNS35 | 2018 | Denmark | Pig Diseased | Systemic | 16 | 9 | [16] | PRJNA1009400 |
| 21219_DNS36 | 2018 | Denmark | Pig Diseased | Systemic | 16 | 9 | [16] | PRJNA1009400 |
| 21220_DNS37 | 2018 | Denmark | Pig Diseased | Systemic | NA | 3 | [16] | PRJNA1009400 |
| 21221_DNS38 | 2018 | Denmark | Pig Diseased | Systemic | 29 | 7 | [16] | PRJNA1009400 |
| 21222_DNS39 | 2018 | Denmark | Pig Diseased | Systemic | 29 | 7 | [16] | PRJNA1009400 |
| 21223_DNS40 | 2018 | Denmark | Pig Diseased | Systemic | 29 | 7 | [16] | PRJNA1009400 |
| 21224_DNS41 | 2018 | Denmark | Pig Diseased | Systemic | 28 | 2 | [16] | PRJNA1009400 |

| | | | | | | | | |
|---|---|---|---|---|---|---|---|---|
| 21225_DNS42 | 2018 | Denmark | Pig Diseased | Systemic | 29 | 7 | [16] | PRJNA1009400 |
| 21227_DNS44 | 2018 | Denmark | Pig Diseased | Systemic | NA | 31 | [16] | PRJNA1009400 |
| 21228_DNS45 | 2018 | Denmark | Pig Diseased | Systemic | 29 | 7 | [16] | PRJNA1009400 |
| 21229_DNS46 | 2018 | Denmark | Pig Diseased | Systemic | 29 | 7 | [16] | PRJNA1009400 |
| 21230_DNS47 | 2018 | Denmark | Pig Diseased | Systemic | 29 | 7 | [16] | PRJNA1009400 |
| 21231_DNS48 | 2018 | Denmark | Pig Diseased | Systemic | 29 | 7 | [16] | PRJNA1009400 |
| 21232_DNS49 | 2018 | Denmark | Pig Diseased | Systemic | 29 | 7 | [16] | PRJNA1009400 |
| 21233_DNS50 | 2018 | Denmark | Pig Diseased | Systemic | NA | 28 | [16] | PRJNA1009400 |
| 21234_DNS51 | 2018 | Denmark | Pig Diseased | Systemic | 54 | 4 | [16] | PRJNA1009400 |
| 21235_DNC1 | 2018 | Denmark | Pig Healthy | Non-Clinical | 54 | 5 | [16] | PRJNA1009400 |
| 21236_DNC2 | 2018 | Denmark | Pig Healthy | Non-Clinical | NA | 10 | [16] | PRJNA1009400 |
| 21237_DNC3 | 2018 | Denmark | Pig Healthy | Non-Clinical | NA | 16 | [16] | PRJNA1009400 |
| 21238_DNC4 | 2018 | Denmark | Pig Healthy | Non-Clinical | NA | NA | [16] | PRJNA1009400 |
| 21239_DNC5 | 2018 | Denmark | Pig Healthy | Non-Clinical | NA | 3 | [16] | PRJNA1009400 |
| 21240_DNC6 | 2018 | Denmark | Pig Healthy | Non-Clinical | NA | 3 | [16] | PRJNA1009400 |
| 21241_DNC7 | 2018 | Denmark | Pig Healthy | Non-Clinical | NA | 3 | [16] | PRJNA1009400 |
| 21242_DNC8 | 2018 | Denmark | Pig Healthy | Non-Clinical | NA | 15 | [16] | PRJNA1009400 |
| 21243_DNC9 | 2018 | Denmark | Pig Healthy | Non-Clinical | NA | 15 | [16] | PRJNA1009400 |
| 21244_DNC10 | 2018 | Denmark | Pig Healthy | Non-Clinical | NA | 31 | [16] | PRJNA1009400 |
| 21245_DNC11 | 2018 | Denmark | Pig Healthy | Non-Clinical | NA | 16 | [16] | PRJNA1009400 |
| 21246_DNC12 | 2018 | Denmark | Pig Healthy | Non-Clinical | NA | 31 | [16] | PRJNA1009400 |
| 21248_DNC14 | 2018 | Denmark | Pig Healthy | Non-Clinical | NA | NA | [16] | PRJNA1009400 |
| 21249_DNC15 | 2018 | Denmark | Pig Healthy | Non-Clinical | NA | 16 | [16] | PRJNA1009400 |
| 21250_DNC16 | 2018 | Denmark | Pig Healthy | Non-Clinical | NA | 11 | [16] | PRJNA1009400 |
| 21251_DNC17 | 2018 | Denmark | Pig Healthy | Non-Clinical | NA | 31 | [16] | PRJNA1009400 |
| 21252_DNC18 | 2018 | Denmark | Pig Healthy | Non-Clinical | 54 | 3 | [16] | PRJNA1009400 |

| | | | | | | | | |
|---|---|---|---|---|---|---|---|---|
| 21254_DNC20 | 2018 | Denmark | Pig Healthy | Non-Clinical | NA | 30 | [16] | PRJNA1009400 |
| 21255_DNC21 | 2018 | Denmark | Pig Healthy | Non-Clinical | NA | 30 | [16] | PRJNA1009400 |
| 21256_DNC22 | 2018 | Denmark | Pig Healthy | Non-Clinical | NA | 16 | [16] | PRJNA1009400 |
| 21257_DNC23 | 2018 | Denmark | Pig Healthy | Non-Clinical | NA | 31 | [16] | PRJNA1009400 |
| 21258_DNC24 | 2018 | Denmark | Pig Healthy | Non-Clinical | NA | 8 | [16] | PRJNA1009400 |
| 21259_DNC25 | 2018 | Denmark | Pig Healthy | Non-Clinical | NA | 8 | [16] | PRJNA1009400 |
| 21260_DNC26 | 2018 | Denmark | Pig Healthy | Non-Clinical | NA | 16 | [16] | PRJNA1009400 |
| 21261_DNC27 | 2018 | Denmark | Pig Healthy | Non-Clinical | NA | NA | [16] | PRJNA1009400 |
| 21262_DNC28 | 2018 | Denmark | Pig Healthy | Non-Clinical | 54 | 5 | [16] | PRJNA1009400 |
| 21263_DNC29 | 2018 | Denmark | Pig Healthy | Non-Clinical | 54 | 5 | [16] | PRJNA1009400 |
| 21264_DNC30 | 2018 | Denmark | Pig Healthy | Non-Clinical | NA | NA | [16] | PRJNA1009400 |
| 21265_DNC31 | 2018 | Denmark | Pig Healthy | Non-Clinical | NA | NA | [16] | PRJNA1009400 |
| 21266_DNC32 | 2018 | Denmark | Pig Healthy | Non-Clinical | 54 | 3 | [16] | PRJNA1009400 |
| 21267_DNC33 | 2018 | Denmark | Pig Healthy | Non-Clinical | 29 | 7 | [16] | PRJNA1009400 |
| 21268_DNC34 | 2018 | Denmark | Pig Healthy | Non-Clinical | NA | 16 | [16] | PRJNA1009400 |
| 21270_DNC36 | 2018 | Denmark | Pig Healthy | Non-Clinical | NA | 28 | [16] | PRJNA1009400 |
| 21271_DNC37 | 2018 | Denmark | Pig Healthy | Non-Clinical | NA | 29 | [16] | PRJNA1009400 |
| 21272_DNC38 | 2018 | Denmark | Pig Healthy | Non-Clinical | NA | 16 | [16] | PRJNA1009400 |
| 21273_DNC39 | 2018 | Denmark | Pig Healthy | Non-Clinical | NA | NA | [16] | PRJNA1009400 |
| 21274_DNC40 | 2018 | Denmark | Pig Healthy | Non-Clinical | NA | 31 | [16] | PRJNA1009400 |
| 21275_DNC41 | 2018 | Denmark | Pig Healthy | Non-Clinical | NA | 16 | [16] | PRJNA1009400 |
| 21276_DNC42 | 2018 | Denmark | Pig Healthy | Non-Clinical | NA | 16 | [16] | PRJNA1009400 |
| 21277_DNC43 | 2018 | Denmark | Pig Healthy | Non-Clinical | NA | NA | [16] | PRJNA1009400 |
| 21278_DNC44 | 2018 | Denmark | Pig Healthy | Non-Clinical | NA | 29 | [16] | PRJNA1009400 |
| 21279_DNC45 | 2018 | Denmark | Pig Healthy | Non-Clinical | 17 | 4 | [16] | PRJNA1009400 |
| 21280_DNC46 | 2018 | Denmark | Pig Healthy | Non-Clinical | NA | 13 | [16] | PRJNA1009400 |

| | | | | | | | | |
|---|---|---|---|---|---|---|---|---|
| 21281_DNC47 | 2018 | Denmark | Pig Healthy | Non-Clinical | NA | 16 | [16] | PRJNA1009400 |
| 21282_DNC48 | 2018 | Denmark | Pig Healthy | Non-Clinical | NA | 31 | [16] | PRJNA1009400 |
| 21283_DNC49 | 2018 | Denmark | Pig Healthy | Non-Clinical | 28 | 2 | [16] | PRJNA1009400 |
| 21284_DNC50 | 2018 | Denmark | Pig Healthy | Non-Clinical | 87 | 8 | [16] | PRJNA1009400 |
| 21285_NLR1 | 2017 | Netherlands | Pig Diseased | Respiratory | 87 | 8 | [16] | PRJNA1009400 |
| 21286_NLR2 | 2017 | Netherlands | Pig Diseased | Respiratory | 16 | 9 | [16] | PRJNA1009400 |
| 21287_NLR3 | 2017 | Netherlands | Pig Diseased | Respiratory | 28 | 2 | [16] | PRJNA1009400 |
| 21288_NLR4 | 2017 | Netherlands | Pig Diseased | Respiratory | 28 | 2 | [16] | PRJNA1009400 |
| 21289_NLR5 | 2017 | Netherlands | Pig Diseased | Respiratory | 17 | 4 | [16] | PRJNA1009400 |
| 21290_NLR6 | 2017 | Netherlands | Pig Diseased | Respiratory | 29 | 7 | [16] | PRJNA1009400 |
| 21291_NLR7 | 2017 | Netherlands | Pig Diseased | Respiratory | 28 | 2 | [16] | PRJNA1009400 |
| 21292_NLR8 | 2017 | Netherlands | Pig Diseased | Respiratory | NA | NA | [16] | PRJNA1009400 |
| 21293_NLR9 | 2017 | Netherlands | Pig Diseased | Respiratory | 16 | 9 | [16] | PRJNA1009400 |
| 21294_NLR10 | 2017 | Netherlands | Pig Diseased | Respiratory | NA | 12 | [16] | PRJNA1009400 |
| 21296_NLR12 | 2017 | Netherlands | Pig Diseased | Respiratory | 15 | 3 | [16] | PRJNA1009400 |
| 21297_NLR13 | 2017 | Netherlands | Pig Diseased | Respiratory | 28 | 2 | [16] | PRJNA1009400 |
| 21298_NLR14 | 2017 | Netherlands | Pig Diseased | Respiratory | 87 | 8 | [16] | PRJNA1009400 |
| 21299_NLR15 | 2017 | Netherlands | Pig Diseased | Respiratory | 16 | 9 | [16] | PRJNA1009400 |
| 21300_NLR16 | 2017 | Netherlands | Pig Diseased | Respiratory | 29 | 7 | [16] | PRJNA1009400 |
| 21301_NLR17 | 2017 | Netherlands | Pig Diseased | Respiratory | 16 | 9 | [16] | PRJNA1009400 |
| 21302_NLR18 | 2017 | Netherlands | Pig Diseased | Respiratory | 1 | 2 | [16] | PRJNA1009400 |
| 21303_NLR19 | 2017 | Netherlands | Pig Diseased | Respiratory | 136 | 9 | [16] | PRJNA1009400 |
| 21304_NLR20 | 2017 | Netherlands | Pig Diseased | Respiratory | 28 | 2 | [16] | PRJNA1009400 |
| 21305_NLR21 | 2017 | Netherlands | Pig Diseased | Respiratory | 28 | 2 | [16] | PRJNA1009400 |
| 21306_NLR22 | 2017 | Netherlands | Pig Diseased | Respiratory | 15 | 3 | [16] | PRJNA1009400 |
| 21307_NLR23 | 2017 | Netherlands | Pig Diseased | Respiratory | 15 | 3 | [16] | PRJNA1009400 |

| | | | | | | | | |
|---|---|---|---|---|---|---|---|---|
| 21308_NLR24 | 2017 | Netherlands | Pig Diseased | Respiratory | 16 | 9 | [16] | PRJNA1009400 |
| 21309_NLR25 | 2017 | Netherlands | Pig Diseased | Respiratory | 29 | 7 | [16] | PRJNA1009400 |
| 21310_NLR26 | 2017 | Netherlands | Pig Diseased | Respiratory | 14 | 8 | [16] | PRJNA1009400 |
| 21311_NLR27 | 2017 | Netherlands | Pig Diseased | Respiratory | 29 | 7 | [16] | PRJNA1009400 |
| 21312_NLR28 | 2017 | Netherlands | Pig Diseased | Respiratory | 28 | 2 | [16] | PRJNA1009400 |
| 21313_NLR29 | 2017 | Netherlands | Pig Diseased | Respiratory | 16 | 9 | [16] | PRJNA1009400 |
| 21314_NLR30 | 2017 | Netherlands | Pig Diseased | Respiratory | NA | 2 | [16] | PRJNA1009400 |
| 21315_NLR31 | 2017 | Netherlands | Pig Diseased | Respiratory | 16 | 9 | [16] | PRJNA1009400 |
| 21316_NLR32 | 2017 | Netherlands | Pig Diseased | Respiratory | NA | 19 | [16] | PRJNA1009400 |
| 21317_NLR33 | 2017 | Netherlands | Pig Diseased | Respiratory | NA | NA | [16] | PRJNA1009400 |
| 21318_NLR34 | 2017 | Netherlands | Pig Diseased | Respiratory | NA | NA | [16] | PRJNA1009400 |
| 21319_NLR35 | 2017 | Netherlands | Pig Diseased | Respiratory | 17 | 4 | [16] | PRJNA1009400 |
| 21320_NLR36 | 2017 | Netherlands | Pig Diseased | Respiratory | 28 | 2 | [16] | PRJNA1009400 |
| 21321_NLR37 | 2017 | Netherlands | Pig Diseased | Respiratory | 28 | 2 | [16] | PRJNA1009400 |
| 21322_NLR38 | 2017 | Netherlands | Pig Diseased | Respiratory | 14 | 8 | [16] | PRJNA1009400 |
| 21323_NLR39 | 2017 | Netherlands | Pig Diseased | Respiratory | 15 | 3 | [16] | PRJNA1009400 |
| 21324_NLR40 | 2017 | Netherlands | Pig Diseased | Respiratory | NA | 2 | [16] | PRJNA1009400 |
| 21325_NLR41 | 2017 | Netherlands | Pig Diseased | Respiratory | 1 | 2 | [16] | PRJNA1009400 |
| 21326_NLR42 | 2017 | Netherlands | Pig Diseased | Respiratory | NA | 4 | [16] | PRJNA1009400 |
| 21327_NLR43 | 2017 | Netherlands | Pig Diseased | Respiratory | NA | 10 | [16] | PRJNA1009400 |
| 21328_NLR44 | 2017 | Netherlands | Pig Diseased | Respiratory | 54 | 5 | [16] | PRJNA1009400 |
| 21329_NLR45 | 2017 | Netherlands | Pig Diseased | Respiratory | 15 | 3 | [16] | PRJNA1009400 |
| 21330_NLR46 | 2017 | Netherlands | Pig Diseased | Respiratory | NA | 18 | [16] | PRJNA1009400 |
| 21331_NLR47 | 2017 | Netherlands | Pig Diseased | Respiratory | 28 | 2 | [16] | PRJNA1009400 |
| 21332_NLR48 | 2017 | Netherlands | Pig Diseased | Respiratory | 16 | 9 | [16] | PRJNA1009400 |
| 21333_NLR49 | 2017 | Netherlands | Pig Diseased | Respiratory | NA | 19 | [16] | PRJNA1009400 |

| | | | | | | | | |
|---|---|---|---|---|---|---|---|---|
| 21334_NLR50 | 2017 | Netherlands | Pig Diseased | Respiratory | 16 | 9 | [16] | PRJNA1009400 |
| 21335_NLS1 | 2017 | Netherlands | Pig Diseased | Systemic | 16 | 9 | [16] | PRJNA1009400 |
| 21336_NLS2 | 2017 | Netherlands | Pig Diseased | Systemic | 16 | 9 | [16] | PRJNA1009400 |
| 21337_NLS3 | 2017 | Netherlands | Pig Diseased | Systemic | 1 | 2 | [16] | PRJNA1009400 |
| 21338_NLS4 | 2017 | Netherlands | Pig Diseased | Systemic | 108 | 23 | [16] | PRJNA1009400 |
| 21340_NLS6 | 2017 | Netherlands | Pig Diseased | Systemic | 20 | 2 | [16] | PRJNA1009400 |
| 21341_NLS7 | 2017 | Netherlands | Pig Diseased | Systemic | 1 | 2 | [16] | PRJNA1009400 |
| 21342_NLS8 | 2017 | Netherlands | Pig Diseased | Systemic | 1 | 2 | [16] | PRJNA1009400 |
| 21343_NLS9 | 2017 | Netherlands | Pig Diseased | Systemic | 16 | 9 | [16] | PRJNA1009400 |
| 21344_NLS10 | 2017 | Netherlands | Pig Diseased | Systemic | 16 | 9 | [16] | PRJNA1009400 |
| 21345_NLS11 | 2017 | Netherlands | Pig Diseased | Systemic | 16 | 9 | [16] | PRJNA1009400 |
| 21347_NLS13 | 2017 | Netherlands | Pig Diseased | Systemic | NA | 31 | [16] | PRJNA1009400 |
| 21348_NLS14 | 2017 | Netherlands | Pig Diseased | Systemic | 14 | 9 | [16] | PRJNA1009400 |
| 21349_NLS15 | 2017 | Netherlands | Pig Diseased | Systemic | 16 | 9 | [16] | PRJNA1009400 |
| 21350_NLS16 | 2017 | Netherlands | Pig Diseased | Systemic | 29 | 7 | [16] | PRJNA1009400 |
| 21351_NLS17 | 2017 | Netherlands | Pig Diseased | Systemic | 108 | 23 | [16] | PRJNA1009400 |
| 21352_NLS18 | 2017 | Netherlands | Pig Diseased | Systemic | 16 | 9 | [16] | PRJNA1009400 |
| 21353_NLS19 | 2017 | Netherlands | Pig Diseased | Systemic | 16 | 9 | [16] | PRJNA1009400 |
| 21356_NLS22 | 2017 | Netherlands | Pig Diseased | Systemic | 16 | 9 | [16] | PRJNA1009400 |
| 21357_NLS23 | 2017 | Netherlands | Pig Diseased | Systemic | 28 | 2 | [16] | PRJNA1009400 |
| 21358_NLS24 | 2017 | Netherlands | Pig Diseased | Systemic | 29 | 7 | [16] | PRJNA1009400 |
| 21359_NLS25 | 2017 | Netherlands | Pig Diseased | Systemic | 16 | 9 | [16] | PRJNA1009400 |
| 21360_NLS26 | 2017 | Netherlands | Pig Diseased | Systemic | NA | 12 | [16] | PRJNA1009400 |
| 21361_NLS27 | 2017 | Netherlands | Pig Diseased | Systemic | 1 | NA | [16] | PRJNA1009400 |
| 21364_NLS30 | 2017 | Netherlands | Pig Diseased | Systemic | 15 | 3 | [16] | PRJNA1009400 |
| 21365_NLS31 | 2017 | Netherlands | Pig Diseased | Systemic | 29 | 7 | [16] | PRJNA1009400 |

| | | | | | | | | |
|---|---|---|---|---|---|---|---|---|
| 21366_NLS32 | 2017 | Netherlands | Pig Diseased | Systemic | 16 | 9 | [16] | PRJNA1009400 |
| 21367_NLS33 | 2017 | Netherlands | Pig Diseased | Systemic | 29 | 7 | [16] | PRJNA1009400 |
| 21368_NLS34 | 2017 | Netherlands | Pig Diseased | Systemic | 1 | 2 | [16] | PRJNA1009400 |
| 21370_NLS36 | 2017 | Netherlands | Pig Diseased | Systemic | 16 | 9 | [16] | PRJNA1009400 |
| 21371_NLS37 | 2017 | Netherlands | Pig Diseased | Systemic | 1 | 2 | [16] | PRJNA1009400 |
| 21372_NLS38 | 2017 | Netherlands | Pig Diseased | Systemic | 16 | 9 | [16] | PRJNA1009400 |
| 21373_NLS39 | 2017 | Netherlands | Pig Diseased | Systemic | 16 | 9 | [16] | PRJNA1009400 |
| 21375_NLS41 | 2017 | Netherlands | Pig Diseased | Systemic | 16 | 9 | [16] | PRJNA1009400 |
| 21377_NLS43 | 2017 | Netherlands | Pig Diseased | Systemic | 16 | 9 | [16] | PRJNA1009400 |
| 21378_NLS44 | 2017 | Netherlands | Pig Diseased | Systemic | NA | NA | [16] | PRJNA1009400 |
| 21379_NLS45 | 2017 | Netherlands | Pig Diseased | Systemic | 16 | 9 | [16] | PRJNA1009400 |
| 21380_NLS46 | 2017 | Netherlands | Pig Diseased | Systemic | 1 | 14 | [16] | PRJNA1009400 |
| 21382_NLS48 | 2017 | Netherlands | Pig Diseased | Systemic | 1 | 14 | [16] | PRJNA1009400 |
| 21383_NLS49 | 2017 | Netherlands | Pig Diseased | Systemic | 16 | 9 | [16] | PRJNA1009400 |
| 21386_NLC2 | 2017 | Netherlands | Pig Healthy | Non-Clinical | NA | 12 | [16] | PRJNA1009400 |
| 21387_NLC3 | 2017 | Netherlands | Pig Healthy | Non-Clinical | NA | 21 | [16] | PRJNA1009400 |
| 21388_NLC4 | 2017 | Netherlands | Pig Healthy | Non-Clinical | NA | 31 | [16] | PRJNA1009400 |
| 21390_NLC6 | 2017 | Netherlands | Pig Healthy | Non-Clinical | NA | 29 | [16] | PRJNA1009400 |
| 21391_NLC7 | 2017 | Netherlands | Pig Healthy | Non-Clinical | 29 | 7 | [16] | PRJNA1009400 |
| 21393_NLC9 | 2017 | Netherlands | Pig Healthy | Non-Clinical | 29 | 7 | [16] | PRJNA1009400 |
| 21394_NLC10 | 2017 | Netherlands | Pig Healthy | Non-Clinical | NA | NA | [16] | PRJNA1009400 |
| 21395_NLC11 | 2017 | Netherlands | Pig Healthy | Non-Clinical | NA | 6 | [16] | PRJNA1009400 |
| 21396_NLC12 | 2017 | Netherlands | Pig Healthy | Non-Clinical | NA | NA | [16] | PRJNA1009400 |
| 21397_NLC13 | 2017 | Netherlands | Pig Healthy | Non-Clinical | NA | 28 | [16] | PRJNA1009400 |
| 21400_NLC16 | 2017 | Netherlands | Pig Healthy | Non-Clinical | NA | 31 | [16] | PRJNA1009400 |
| 21401_NLC17 | 2017 | Netherlands | Pig Healthy | Non-Clinical | NA | 5 | [16] | PRJNA1009400 |

| | | | | | | | | |
|---|---|---|---|---|---|---|---|---|
| 21402_NLC18 | 2017 | Netherlands | Pig Healthy | Non-Clinical | NA | 2 | [16] | PRJNA1009400 |
| 21403_NLC19 | 2017 | Netherlands | Pig Healthy | Non-Clinical | NA | NA | [16] | PRJNA1009400 |
| 21404_NLC20 | 2017 | Netherlands | Pig Healthy | Non-Clinical | NA | 16 | [16] | PRJNA1009400 |
| 21408_NLC24 | 2017 | Netherlands | Pig Healthy | Non-Clinical | NA | NA | [16] | PRJNA1009400 |
| 21410_NLC44 | 2017 | Netherlands | Pig Healthy | Non-Clinical | NA | 12 | [16] | PRJNA1009400 |
| 21413_NLC29 | 2017 | Netherlands | Pig Healthy | Non-Clinical | 28 | 2 | [16] | PRJNA1009400 |
| 21414_NLC30 | 2017 | Netherlands | Pig Healthy | Non-Clinical | NA | 21 | [16] | PRJNA1009400 |
| 21416_NLC32 | 2017 | Netherlands | Pig Healthy | Non-Clinical | 14 | 8 | [16] | PRJNA1009400 |
| 21417_NLC33 | 2017 | Netherlands | Pig Healthy | Non-Clinical | 17 | 4 | [16] | PRJNA1009400 |
| 21418_NLC34 | 2017 | Netherlands | Pig Healthy | Non-Clinical | NA | 29 | [16] | PRJNA1009400 |
| 21422_NLC38 | 2017 | Netherlands | Pig Healthy | Non-Clinical | NA | 30 | [16] | PRJNA1009400 |
| 21423_NLC39 | 2017 | Netherlands | Pig Healthy | Non-Clinical | 16 | 9 | [16] | PRJNA1009400 |
| 21424_NLC40 | 2017 | Netherlands | Pig Healthy | Non-Clinical | NA | 31 | [16] | PRJNA1009400 |
| 21425_NLC41 | 2017 | Netherlands | Pig Healthy | Non-Clinical | NA | 9 | [16] | PRJNA1009400 |
| 21431_NLC47 | 2017 | Netherlands | Pig Healthy | Non-Clinical | NA | 21 | [16] | PRJNA1009400 |
| 21483_PH005 | 2016 | Germany | Pig Diseased | Systemic | NA | 5 | [16] | PRJNA1009400 |
| 21484_PH013 | 2016 | Germany | Pig Diseased | Systemic | NA | 28 | [16] | PRJNA1009400 |
| 21485_PH015 | 2016 | Germany | Pig Diseased | Systemic | 16 | 9 | [16] | PRJNA1009400 |
| 21486_PH023 | 2016 | Germany | Pig Diseased | Systemic | NA | 9 | [16] | PRJNA1009400 |
| 21487_PH032 | 2016 | Germany | Pig Diseased | Systemic | 16 | 9 | [16] | PRJNA1009400 |
| 21488_PH036 | 2016 | Germany | Pig Diseased | Systemic | NA | 11 | [16] | PRJNA1009400 |
| 21489_PH050 | 2016 | Germany | Pig Diseased | Systemic | 1 | 14 | [16] | PRJNA1009400 |
| 21490_PH058 | 2016 | Germany | Pig Diseased | Respiratory | 308 | 8 | [16] | PRJNA1009400 |
| 21491_PH085 | 2016 | Germany | Pig Diseased | Respiratory | NA | 31 | [16] | PRJNA1009400 |
| 21492_PH103 | 2018 | Germany | Pig Healthy | Non-Clinical | NA | NA | [16] | PRJNA1009400 |
| 21493_PH122 | 2018 | Germany | Pig Healthy | Non-Clinical | NA | NA | [16] | PRJNA1009400 |

| | | | | | | | | |
|---|---|---|---|---|---|---|---|---|
| 21494_SPR16 | 2017 | Spain | Pig Diseased | Respiratory | NA | NA | [16] | PRJNA1009400 |
| 22303_SPC3 | 2014 | Spain | Pig Healthy | Non-Clinical | NA | NA | [16] | PRJNA1009400 |
| 22304_SPC4 | 2014 | Spain | Pig Healthy | Non-Clinical | NA | NA | [16] | PRJNA1009400 |
| 22305_SPC6 | 2014 | Spain | Pig Healthy | Non-Clinical | NA | NA | [16] | PRJNA1009400 |
| 22308_SPC11 | 2014 | Spain | Pig Healthy | Non-Clinical | NA | NA | [16] | PRJNA1009400 |
| 22309_SPC14 | 2015 | Spain | Pig Healthy | Non-Clinical | NA | NA | [16] | PRJNA1009400 |
| 22311_SPC16 | 2015 | Spain | Pig Healthy | Non-Clinical | NA | NA | [16] | PRJNA1009400 |
| 22312_SPC17 | 2014 | Spain | Pig Healthy | Non-Clinical | NA | NA | [16] | PRJNA1009400 |
| 22314_SPC20 | 2014 | Spain | Pig Healthy | Non-Clinical | NA | NA | [16] | PRJNA1009400 |
| 22315_2606-1D15 | 2017 | Spain | Pig Healthy | Non-Clinical | NA | NA | [16] | PRJNA1009400 |
| 22316_45-1D7 | 2017 | Spain | Pig Healthy | Non-Clinical | NA | NA | [16] | PRJNA1009400 |
| 22317_73-4D7 | 2017 | Spain | Pig Healthy | Non-Clinical | NA | NA | [16] | PRJNA1009400 |
| 22318_73-1D7 | 2017 | Spain | Pig Healthy | Non-Clinical | NA | NA | [16] | PRJNA1009400 |
| 22319_7-1D7 | 2017 | Spain | Pig Healthy | Non-Clinical | NA | NA | [16] | PRJNA1009400 |
| 22320_326-1D7 | 2017 | Spain | Pig Healthy | Non-Clinical | NA | NA | [16] | PRJNA1009400 |
| 22321_816-2D21 | 2017 | Spain | Pig Healthy | Non-Clinical | NA | NA | [16] | PRJNA1009400 |
| 22322_255-1D7 | 2017 | Spain | Pig Healthy | Non-Clinical | NA | NA | [16] | PRJNA1009400 |
| 22323_895-LCR | 2017 | Spain | Pig Diseased | Systemic | NA | NA | [16] | PRJNA1009400 |
| 22324_613-1D49 | 2017 | Spain | Pig Healthy | Non-Clinical | NA | NA | [16] | PRJNA1009400 |
| 22325_559-1D21 | 2017 | Spain | Pig Healthy | Non-Clinical | NA | NA | [16] | PRJNA1009400 |
| 22327_328-3D49 | 2017 | Spain | Pig Healthy | Non-Clinical | NA | NA | [16] | PRJNA1009400 |
| 22329_783-3D21 | 2017 | Spain | Pig Healthy | Non-Clinical | NA | NA | [16] | PRJNA1009400 |
| 22332_751-2D21 | 2017 | Spain | Pig Healthy | Non-Clinical | NA | NA | [16] | PRJNA1009400 |
| 22333_Z2-1 | 2017 | Spain | Pig Healthy | Non-Clinical | NA | NA | [16] | PRJNA1009400 |
| 22334_805-4D21 | 2017 | Spain | Pig Healthy | Non-Clinical | NA | NA | [16] | PRJNA1009400 |
| 22335_73-1D15 | 2017 | Spain | Pig Healthy | Non-Clinical | NA | NA | [16] | PRJNA1009400 |

| | | | | | | | | |
|---|---|---|---|---|---|---|---|---|
| 22336_326-2D7 | 2017 | Spain | Pig Healthy | Non-Clinical | NA | NA | [16] | PRJNA1009400 |
| 25301_PH1993199827 | 1997 | Germany | Pig Unknown | Unknown | NA | 5 | [16] | PRJNA1009400 |
| 25302_PH1993199865 | 1998 | Germany | Pig Unknown | Unknown | 28 | 2 | [16] | PRJNA1009400 |
| 25303_PH1993199811 | 1996 | Germany | Pig Unknown | Unknown | 28 | 2 | [16] | PRJNA1009400 |
| 25304_PH1993199836 | 1997 | Germany | Pig Unknown | Unknown | 28 | 2 | [16] | PRJNA1009400 |
| 25305_PH1993199812 | 1996 | Germany | Pig Unknown | Unknown | 94 | 4 | [16] | PRJNA1009400 |
| 25306_PH1993199867 | 1998 | Germany | Pig Unknown | Unknown | 16 | 9 | [16] | PRJNA1009400 |
| 25307_PH1993199814 | 1996 | Germany | Pig Unknown | Unknown | NA | 5 | [16] | PRJNA1009400 |
| 25308_PH199319988 | 1994 | Germany | Pig Unknown | Unknown | 16 | 9 | [16] | PRJNA1009400 |
| 25309_PH1993199820 | 1996 | Germany | Pig Unknown | Unknown | 16 | 9 | [16] | PRJNA1009400 |
| 25311_PH1993199845 | 1996 | Germany | Pig Unknown | Unknown | NA | 5 | [16] | PRJNA1009400 |
| 25312_PH1993199822 | 1996 | Germany | Pig Unknown | Unknown | 27 | 3 | [16] | PRJNA1009400 |
| 25313_PH1993199830 | 1997 | Germany | Pig Unknown | Unknown | 27 | 3 | [16] | PRJNA1009400 |
| 25314_PH1993199832 | 1997 | Germany | Pig Unknown | Unknown | 15 | 3 | [16] | PRJNA1009400 |
| 25316_PH1993199837 | 1997 | Germany | Pig Unknown | Unknown | NA | NA | [16] | PRJNA1009400 |
| 25317_PH1993199852 | 1997 | Germany | Pig Unknown | Unknown | 29 | 7 | [16] | PRJNA1009400 |
| 25319_PH1993199854 | 1997 | Germany | Pig Unknown | Unknown | 17 | 4 | [16] | PRJNA1009400 |
| 25320_PH1993199818 | 1996 | Germany | Pig Unknown | Unknown | 16 | 9 | [16] | PRJNA1009400 |
| 25321_PH1993199828 | 1997 | Germany | Pig Unknown | Unknown | NA | 5 | [16] | PRJNA1009400 |
| 25322_PH1993199855 | 1997 | Germany | Pig Unknown | Unknown | 94 | 4 | [16] | PRJNA1009400 |
| 25323_PH1993199863 | 1998 | Germany | Pig Unknown | Unknown | 29 | 7 | [16] | PRJNA1009400 |
| 28283_HSPI171 | 1997 | Spain | Pig Unknown | Unknown | NA | 11 | [16] | PRJNA1009400 |
| 28284_HSPI233 | 1997 | Spain | Pig Unknown | Unknown | 28 | 2 | [16] | PRJNA1009400 |
| 28285_HSPI245 | 1997 | Spain | Pig Unknown | Unknown | NA | 16 | [16] | PRJNA1009400 |
| 28286_HSPI11 | 1995 | Spain | Pig Unknown | Unknown | NA | 28 | [16] | PRJNA1009400 |
| 28287_HSPI35 | 1996 | Spain | Pig Unknown | Unknown | 123 | 9 | [16] | PRJNA1009400 |

| | | | | | | | | |
|---|---|---|---|---|---|---|---|---|
| 28288_HSPI47 | 1996 | Spain | Pig Unknown | Unknown | NA | 7 | [16] | PRJNA1009400 |
| 28289_HSPI143 | 1996 | Spain | Pig Unknown | Unknown | NA | 31 | [16] | PRJNA1009400 |
| 28290_HSPI145 | 1996 | Spain | Pig Unknown | Unknown | NA | NA | [16] | PRJNA1009400 |
| 28291_HSPI177 | 1997 | Spain | Pig Unknown | Unknown | 231 | 3 | [16] | PRJNA1009400 |
| 28292_HSPI275 | 1999 | Spain | Pig Unknown | Unknown | NA | NA | [16] | PRJNA1009400 |
| 28293_HSPI46 | 1996 | Spain | Pig Unknown | Unknown | NA | 21 | [16] | PRJNA1009400 |
| 28294_HSPI83 | 1996 | Spain | Pig Unknown | Unknown | NA | 29 | [16] | PRJNA1009400 |
| 28295_HSPI88 | 1996 | Spain | Pig Unknown | Unknown | 14 | 8 | [16] | PRJNA1009400 |
| 28296_HSPI89 | 1996 | Spain | Pig Unknown | Unknown | NA | NA | [16] | PRJNA1009400 |
| 28297_HSPI141 | 1996 | Spain | Pig Unknown | Unknown | NA | 19 | [16] | PRJNA1009400 |
| 28298_HSPI158 | 1997 | Spain | Pig Unknown | Unknown | 1 | 14 | [16] | PRJNA1009400 |
| 28299_HSPI222 | 1997 | Spain | Pig Unknown | Unknown | NA | 5 | [16] | PRJNA1009400 |
| 28300_HSPI252 | 1998 | Spain | Pig Unknown | Unknown | 27 | 3 | [16] | PRJNA1009400 |
| 28301_HSPGP439 | 2005 | Spain | Pig Unknown | Unknown | 28 | 2 | [16] | PRJNA1009400 |
| 28302_HNL1463 | 1983 | Netherlands | Pig Diseased | Systemic | 15 | 3 | [16] | PRJNA1009400 |
| 28303_HNL1468 | 1983 | Netherlands | Pig Diseased | Respiratory | 108 | 4 | [16] | PRJNA1009400 |
| 28304_HNL2467 | 1983 | Netherlands | Pig Diseased | Systemic | NA | 3 | [16] | PRJNA1009400 |
| 28305_HNL2490 | 1983 | Netherlands | Pig Diseased | Systemic | 16 | 9 | [16] | PRJNA1009400 |
| 28306_HNL2492 | 1983 | Netherlands | Pig Diseased | Systemic | 56 | 2 | [16] | PRJNA1009400 |
| 28307_HNL2499 | 1983 | Netherlands | Pig Diseased | Systemic | 29 | 7 | [16] | PRJNA1009400 |
| 28308_HNL2669 | 1983 | Netherlands | Pig Diseased | Systemic | 13 | 14 | [16] | PRJNA1009400 |
| 28309_HNL2728 | 1983 | Netherlands | Pig Diseased | Respiratory | 16 | 9 | [16] | PRJNA1009400 |
| 28310_HNL2737 | 1983 | Netherlands | Pig Diseased | Respiratory | 87 | 8 | [16] | PRJNA1009400 |
| 28311_HNL2824 | 1983 | Netherlands | Pig Diseased | Systemic | 16 | 9 | [16] | PRJNA1009400 |
| 28312_HNL2850 | 1983 | Netherlands | Pig Diseased | Respiratory | 108 | 4 | [16] | PRJNA1009400 |
| 28313_HNL2852 | 1983 | Netherlands | Pig Diseased | Systemic | 20 | 2 | [16] | PRJNA1009400 |

| | | | | | | | | |
|---|---|---|---|---|---|---|---|---|
| 28314_HNL2897 | 1983 | Netherlands | Pig Diseased | Respiratory | 87 | 8 | [16] | PRJNA1009400 |
| 28315_HNL3561 | 1984 | Netherlands | Pig Diseased | Systemic | 29 | 7 | [16] | PRJNA1009400 |
| 28316_HNL5993 | 1992 | Netherlands | Pig Diseased | Systemic | 13 | 14 | [16] | PRJNA1009400 |
| 28317_HNL6011 | 1992 | Netherlands | Pig Diseased | Systemic | 16 | 9 | [16] | PRJNA1009400 |
| 28318_HNL6051 | 1992 | Netherlands | Pig Diseased | Systemic | 15 | 3 | [16] | PRJNA1009400 |
| 28319_HNL6070 | 1992 | Netherlands | Pig Diseased | Respiratory | NA | 5 | [16] | PRJNA1009400 |
| 28320_HNL6097 | 1992 | Netherlands | Pig Diseased | Systemic | NA | 9 | [16] | PRJNA1009400 |
| 28321_HNL6547 | 1993 | Netherlands | Pig Diseased | Respiratory | NA | 8 | [16] | PRJNA1009400 |
| 28322_HNL6620 | 1993 | Netherlands | Pig Diseased | Respiratory | 29 | 7 | [16] | PRJNA1009400 |
| 28323_HNL6706 | 1993 | Netherlands | Pig Diseased | Systemic | 13 | 14 | [16] | PRJNA1009400 |
| 28324_HNL6738 | 1993 | Netherlands | Pig Diseased | Systemic | 16 | 9 | [16] | PRJNA1009400 |
| 28325_HNL6763 | 1993 | Netherlands | Pig Diseased | Systemic | 13 | 14 | [16] | PRJNA1009400 |
| 28326_HNL6766 | 1993 | Netherlands | Pig Diseased | Systemic | NA | 7 | [16] | PRJNA1009400 |
| 28327_HNL6792 | 1993 | Netherlands | Pig Diseased | Systemic | 108 | 4 | [16] | PRJNA1009400 |
| 28328_HNL16560 | 1960 | Netherlands | Pig Unknown | Unknown | 13 | 14 | [16] | PRJNA1009400 |
| 28329_HNL26780 | 1981 | Netherlands | Pig Unknown | Unknown | 29 | 7 | [16] | PRJNA1009400 |
| 28330_201988791 | 2004 | Denmark | Pig Diseased | Unknown | 1 | 2 | [16] | PRJNA1009400 |
| 28331_201988792 | 2004 | Denmark | Pig Diseased | Unknown | 27 | 3 | [16] | PRJNA1009400 |
| 28332_201988793 | 2004 | Denmark | Pig Diseased | Unknown | 17 | 4 | [16] | PRJNA1009400 |
| 28333_201988794 | 2004 | Denmark | Pig Diseased | Unknown | 87 | 8 | [16] | PRJNA1009400 |
| 28334_201988795 | 2004 | Denmark | Pig Diseased | Unknown | 54 | 5 | [16] | PRJNA1009400 |
| 28335_201988796 | 2004 | Denmark | Pig Diseased | Unknown | 35 | 3 | [16] | PRJNA1009400 |
| 28336_201988797 | 2004 | Denmark | Pig Diseased | Unknown | 54 | 4 | [16] | PRJNA1009400 |
| 28337_201988798 | 2004 | Denmark | Pig Diseased | Unknown | 29 | 7 | [16] | PRJNA1009400 |
| 28338_201988799 | 2004 | Denmark | Pig Diseased | Unknown | 94 | 4 | [16] | PRJNA1009400 |
| 28339_2019887910 | 2004 | Denmark | Pig Diseased | Unknown | 54 | 5 | [16] | PRJNA1009400 |

| | | | | | | | | |
|---|---|---|---|---|---|---|---|---|
| 28340_2019887911 | 2004 | Denmark | Pig Diseased | Unknown | 27 | 3 | [16] | PRJNA1009400 |
| 28341_2019887912 | 2004 | Denmark | Pig Diseased | Unknown | 54 | 4 | [16] | PRJNA1009400 |
| 28342_2019887913 | 2004 | Denmark | Pig Diseased | Unknown | 17 | 4 | [16] | PRJNA1009400 |
| 28343_2019887914 | 2004 | Denmark | Pig Diseased | Unknown | NA | 3 | [16] | PRJNA1009400 |
| 28344_2019887915 | 2004 | Denmark | Pig Diseased | Unknown | 54 | 4 | [16] | PRJNA1009400 |
| 28345_2019887916 | 2004 | Denmark | Pig Diseased | Unknown | 87 | 8 | [16] | PRJNA1009400 |
| 28346_2019887917 | 2004 | Denmark | Pig Diseased | Unknown | 35 | 3 | [16] | PRJNA1009400 |
| 28347_2019887918 | 2004 | Denmark | Pig Diseased | Unknown | 643 | 2 | [16] | PRJNA1009400 |
| 28348_2019887919 | 2004 | Denmark | Pig Diseased | Unknown | 94 | 4 | [16] | PRJNA1009400 |
| 28349_2019887920 | 2005 | Denmark | Pig Diseased | Unknown | 1 | 2 | [16] | PRJNA1009400 |
| 28350_2019887921 | 2007 | Denmark | Pig Diseased | Unknown | 1 | 14 | [16] | PRJNA1009400 |
| 28351_2019887922 | 2007 | Denmark | Pig Diseased | Unknown | 1 | NA | [16] | PRJNA1009400 |
| 28352_2019887923 | 2001 | Denmark | Pig Diseased | Unknown | 13 | 14 | [16] | PRJNA1009400 |
| 28353_2019887924 | 2001 | Denmark | Pig Diseased | Unknown | 1 | 14 | [16] | PRJNA1009400 |
| 28375_SS15064T1 | 2015 | Spain | Wild Boar Healthy | Non-Clinical | NA | 4 | [16] | PRJNA1009400 |
| 28376_SS15079N3 | 2015 | Spain | Wild Boar Healthy | Non-Clinical | NA | 4 | [16] | PRJNA1009400 |
| 28377_SS15080T3 | 2015 | Spain | Wild Boar Healthy | Non-Clinical | NA | 4 | [16] | PRJNA1009400 |
| 28378_SS15120T1 | 2015 | Spain | Wild Boar Healthy | Non-Clinical | NA | 4 | [16] | PRJNA1009400 |
| 28379_SS15125T3 | 2015 | Spain | Wild Boar Healthy | Non-Clinical | NA | 4 | [16] | PRJNA1009400 |
| 28380_SS15149T1 | 2015 | Spain | Wild Boar Healthy | Non-Clinical | NA | 4 | [16] | PRJNA1009400 |
| 28382_SS15095N3 | 2015 | Spain | Wild Boar Healthy | Non-Clinical | NA | 8 | [16] | PRJNA1009400 |
| 28383_SS15127N2 | 2015 | Spain | Wild Boar Healthy | Non-Clinical | NA | 8 | [16] | PRJNA1009400 |
| 28384_SS15147T1 | 2015 | Spain | Wild Boar Healthy | Non-Clinical | NA | 8 | [16] | PRJNA1009400 |
| 28387_SS15105T2 | 2015 | Spain | Wild Boar Healthy | Non-Clinical | NA | 9 | [16] | PRJNA1009400 |
| 28388_SS15109T2 | 2015 | Spain | Wild Boar Healthy | Non-Clinical | NA | 9 | [16] | PRJNA1009400 |
| 28389_SS15112T2 | 2015 | Spain | Wild Boar Healthy | Non-Clinical | NA | NA | [16] | PRJNA1009400 |

| | | | | | | | | |
|---|---|---|---|---|---|---|---|---|
| 28390_SS15150N1 | 2015 | Spain | Wild Boar Healthy | Non-Clinical | NA | 9 | [16] | PRJNA1009400 |
| 28391_SS15151T4 | 2015 | Spain | Wild Boar Healthy | Non-Clinical | NA | 9 | [16] | PRJNA1009400 |
| 28392_SS15048T2 | 2015 | Spain | Wild Boar Healthy | Non-Clinical | NA | 10 | [16] | PRJNA1009400 |
| 28393_SS150302T | 2015 | Spain | Wild Boar Healthy | Non-Clinical | NA | 15 | [16] | PRJNA1009400 |
| 28394_SS15045N4 | 2015 | Spain | Wild Boar Healthy | Non-Clinical | NA | 16 | [16] | PRJNA1009400 |
| 28395_SS15153T4 | 2015 | Spain | Wild Boar Healthy | Non-Clinical | NA | 16 | [16] | PRJNA1009400 |
| 28397_SS15072N2 | 2015 | Spain | Wild Boar Healthy | Non-Clinical | NA | 21 | [16] | PRJNA1009400 |
| 28398_SS15130N2 | 2015 | Spain | Wild Boar Healthy | Non-Clinical | NA | NA | [16] | PRJNA1009400 |
| 28401_SS15052T1 | 2015 | Spain | Wild Boar Healthy | Non-Clinical | NA | 31 | [16] | PRJNA1009400 |
| 28402_SS15061T1 | 2015 | Spain | Wild Boar Healthy | Non-Clinical | NA | 31 | [16] | PRJNA1009400 |
| 28403_SS15124N3 | 2015 | Spain | Wild Boar Healthy | Non-Clinical | 209 | 10 | [16] | PRJNA1009400 |
| 28404_SS15126N1 | 2015 | Spain | Wild Boar Healthy | Non-Clinical | NA | 10 | [16] | PRJNA1009400 |
| 28405_SS150295N | 2015 | Spain | Wild Boar Healthy | Non-Clinical | NA | NA | [16] | PRJNA1009400 |
| 28407_SS15059N1 | 2015 | Spain | Wild Boar Healthy | Non-Clinical | NA | NA | [16] | PRJNA1009400 |
| 28410_SS15128N4 | 2015 | Spain | Wild Boar Healthy | Non-Clinical | NA | NA | [16] | PRJNA1009400 |
| 2957-1_S1_L001 | 2018 | USA | Pig Diseased | Unknown | 108 | 23 | [16] | PRJNA972671 |
| 2957-10_S10_L001 | 2018 | USA | Pig Diseased | Unknown | 108 | 23 | [16] | PRJNA972671 |
| 2957-11_S11_L001 | 2018 | USA | Pig Diseased | Unknown | 108 | 23 | [16] | PRJNA972671 |
| 2957-12_S12_L001 | 2018 | USA | Pig Diseased | Unknown | 108 | 23 | [16] | PRJNA972671 |
| 2957-13_S13_L001 | 2018 | USA | Pig Diseased | Unknown | 108 | 23 | [16] | PRJNA972671 |
| 2957-15_S15_L001 | 2018 | USA | Pig Diseased | Unknown | 108 | 5 | [16] | PRJNA972671 |
| 2957-16_S16_L001 | 2018 | USA | Pig Diseased | Unknown | NA | NA | [16] | PRJNA972671 |
| 2957-17_S17_L001 | 2018 | USA | Pig Diseased | Unknown | 28 | NA | [16] | PRJNA972671 |
| 2957-2_S2_L001 | 2018 | USA | Pig Diseased | Unknown | 87 | 8 | [16] | PRJNA972671 |
| 2957-20_S20_L001 | 2018 | USA | Pig Diseased | Unknown | 28 | NA | [16] | PRJNA972671 |
| 2957-21_S21_L001 | 2018 | USA | Pig Diseased | Unknown | NA | NA | [16] | PRJNA972671 |

| | | | | | | | | |
|---|---|---|---|---|---|---|---|---|
| 2957-22_S22_L001 | 2019 | USA | Pig Diseased | Unknown | 485 | 4 | [16] | PRJNA972671 |
| 2957-23_S23_L001 | 2019 | USA | Pig Diseased | Unknown | 485 | 4 | [16] | PRJNA972671 |
| 2957-24_S24_L001 | 2019 | USA | Pig Diseased | Unknown | NA | 9 | [16] | PRJNA972671 |
| 2957-25_S25_L001 | 2018 | USA | Pig Diseased | Unknown | 108 | 23 | [16] | PRJNA972671 |
| 2957-26_S26_L001 | 2018 | USA | Pig Diseased | Unknown | NA | 5 | [16] | PRJNA972671 |
| 2957-27_S27_L001 | 2018 | USA | Pig Diseased | Unknown | NA | 31 | [16] | PRJNA972671 |
| 2957-28_S28_L001 | 2019 | USA | Pig Diseased | Unknown | 1 | NA | [16] | PRJNA972671 |
| 2957-29_S29_L001 | 2018 | USA | Pig Diseased | Unknown | NA | 2 | [16] | PRJNA972671 |
| 2957-3_S3_L001 | 2018 | USA | Pig Diseased | Unknown | 108 | NA | [16] | PRJNA972671 |
| 2957-30_S30_L001 | 2019 | USA | Pig Diseased | Unknown | 23 | 4 | [16] | PRJNA972671 |
| 2957-31_S31_L001 | 2019 | USA | Pig Diseased | Unknown | NA | 9 | [16] | PRJNA972671 |
| 2957-32_S32_L001 | 2019 | USA | Pig Diseased | Unknown | NA | 9 | [16] | PRJNA972671 |
| 2957-33_S33_L001 | 2019 | USA | Pig Diseased | Unknown | 17 | NA | [16] | PRJNA972671 |
| 2957-34_S34_L001 | 2019 | USA | Pig Diseased | Unknown | 28 | NA | [16] | PRJNA972671 |
| 2957-35_S35_L001 | 2018 | USA | Pig Diseased | Unknown | 28 | 2 | [16] | PRJNA972671 |
| 2957-36_S36_L001 | 2019 | USA | Pig Diseased | Unknown | 108 | 23 | [16] | PRJNA972671 |
| 2957-39_S39_L001 | 2018 | USA | Pig Diseased | Unknown | 1 | NA | [16] | PRJNA972671 |
| 2957-4_S4_L001 | 2018 | USA | Pig Diseased | Unknown | 225 | 7 | [16] | PRJNA972671 |
| 2957-40_S40_L001 | 2019 | USA | Pig Diseased | Unknown | 108 | 23 | [16] | PRJNA972671 |
| 2957-41_S41_L001 | 2018 | USA | Pig Diseased | Unknown | 28 | 2 | [16] | PRJNA972671 |
| 2957-43_S43_L001 | 2019 | USA | Pig Diseased | Unknown | 1 | 14 | [16] | PRJNA972671 |
| 2957-50_S50_L001 | 2019 | USA | Pig Diseased | Unknown | 108 | 23 | [16] | PRJNA972671 |
| 2957-6_S6_L001 | 2018 | USA | Pig Diseased | Unknown | 1 | 14 | [16] | PRJNA972671 |
| 2957-7_S7_L001 | 2018 | USA | Pig Diseased | Unknown | 28 | 2 | [16] | PRJNA972671 |
| 2957-8_S8_L001 | 2019 | USA | Pig Diseased | Unknown | 28 | 2 | [16] | PRJNA972671 |
| 2957-9_S9_L001 | 2018 | USA | Pig Diseased | Unknown | 29 | 7 | [16] | PRJNA972671 |

| | | | | | | | | |
|---|---|---|---|---|---|---|---|---|
| 2972-1_S26_L001 | 2019 | USA | Pig Diseased | Unknown | 87 | 8 | [16] | PRJNA972671 |
| 2972-2_S27_L001 | 2019 | USA | Pig Diseased | Unknown | 1 | 14 | [16] | PRJNA972671 |
| 2972-3_S28_L001 | 2019 | USA | Pig Diseased | Unknown | 1 | NA | [16] | PRJNA972671 |
| 2972-4_S29_L001 | 2019 | USA | Pig Diseased | Unknown | NA | 9 | [16] | PRJNA972671 |
| 2972-5_S30_L001 | 2019 | USA | Pig Diseased | Unknown | 1 | NA | [16] | PRJNA972671 |
| 33135_Y0506 | 1988 | UK | Pig Diseased | Systemic | 1 | 2 | [16] | PRJNA1009400 |
| 33136_Y0498 | 1990 | UK | Pig Diseased | Systemic | NA | 2 | [16] | PRJNA1009400 |
| 33137_Y0497 | 1990 | UK | Pig Diseased | Systemic | 1 | 2 | [16] | PRJNA1009400 |
| 33140_Y0491 | 1989 | UK | Pig Diseased | Systemic | 1 | 2 | [16] | PRJNA1009400 |
| 33141_Y0473 | 1987 | UK | Pig Diseased | Systemic | 11 | 14 | [16] | PRJNA1009400 |
| 33142_Y0472 | 1987 | UK | Pig Diseased | Systemic | NA | NA | [16] | PRJNA1009400 |
| 33143_Y0471 | 1987 | UK | Pig Diseased | Systemic | 30 | 3 | [16] | PRJNA1009400 |
| 33145_Y0509 | 1989 | UK | Pig Diseased | Systemic | 1 | 2 | [16] | PRJNA1009400 |
| 33146_Y0510 | 1989 | UK | Pig Diseased | Systemic | 1 | 2 | [16] | PRJNA1009400 |
| 33147_Y0514 | 1989 | UK | Pig Diseased | Systemic | 25 | 2 | [16] | PRJNA1009400 |
| 33148_Y0515 | 1989 | UK | Pig Diseased | Systemic | 1 | 2 | [16] | PRJNA1009400 |
| 33149_Y0517 | 1989 | UK | Pig Diseased | Systemic | 1 | 2 | [16] | PRJNA1009400 |
| 33150_Y0518 | 1990 | UK | Pig Diseased | Systemic | 1 | 2 | [16] | PRJNA1009400 |
| 33151_Y0642 | 1990 | UK | Pig Diseased | Systemic | 25 | 2 | [16] | PRJNA1009400 |
| 33152_Y1579 | 1998 | UK | Pig Diseased | Systemic | NA | 2 | [16] | PRJNA1009400 |
| 33153_Y1589 | 1998 | UK | Pig Diseased | Respiratory | 23 | 4 | [16] | PRJNA1009400 |
| 33154_Y1590 | 1998 | UK | Pig Diseased | Respiratory | 1 | 2 | [16] | PRJNA1009400 |
| 33155_Y1799 | 1998 | UK | Pig Diseased | Systemic | 1 | 2 | [16] | PRJNA1009400 |
| 33156_Y1870 | 1998 | UK | Pig Diseased | Respiratory | 27 | 3 | [16] | PRJNA1009400 |
| 33157_Y1965 | 1998 | UK | Pig Diseased | Systemic | 1 | 14 | [16] | PRJNA1009400 |
| 33158_Y2002 | 1995 | UK | Pig Diseased | Systemic | 29 | 7 | [16] | PRJNA1009400 |

| | | | | | | | | |
|---|---|---|---|---|---|---|---|---|
| 33159_Y2006 | 1995 | UK | Pig Diseased | Respiratory | 29 | 7 | [16] | PRJNA1009400 |
| 33160_Y2009 | 1995 | UK | Pig Diseased | Systemic | 44 | 5 | [16] | PRJNA1009400 |
| 33161_Y2019 | 1995 | UK | Pig Diseased | Respiratory | NA | 3 | [16] | PRJNA1009400 |
| 33162_Y2105 | 1996 | UK | Pig Diseased | Respiratory | 31 | 3 | [16] | PRJNA1009400 |
| 33163_Y2123 | 1996 | UK | Pig Diseased | Respiratory | 33 | 3 | [16] | PRJNA1009400 |
| 33164_Y2125 | 1996 | UK | Pig Diseased | Respiratory | 34 | 7 | [16] | PRJNA1009400 |
| 33165_Y2126 | 1996 | UK | Pig Diseased | Respiratory | 29 | 7 | [16] | PRJNA1009400 |
| 33166_Y2136 | 1996 | UK | Pig Diseased | Systemic | 23 | 4 | [16] | PRJNA1009400 |
| 33167_Y2148 | 1996 | UK | Pig Diseased | Systemic | 29 | 7 | [16] | PRJNA1009400 |
| 33168_Y2167 | 1998 | UK | Pig Diseased | Systemic | 29 | 7 | [16] | PRJNA1009400 |
| 33170_Y2366 | 1997 | UK | Pig Diseased | Systemic | 1 | 2 | [16] | PRJNA1009400 |
| 33174_Y2440 | 1998 | UK | Pig Diseased | Respiratory | 29 | 7 | [16] | PRJNA1009400 |
| 33175_Y2567 | 1997 | UK | Pig Diseased | Respiratory | 87 | 8 | [16] | PRJNA1009400 |
| 33176_Y2573 | 1997 | UK | Pig Diseased | Systemic | NA | NA | [16] | PRJNA1009400 |
| 33177_Y2585 | 1997 | UK | Pig Diseased | Systemic | 29 | 4 | [16] | PRJNA1009400 |
| 33178_Y2590 | 1997 | UK | Pig Diseased | Systemic | 1 | NA | [16] | PRJNA1009400 |
| 33179_Y2598 | 1997 | UK | Pig Diseased | Respiratory | 87 | 8 | [16] | PRJNA1009400 |
| 33182_Y2701 | 1997 | UK | Pig Diseased | Systemic | NA | 7 | [16] | PRJNA1009400 |
| 4057-10_S10_L001 | 2019 | USA | Pig Diseased | Unknown | 1 | NA | [16] | PRJNA972671 |
| 4057-11_S11_L001 | 2019 | USA | Pig Diseased | Unknown | NA | 5 | [16] | PRJNA972671 |
| 4057-12_S12_L001 | 2019 | USA | Pig Diseased | Unknown | 108 | 23 | [16] | PRJNA972671 |
| 4057-13_S13_L001 | 2019 | USA | Pig Diseased | Unknown | NA | 21 | [16] | PRJNA972671 |
| 4057-14_S14_L001 | 2019 | USA | Pig Diseased | Unknown | 1 | NA | [16] | PRJNA972671 |
| 4057-15_S15_L001 | 2019 | USA | Pig Diseased | Unknown | 1 | NA | [16] | PRJNA972671 |
| 4057-16_S16_L001 | 2019 | USA | Pig Diseased | Unknown | 1 | 14 | [16] | PRJNA972671 |
| 4057-17_S17_L001 | 2019 | USA | Pig Diseased | Unknown | 1 | NA | [16] | PRJNA972671 |

| | | | | | | | | |
|---|---|---|---|---|---|---|---|---|
| 4057-18_S18_L001 | 2019 | USA | Pig Diseased | Unknown | 1 | NA | [16] | PRJNA972671 |
| 4057-19_S19_L001 | 2019 | USA | Pig Diseased | Unknown | 1 | 14 | [16] | PRJNA972671 |
| 4057-20_S20_L001 | 2019 | USA | Pig Diseased | Unknown | 1 | NA | [16] | PRJNA972671 |
| 4057-21_S21_L001 | 2019 | USA | Pig Diseased | Unknown | 1 | NA | [16] | PRJNA972671 |
| 4057-22_S22_L001 | 2019 | USA | Pig Diseased | Unknown | NA | 9 | [16] | PRJNA972671 |
| 4057-23_S23_L001 | 2019 | USA | Pig Diseased | Unknown | NA | 8 | [16] | PRJNA972671 |
| 4057-24_S24_L001 | 2019 | USA | Pig Diseased | Unknown | 29 | 7 | [16] | PRJNA972671 |
| 4057-25_S25_L001 | 2019 | USA | Pig Diseased | Unknown | 13 | NA | [16] | PRJNA972671 |
| 4057-26_S26_L001 | 2019 | USA | Pig Diseased | Unknown | 87 | 8 | [16] | PRJNA972671 |
| 4057-27_S27_L001 | 2019 | USA | Pig Diseased | Unknown | 28 | NA | [16] | PRJNA972671 |
| 4057-28_S28_L001 | 2019 | USA | Pig Diseased | Unknown | 25 | 2 | [16] | PRJNA972671 |
| 4057-30_S30_L001 | 2019 | USA | Pig Diseased | Unknown | NA | 9 | [16] | PRJNA972671 |
| 4057-31_S31_L001 | 2019 | USA | Pig Diseased | Unknown | 28 | 2 | [16] | PRJNA972671 |
| 4057-32_S32_L001 | 2019 | USA | Pig Diseased | Unknown | 119 | NA | [16] | PRJNA972671 |
| 4057-33_S33_L001 | 2019 | USA | Pig Diseased | Unknown | NA | 9 | [16] | PRJNA972671 |
| 4057-34_S34_L001 | 2019 | USA | Pig Diseased | Unknown | 25 | 2 | [16] | PRJNA972671 |
| 4057-35_S35_L001 | 2019 | USA | Pig Diseased | Unknown | 28 | 2 | [16] | PRJNA972671 |
| 4057-36_S36_L001 | 2019 | USA | Pig Diseased | Unknown | NA | 13 | [16] | PRJNA972671 |
| 4057-38_S38_L001 | 2019 | USA | Pig Diseased | Unknown | 94 | NA | [16] | PRJNA972671 |
| 4057-39_S39_L001 | 2019 | USA | Pig Diseased | Unknown | 17 | NA | [16] | PRJNA972671 |
| 4057-4_S4_L001 | 2019 | USA | Pig Diseased | Unknown | 1 | NA | [16] | PRJNA972671 |
| 4057-40_S40_L001 | 2019 | USA | Pig Diseased | Unknown | 119 | NA | [16] | PRJNA972671 |
| 4057-41_S41_L001 | 2019 | USA | Pig Diseased | Unknown | 119 | 5 | [16] | PRJNA972671 |
| 4057-42_S42_L001 | 2019 | USA | Pig Diseased | Unknown | 28 | 2 | [16] | PRJNA972671 |
| 4057-43_S43_L001 | 2019 | USA | Pig Diseased | Unknown | 28 | 2 | [16] | PRJNA972671 |
| 4057-45_S45_L001 | 2019 | USA | Pig Diseased | Unknown | 28 | NA | [16] | PRJNA972671 |

| | | | | | | | | |
|---|---|---|---|---|---|---|---|---|
| 4057-46_S46_L001 | 2019 | USA | Pig Diseased | Unknown | 25 | 2 | [16] | PRJNA972671 |
| 4057-47_S47_L001 | 2019 | USA | Pig Diseased | Unknown | 28 | 2 | [16] | PRJNA972671 |
| 4057-48_S48_L001 | 2019 | USA | Pig Diseased | Unknown | 25 | 2 | [16] | PRJNA972671 |
| 4057-49_S49_L001 | 2019 | USA | Pig Diseased | Unknown | 25 | 2 | [16] | PRJNA972671 |
| 4057-5_S5_L001 | 2019 | USA | Pig Diseased | Unknown | 1 | NA | [16] | PRJNA972671 |
| 4057-50_S50_L001 | 2019 | USA | Pig Diseased | Unknown | NA | 23 | [16] | PRJNA972671 |
| 4057-51_S51_L001 | 2019 | USA | Pig Diseased | Unknown | 25 | 2 | [16] | PRJNA972671 |
| 4057-52_S52_L001 | 2019 | USA | Pig Diseased | Unknown | 119 | 5 | [16] | PRJNA972671 |
| 4057-54_S54_L001 | 2019 | USA | Pig Diseased | Unknown | 119 | 5 | [16] | PRJNA972671 |
| 4057-55_S55_L001 | 2019 | USA | Pig Diseased | Unknown | 1 | NA | [16] | PRJNA972671 |
| 4057-56_S56_L001 | 2019 | USA | Pig Diseased | Unknown | 119 | NA | [16] | PRJNA972671 |
| 4057-57_S57_L001 | 2019 | USA | Pig Diseased | Unknown | 119 | NA | [16] | PRJNA972671 |
| 4057-6_S6_L001 | 2019 | USA | Pig Diseased | Unknown | 1 | NA | [16] | PRJNA972671 |
| 4057-60_S60_L001 | 2019 | USA | Pig Diseased | Unknown | NA | NA | [16] | PRJNA972671 |
| 4057-61_S61_L001 | 2019 | USA | Pig Diseased | Unknown | NA | NA | [16] | PRJNA972671 |
| 4057-62_S62_L001 | 2019 | USA | Pig Diseased | Unknown | NA | NA | [16] | PRJNA972671 |
| 4057-63_S63_L001 | 2019 | USA | Pig Diseased | Unknown | 28 | NA | [16] | PRJNA972671 |
| 4057-64_S64_L001 | 2019 | USA | Pig Diseased | Unknown | 28 | NA | [16] | PRJNA972671 |
| 4057-65_S65_L001 | 2019 | USA | Pig Diseased | Unknown | 87 | 8 | [16] | PRJNA972671 |
| 4057-66_S66_L001 | 2019 | USA | Pig Diseased | Unknown | 28 | NA | [16] | PRJNA972671 |
| 4057-67_S67_L001 | 2019 | USA | Pig Diseased | Unknown | 1 | 14 | [16] | PRJNA972671 |
| 4057-68_S68_L001 | 2019 | USA | Pig Diseased | Unknown | 1 | NA | [16] | PRJNA972671 |
| 4057-69_S69_L001 | 2019 | USA | Pig Diseased | Unknown | NA | NA | [16] | PRJNA972671 |
| 4057-7_S7_L001 | 2019 | USA | Pig Diseased | Unknown | NA | 2 | [16] | PRJNA972671 |
| 4057-70_S70_L001 | 2019 | USA | Pig Diseased | Unknown | 28 | NA | [16] | PRJNA972671 |
| 4057-71_S71_L001 | 2019 | USA | Pig Diseased | Unknown | 28 | NA | [16] | PRJNA972671 |

| | | | | | | | | |
|---|---|---|---|---|---|---|---|---|
| 4057-72_S72_L001 | 2019 | USA | Pig Diseased | Unknown | 87 | 8 | [16] | PRJNA972671 |
| 4057-73_S73_L001 | 2019 | USA | Pig Diseased | Unknown | 94 | NA | [16] | PRJNA972671 |
| 4057-74_S74_L001 | 2019 | USA | Pig Diseased | Unknown | NA | 10 | [16] | PRJNA972671 |
| 4057-75_S75_L001 | 2019 | USA | Pig Diseased | Unknown | NA | 9 | [16] | PRJNA972671 |
| 4057-76_S76_L001 | 2019 | USA | Pig Diseased | Unknown | NA | 9 | [16] | PRJNA972671 |
| 4057-77_S77_L001 | 2019 | USA | Pig Diseased | Unknown | NA | 7 | [16] | PRJNA972671 |
| 4057-78_S78_L001 | 2019 | USA | Pig Diseased | Unknown | NA | 8 | [16] | PRJNA972671 |
| 4057-79_S79_L001 | 2019 | USA | Pig Diseased | Unknown | 54 | NA | [16] | PRJNA972671 |
| 4057-8_S8_L001 | 2019 | USA | Pig Diseased | Unknown | 87 | 8 | [16] | PRJNA972671 |
| 4057-80_S80_L001 | 2019 | USA | Pig Diseased | Unknown | NA | NA | [16] | PRJNA972671 |
| 4057-82_S82_L001 | 2019 | USA | Pig Diseased | Unknown | 54 | NA | [16] | PRJNA972671 |
| 4057-9_S9_L001 | 2019 | USA | Pig Diseased | Unknown | 1 | NA | [16] | PRJNA972671 |
| A-04-3-TT-01-SS-C2 | NA | Myanmar | Pig Unknown | Unknown | NA | 30 | [16] | PRJNA1012585 |
| A-05-4-BL-01-SS-C1 | NA | Myanmar | Pig Unknown | Unknown | NA | 15 | [16] | PRJNA1012585 |
| A-05-5-TT-01-SS-C1 | NA | Myanmar | Pig Unknown | Unknown | NA | 28 | [16] | PRJNA1012585 |
| A-05-5-TT-02-SS-C5 | NA | Myanmar | Pig Unknown | Unknown | NA | 15 | [16] | PRJNA1012585 |
| A-07-3-TT-01-SS-C2 | NA | Myanmar | Pig Unknown | Unknown | NA | 31 | [16] | PRJNA1012585 |
| A-07-3-TT-01-SS-C3 | NA | Myanmar | Pig Unknown | Unknown | NA | 31 | [16] | PRJNA1012585 |
| A-07-3-TT-01-SS-C4 | NA | Myanmar | Pig Unknown | Unknown | NA | 31 | [16] | PRJNA1012585 |
| A-07-3-TT-02-SS-C1 | NA | Myanmar | Pig Unknown | Unknown | NA | 31 | [16] | PRJNA1012585 |
| A-07-3-TT-02-SS-C2 | NA | Myanmar | Pig Unknown | Unknown | NA | 31 | [16] | PRJNA1012585 |
| A-07-3-TT-02-SS-C3 | NA | Myanmar | Pig Unknown | Unknown | NA | 31 | [16] | PRJNA1012585 |
| A-07-3-TT-02-SS-C4 | NA | Myanmar | Pig Unknown | Unknown | NA | 31 | [16] | PRJNA1012585 |
| A-07-3-TT-02-SS-C5 | NA | Myanmar | Pig Unknown | Unknown | NA | 29 | [16] | PRJNA1012585 |
| A-08-1-TT-01-SS-C4 | NA | Myanmar | Pig Unknown | Unknown | 225 | 7 | [16] | PRJNA1012585 |
| A-09-1-TT-09-SS-C3 | NA | Myanmar | Pig Unknown | Unknown | NA | 29 | [16] | PRJNA1012585 |

| | | | | | | | | |
|---|---|---|---|---|---|---|---|---|
| A-09-7-TT-04-SS-C1 | NA | Myanmar | Pig Unknown | Unknown | NA | 31 | [16] | PRJNA1012585 |
| A-09-7-TT-05-SS-C2 | NA | Myanmar | Pig Unknown | Unknown | NA | 5 | [16] | PRJNA1012585 |
| A-09-7-TT-05-SS-C3 | NA | Myanmar | Pig Unknown | Unknown | NA | 5 | [16] | PRJNA1012585 |
| A-09-8-LN-03-SS-C3 | NA | Myanmar | Pig Unknown | Unknown | NA | NA | [16] | PRJNA1012585 |
| A-09-9-BL-03-SS-C2 | NA | Myanmar | Pig Unknown | Unknown | NA | NA | [16] | PRJNA1012585 |
| A-09-9-TT-02-SS-C5 | NA | Myanmar | Pig Unknown | Unknown | NA | 15 | [16] | PRJNA1012585 |
| A-09-9-TT-04-SS-C2 | NA | Myanmar | Pig Unknown | Unknown | NA | 31 | [16] | PRJNA1012585 |
| A-09-9-TT-04-SS-C4 | NA | Myanmar | Pig Unknown | Unknown | NA | 31 | [16] | PRJNA1012585 |
| DNS31_ref | 2018 | Denmark | Pig Diseased | Systemic | 28 | 2 | [16] | PRJNA763404 |
| F-01-2-TS-01-SS-C1 | NA | Myanmar | Pig Healthy | Non-Clinical | NA | NA | [16] | PRJNA1012585 |
| F-01-2-TS-01-SS-C4 | NA | Myanmar | Pig Healthy | Non-Clinical | NA | 19 | [16] | PRJNA1012585 |
| F-01-2-TS-01-SS-C5 | NA | Myanmar | Pig Healthy | Non-Clinical | NA | NA | [16] | PRJNA1012585 |
| F-01-2-TS-02-SS-C1 | NA | Myanmar | Pig Healthy | Non-Clinical | NA | NA | [16] | PRJNA1012585 |
| F-01-2-TS-03-SS-C3 | NA | Myanmar | Pig Healthy | Non-Clinical | NA | 19 | [16] | PRJNA1012585 |
| F-01-2-TS-03-SS-C4 | NA | Myanmar | Pig Healthy | Non-Clinical | NA | NA | [16] | PRJNA1012585 |
| F-01-2-TS-07-SS-C4 | NA | Myanmar | Pig Healthy | Non-Clinical | NA | NA | [16] | PRJNA1012585 |
| F-01-2-TS-09-SS-C1 | NA | Myanmar | Pig Healthy | Non-Clinical | NA | NA | [16] | PRJNA1012585 |
| F-01-2-TS-10-SS-C1 | NA | Myanmar | Pig Healthy | Non-Clinical | NA | 19 | [16] | PRJNA1012585 |
| F-01-2-TS-10-SS-C4 | NA | Myanmar | Pig Healthy | Non-Clinical | NA | NA | [16] | PRJNA1012585 |
| F-01-3-TS-08-SS-C1 | NA | Myanmar | Pig Healthy | Non-Clinical | NA | 29 | [16] | PRJNA1012585 |
| F-02-1-TS-01-SS-C5 | NA | Myanmar | Pig Healthy | Non-Clinical | NA | NA | [16] | PRJNA1012585 |
| F-02-2-TS-07-SS-C4 | NA | Myanmar | Pig Healthy | Non-Clinical | NA | NA | [16] | PRJNA1012585 |
| F-02-2-TS-08-SS-C1 | NA | Myanmar | Pig Healthy | Non-Clinical | NA | NA | [16] | PRJNA1012585 |
| F-02-3-TS-06-SS-C1 | NA | Myanmar | Pig Healthy | Non-Clinical | NA | NA | [16] | PRJNA1012585 |
| F-02-3-TS-06-SS-C4 | NA | Myanmar | Pig Healthy | Non-Clinical | NA | NA | [16] | PRJNA1012585 |
| F-02-3-TS-07-SS-C3 | NA | Myanmar | Pig Healthy | Non-Clinical | NA | NA | [16] | PRJNA1012585 |

| | | | | | | | | |
|---|---|---|---|---|---|---|---|---|
| F-02-3-TS-08-SS-C3 | NA | Myanmar | Pig Healthy | Non-Clinical | NA | 21 | [16] | PRJNA1012585 |
| F-02-3-TS-09-SS-C5 | NA | Myanmar | Pig Healthy | Non-Clinical | NA | NA | [16] | PRJNA1012585 |
| F-02-3-TS-10-SS-C2 | NA | Myanmar | Pig Healthy | Non-Clinical | NA | 29 | [16] | PRJNA1012585 |
| F-02-3-TS-10-SS-C4 | NA | Myanmar | Pig Healthy | Non-Clinical | NA | NA | [16] | PRJNA1012585 |
| F-02-3-TS-10-SS-C5 | NA | Myanmar | Pig Healthy | Non-Clinical | NA | NA | [16] | PRJNA1012585 |
| F-03-3-TS-04-SS-C1 | NA | Myanmar | Pig Healthy | Non-Clinical | NA | 5 | [16] | PRJNA1012585 |
| F-03-3-TS-09-SS-C1 | NA | Myanmar | Pig Healthy | Non-Clinical | NA | 5 | [16] | PRJNA1012585 |
| F-04-2-TS-05-SS-C1 | NA | Myanmar | Pig Healthy | Non-Clinical | NA | NA | [16] | PRJNA1012585 |
| F-04-2-TS-06-SS-C3 | NA | Myanmar | Pig Healthy | Non-Clinical | NA | 24 | [16] | PRJNA1012585 |
| F-04-2-TS-06-SS-C4 | NA | Myanmar | Pig Healthy | Non-Clinical | NA | 24 | [16] | PRJNA1012585 |
| F-04-2-TS-08-SS-C5 | NA | Myanmar | Pig Healthy | Non-Clinical | NA | 24 | [16] | PRJNA1012585 |
| F-04-2-TS-09-SS-C4 | NA | Myanmar | Pig Healthy | Non-Clinical | NA | NA | [16] | PRJNA1012585 |
| F-04-2-TS-10-SS-C1 | NA | Myanmar | Pig Healthy | Non-Clinical | NA | NA | [16] | PRJNA1012585 |
| F-04-3-TS-07-SS-C4 | NA | Myanmar | Pig Healthy | Non-Clinical | NA | 30 | [16] | PRJNA1012585 |
| F-04-3-TS-07-SS-C5 | NA | Myanmar | Pig Healthy | Non-Clinical | NA | 31 | [16] | PRJNA1012585 |
| F-05-1-TS-01-SS-C4 | NA | Myanmar | Pig Healthy | Non-Clinical | NA | 30 | [16] | PRJNA1012585 |
| F-05-1-TS-06-SS-C2 | NA | Myanmar | Pig Healthy | Non-Clinical | NA | 30 | [16] | PRJNA1012585 |
| F-05-1-TS-08-SS-C3 | NA | Myanmar | Pig Healthy | Non-Clinical | NA | 30 | [16] | PRJNA1012585 |
| F-05-3-TS-02-SS-C5 | NA | Myanmar | Pig Healthy | Non-Clinical | NA | 30 | [16] | PRJNA1012585 |
| F-07-3-TS-07-SS-C3 | NA | Myanmar | Pig Healthy | Non-Clinical | NA | 29 | [16] | PRJNA1012585 |
| F-09-1-TS-05-SS-C2 | NA | Myanmar | Pig Healthy | Non-Clinical | NA | NA | [16] | PRJNA1012585 |
| F-09-2-TS-05-SS-C4 | NA | Myanmar | Pig Healthy | Non-Clinical | NA | NA | [16] | PRJNA1012585 |
| F-09-2-TS-07-SS-C1 | NA | Myanmar | Pig Healthy | Non-Clinical | NA | NA | [16] | PRJNA1012585 |
| F-09-2-TS-07-SS-C2 | NA | Myanmar | Pig Healthy | Non-Clinical | NA | 30 | [16] | PRJNA1012585 |
| F-09-2-TS-10-SS-C4 | NA | Myanmar | Pig Healthy | Non-Clinical | NA | 30 | [16] | PRJNA1012585 |
| F-09-3-TS-04-SS-C3 | NA | Myanmar | Pig Healthy | Non-Clinical | NA | NA | [16] | PRJNA1012585 |

| | | | | | | | | |
|---|---|---|---|---|---|---|---|---|
| F-09-3-TS-05-SS-C2 | NA | Myanmar | Pig Healthy | Non-Clinical | NA | 29 | [16] | PRJNA1012585 |
| F-09-3-TS-05-SS-C5 | NA | Myanmar | Pig Healthy | Non-Clinical | NA | 29 | [16] | PRJNA1012585 |
| F-09-3-TS-06-SS-C3 | NA | Myanmar | Pig Healthy | Non-Clinical | NA | NA | [16] | PRJNA1012585 |
| F-09-3-TS-08-SS-C5 | NA | Myanmar | Pig Healthy | Non-Clinical | NA | NA | [16] | PRJNA1012585 |
| F-09-3-TS-09-SS-C1 | NA | Myanmar | Pig Healthy | Non-Clinical | NA | 29 | [16] | PRJNA1012585 |
| F-09-3-TS-09-SS-C3 | NA | Myanmar | Pig Healthy | Non-Clinical | NA | NA | [16] | PRJNA1012585 |
| F-11-1-TS-03-SS-C5 | NA | Myanmar | Pig Healthy | Non-Clinical | NA | NA | [16] | PRJNA1012585 |
| F-11-1-TS-08-SS-C1 | NA | Myanmar | Pig Healthy | Non-Clinical | NA | NA | [16] | PRJNA1012585 |
| F-11-1-TS-08-SS-C3 | NA | Myanmar | Pig Healthy | Non-Clinical | NA | NA | [16] | PRJNA1012585 |
| F-11-2-TS-01-SS-C1 | NA | Myanmar | Pig Healthy | Non-Clinical | NA | NA | [16] | PRJNA1012585 |
| F-11-2-TS-03-SS-C1 | NA | Myanmar | Pig Healthy | Non-Clinical | NA | NA | [16] | PRJNA1012585 |
| F-11-2-TS-05-SS-C2 | NA | Myanmar | Pig Healthy | Non-Clinical | NA | NA | [16] | PRJNA1012585 |
| F-11-2-TS-05-SS-C3 | NA | Myanmar | Pig Healthy | Non-Clinical | NA | NA | [16] | PRJNA1012585 |
| F-11-2-TS-05-SS-C5 | NA | Myanmar | Pig Healthy | Non-Clinical | NA | NA | [16] | PRJNA1012585 |
| F-11-2-TS-08-SS-C1 | NA | Myanmar | Pig Healthy | Non-Clinical | NA | 15 | [16] | PRJNA1012585 |
| F-11-2-TS-08-SS-C4 | NA | Myanmar | Pig Healthy | Non-Clinical | NA | 15 | [16] | PRJNA1012585 |
| F-11-2-TS-08-SS-C5 | NA | Myanmar | Pig Healthy | Non-Clinical | NA | 15 | [16] | PRJNA1012585 |
| F-11-2-TS-10-SS-C2 | NA | Myanmar | Pig Healthy | Non-Clinical | NA | 30 | [16] | PRJNA1012585 |
| F-11-2-TS-10-SS-C5 | NA | Myanmar | Pig Healthy | Non-Clinical | NA | NA | [16] | PRJNA1012585 |
| F-12-1-TS-07-SS-C1 | NA | Myanmar | Pig Healthy | Non-Clinical | NA | 30 | [16] | PRJNA1012585 |
| F-12-1-TS-09-SS-C4 | NA | Myanmar | Pig Healthy | Non-Clinical | NA | NA | [16] | PRJNA1012585 |
| F-12-2-TS-02-SS-C4 | NA | Myanmar | Pig Healthy | Non-Clinical | NA | 31 | [16] | PRJNA1012585 |
| F-12-3-TS-02-SS-C1 | NA | Myanmar | Pig Healthy | Non-Clinical | NA | NA | [16] | PRJNA1012585 |
| F-12-3-TS-03-SS-C2 | NA | Myanmar | Pig Healthy | Non-Clinical | NA | 15 | [16] | PRJNA1012585 |
| F-12-3-TS-04-SS-C1 | NA | Myanmar | Pig Healthy | Non-Clinical | NA | 16 | [16] | PRJNA1012585 |
| F-12-3-TS-04-SS-C2 | NA | Myanmar | Pig Healthy | Non-Clinical | NA | 15 | [16] | PRJNA1012585 |

| | | | | | | | | |
|---|---|---|---|---|---|---|---|---|
| F-12-3-TS-04-SS-C3 | NA | Myanmar | Pig Healthy | Non-Clinical | NA | 15 | 16 | PRJNA1012585 |
| F-12-3-TS-04-SS-C4 | NA | Myanmar | Pig Healthy | Non-Clinical | NA | 15 | 16 | PRJNA1012585 |
| F-12-3-TS-04-SS-C5 | NA | Myanmar | Pig Healthy | Non-Clinical | NA | NA | 16 | PRJNA1012585 |
| F-12-3-TS-05-SS-C1 | NA | Myanmar | Pig Healthy | Non-Clinical | NA | 29 | 16 | PRJNA1012585 |
| F-13-1-TS-02-SS-C1 | NA | Myanmar | Pig Healthy | Non-Clinical | NA | NA | 16 | PRJNA1012585 |
| F-13-2-TS-01-SS-C4 | NA | Myanmar | Pig Healthy | Non-Clinical | NA | NA | 16 | PRJNA1012585 |
| F-13-2-TS-02-SS-C2 | NA | Myanmar | Pig Healthy | Non-Clinical | NA | NA | 16 | PRJNA1012585 |
| F-16-1-TS-03-SS-C3 | NA | Myanmar | Pig Healthy | Non-Clinical | NA | NA | 16 | PRJNA1012585 |
| F-16-1-TS-03-SS-C4 | NA | Myanmar | Pig Healthy | Non-Clinical | NA | NA | 16 | PRJNA1012585 |
| F-17-2-TS-02-SS-C3 | NA | Myanmar | Pig Healthy | Non-Clinical | NA | NA | 16 | PRJNA1012585 |
| F-17-2-TS-02-SS-C4 | NA | Myanmar | Pig Healthy | Non-Clinical | NA | NA | 16 | PRJNA1012585 |
| F-17-2-TS-02-SS-C5 | NA | Myanmar | Pig Healthy | Non-Clinical | NA | NA | 16 | PRJNA1012585 |
| F-17-2-TS-04-SS-C3 | NA | Myanmar | Pig Healthy | Non-Clinical | NA | NA | 16 | PRJNA1012585 |
| F-17-2-TS-07-SS-C5 | NA | Myanmar | Pig Healthy | Non-Clinical | NA | NA | 16 | PRJNA1012585 |
| F-27-2-TS-04-SS-C2 | NA | Myanmar | Pig Healthy | Non-Clinical | NA | 11 | 16 | PRJNA1012585 |
| F-28-2-TS-01-SS-C3 | NA | Myanmar | Pig Healthy | Non-Clinical | NA | NA | 16 | PRJNA1012585 |
| F-28-2-TS-02-SS-C1 | NA | Myanmar | Pig Healthy | Non-Clinical | NA | NA | 16 | PRJNA1012585 |
| F-28-2-TS-02-SS-C3 | NA | Myanmar | Pig Healthy | Non-Clinical | NA | NA | 16 | PRJNA1012585 |
| F-28-2-TS-02-SS-C5 | NA | Myanmar | Pig Healthy | Non-Clinical | NA | NA | 16 | PRJNA1012585 |
| F-28-2-TS-03-SS-C1 | NA | Myanmar | Pig Healthy | Non-Clinical | NA | NA | 16 | PRJNA1012585 |
| F-28-2-TS-03-SS-C2 | NA | Myanmar | Pig Healthy | Non-Clinical | NA | NA | 16 | PRJNA1012585 |
| F-28-2-TS-03-SS-C3 | NA | Myanmar | Pig Healthy | Non-Clinical | NA | NA | 16 | PRJNA1012585 |
| F-28-2-TS-05-SS-C3 | NA | Myanmar | Pig Healthy | Non-Clinical | NA | NA | 16 | PRJNA1012585 |
| F-28-2-TS-06-SS-C4 | NA | Myanmar | Pig Healthy | Non-Clinical | NA | NA | 16 | PRJNA1012585 |
| F-28-2-TS-07-SS-C1 | NA | Myanmar | Pig Healthy | Non-Clinical | NA | 29 | 16 | PRJNA1012585 |
| F-28-2-TS-07-SS-C3 | NA | Myanmar | Pig Healthy | Non-Clinical | NA | NA | 16 | PRJNA1012585 |

| | | | | | | | | |
|---|---|---|---|---|---|---|---|---|
| M104170_C2_ref | 2017 | Spain | Pig Healthy | Non-Clinical | NA | 21 | [16] | PRJNA1010137 |
| NLR11_ref | 2017 | Netherlands | Pig Diseased | Respiratory | 87 | 8 | [16] | PRJNA1009400 |
| NLS40_ref | 2017 | Netherlands | Pig Diseased | Systemic | 13 | 14 | [16] | PRJNA1009400 |
| NLS47_ref | 2017 | Netherlands | Pig Diseased | Systemic | 17 | 4 | [16] | PRJNA763404 |
| NLS50_ref | 2017 | Netherlands | Pig Diseased | Systemic | 16 | 9 | [16] | PRJNA1009400 |
| PH2016-081_ref | 2016 | Germany | Pig Diseased | Respiratory | 15 | 3 | [16] | PRJNA1009400 |
| SPC15_ref | 2015 | Spain | Pig Healthy | Non-Clinical | NA | NA | [16] | PRJNA763404 |
| SS001 | 2019 | Myanmar | Pig Healthy | Non-Clinical | NA | NA | [16] | PRJNA1012585 |
| SS004 | 2019 | Myanmar | Pig Healthy | Non-Clinical | NA | 29 | [16] | PRJNA1012585 |
| SS007 | 2019 | Myanmar | Pig Healthy | Non-Clinical | NA | NA | [16] | PRJNA1012585 |
| SS013 | 2019 | Myanmar | Pig Healthy | Non-Clinical | NA | 29 | [16] | PRJNA1012585 |
| SS059 | 2019 | Myanmar | Pig Healthy | Non-Clinical | NA | 29 | [16] | PRJNA1012585 |
| SS063 | 2019 | Myanmar | Pig Healthy | Non-Clinical | NA | NA | [16] | PRJNA1012585 |
| SS064 | 2019 | Myanmar | Pig Healthy | Non-Clinical | NA | 29 | [16] | PRJNA1012585 |
| SS065 | 2019 | Myanmar | Pig Healthy | Non-Clinical | NA | 29 | [16] | PRJNA1012585 |
| SS066 | 2019 | Myanmar | Pig Healthy | Non-Clinical | NA | 4 | [16] | PRJNA1012585 |
| SS070 | 2019 | Myanmar | Pig Healthy | Non-Clinical | NA | 30 | [16] | PRJNA1012585 |
| SS071 | 2019 | Myanmar | Pig Healthy | Non-Clinical | NA | NA | [16] | PRJNA1012585 |
| SS080 | 2019 | Myanmar | Pig Healthy | Non-Clinical | NA | 5 | [16] | PRJNA1012585 |
| SS082 | 2019 | Myanmar | Pig Healthy | Non-Clinical | NA | 29 | [16] | PRJNA1012585 |
| SS086 | 2019 | Myanmar | Pig Healthy | Non-Clinical | NA | NA | [16] | PRJNA1012585 |
| SS087 | 2019 | Myanmar | Pig Healthy | Non-Clinical | NA | NA | [16] | PRJNA1012585 |
| SS089 | 2019 | Myanmar | Pig Healthy | Non-Clinical | NA | NA | [16] | PRJNA1012585 |
| SS101m | 2019 | Myanmar | Pig Healthy | Non-Clinical | NA | NA | [16] | PRJNA1012585 |
| SS108 | 2019 | Myanmar | Pig Healthy | Non-Clinical | NA | 4 | [16] | PRJNA1012585 |
| SS113 | 2019 | Myanmar | Pig Healthy | Non-Clinical | 117 | 3 | [16] | PRJNA1012585 |

| | | | | | | | | |
|---|---|---|---|---|---|---|---|---|
| SS115 | 2019 | Myanmar | Pig Healthy | Non-Clinical | NA | 5 | [16] | PRJNA1012585 |
| SS149 | 2019 | Myanmar | Pig Healthy | Non-Clinical | NA | 21 | [16] | PRJNA1012585 |
| SS150 | 2019 | Myanmar | Pig Healthy | Non-Clinical | NA | NA | [16] | PRJNA1012585 |
| SS154 | 2019 | Myanmar | Pig Healthy | Non-Clinical | NA | NA | [16] | PRJNA1012585 |
| SS155 | 2019 | Myanmar | Pig Healthy | Non-Clinical | NA | NA | [16] | PRJNA1012585 |
| SS160 | 2019 | Myanmar | Pig Healthy | Non-Clinical | NA | NA | [16] | PRJNA1012585 |
| SS161 | 2019 | Myanmar | Pig Healthy | Non-Clinical | NA | 11 | [16] | PRJNA1012585 |
| SS163 | 2019 | Myanmar | Pig Healthy | Non-Clinical | NA | NA | [16] | PRJNA1012585 |
| SS173 | 2019 | Myanmar | Pig Healthy | Non-Clinical | NA | NA | [16] | PRJNA1012585 |
| SS174 | 2019 | Myanmar | Pig Healthy | Non-Clinical | NA | NA | [16] | PRJNA1012585 |
| SS176 | 2019 | Myanmar | Pig Healthy | Non-Clinical | NA | 4 | [16] | PRJNA1012585 |
| SS177 | 2019 | Myanmar | Pig Healthy | Non-Clinical | NA | 4 | [16] | PRJNA1012585 |
| SS179 | 2019 | Myanmar | Pig Healthy | Non-Clinical | NA | NA | [16] | PRJNA1012585 |
| SS183 | 2019 | Myanmar | Pig Healthy | Non-Clinical | NA | NA | [16] | PRJNA1012585 |
| SS184 | 2019 | Myanmar | Pig Healthy | Non-Clinical | NA | NA | [16] | PRJNA1012585 |
| SS185 | 2019 | Myanmar | Pig Healthy | Non-Clinical | NA | 29 | [16] | PRJNA1012585 |
| SS186 | 2019 | Myanmar | Pig Healthy | Non-Clinical | NA | NA | [16] | PRJNA1012585 |
| SS189 | 2019 | Myanmar | Pig Healthy | Non-Clinical | NA | NA | [16] | PRJNA1012585 |
| SS190 | 2019 | Myanmar | Pig Healthy | Non-Clinical | NA | 29 | [16] | PRJNA1012585 |
| SS199 | 2019 | Myanmar | Pig Healthy | Non-Clinical | NA | 30 | [16] | PRJNA1012585 |
| SS202 | 2019 | Myanmar | Pig Healthy | Non-Clinical | NA | 15 | [16] | PRJNA1012585 |
| SS208 | 2019 | Myanmar | Pig Healthy | Non-Clinical | NA | 24 | [16] | PRJNA1012585 |
| SS213 | 2019 | Myanmar | Pig Healthy | Non-Clinical | NA | NA | [16] | PRJNA1012585 |
| SS217 | 2019 | Myanmar | Pig Healthy | Non-Clinical | NA | 4 | [16] | PRJNA1012585 |
| SS220 | 2019 | Myanmar | Pig Healthy | Non-Clinical | NA | 31 | [16] | PRJNA1012585 |
| SS221 | 2019 | Myanmar | Pig Healthy | Non-Clinical | NA | 29 | [16] | PRJNA1012585 |

| | | | | | | | | |
|---|---|---|---|---|---|---|---|---|
| SS222 | 2019 | Myanmar | Pig Healthy | Non-Clinical | NA | NA | [16] | PRJNA1012585 |
| SS223 | 2019 | Myanmar | Pig Healthy | Non-Clinical | NA | 31 | [16] | PRJNA1012585 |
| SS224 | 2019 | Myanmar | Pig Healthy | Non-Clinical | NA | NA | [16] | PRJNA1012585 |
| SS225 | 2019 | Myanmar | Pig Healthy | Non-Clinical | NA | 29 | [16] | PRJNA1012585 |
| SS229 | 2019 | Myanmar | Pig Healthy | Non-Clinical | NA | 16 | [16] | PRJNA1012585 |
| SS232 | 2019 | Myanmar | Pig Healthy | Non-Clinical | NA | 29 | [16] | PRJNA1012585 |
| SS236 | 2019 | Myanmar | Pig Healthy | Non-Clinical | NA | 29 | [16] | PRJNA1012585 |
| SS238 | 2019 | Myanmar | Pig Healthy | Non-Clinical | 221 | 24 | [16] | PRJNA1012585 |
| SS241 | 2019 | Myanmar | Pig Healthy | Non-Clinical | NA | NA | [16] | PRJNA1012585 |
| Y0191 | 1999 | UK | Pig Diseased | Respiratory | 30 | 3 | [16] | PRJNA1009400 |
| Y0493 | 1990 | UK | Pig Diseased | Systemic | 1 | 2 | [16] | PRJNA1009400 |
| Y0494 | 1989 | UK | Pig Diseased | Systemic | 1 | 2 | [16] | PRJNA1009400 |
| Y2001 | 1995 | UK | Pig Diseased | Respiratory | 27 | 3 | [16] | PRJNA1009400 |
| Y2116 | 1996 | UK | Pig Diseased | Respiratory | 23 | 4 | [16] | PRJNA1009400 |
| Y2403 | 1998 | UK | Pig Diseased | Systemic | 27 | 3 | [16] | PRJNA1009400 |
| Y2576 | 1997 | UK | Pig Diseased | Systemic | 1 | 14 | [16] | PRJNA1009400 |
| Y2587 | 1997 | UK | Pig Diseased | Systemic | 27 | 3 | [16] | PRJNA1009400 |
| SRR7369571 | 2017 | Australia | Pig Unknown | Unknown | 28 | 2 | [33] | PRJNA476804 |
| SRR7369572 | 2017 | Australia | Pig Unknown | Unknown | NA | 21 | [33] | PRJNA476804 |
| SRR7369573 | 2017 | Australia | Pig Unknown | Unknown | NA | 16 | [33] | PRJNA476804 |
| SRR7369574 | 2017 | Australia | Pig Unknown | Unknown | NA | 21 | [33] | PRJNA476804 |
| SRR7369575 | 2017 | Australia | Pig Unknown | Unknown | 27 | 3 | [33] | PRJNA476804 |
| SRR7369576 | 2017 | Australia | Pig Unknown | Unknown | NA | 21 | [33] | PRJNA476804 |
| SRR7369577 | 2017 | Australia | Pig Unknown | Unknown | NA | 9 | [33] | PRJNA476804 |
| SRR7369578 | 2017 | Australia | Pig Unknown | Unknown | NA | 31 | [33] | PRJNA476804 |
| SRR7369579 | 2017 | Australia | Pig Unknown | Unknown | NA | 2 | [33] | PRJNA476804 |

| | | | | | | | | |
|---|---|---|---|---|---|---|---|---|
| SRR7369580 | 2017 | Australia | Pig Unknown | Unknown | NA | 4 | [33] | PRJNA476804 |
| SRR7369581 | 2013 | Australia | Pig Unknown | Unknown | NA | 16 | [33] | PRJNA476804 |
| SRR7369582 | 2013 | Australia | Pig Unknown | Unknown | 1 | 2 | [33] | PRJNA476804 |
| SRR7369583 | 2015 | Australia | Pig Unknown | Unknown | 1 | 2 | [33] | PRJNA476804 |
| SRR7369585 | 2015 | Australia | Pig Unknown | Unknown | 1 | 2 | [33] | PRJNA476804 |
| SRR7369586 | 2015 | Australia | Pig Unknown | Unknown | 1 | 2 | [33] | PRJNA476804 |
| SRR7369587 | 2010 | Australia | Pig Unknown | Unknown | 1 | 2 | [33] | PRJNA476804 |
| SRR7369588 | 2015 | Australia | Pig Unknown | Unknown | 483 | 5 | [33] | PRJNA476804 |
| SRR7369589 | NA | Australia | Pig Unknown | Unknown | 1 | 2 | [33] | PRJNA476804 |
| SRR7369590 | 2016 | Australia | Pig Unknown | Unknown | 94 | 4 | [33] | PRJNA476804 |
| SRR7369591 | 2017 | Australia | Pig Unknown | Unknown | NA | NA | [33] | PRJNA476804 |
| SRR7369592 | 2017 | Australia | Pig Unknown | Unknown | 87 | 8 | [33] | PRJNA476804 |
| SRR7369593 | 2017 | Australia | Pig Unknown | Unknown | NA | 12 | [33] | PRJNA476804 |
| SRR7369594 | 2017 | Australia | Pig Unknown | Unknown | NA | 16 | [33] | PRJNA476804 |
| SRR7369595 | 2017 | Australia | Pig Unknown | Unknown | 27 | 3 | [33] | PRJNA476804 |
| SRR7369596 | 2017 | Australia | Pig Unknown | Unknown | 483 | NA | [33] | PRJNA476804 |
| SRR7369597 | 2017 | Australia | Pig Unknown | Unknown | NA | 16 | [33] | PRJNA476804 |
| SRR7369598 | 2017 | Australia | Pig Unknown | Unknown | 483 | NA | [33] | PRJNA476804 |
| SRR7369599 | 2017 | Australia | Pig Unknown | Unknown | NA | 31 | [33] | PRJNA476804 |
| SRR7369600 | 2017 | Australia | Pig Unknown | Unknown | 27 | 3 | [33] | PRJNA476804 |
| SRR7369601 | 2014 | Australia | Pig Unknown | Unknown | 28 | 2 | [33] | PRJNA476804 |
| SRR7369602 | 2014 | Australia | Pig Unknown | Unknown | NA | 31 | [33] | PRJNA476804 |
| SRR7369603 | 2014 | Australia | Pig Unknown | Unknown | 28 | 2 | [33] | PRJNA476804 |
| SRR7369604 | 2014 | Australia | Pig Unknown | Unknown | NA | 4 | [33] | PRJNA476804 |
| SRR7369606 | 2014 | Australia | Pig Unknown | Unknown | NA | NA | [33] | PRJNA476804 |
| SRR7369607 | 2014 | Australia | Pig Unknown | Unknown | NA | 11 | [33] | PRJNA476804 |

| | | | | | | | | |
|---|---|---|---|---|---|---|---|---|
| SRR7369608 | 2014 | Australia | Pig Unknown | Unknown | 27 | 3 | [33] | PRJNA476804 |
| SRR7369609 | 2014 | Australia | Pig Unknown | Unknown | 13 | 14 | [33] | PRJNA476804 |
| SRR7369610 | 2014 | Australia | Pig Unknown | Unknown | NA | 15 | [33] | PRJNA476804 |
| SRR7369611 | NA | Australia | Pig Unknown | Unknown | 483 | 5 | [33] | PRJNA476804 |
| SRR7369612 | 2014 | Australia | Pig Unknown | Unknown | 25 | NA | [33] | PRJNA476804 |
| SRR7369613 | 2014 | Australia | Pig Unknown | Unknown | 117 | 3 | [33] | PRJNA476804 |
| SRR7369614 | 2014 | Australia | Pig Unknown | Unknown | 28 | 2 | [33] | PRJNA476804 |
| SRR7369615 | 2017 | Australia | Pig Unknown | Unknown | 25 | NA | [33] | PRJNA476804 |
| SRR7369616 | 2017 | Australia | Pig Unknown | Unknown | 25 | 2 | [33] | PRJNA476804 |
| SRR7369617 | 2017 | Australia | Pig Unknown | Unknown | NA | 2 | [33] | PRJNA476804 |
| SRR7369618 | 2017 | Australia | Pig Unknown | Unknown | NA | 2 | [33] | PRJNA476804 |
| SRR7369619 | 2017 | Australia | Pig Unknown | Unknown | 25 | 2 | [33] | PRJNA476804 |
| SRR7369620 | 2017 | Australia | Pig Unknown | Unknown | 25 | 2 | [33] | PRJNA476804 |
| SRR7369621 | 2010 | Australia | Pig Unknown | Unknown | 27 | 3 | [33] | PRJNA476804 |
| SRR7369622 | 2013 | Australia | Pig Unknown | Unknown | 87 | 8 | [33] | PRJNA476804 |
| SRR7369623 | 2016 | Australia | Pig Unknown | Unknown | 25 | 2 | [33] | PRJNA476804 |
| SRR7369624 | 2016 | Australia | Pig Unknown | Unknown | 28 | 2 | [33] | PRJNA476804 |
| SRR7369625 | 2016 | Australia | Pig Unknown | Unknown | NA | 2 | [33] | PRJNA476804 |
| SRR7369626 | 2016 | Australia | Pig Unknown | Unknown | NA | 3 | [33] | PRJNA476804 |
| SRR7369627 | 2016 | Australia | Pig Unknown | Unknown | 25 | 2 | [33] | PRJNA476804 |
| SRR7369628 | 2016 | Australia | Pig Unknown | Unknown | 25 | 2 | [33] | PRJNA476804 |
| SRR7369629 | 2016 | Australia | Pig Unknown | Unknown | 25 | 2 | [33] | PRJNA476804 |
| SRR7369630 | 2016 | Australia | Pig Unknown | Unknown | 25 | 2 | [33] | PRJNA476804 |
| SRR7369631 | 2016 | Australia | Pig Unknown | Unknown | 483 | 23 | [33] | PRJNA476804 |
| SRR7369632 | 2016 | Australia | Pig Unknown | Unknown | 25 | 2 | [33] | PRJNA476804 |
| SRR7369633 | 2016 | Australia | Pig Unknown | Unknown | 27 | NA | [33] | PRJNA476804 |

| | | | | | | | | |
|---|---|---|---|---|---|---|---|---|
| SRR7369634 | 2017 | Australia | Pig Unknown | Unknown | 25 | 2 | [33] | PRJNA476804 |
| SRR7369635 | 2016 | Australia | Pig Unknown | Unknown | 117 | 3 | [33] | PRJNA476804 |
| SRR7369636 | 2013 | Australia | Pig Unknown | Unknown | 483 | 5 | [33] | PRJNA476804 |
| SRR7369637 | NA | Australia | Pig Unknown | Unknown | 25 | 2 | [33] | PRJNA476804 |
| SRR7369638 | 2016 | Australia | Pig Unknown | Unknown | 117 | 3 | [33] | PRJNA476804 |
| SRR7369639 | 2016 | Australia | Pig Unknown | Unknown | NA | 19 | [33] | PRJNA476804 |
| SRR7369640 | 2016 | Australia | Pig Unknown | Unknown | NA | 16 | [33] | PRJNA476804 |
| SRR7369641 | 2016 | Australia | Pig Unknown | Unknown | 27 | 3 | [33] | PRJNA476804 |
| SRR7369642 | 2016 | Australia | Pig Unknown | Unknown | 76 | 19 | [33] | PRJNA476804 |
| SRR7369643 | 2013 | Australia | Pig Unknown | Unknown | 87 | 8 | [33] | PRJNA476804 |
| SRR7369644 | 2014 | Australia | Pig Unknown | Unknown | 25 | 2 | [33] | PRJNA476804 |
| SRR7369645 | 2013 | Australia | Pig Unknown | Unknown | 27 | 3 | [33] | PRJNA476804 |
| SRR7369646 | 2013 | Australia | Pig Unknown | Unknown | NA | 10 | [33] | PRJNA476804 |
| SRR7369647 | 2017 | Australia | Pig Unknown | Unknown | 25 | 2 | [33] | PRJNA476804 |
| SRR7369648 | 2016 | Australia | Pig Unknown | Unknown | 27 | 3 | [33] | PRJNA476804 |
| SRR7369649 | 2017 | Australia | Pig Unknown | Unknown | 25 | 2 | [33] | PRJNA476804 |
| SRR7369650 | 2016 | Australia | Pig Unknown | Unknown | 25 | 2 | [33] | PRJNA476804 |
| SRR7369651 | 2016 | Australia | Pig Unknown | Unknown | 25 | 2 | [33] | PRJNA476804 |
| SRR7369652 | 2016 | Australia | Pig Unknown | Unknown | 27 | 3 | [33] | PRJNA476804 |
| SRR7369653 | 2016 | Australia | Pig Unknown | Unknown | 27 | 3 | [33] | PRJNA476804 |
| SRR7369654 | 2016 | Australia | Pig Unknown | Unknown | NA | 2 | [33] | PRJNA476804 |
| SRR7369655 | 2016 | Australia | Pig Unknown | Unknown | 27 | 3 | [33] | PRJNA476804 |
| SRR7369656 | 2016 | Australia | Pig Unknown | Unknown | 483 | 4 | [33] | PRJNA476804 |
| SRR7369657 | 2016 | Australia | Pig Unknown | Unknown | 483 | 4 | [33] | PRJNA476804 |
| SRR7369658 | 2016 | Australia | Pig Unknown | Unknown | 27 | 3 | [33] | PRJNA476804 |
| SRR7369659 | 2016 | Australia | Pig Unknown | Unknown | 25 | 2 | [33] | PRJNA476804 |

| | | | | | | | | |
|---|---|---|---|---|---|---|---|---|
| SRR7369660 | 2016 | Australia | Pig Unknown | Unknown | 76 | 19 | [33] | PRJNA476804 |
| SRR7369661 | 2013 | Australia | Pig Unknown | Unknown | NA | 16 | [33] | PRJNA476804 |
| SRR7369662 | 2013 | Australia | Pig Unknown | Unknown | NA | 31 | [33] | PRJNA476804 |
| SRR7369663 | 2013 | Australia | Pig Unknown | Unknown | 27 | 3 | [33] | PRJNA476804 |
| SRR7369664 | 2013 | Australia | Pig Unknown | Unknown | 87 | 8 | [33] | PRJNA476804 |
| SRR7369665 | 2013 | Australia | Pig Unknown | Unknown | 117 | 3 | [33] | PRJNA476804 |
| SRR7369666 | 2013 | Australia | Pig Unknown | Unknown | 483 | 23 | [33] | PRJNA476804 |
| SRR7369667 | 2013 | Australia | Pig Unknown | Unknown | 76 | 19 | [33] | PRJNA476804 |
| SRR7369668 | 2013 | Australia | Pig Unknown | Unknown | NA | 16 | [33] | PRJNA476804 |
| SRR7369669 | 2013 | Australia | Pig Unknown | Unknown | NA | 15 | [33] | PRJNA476804 |
| SRR7369670 | 2013 | Australia | Pig Unknown | Unknown | 117 | 3 | [33] | PRJNA476804 |
| SRR7369671 | 2016 | Australia | Pig Unknown | Unknown | NA | NA | [33] | PRJNA476804 |
| SRR7369673 | 2016 | Australia | Pig Unknown | Unknown | 27 | 3 | [33] | PRJNA476804 |
| SRR7369674 | 2016 | Australia | Pig Unknown | Unknown | NA | 19 | [33] | PRJNA476804 |
| SRR7369675 | 2016 | Australia | Pig Unknown | Unknown | 28 | 2 | [33] | PRJNA476804 |
| SRR7369676 | 2016 | Australia | Pig Unknown | Unknown | NA | 2 | [33] | PRJNA476804 |
| SRR7369677 | 2016 | Australia | Pig Unknown | Unknown | NA | NA | [33] | PRJNA476804 |
| SRR7369678 | 2016 | Australia | Pig Unknown | Unknown | NA | 3 | [33] | PRJNA476804 |
| SRR7369679 | 2016 | Australia | Pig Unknown | Unknown | NA | 19 | [33] | PRJNA476804 |
| SRR7369680 | 2016 | Australia | Pig Unknown | Unknown | NA | 18 | [33] | PRJNA476804 |
| SRR7369681 | 2011 | Australia | Pig Unknown | Unknown | 1 | 14 | [33] | PRJNA476804 |
| SRR7369682 | 2013 | Australia | Pig Unknown | Unknown | NA | 31 | [33] | PRJNA476804 |
| SRR7369683 | 2014 | Australia | Pig Unknown | Unknown | NA | 29 | [33] | PRJNA476804 |
| SRR7369684 | 2014 | Australia | Pig Unknown | Unknown | 87 | 8 | [33] | PRJNA476804 |
| SRR7369685 | 2014 | Australia | Pig Unknown | Unknown | 27 | 3 | [33] | PRJNA476804 |
| SRR7369686 | 2014 | Australia | Pig Unknown | Unknown | NA | 5 | [33] | PRJNA476804 |

| | | | | | | | | |
|---|---|---|---|---|---|---|---|---|
| SRR7369687 | 2014 | Australia | Pig Unknown | Unknown | 27 | 3 | [33] | PRJNA476804 |
| SRR7369688 | 2014 | Australia | Pig Unknown | Unknown | 27 | 3 | [33] | PRJNA476804 |
| SRR7369689 | 2013 | Australia | Pig Unknown | Unknown | 117 | 3 | [33] | PRJNA476804 |
| SRR7369690 | 2013 | Australia | Pig Unknown | Unknown | 483 | 4 | [33] | PRJNA476804 |
| SRR7369691 | 2013 | Australia | Pig Unknown | Unknown | 117 | 3 | [33] | PRJNA476804 |
| SRR7369692 | 2013 | Australia | Pig Unknown | Unknown | 27 | 3 | [33] | PRJNA476804 |
| SRR7369693 | 2013 | Australia | Pig Unknown | Unknown | 25 | 2 | [33] | PRJNA476804 |
| SRR7369695 | 2013 | Australia | Pig Unknown | Unknown | 27 | 3 | [33] | PRJNA476804 |
| SRR7369696 | 2013 | Australia | Pig Unknown | Unknown | 87 | 8 | [33] | PRJNA476804 |
| SRR7369697 | 2013 | Australia | Pig Unknown | Unknown | 1 | 2 | [33] | PRJNA476804 |
| SRR7369698 | 2013 | Australia | Pig Unknown | Unknown | NA | 12 | [33] | PRJNA476804 |
| SRR7369699 | 2015 | Australia | Pig Unknown | Unknown | 76 | 19 | [33] | PRJNA476804 |
| SRR7369700 | 2015 | Australia | Pig Unknown | Unknown | 76 | 19 | [33] | PRJNA476804 |
| SRR7369701 | 2015 | Australia | Pig Unknown | Unknown | 27 | 3 | [33] | PRJNA476804 |
| SRR7369702 | 2014 | Australia | Pig Unknown | Unknown | NA | 7 | [33] | PRJNA476804 |
| SRR7369703 | 2014 | Australia | Pig Unknown | Unknown | 483 | 4 | [33] | PRJNA476804 |
| SRR7369704 | 2014 | Australia | Pig Unknown | Unknown | 1 | 2 | [33] | PRJNA476804 |
| SRR7369705 | 2014 | Australia | Pig Unknown | Unknown | 1 | 2 | [33] | PRJNA476804 |
| SRR7369706 | 2014 | Australia | Pig Unknown | Unknown | 25 | 2 | [33] | PRJNA476804 |
| SRR7369707 | 2014 | Australia | Pig Unknown | Unknown | NA | 12 | [33] | PRJNA476804 |
| SRR7369709 | 2016 | Australia | Pig Unknown | Unknown | 28 | 2 | [33] | PRJNA476804 |
| SRR7369710 | 2016 | Australia | Pig Unknown | Unknown | 28 | 2 | [33] | PRJNA476804 |
| SRR7369711 | 2015 | Australia | Pig Unknown | Unknown | NA | 21 | [33] | PRJNA476804 |
| SRR7369712 | 2015 | Australia | Pig Unknown | Unknown | 27 | 3 | [33] | PRJNA476804 |
| SRR7369713 | 2016 | Australia | Pig Unknown | Unknown | NA | 2 | [33] | PRJNA476804 |
| SRR7369714 | 2016 | Australia | Pig Unknown | Unknown | NA | 3 | [33] | PRJNA476804 |

| | | | | | | | | |
|---|---|---|---|---|---|---|---|---|
| SRR7369715 | 2016 | Australia | Pig Unknown | Unknown | 27 | 3 | [33] | PRJNA476804 |
| SRR7369716 | 2016 | Australia | Pig Unknown | Unknown | 87 | 8 | [33] | PRJNA476804 |
| SRR7369717 | 2016 | Australia | Pig Unknown | Unknown | 27 | 3 | [33] | PRJNA476804 |
| SRR7369718 | 2016 | Australia | Pig Unknown | Unknown | 28 | 2 | [33] | PRJNA476804 |
| 1093404 | 2008 | Canada | Pig Diseased | Systemic | NA | 27 | [20] | PRJNA628943 |
| 1112943 | 2008 | Canada | Pig Unknown | Unknown | NA | NA | [20] | PRJNA628943 |
| 1126699 | 2008 | Canada | Pig Unknown | Unknown | NA | 17 | [20] | PRJNA628943 |
| 1129711 | 2008 | Canada | Pig Diseased | Systemic | NA | 29 | [20] | PRJNA628943 |
| 1148794 | 2009 | Canada | Pig Diseased | Systemic | NA | NA | [20] | PRJNA628943 |
| 1152130 | 2009 | Canada | Pig Unknown | Unknown | NA | 21 | [20] | PRJNA628943 |
| 1158584 | 2009 | Canada | Pig Unknown | Unknown | NA | 31 | [20] | PRJNA628943 |
| 1160410 | 2009 | Canada | Pig Unknown | Unknown | NA | NA | [20] | PRJNA628943 |
| 1162351 | 2009 | Canada | Pig Diseased | Respiratory | NA | 24 | [20] | PRJNA628943 |
| 1171239 | 2009 | Canada | Pig Unknown | Unknown | NA | NA | [20] | PRJNA628943 |
| 1193601 | 2009 | Canada | Pig Diseased | Systemic | NA | NA | [20] | PRJNA628943 |
| 1197165 | 2009 | Canada | Pig Diseased | Systemic | NA | 9 | [20] | PRJNA628943 |
| 1197378 | 2009 | Canada | Pig Unknown | Unknown | NA | 9 | [20] | PRJNA628943 |
| 1209618 | 2010 | Canada | Pig Unknown | Unknown | NA | 9 | [20] | PRJNA628943 |
| 1231073 | 2010 | Canada | Pig Diseased | Systemic | NA | 31 | [20] | PRJNA628943 |
| 1231255 | 2010 | Canada | Pig Diseased | Respiratory | 76 | 19 | [20] | PRJNA628943 |
| 1248867 | 2010 | Canada | Pig Diseased | Respiratory | NA | NA | [20] | PRJNA628943 |
| 1269330 | 2011 | Canada | Pig Diseased | Systemic | NA | 21 | [20] | PRJNA628943 |
| 1270935 | 2011 | Canada | Pig Unknown | Unknown | NA | 28 | [20] | PRJNA628943 |
| 1348170 | 2011 | Canada | Pig Unknown | Unknown | NA | 13 | [20] | PRJNA628943 |
| 1378494 | 2012 | Canada | Pig Unknown | Unknown | NA | 25 | [20] | PRJNA628943 |
| 1442765 | 2012 | Canada | Pig Unknown | Unknown | NA | 16 | [20] | PRJNA628943 |

| | | | | | | | | |
|---|---|---|---|---|---|---|---|---|
| 1456898 | 2013 | Canada | Pig Unknown | Unknown | NA | NA | [20] | PRJNA628943 |
| 1462648 | 2013 | Canada | Pig Unknown | Unknown | 1 | 14 | [20] | PRJNA628943 |
| 1463646 | 2013 | Canada | Pig Diseased | Systemic | 1 | 14 | [20] | PRJNA628943 |
| 1464747 | 2013 | Canada | Pig Unknown | Unknown | 87 | 8 | [20] | PRJNA628943 |
| 1464749 | 2013 | Canada | Pig Unknown | Unknown | 1 | 14 | [20] | PRJNA628943 |
| 1465473 | 2013 | Canada | Pig Unknown | Unknown | 1 | 14 | [20] | PRJNA628943 |
| 1466571 | 2013 | Canada | Pig Diseased | Systemic | NA | NA | [20] | PRJNA628943 |
| 1475186 | 2013 | Canada | Pig Unknown | Unknown | 27 | 3 | [20] | PRJNA628943 |
| 1477528 | 2013 | Canada | Pig Unknown | Unknown | 27 | 3 | [20] | PRJNA628943 |
| 1478991 | 2013 | Canada | Pig Diseased | Respiratory | 87 | 8 | [20] | PRJNA628943 |
| 1481256 | 2013 | Canada | Pig Unknown | Unknown | 87 | 8 | [20] | PRJNA628943 |
| 1481257 | 2013 | Canada | Pig Unknown | Unknown | 87 | 8 | [20] | PRJNA628943 |
| 1481965 | 2013 | Canada | Pig Diseased | Systemic | NA | NA | [20] | PRJNA628943 |
| 1491188 | 2013 | Canada | Pig Diseased | Respiratory | NA | 23 | [20] | PRJNA628943 |
| 1493634 | 2013 | Canada | Pig Unknown | Unknown | NA | NA | [20] | PRJNA628943 |
| 1505307 | 2013 | Canada | Pig Unknown | Unknown | NA | 16 | [20] | PRJNA628943 |
| 1506978 | 2013 | Canada | Pig Diseased | Systemic | 32 | 7 | [20] | PRJNA628943 |
| 1506989 | 2013 | Canada | Pig Unknown | Unknown | NA | NA | [20] | PRJNA628943 |
| 1508445 | 2013 | Canada | Pig Diseased | Systemic | 108 | 23 | [20] | PRJNA628943 |
| 1514647 | 2013 | Canada | Pig Diseased | Respiratory | NA | 28 | [20] | PRJNA628943 |
| 1521251 | 2013 | Canada | Pig Diseased | Systemic | NA | NA | [20] | PRJNA628943 |
| 1588531 | 2014 | Canada | Pig Unknown | Unknown | 1 | 14 | [20] | PRJNA628943 |
| 1596165 | 2014 | Canada | Pig Diseased | Systemic | 28 | 14 | [20] | PRJNA628943 |
| 1596201 | 2014 | Canada | Pig Diseased | Systemic | 1 | 14 | [20] | PRJNA628943 |
| 1596217 | 2014 | Canada | Pig Unknown | Unknown | 1 | 14 | [20] | PRJNA628943 |
| 1597776 | 2014 | Canada | Pig Unknown | Unknown | 28 | 2 | [20] | PRJNA628943 |

| | | | | | | | | |
|---|---|---|---|---|---|---|---|---|
| 1597778 | 2014 | Canada | Pig Unknown | Unknown | 28 | 2 | [20] | PRJNA628943 |
| 1600268 | 2014 | Canada | Pig Unknown | Unknown | 94 | NA | [20] | PRJNA628943 |
| 1601399 | 2014 | Canada | Pig Unknown | Unknown | 28 | 2 | [20] | PRJNA628943 |
| 1601827 | 2014 | Canada | Pig Unknown | Unknown | 94 | NA | [20] | PRJNA628943 |
| 1602949 | 2014 | Canada | Pig Unknown | Unknown | 13 | 14 | [20] | PRJNA628943 |
| 1602951 | 2014 | Canada | Pig Diseased | Respiratory | 25 | 2 | [20] | PRJNA628943 |
| 1602952 | 2014 | Canada | Pig Diseased | Respiratory | NA | NA | [20] | PRJNA628943 |
| 1602954 | 2014 | Canada | Pig Diseased | Respiratory | 87 | 8 | [20] | PRJNA628943 |
| 1602956 | 2014 | Canada | Pig Unknown | Unknown | 94 | 4 | [20] | PRJNA628943 |
| 1603813 | 2014 | Canada | Pig Unknown | Unknown | 28 | 2 | [20] | PRJNA628943 |
| 1606517 | 2014 | Canada | Pig Unknown | Unknown | NA | 3 | [20] | PRJNA628943 |
| 1606529 | 2014 | Canada | Pig Unknown | Unknown | 28 | 2 | [20] | PRJNA628943 |
| 1607743 | 2014 | Canada | Pig Unknown | Unknown | NA | 4 | [20] | PRJNA628943 |
| 1607744 | 2014 | Canada | Pig Unknown | Unknown | 89 | NA | [20] | PRJNA628943 |
| 1610743 | 2014 | Canada | Pig Unknown | Unknown | NA | 3 | [20] | PRJNA628943 |
| 1611505 | 2014 | Canada | Pig Unknown | Unknown | 1 | 14 | [20] | PRJNA628943 |
| 1614720 | 2014 | Canada | Pig Unknown | Unknown | NA | 16 | [20] | PRJNA628943 |
| 1615460 | 2014 | Canada | Pig Unknown | Unknown | 225 | 7 | [20] | PRJNA628943 |
| 1615461 | 2014 | Canada | Pig Diseased | Systemic | NA | 9 | [20] | PRJNA628943 |
| 1615892 | 2014 | Canada | Pig Unknown | Unknown | NA | 9 | [20] | PRJNA628943 |
| 1621705 | 2014 | Canada | Pig Unknown | Unknown | NA | NA | [20] | PRJNA628943 |
| 1622145 | 2014 | Canada | Pig Diseased | Respiratory | NA | 28 | [20] | PRJNA628943 |
| 1628823 | 2014 | Canada | Pig Diseased | Systemic | 87 | 8 | [20] | PRJNA628943 |
| 1633180 | 2014 | Canada | Pig Diseased | Systemic | 108 | 23 | [20] | PRJNA628943 |
| 1636820 | 2014 | Canada | Pig Unknown | Unknown | 25 | 2 | [20] | PRJNA628943 |
| 1637946 | 2014 | Canada | Pig Unknown | Unknown | 94 | NA | [20] | PRJNA628943 |

| | | | | | | | | |
|---|---|---|---|---|---|---|---|---|
| 1644048 | 2014 | Canada | Pig Unknown | Unknown | 1 | 14 | [20] | PRJNA628943 |
| 1650131 | 2014 | Canada | Pig Unknown | Unknown | 28 | 2 | [20] | PRJNA628943 |
| 1650132 | 2014 | Canada | Pig Unknown | Unknown | 28 | 2 | [20] | PRJNA628943 |
| 1651929 | 2014 | Canada | Pig Diseased | Respiratory | 28 | 2 | [20] | PRJNA628943 |
| 1652716 | 2014 | Canada | Pig Diseased | Systemic | NA | 4 | [20] | PRJNA628943 |
| 1654537 | 2014 | Canada | Pig Diseased | Respiratory | 28 | 2 | [20] | PRJNA628943 |
| 1656890 | 2014 | Canada | Pig Unknown | Unknown | 108 | 3 | [20] | PRJNA628943 |
| 1659834 | 2014 | Canada | Pig Unknown | Unknown | 1 | 14 | [20] | PRJNA628943 |
| 1661949 | 2014 | Canada | Pig Unknown | Unknown | 28 | 2 | [20] | PRJNA628943 |
| 1665814 | 2014 | Canada | Pig Unknown | Unknown | NA | 4 | [20] | PRJNA628943 |
| 1666725 | 2014 | Canada | Pig Unknown | Unknown | 25 | 2 | [20] | PRJNA628943 |
| 1666727 | 2014 | Canada | Pig Unknown | Unknown | 1 | 14 | [20] | PRJNA628943 |
| 1667796 | 2014 | Canada | Pig Diseased | Systemic | NA | 2 | [20] | PRJNA628943 |
| 1671154 | 2014 | Canada | Pig Unknown | Unknown | NA | 27 | [20] | PRJNA628943 |
| 1801887 | 2016 | Canada | Pig Unknown | Unknown | 28 | 2 | [20] | PRJNA628943 |
| 1803662 | 2016 | Canada | Pig Unknown | Unknown | 622 | 9 | [20] | PRJNA628943 |
| 00-3638-4B | 2000 | Canada | Pig Diseased | Systemic | NA | NA | [20] | PRJNA628943 |
| 11.CEREBRO | 2016 | Spain | Pig Diseased | Systemic | NA | 9 | [20] | PRJNA628943 |
| 1191316_ref | 2009 | Canada | Pig Diseased | Systemic | NA | 28 | [20] | PRJNA628943 |
| 12.CEREBRO | 2016 | Spain | Pig Diseased | Systemic | NA | 9 | [20] | PRJNA628943 |
| 13.CEREBRO | 2016 | Spain | Pig Diseased | Systemic | NA | 9 | [20] | PRJNA628943 |
| 1522228_ref | 2013 | Canada | Pig Diseased | Systemic | 483 | 23 | [20] | PRJNA1010137 |
| 1619952_ref | 2014 | Canada | Pig Diseased | Systemic | 1 | 14 | [20] | PRJNA628943 |
| 3.ARTICULATION | 2016 | Spain | Pig Diseased | Systemic | 1 | 14 | [20] | PRJNA628943 |
| 3.CEREBRO | 2016 | Spain | Pig Diseased | Systemic | 1 | 14 | [20] | PRJNA628943 |
| 4.2CEREBRO | 2016 | Spain | Pig Diseased | Systemic | 1 | 14 | [20] | PRJNA628943 |

| | | | | | | | | |
|---|---|---|---|---|---|---|---|---|
| 4.ARTICULATION | 2016 | Spain | Pig Diseased | Systemic | 1 | 14 | [20] | PRJNA628943 |
| 87-4647 | 1987 | Canada | Pig Unknown | Unknown | 225 | 7 | [20] | PRJNA628943 |
| 88-2775 | 1988 | Canada | Pig Diseased | Respiratory | NA | 13 | [20] | PRJNA628943 |
| 88-5454A | 1988 | Canada | Pig Diseased | Respiratory | 117 | 3 | [20] | PRJNA628943 |
| 89-1591 | 1989 | Canada | Pig Unknown | Unknown | 25 | 2 | [20] | PRJNA628943 |
| 89-1824 | 1989 | Canada | Pig Diseased | Systemic | 76 | 19 | [20] | PRJNA628943 |
| 89-289 | 1989 | Canada | Pig Unknown | Unknown | NA | 9 | [20] | PRJNA628943 |
| 89-4004 | 1989 | Canada | Pig Diseased | Respiratory | 28 | 2 | [20] | PRJNA628943 |
| 89-4223 | 1989 | Canada | Pig Diseased | Respiratory | 28 | 2 | [20] | PRJNA628943 |
| 89-4294_ref | 1989 | Canada | Pig Diseased | Respiratory | NA | 19 | [20] | PRJNA628943 |
| 89-5046 | 1989 | Canada | Pig Diseased | Systemic | 25 | 2 | [20] | PRJNA628943 |
| 89-5189 | 1989 | Canada | Pig Unknown | Unknown | 225 | 7 | [20] | PRJNA628943 |
| 89-5197 | 1989 | Canada | Pig Diseased | Respiratory | NA | 16 | [20] | PRJNA628943 |
| 89-6891-2 | 1989 | Canada | Pig Diseased | Respiratory | 25 | 2 | [20] | PRJNA628943 |
| 89-88-422 | 1989 | Canada | Pig Diseased | Respiratory | NA | 31 | [20] | PRJNA628943 |
| 90-1145 | 1990 | Canada | Pig Unknown | Unknown | NA | 31 | [20] | PRJNA628943 |
| 90-2741-7 | 1990 | Canada | Pig Diseased | Systemic | 25 | 2 | [20] | PRJNA628943 |
| 90-3615-2 | 1990 | Canada | Pig Unknown | Unknown | NA | 11 | [20] | PRJNA628943 |
| 90-4400 | 1990 | Canada | Pig Diseased | Respiratory | NA | NA | [20] | PRJNA628943 |
| 90-4517-4 | 1990 | Canada | Pig Diseased | Respiratory | NA | 11 | [20] | PRJNA628943 |
| 90-4582 | 1990 | Canada | Pig Diseased | Respiratory | NA | 19 | [20] | PRJNA628943 |
| 90-546 | 1990 | Canada | Pig Diseased | Respiratory | NA | NA | [20] | PRJNA628943 |
| 91-1174 | 1991 | Canada | Pig Diseased | Respiratory | 13 | 14 | [20] | PRJNA628943 |
| 91-173B-752 | 1991 | Canada | Pig Diseased | Systemic | 13 | 14 | [20] | PRJNA628943 |
| 91-175B-2156 | 1991 | Canada | Pig Diseased | Respiratory | NA | 2 | [20] | PRJNA628943 |
| 91-178-2215 | 1991 | Canada | Pig Unknown | Unknown | NA | 4 | [20] | PRJNA628943 |

| | | | | | | | | |
|---|---|---|---|---|---|---|---|---|
| 91-178-2469 | 1991 | Canada | Pig Diseased | Respiratory | 117 | 3 | [20] | PRJNA628943 |
| 92-1130 | 1992 | Canada | Pig Diseased | Systemic | 13 | 14 | [20] | PRJNA628943 |
| 92-2402-1119 | 1992 | Canada | Pig Diseased | Respiratory | NA | NA | [20] | PRJNA628943 |
| 93-1320 | 1993 | Canada | Pig Unknown | Unknown | NA | NA | [20] | PRJNA628943 |
| 93-2042-4514 | 1993 | Canada | Pig Diseased | Respiratory | NA | NA | [20] | PRJNA628943 |
| 94-1713 | 1994 | Canada | Pig Unknown | Unknown | NA | 11 | [20] | PRJNA628943 |
| 94-2571 | 1994 | Canada | Pig Diseased | Systemic | NA | 12 | [20] | PRJNA628943 |
| 94-3370-6480 | 1994 | Canada | Pig Unknown | Unknown | NA | 15 | [20] | PRJNA628943 |
| 94-8576-4887 | 1994 | Canada | Pig Diseased | Respiratory | 28 | 14 | [20] | PRJNA628943 |
| 99-C561-8614-2 | 1999 | Canada | Pig Diseased | Systemic | 483 | 23 | [20] | PRJNA628943 |
| BE4B3-2A | 2016 | Canada | Pig Healthy | Non-Clinical | 108 | 23 | [20] | PRJNA628943 |
| BE4B3-4A | 2016 | Canada | Pig Healthy | Non-Clinical | NA | 16 | [20] | PRJNA628943 |
| BE4B3-4B | 2016 | Canada | Pig Healthy | Non-Clinical | NA | 16 | [20] | PRJNA628943 |
| CF2D3-1C | 2016 | Canada | Pig Healthy | Non-Clinical | NA | 28 | [20] | PRJNA628943 |
| CF2D3-2C | 2016 | Canada | Pig Healthy | Non-Clinical | NA | 28 | [20] | PRJNA628943 |
| CF2D3-2D | 2016 | Canada | Pig Healthy | Non-Clinical | NA | NA | [20] | PRJNA628943 |
| CF2D3-2E | 2016 | Canada | Pig Healthy | Non-Clinical | NA | NA | [20] | PRJNA628943 |
| CF2D3-4A | 2016 | Canada | Pig Healthy | Non-Clinical | NA | 5 | [20] | PRJNA628943 |
| CF2D3-4B | 2016 | Canada | Pig Healthy | Non-Clinical | NA | NA | [20] | PRJNA628943 |
| CF2D3-4C | 2016 | Canada | Pig Healthy | Non-Clinical | NA | NA | [20] | PRJNA628943 |
| CF2D3-4D | 2016 | Canada | Pig Healthy | Non-Clinical | NA | NA | [20] | PRJNA628943 |
| CG3S3-2A | 2016 | Canada | Pig Healthy | Non-Clinical | NA | 29 | [20] | PRJNA628943 |
| CG3S3-2B | 2016 | Canada | Pig Healthy | Non-Clinical | NA | 29 | [20] | PRJNA628943 |
| D15-036307 | NA | USA | Pig Unknown | Unknown | 28 | 2 | [20] | PRJNA628943 |
| D15-036308 | NA | USA | Pig Diseased | Systemic | 108 | 23 | [20] | PRJNA628943 |
| D15-036753 | NA | USA | Pig Unknown | Unknown | NA | 3 | [20] | PRJNA628943 |

| | | | | | | | | |
|---|---|---|---|---|---|---|---|---|
| D15-036755 | NA | USA | Pig Unknown | Unknown | 28 | NA | [20] | PRJNA628943 |
| D15-036840 | NA | USA | Pig Unknown | Unknown | NA | 9 | [20] | PRJNA628943 |
| D15-037100 | NA | USA | Pig Diseased | Systemic | 108 | 23 | [20] | PRJNA628943 |
| D15-037104 | NA | USA | Pig Unknown | Unknown | 28 | 2 | [20] | PRJNA628943 |
| D15-049776 | NA | USA | Pig Diseased | Respiratory | NA | 9 | [20] | PRJNA628943 |
| D15-049887 | NA | USA | Pig Unknown | Unknown | 28 | 2 | [20] | PRJNA628943 |
| D15-055799 | NA | USA | Pig Diseased | Respiratory | 28 | 2 | [20] | PRJNA628943 |
| D15-061569 | NA | USA | Pig Unknown | Unknown | NA | 31 | [20] | PRJNA628943 |
| D16-009258 | NA | USA | Pig Diseased | Respiratory | NA | NA | [20] | PRJNA628943 |
| D16-010263 | NA | USA | Pig Diseased | Respiratory | 28 | 2 | [20] | PRJNA628943 |
| D16-010378 | NA | USA | Pig Unknown | Unknown | NA | 4 | [20] | PRJNA628943 |
| D16-017871 | NA | USA | Pig Diseased | Systemic | NA | 9 | [20] | PRJNA628943 |
| DB1V3-4A_ref | 2016 | Canada | Pig Healthy | Non-Clinical | NA | 4 | [20] | PRJNA628943 |
| EJ2T3-1A | 2016 | Canada | Pig Healthy | Non-Clinical | NA | NA | [20] | PRJNA628943 |
| EJ2T3-1B | 2016 | Canada | Pig Healthy | Non-Clinical | NA | NA | [20] | PRJNA628943 |
| EJ2T3-2A | 2016 | Canada | Pig Healthy | Non-Clinical | 89 | NA | [20] | PRJNA628943 |
| EJ2T3-2B | 2016 | Canada | Pig Healthy | Non-Clinical | NA | NA | [20] | PRJNA628943 |
| EJ2T3-2C | 2016 | Canada | Pig Healthy | Non-Clinical | 89 | NA | [20] | PRJNA628943 |
| EJ2T3-2D | 2016 | Canada | Pig Healthy | Non-Clinical | 89 | NA | [20] | PRJNA628943 |
| EJ2T3-2E | 2016 | Canada | Pig Healthy | Non-Clinical | 89 | NA | [20] | PRJNA628943 |
| EJ2T3-3B | 2016 | Canada | Pig Healthy | Non-Clinical | NA | 15 | [20] | PRJNA628943 |
| MA4T3-1A | 2016 | Canada | Pig Healthy | Non-Clinical | NA | NA | [20] | PRJNA628943 |
| MA4T3-1B | 2016 | Canada | Pig Healthy | Non-Clinical | NA | 31 | [20] | PRJNA628943 |
| MA4T3-2B | 2016 | Canada | Pig Healthy | Non-Clinical | 87 | 8 | [20] | PRJNA628943 |
| MA4T3-3A | 2016 | Canada | Pig Healthy | Non-Clinical | 28 | 2 | [20] | PRJNA628943 |
| MA4T3-3B | 2016 | Canada | Pig Healthy | Non-Clinical | 28 | 2 | [20] | PRJNA628943 |

| | | | | | | | | |
|---|---|---|---|---|---|---|---|---|
| MA4T3-3C | 2016 | Canada | Pig Healthy | Non-Clinical | NA | 31 | [20] | PRJNA628943 |
| MA4T3-4A | 2016 | Canada | Pig Healthy | Non-Clinical | NA | 5 | [20] | PRJNA628943 |
| MA4T3-4B | 2016 | Canada | Pig Healthy | Non-Clinical | NA | NA | [20] | PRJNA628943 |
| MA4T3-4C | 2016 | Canada | Pig Healthy | Non-Clinical | NA | 4 | [20] | PRJNA628943 |
| MA4T3-4D | 2016 | Canada | Pig Healthy | Non-Clinical | NA | NA | [20] | PRJNA628943 |
| ME1D3-3A | 2016 | Canada | Pig Healthy | Non-Clinical | NA | 31 | [20] | PRJNA628943 |
| MF1S3-4a_LW | NA | Canada | NA | Unknown | NA | 2 | [20] | PRJNA628943 |
| MY1C3_3B_ref | 2016 | Canada | Pig Healthy | Non-Clinical | NA | 3 | [20] | PRJNA628943 |
| MY1C3-1A | 2016 | Canada | Pig Healthy | Non-Clinical | 28 | 2 | [20] | PRJNA628943 |
| MY1C3-1B | 2016 | Canada | Pig Healthy | Non-Clinical | 28 | 2 | [20] | PRJNA628943 |
| MY1C3-2B | 2016 | Canada | Pig Healthy | Non-Clinical | NA | 3 | [20] | PRJNA628943 |
| MY1C3-2C | 2016 | Canada | Pig Healthy | Non-Clinical | NA | 3 | [20] | PRJNA628943 |
| MY1C3-3C | 2016 | Canada | Pig Healthy | Non-Clinical | NA | 4 | [20] | PRJNA628943 |
| MZ1B3_4E_ref | 2016 | Canada | Pig Healthy | Non-Clinical | 28 | 14 | [20] | PRJNA763404 |
| MZ1B3-4A | 2016 | Canada | Pig Healthy | Non-Clinical | 28 | 14 | [20] | PRJNA628943 |
| MZ1B3-4B | 2016 | Canada | Pig Healthy | Non-Clinical | 28 | 14 | [20] | PRJNA628943 |
| MZ1B3-4C | 2016 | Canada | Pig Healthy | Non-Clinical | 28 | 14 | [20] | PRJNA628943 |
| MZ1B3-4D | 2016 | Canada | Pig Healthy | Non-Clinical | 28 | 14 | [20] | PRJNA628943 |
| NH1E3-10A | 2016 | Canada | Pig Healthy | Non-Clinical | NA | NA | [20] | PRJNA628943 |
| NH1E3-10B | 2016 | Canada | Pig Healthy | Non-Clinical | NA | NA | [20] | PRJNA628943 |
| NH1E3-9A | 2016 | Canada | Pig Healthy | Non-Clinical | NA | NA | [20] | PRJNA628943 |
| NH1E3-9C | 2016 | Canada | Pig Healthy | Non-Clinical | NA | NA | [20] | PRJNA628943 |
| NH1E3-9D | 2016 | Canada | Pig Healthy | Non-Clinical | NA | NA | [20] | PRJNA628943 |
| P83-3336 | 1983 | Canada | Pig Diseased | Systemic | NA | NA | [20] | PRJNA628943 |
| TWLWHOG | NA | UK | NA | Unknown | 1 | 2 | [20] | PRJNA628943 |
| VALENCIA_ST2 | 2016 | Spain | Pig Unknown | Unknown | 29 | NA | [20] | PRJNA628943 |

| | | | | | | | | |
|---|---|---|---|---|---|---|---|---|
| VALENCIA_ST4 | 2016 | Spain | Pig Unknown | Unknown | NA | NA | [20] | PRJNA628943 |
| VALENCIA_ST6 | 2016 | Spain | Pig Unknown | Unknown | 29 | NA | [20] | PRJNA628943 |
| 01ZN-ST222 | 2010 | Vietnam | Pig Unknown | Unknown | 1 | 2 | [34] | PRJEB2670 |
| 01ZN-TG183 | 2010 | Vietnam | Pig Unknown | Unknown | 1 | 2 | [34] | PRJEB2670 |
| 01ZN-TG187 | NA | Vietnam | Pig Unknown | Unknown | 1 | 2 | [34] | PRJEB2670 |
| 01ZN-TG189 | 2010 | Vietnam | Pig Unknown | Unknown | 1 | 2 | [34] | PRJEB2670 |
| BM178a | 2000 | Vietnam | Human | Systemic | 1 | 2 | [34] | PRJEB2670 |
| BM190a | 2000 | Vietnam | Human | Unknown | 1 | 2 | [34] | PRJEB2670 |
| BM191a | 2000 | Vietnam | Human | Systemic | 1 | 2 | [34] | PRJEB2670 |
| BM197a | 2000 | Vietnam | Human | Systemic | 1 | 2 | [34] | PRJEB2670 |
| BM198a | 2000 | Vietnam | Human | Systemic | 1 | 2 | [34] | PRJEB2670 |
| BM203a | 2000 | Vietnam | Human | Systemic | 1 | 2 | [34] | PRJEB2670 |
| BM209a | 2000 | Vietnam | Human | Systemic | 1 | 2 | [34] | PRJEB2670 |
| BM211a | 2000 | Vietnam | Human | Systemic | 1 | 2 | [34] | PRJEB2670 |
| BM213a | 2000 | Vietnam | Human | Systemic | 1 | 2 | [34] | PRJEB2670 |
| BM216a | 2001 | Vietnam | Human | Systemic | 1 | 2 | [34] | PRJEB2670 |
| BM224C | 2001 | Vietnam | Human | Systemic | 1 | 2 | [34] | PRJEB2670 |
| BM228C | 2001 | Vietnam | Human | Systemic | 1 | 2 | [34] | PRJEB2670 |
| BM231a | 2001 | Vietnam | Human | Systemic | 1 | 2 | [34] | PRJEB2670 |
| BM237a | 2001 | Vietnam | Human | Systemic | 1 | 2 | [34] | PRJEB2670 |
| BM241a | 2001 | Vietnam | Human | Systemic | 1 | 2 | [34] | PRJEB2670 |
| BM251C | 2001 | Vietnam | Human | Systemic | 1 | 2 | [34] | PRJEB2670 |
| BM253a | 2001 | Vietnam | Human | Systemic | 1 | 2 | [34] | PRJEB2670 |
| BM257a | 2001 | Vietnam | Human | Systemic | 1 | 2 | [34] | PRJEB2670 |
| BM264a | 2002 | Vietnam | Human | Systemic | 1 | 2 | [34] | PRJEB2670 |
| BM272C | 2002 | Vietnam | Human | Systemic | 1 | 2 | [34] | PRJEB2670 |

| | | | | | | | | |
|---|---|---|---|---|---|---|---|---|
| BM278a | 2002 | Vietnam | Human | Systemic | 1 | 2 | [34] | PRJEB2670 |
| BM303a | 2002 | Vietnam | Human | Systemic | 1 | 2 | [34] | PRJEB2670 |
| BM308a | 2002 | Vietnam | Human | Systemic | 1 | 2 | [34] | PRJEB2670 |
| BM315C | 2002 | Vietnam | Human | Systemic | 1 | 2 | [34] | PRJEB2670 |
| BM321C | 2002 | Vietnam | Human | Systemic | 1 | 2 | [34] | PRJEB2670 |
| BM329B | 2003 | Vietnam | Human | Systemic | 1 | 2 | [34] | PRJEB2670 |
| BM334B | 2003 | Vietnam | Human | Systemic | 1 | 2 | [34] | PRJEB2670 |
| BM345B | 2003 | Vietnam | Human | Systemic | 1 | 2 | [34] | PRJEB2670 |
| BM346B | 2003 | Vietnam | Human | Systemic | 1 | 2 | [34] | PRJEB2670 |
| BM357B | 2003 | Vietnam | Human | Systemic | 1 | 2 | [34] | PRJEB2670 |
| BM358a | 2003 | Vietnam | Human | Systemic | 1 | 2 | [34] | PRJEB2670 |
| BM362a | 2003 | Vietnam | Human | Systemic | 1 | 2 | [34] | PRJEB2670 |
| BM366a | 2003 | Vietnam | Human | Systemic | 1 | 2 | [34] | PRJEB2670 |
| BM373a | 2003 | Vietnam | Human | Systemic | 1 | 2 | [34] | PRJEB2670 |
| BM411a | 2004 | Vietnam | Human | Systemic | 1 | 2 | [34] | PRJEB2670 |
| BM418a | 2004 | Vietnam | Human | Systemic | 1 | 2 | [34] | PRJEB2670 |
| BM424a | 2004 | Vietnam | Human | Systemic | 1 | 2 | [34] | PRJEB2670 |
| BM436a | 2004 | Vietnam | Human | Systemic | 1 | 2 | [34] | PRJEB2670 |
| BM441a | 2004 | Vietnam | Human | Systemic | 1 | 2 | [34] | PRJEB2670 |
| BM444a | 2004 | Vietnam | Human | Systemic | 1 | 2 | [34] | PRJEB2670 |
| BM452C | 2004 | Vietnam | Human | Systemic | 1 | 2 | [34] | PRJEB2670 |
| BM460 | 2005 | Vietnam | Human | Systemic | 1 | 2 | [34] | PRJEB2670 |
| BM461 | 2005 | Vietnam | Human | Systemic | 1 | 2 | [34] | PRJEB2670 |
| BM465 | 2005 | Vietnam | Human | Systemic | 1 | 2 | [34] | PRJEB2670 |
| BM466 | 2005 | Vietnam | Human | Systemic | 1 | 2 | [34] | PRJEB2670 |
| BM471 | 2005 | Vietnam | Human | Systemic | 1 | 2 | [34] | PRJEB2670 |

| | | | | | | | | |
|---|---|---|---|---|---|---|---|---|
| BM478 | 2005 | Vietnam | Human | Systemic | 1 | 2 | [34] | PRJEB2670 |
| CTU02SS-DN025 | 2009 | Vietnam | Pig Unknown | Unknown | 1 | 2 | [34] | PRJEB2670 |
| CTU02SS-DN026 | 2009 | Vietnam | Pig Unknown | Unknown | 1 | 2 | [34] | PRJEB2670 |
| EN003 | 2006 | Vietnam | Human | Systemic | 1 | 2 | [34] | PRJEB2670 |
| EN010 | 2006 | Vietnam | Human | Systemic | 1 | 2 | [34] | PRJEB2670 |
| EN0114 | 2006 | Vietnam | Human | Systemic | 1 | 2 | [34] | PRJEB2670 |
| EN015 | 2006 | Vietnam | Human | Systemic | 1 | 2 | [34] | PRJEB2670 |
| EN020 | 2006 | Vietnam | Human | Systemic | 1 | 2 | [34] | PRJEB2670 |
| EN023 | 2006 | Vietnam | Human | Systemic | 1 | 2 | [34] | PRJEB2670 |
| EN024 | 2006 | Vietnam | Human | Systemic | 1 | 2 | [34] | PRJEB2670 |
| EN031 | 2006 | Vietnam | Human | Systemic | 1 | 2 | [34] | PRJEB2670 |
| EN044 | 2006 | Vietnam | Human | Systemic | 1 | 2 | [34] | PRJEB2670 |
| EN050 | 2006 | Vietnam | Human | Systemic | 1 | 2 | [34] | PRJEB2670 |
| EN068 | 2006 | Vietnam | Human | Systemic | 1 | 2 | [34] | PRJEB2670 |
| EN124 | 2006 | Vietnam | Human | Systemic | 1 | 2 | [34] | PRJEB2670 |
| EN170 | 2006 | Vietnam | Human | Systemic | 1 | 2 | [34] | PRJEB2670 |
| EN171 | 2006 | Vietnam | Human | Systemic | 1 | 2 | [34] | PRJEB2670 |
| EN177 | 2006 | Vietnam | Human | Systemic | 1 | 2 | [34] | PRJEB2670 |
| EN191 | 2006 | Vietnam | Human | Systemic | 105 | 14 | [34] | PRJEB2670 |
| EN251 | 2007 | Vietnam | Human | Systemic | 1 | 2 | [34] | PRJEB2670 |
| EN253 | 2007 | Vietnam | Human | Systemic | 1 | 2 | [34] | PRJEB2670 |
| EN264 | 2007 | Vietnam | Human | Systemic | 1 | 2 | [34] | PRJEB2670 |
| EN271 | 2007 | Vietnam | Human | Systemic | 1 | 2 | [34] | PRJEB2670 |
| EN281 | 2007 | Vietnam | Human | Systemic | 1 | 2 | [34] | PRJEB2670 |
| EN314 | 2007 | Vietnam | Human | Systemic | 105 | 14 | [34] | PRJEB2670 |
| EN326 | 2007 | Vietnam | Human | Systemic | 1 | 2 | [34] | PRJEB2670 |

| | | | | | | | | |
|---|---|---|---|---|---|---|---|---|
| EN330 | 2007 | Vietnam | Human | Systemic | 1 | 2 | [34] | PRJEB2670 |
| EN370 | 2007 | Vietnam | Human | Systemic | 1 | 2 | [34] | PRJEB2670 |
| EN399 | 2007 | Vietnam | Human | Systemic | 1 | 2 | [34] | PRJEB2670 |
| EN506 | 2008 | Vietnam | Human | Systemic | 1 | 2 | [34] | PRJEB2670 |
| EN507 | 2008 | Vietnam | Human | Systemic | 1 | 2 | [34] | PRJEB2670 |
| EN508 | 2008 | Vietnam | Human | Systemic | 1 | 2 | [34] | PRJEB2670 |
| EN515 | 2008 | Vietnam | Human | Systemic | 1 | 2 | [34] | PRJEB2670 |
| EN633 | 2009 | Vietnam | Human | Systemic | 1 | 2 | [34] | PRJEB2670 |
| EN636 | 2009 | Vietnam | Human | Systemic | 1 | 2 | [34] | PRJEB2670 |
| EN652 | 2009 | Vietnam | Human | Systemic | 1 | 14 | [34] | PRJEB2670 |
| EN660 | 2009 | Vietnam | Human | Systemic | 1 | 2 | [34] | PRJEB2670 |
| FX109 | 2006 | Vietnam | Pig Healthy | Non-Clinical | 1 | 2 | [34] | PRJEB2670 |
| FX14 | 2006 | Vietnam | Pig Healthy | Non-Clinical | 1 | 2 | [34] | PRJEB2670 |
| FX147_3 | 2006 | Vietnam | Pig Healthy | Non-Clinical | 1 | 2 | [34] | PRJEB2670 |
| FX18 | 2006 | Vietnam | Pig Healthy | Non-Clinical | 1 | 2 | [34] | PRJEB2670 |
| FX184_2 | 2006 | Vietnam | Pig Healthy | Non-Clinical | 1 | 2 | [34] | PRJEB2670 |
| FX211 | 2007 | Vietnam | Pig Healthy | Non-Clinical | 1 | 2 | [34] | PRJEB2670 |
| FX22 | 2006 | Vietnam | Pig Healthy | Non-Clinical | 1 | 2 | [34] | PRJEB2670 |
| FX230 | 2007 | Vietnam | Pig Healthy | Non-Clinical | 1 | 2 | [34] | PRJEB2670 |
| FX275 | 2007 | Vietnam | Pig Healthy | Non-Clinical | 1 | 2 | [34] | PRJEB2670 |
| FX28_2 | 2006 | Vietnam | Pig Healthy | Non-Clinical | 1 | 2 | [34] | PRJEB2670 |
| FX328_4 | 2007 | Vietnam | Pig Healthy | Non-Clinical | 1 | 2 | [34] | PRJEB2670 |
| FX336_2 | 2007 | Vietnam | Pig Healthy | Non-Clinical | 1 | 2 | [34] | PRJEB2670 |
| FX40_1 | 2006 | Vietnam | Pig Healthy | Non-Clinical | 1 | 2 | [34] | PRJEB2670 |
| FX415_6 | 2007 | Vietnam | Pig Healthy | Non-Clinical | 1 | 2 | [34] | PRJEB2670 |
| FX419 | 2007 | Vietnam | Pig Healthy | Non-Clinical | 1 | 2 | [34] | PRJEB2670 |

| | | | | | | | | |
|---|---|---|---|---|---|---|---|---|
| FX428_1 | 2007 | Vietnam | Pig Healthy | Non-Clinical | 1 | 2 | [34] | PRJEB2670 |
| FX431 | 2007 | Vietnam | Pig Healthy | Non-Clinical | 1 | 2 | [34] | PRJEB2670 |
| FX478 | 2007 | Vietnam | Pig Healthy | Non-Clinical | 1 | 2 | [34] | PRJEB2670 |
| FX483_3 | 2007 | Vietnam | Pig Healthy | Non-Clinical | 1 | 2 | [34] | PRJEB2670 |
| FX50_2 | 2006 | Vietnam | Pig Healthy | Non-Clinical | 1 | 2 | [34] | PRJEB2670 |
| FX501 | 2007 | Vietnam | Pig Healthy | Non-Clinical | 1 | 2 | [34] | PRJEB2670 |
| FX507_1 | 2007 | Vietnam | Pig Healthy | Non-Clinical | 1 | 2 | [34] | PRJEB2670 |
| FX51_2 | NA | Vietnam | Pig Healthy | Non-Clinical | 1 | 2 | [34] | PRJEB2670 |
| FX527_2 | 2007 | Vietnam | Pig Healthy | Non-Clinical | 1 | 2 | [34] | PRJEB2670 |
| FX535_2 | 2007 | Vietnam | Pig Healthy | Non-Clinical | 1 | 2 | [34] | PRJEB2670 |
| FX540_1 | 2007 | Vietnam | Pig Healthy | Non-Clinical | 1 | 2 | [34] | PRJEB2670 |
| FX543_1 | 2007 | Vietnam | Pig Healthy | Non-Clinical | 1 | 2 | [34] | PRJEB2670 |
| FX59_4 | 2006 | Vietnam | Pig Healthy | Non-Clinical | 1 | 2 | [34] | PRJEB2670 |
| FX73_1 | 2006 | Vietnam | Pig Healthy | Non-Clinical | 1 | 2 | [34] | PRJEB2670 |
| FX74 | 2006 | Vietnam | Pig Healthy | Non-Clinical | 1 | 2 | [34] | PRJEB2670 |
| FX77_1 | 2006 | Vietnam | Pig Healthy | Non-Clinical | 1 | 2 | [34] | PRJEB2670 |
| FX90_2 | 2006 | Vietnam | Pig Healthy | Non-Clinical | 1 | 2 | [34] | PRJEB2670 |
| LOLA-SS001 | 2010 | UK | Pig Diseased | Systemic | NA | 2 | [34] | PRJEB8392 |
| LOLA-SS002 | 2010 | UK | Pig Unknown | Unknown | 1 | 14 | [34] | PRJEB8392 |
| LOLA-SS003 | 2010 | UK | Pig Unknown | Unknown | 1 | 14 | [34] | PRJEB8392 |
| LOLA-SS004 | 2010 | UK | Pig Diseased | Respiratory | 27 | 3 | [34] | PRJEB8392 |
| LOLA-SS005 | 2010 | UK | Pig Diseased | Systemic | NA | 3 | [34] | PRJEB8392 |
| LOLA-SS006 | 2010 | UK | Pig Diseased | Respiratory | NA | 4 | [34] | PRJEB8392 |
| LOLA-SS008 | 2010 | UK | Pig Unknown | Unknown | 29 | 7 | [34] | PRJEB8392 |
| LOLA-SS009 | 2010 | UK | Pig Diseased | Systemic | 29 | 7 | [34] | PRJEB8392 |
| LOLA-SS010 | 2010 | UK | Pig Unknown | Unknown | 1 | 14 | [34] | PRJEB8392 |

| | | | | | | | | |
|---|---|---|---|---|---|---|---|---|
| LOLA-SS011 | 2010 | UK | Pig Diseased | Systemic | 1 | 14 | [34] | PRJEB8392 |
| LSS1 | 2010 | UK | Pig Healthy | Non-Clinical | 28 | 2 | [34] | PRJEB8392 |
| LSS10 | 2010 | UK | Pig Unknown | Unknown | NA | NA | [34] | PRJEB8392 |
| LSS100 | 2011 | UK | Pig Unknown | Unknown | NA | 15 | [34] | PRJEB8392 |
| LSS11 | 2011 | UK | Pig Healthy | Non-Clinical | NA | 31 | [34] | PRJEB8392 |
| LSS12 | 2009 | UK | Pig Diseased | Respiratory | NA | 25 | [34] | PRJEB8392 |
| LSS13 | 2009 | UK | Pig Diseased | Respiratory | NA | 16 | [34] | PRJEB8392 |
| LSS16 | 2009 | UK | Pig Unknown | Unknown | NA | NA | [34] | PRJEB8392 |
| LSS18 | 2010 | UK | Pig Healthy | Non-Clinical | NA | 16 | [34] | PRJEB8392 |
| LSS2 | 2010 | UK | Pig Unknown | Unknown | NA | 15 | [34] | PRJEB8392 |
| LSS20 | 2010 | UK | Pig Healthy | Non-Clinical | 23 | 4 | [34] | PRJEB8392 |
| LSS21 | 2011 | UK | Pig Unknown | Unknown | 1 | 2 | [34] | PRJEB8392 |
| LSS22 | 2011 | UK | Pig Unknown | Unknown | 1 | 2 | [34] | PRJEB8392 |
| LSS23 | 2011 | UK | Pig Healthy | Non-Clinical | NA | 24 | [34] | PRJEB8392 |
| LSS24 | 2011 | UK | Pig Unknown | Unknown | 28 | 2 | [34] | PRJEB8392 |
| LSS25 | 2011 | UK | Pig Healthy | Non-Clinical | 28 | 2 | [34] | PRJEB8392 |
| LSS26 | 2011 | UK | Pig Unknown | Unknown | 28 | 2 | [34] | PRJEB8392 |
| LSS27 | 2011 | UK | Pig Healthy | Non-Clinical | NA | 10 | [34] | PRJEB8392 |
| LSS28 | 2011 | UK | Pig Unknown | Unknown | 1 | 2 | [34] | PRJEB8392 |
| LSS29 | 2011 | UK | Pig Unknown | Unknown | 28 | 2 | [34] | PRJEB8392 |
| LSS3 | 2010 | UK | Pig Healthy | Non-Clinical | 28 | 2 | [34] | PRJEB8392 |
| LSS30 | 2011 | UK | Pig Healthy | Non-Clinical | NA | 31 | [34] | PRJEB8392 |
| LSS31 | 2011 | UK | Pig Healthy | Non-Clinical | NA | 24 | [34] | PRJEB8392 |
| LSS32 | 2011 | UK | Pig Healthy | Non-Clinical | NA | 24 | [34] | PRJEB8392 |
| LSS33 | 2011 | UK | Pig Unknown | Unknown | NA | 4 | [34] | PRJEB8392 |
| LSS34 | 2011 | UK | Pig Healthy | Non-Clinical | 87 | 8 | [34] | PRJEB8392 |

| | | | | | | | | |
|---|---|---|---|---|---|---|---|---|
| LSS35 | 2011 | UK | Pig Unknown | Unknown | NA | 2 | [34] | PRJEB8392 |
| LSS36 | 2011 | UK | Pig Unknown | Unknown | NA | NA | [34] | PRJEB8392 |
| LSS37 | 2011 | UK | Pig Healthy | Non-Clinical | NA | 24 | [34] | PRJEB8392 |
| LSS38 | 2011 | UK | Pig Healthy | Non-Clinical | NA | NA | [34] | PRJEB8392 |
| LSS39 | 2011 | UK | Pig Healthy | Non-Clinical | NA | 31 | [34] | PRJEB8392 |
| LSS4 | 2010 | UK | Pig Unknown | Unknown | NA | 2 | [34] | PRJEB8392 |
| LSS40 | 2011 | UK | Pig Healthy | Non-Clinical | NA | 6 | [34] | PRJEB8392 |
| LSS41 | 2011 | UK | Pig Healthy | Non-Clinical | NA | 21 | [34] | PRJEB8392 |
| LSS42 | 2011 | UK | Pig Healthy | Non-Clinical | NA | 16 | [34] | PRJEB8392 |
| LSS43 | 2011 | UK | Pig Unknown | Unknown | NA | 16 | [34] | PRJEB8392 |
| LSS44 | 2011 | UK | Pig Healthy | Non-Clinical | NA | 15 | [34] | PRJEB8392 |
| LSS47 | 2011 | UK | Pig Healthy | Non-Clinical | NA | 16 | [34] | PRJEB8392 |
| LSS48 | 2011 | UK | Pig Healthy | Non-Clinical | NA | 31 | [34] | PRJEB8392 |
| LSS49 | 2011 | UK | Pig Healthy | Non-Clinical | NA | NA | [34] | PRJEB8392 |
| LSS5 | 2010 | UK | Pig Unknown | Unknown | NA | 2 | [34] | PRJEB8392 |
| LSS50 | 2011 | UK | Pig Unknown | Unknown | NA | 2 | [34] | PRJEB8392 |
| LSS51 | 2011 | UK | Pig Unknown | Unknown | 1 | 2 | [34] | PRJEB8392 |
| LSS52 | 2011 | UK | Pig Unknown | Unknown | NA | 19 | [34] | PRJEB8392 |
| LSS53 | 2011 | UK | Pig Healthy | Non-Clinical | NA | 21 | [34] | PRJEB8392 |
| LSS54 | 2011 | UK | Pig Healthy | Non-Clinical | NA | NA | [34] | PRJEB8392 |
| LSS55 | 2011 | UK | Pig Healthy | Non-Clinical | NA | 9 | [34] | PRJEB8392 |
| LSS56 | 2011 | UK | Pig Unknown | Unknown | 29 | 7 | [34] | PRJEB8392 |
| LSS57 | 2011 | UK | Pig Healthy | Non-Clinical | NA | NA | [34] | PRJEB8392 |
| LSS58 | 2011 | UK | Pig Healthy | Non-Clinical | NA | 31 | [34] | PRJEB8392 |
| LSS59 | 2011 | UK | Pig Healthy | Non-Clinical | NA | 10 | [34] | PRJEB8392 |
| LSS60 | 2011 | UK | Pig Healthy | Non-Clinical | NA | 31 | [34] | PRJEB8392 |

| | | | | | | | | |
|---|---|---|---|---|---|---|---|---|
| LSS61 | 2011 | UK | Pig Healthy | Non-Clinical | 28 | 2 | [34] | PRJEB8392 |
| LSS62 | 2011 | UK | Pig Healthy | Non-Clinical | 1 | 2 | [34] | PRJEB8392 |
| LSS63 | 2011 | UK | Pig Healthy | Non-Clinical | 1 | 2 | [34] | PRJEB8392 |
| LSS64 | 2011 | UK | Pig Healthy | Non-Clinical | NA | 21 | [34] | PRJEB8392 |
| LSS65 | 2011 | UK | Pig Healthy | Non-Clinical | NA | 10 | [34] | PRJEB8392 |
| LSS66 | 2011 | UK | Pig Unknown | Unknown | NA | 9 | [34] | PRJEB8392 |
| LSS67 | 2011 | UK | Pig Healthy | Non-Clinical | 87 | 8 | [34] | PRJEB8392 |
| LSS68 | 2011 | UK | Pig Healthy | Non-Clinical | 87 | 8 | [34] | PRJEB8392 |
| LSS69 | 2011 | UK | Pig Healthy | Non-Clinical | NA | 31 | [34] | PRJEB8392 |
| LSS70 | 2011 | UK | Pig Healthy | Non-Clinical | 87 | 9 | [34] | PRJEB8392 |
| LSS71 | 2011 | UK | Pig Healthy | Non-Clinical | 28 | 2 | [34] | PRJEB8392 |
| LSS72 | 2011 | UK | Pig Healthy | Non-Clinical | 28 | 2 | [34] | PRJEB8392 |
| LSS73 | 2011 | UK | Pig Unknown | Unknown | 94 | 3 | [34] | PRJEB8392 |
| LSS74 | 2011 | UK | Pig Unknown | Unknown | 94 | 3 | [34] | PRJEB8392 |
| LSS75 | 2011 | UK | Pig Healthy | Non-Clinical | NA | 31 | [34] | PRJEB8392 |
| LSS76 | 2011 | UK | Pig Healthy | Non-Clinical | 87 | 8 | [34] | PRJEB8392 |
| LSS77 | 2011 | UK | Pig Healthy | Non-Clinical | 28 | 2 | [34] | PRJEB8392 |
| LSS78 | 2011 | UK | Pig Healthy | Non-Clinical | NA | 12 | [34] | PRJEB8392 |
| LSS79 | 2011 | UK | Pig Healthy | Non-Clinical | NA | 15 | [34] | PRJEB8392 |
| LSS8 | 2010 | UK | Pig Unknown | Unknown | NA | 16 | [34] | PRJEB8392 |
| LSS80 | 2011 | UK | Pig Healthy | Non-Clinical | NA | 31 | [34] | PRJEB8392 |
| LSS81 | 2011 | UK | Pig Healthy | Non-Clinical | NA | 15 | [34] | PRJEB8392 |
| LSS82 | 2011 | UK | Pig Healthy | Non-Clinical | NA | 16 | [34] | PRJEB8392 |
| LSS83 | 2011 | UK | Pig Healthy | Non-Clinical | 29 | 7 | [34] | PRJEB8392 |
| LSS84 | 2011 | UK | Pig Healthy | Non-Clinical | 29 | 7 | [34] | PRJEB8392 |
| LSS85 | 2011 | UK | Pig Healthy | Non-Clinical | 29 | 7 | [34] | PRJEB8392 |

| | | | | | | | | |
|---|---|---|---|---|---|---|---|---|
| LSS86 | 2011 | UK | Pig Healthy | Non-Clinical | 28 | 2 | [34] | PRJEB8392 |
| LSS87 | 2011 | UK | Pig Unknown | Unknown | 28 | 2 | [34] | PRJEB8392 |
| LSS88 | 2011 | UK | Pig Healthy | Non-Clinical | NA | NA | [34] | PRJEB8392 |
| LSS89 | 2011 | UK | Pig Healthy | Non-Clinical | 87 | 8 | [34] | PRJEB8392 |
| LSS9 | 2010 | UK | Pig Healthy | Non-Clinical | NA | 10 | [34] | PRJEB8392 |
| LSS90 | 2011 | UK | Pig Healthy | Non-Clinical | NA | 31 | [34] | PRJEB8392 |
| LSS91 | 2011 | UK | Pig Healthy | Non-Clinical | NA | 6 | [34] | PRJEB8392 |
| LSS92 | 2011 | UK | Pig Healthy | Non-Clinical | 46 | 9 | [34] | PRJEB8392 |
| LSS94 | 2011 | UK | Pig Healthy | Non-Clinical | NA | 4 | [34] | PRJEB8392 |
| LSS95 | 2011 | UK | Pig Healthy | Non-Clinical | NA | 11 | [34] | PRJEB8392 |
| LSS96 | 2011 | UK | Pig Healthy | Non-Clinical | NA | 11 | [34] | PRJEB8392 |
| LSS97 | 2011 | UK | Pig Unknown | Unknown | 1 | 2 | [34] | PRJEB8392 |
| LSS98 | 2011 | UK | Pig Unknown | Unknown | 1 | 2 | [34] | PRJEB8392 |
| LSS99 | 2011 | UK | Pig Unknown | Unknown | NA | 30 | [34] | PRJEB8392 |
| SS1000 | 2010 | UK | Pig Diseased | Respiratory | 29 | 7 | [34] | PRJEB8392 |
| SS1001 | 2010 | UK | Pig Unknown | Unknown | 1 | 2 | [34] | PRJEB8392 |
| SS1002 | 2010 | UK | Pig Diseased | Systemic | 1 | 2 | [34] | PRJEB8392 |
| SS1004 | 2010 | UK | Pig Unknown | Unknown | NA | 2 | [34] | PRJEB8392 |
| SS1005 | 2010 | UK | Pig Diseased | Respiratory | 31 | 3 | [34] | PRJEB8392 |
| SS1006 | 2010 | UK | Pig Unknown | Unknown | 1 | 2 | [34] | PRJEB8392 |
| SS1007 | 2010 | UK | Pig Diseased | Systemic | 29 | 7 | [34] | PRJEB8392 |
| SS1008 | 2010 | UK | Pig Diseased | Systemic | NA | 2 | [34] | PRJEB8392 |
| SS1009 | 2010 | UK | Pig Diseased | Respiratory | 28 | 2 | [34] | PRJEB8392 |
| SS101 | 2010 | Vietnam | Human | Systemic | 1 | 2 | [34] | PRJEB2670 |
| SS1010 | 2010 | UK | Pig Unknown | Unknown | 1 | 2 | [34] | PRJEB8392 |
| SS1011 | 2011 | UK | Pig Diseased | Systemic | 1 | 14 | [34] | PRJEB8392 |

| | | | | | | | | |
|---|---|---|---|---|---|---|---|---|
| SS1012 | 2010 | UK | Pig Diseased | Systemic | 1 | 2 | [34] | PRJEB8392 |
| SS1013 | 2011 | UK | Pig Diseased | Respiratory | NA | NA | [34] | PRJEB8392 |
| SS1014 | 2011 | UK | Pig Diseased | Respiratory | 27 | 3 | [34] | PRJEB8392 |
| SS1015 | 2010 | UK | Pig Diseased | Respiratory | 27 | 3 | [34] | PRJEB8392 |
| SS1016 | 2010 | UK | Pig Diseased | Respiratory | 87 | 8 | [34] | PRJEB8392 |
| SS1017 | 2010 | UK | Pig Diseased | Respiratory | 1 | 2 | [34] | PRJEB8392 |
| SS1018 | 2010 | UK | Pig Diseased | Respiratory | 29 | 7 | [34] | PRJEB8392 |
| SS102 | 2010 | Vietnam | Human | Systemic | 144 | 2 | [34] | PRJEB2670 |
| SS1021 | 2010 | UK | Pig Diseased | Respiratory | 27 | 3 | [34] | PRJEB8392 |
| SS1022 | 2010 | UK | Pig Diseased | Respiratory | 32 | 7 | [34] | PRJEB8392 |
| SS1023 | 2010 | UK | Pig Unknown | Unknown | 1 | 2 | [34] | PRJEB8392 |
| SS1024 | 2010 | UK | Pig Diseased | Systemic | 1 | 2 | [34] | PRJEB8392 |
| SS1025 | 2010 | UK | Pig Diseased | Respiratory | 1 | 2 | [34] | PRJEB8392 |
| SS1026 | 2010 | UK | Pig Diseased | Systemic | 1 | 2 | [34] | PRJEB8392 |
| SS1027 | 2010 | UK | Pig Diseased | Respiratory | 1 | 14 | [34] | PRJEB8392 |
| SS1028 | 2010 | UK | Pig Diseased | Systemic | NA | 4 | [34] | PRJEB8392 |
| SS1029 | 2010 | UK | Pig Unknown | Unknown | 1 | 14 | [34] | PRJEB8392 |
| SS1030 | 2010 | UK | Pig Unknown | Unknown | 27 | 3 | [34] | PRJEB8392 |
| SS1032 | 2010 | UK | Pig Unknown | Unknown | 1 | 2 | [34] | PRJEB8392 |
| SS1033 | 2010 | UK | Pig Diseased | Systemic | 1 | 14 | [34] | PRJEB8392 |
| SS1034 | 2010 | UK | Pig Diseased | Respiratory | 28 | 2 | [34] | PRJEB8392 |
| SS1035 | 2010 | UK | Pig Diseased | Systemic | NA | 8 | [34] | PRJEB8392 |
| SS1036 | 2010 | UK | Pig Unknown | Unknown | 1 | 2 | [34] | PRJEB8392 |
| SS1037 | 2010 | UK | Pig Diseased | Respiratory | 87 | 8 | [34] | PRJEB8392 |
| SS1038 | 2010 | UK | Pig Diseased | Systemic | 1 | 14 | [34] | PRJEB8392 |
| SS1039 | 2010 | UK | Pig Unknown | Unknown | 1 | 14 | [34] | PRJEB8392 |

| | | | | | | | | |
|---|---|---|---|---|---|---|---|---|
| SS1040 | 2010 | UK | Pig Diseased | Respiratory | 94 | 4 | [34] | PRJEB8392 |
| SS1041 | 2010 | UK | Pig Diseased | Respiratory | 94 | 4 | [34] | PRJEB8392 |
| SS1042 | 2010 | UK | Pig Diseased | Respiratory | 94 | 4 | [34] | PRJEB8392 |
| SS1043 | 2010 | UK | Pig Diseased | Respiratory | 28 | 2 | [34] | PRJEB8392 |
| SS1046 | 2010 | UK | Pig Unknown | Unknown | NA | 10 | [34] | PRJEB8392 |
| SS1048 | 2010 | UK | Pig Diseased | Respiratory | 23 | 4 | [34] | PRJEB8392 |
| SS1051 | 2010 | UK | Pig Diseased | Systemic | 29 | 7 | [34] | PRJEB8392 |
| SS1052 | 2010 | UK | Pig Diseased | Respiratory | NA | 3 | [34] | PRJEB8392 |
| SS1053 | 2010 | UK | Pig Diseased | Systemic | 25 | 2 | [34] | PRJEB8392 |
| SS1054 | NA | UK | Pig Unknown | Unknown | 1 | 2 | [34] | PRJEB8392 |
| SS1055 | 2010 | UK | Pig Diseased | Systemic | 1 | 2 | [34] | PRJEB8392 |
| SS1056 | 2010 | UK | Pig Unknown | Unknown | NA | 8 | [34] | PRJEB8392 |
| SS1057 | 2010 | UK | Pig Diseased | Respiratory | 31 | 3 | [34] | PRJEB8392 |
| SS1058 | 2010 | UK | Pig Unknown | Unknown | 1 | 14 | [34] | PRJEB8392 |
| SS1061 | 2010 | UK | Pig Diseased | Respiratory | 87 | 8 | [34] | PRJEB8392 |
| SS1062 | 2010 | UK | Pig Diseased | Respiratory | NA | 15 | [34] | PRJEB8392 |
| SS1063 | 2010 | UK | Pig Diseased | Systemic | 1 | 2 | [34] | PRJEB8392 |
| SS1064 | 2010 | UK | Pig Diseased | Respiratory | 25 | 2 | [34] | PRJEB8392 |
| SS1065 | 2010 | UK | Pig Diseased | Systemic | 1 | 2 | [34] | PRJEB8392 |
| SS1066 | 2010 | UK | Pig Diseased | Systemic | 1 | 2 | [34] | PRJEB8392 |
| SS1067 | 2010 | UK | Pig Unknown | Unknown | NA | 16 | [34] | PRJEB8392 |
| SS1070 | 2010 | UK | Pig Diseased | Systemic | 1 | 2 | [34] | PRJEB8392 |
| SS1072 | 2010 | UK | Pig Diseased | Systemic | NA | 2 | [34] | PRJEB8392 |
| SS1073 | 2010 | UK | Pig Diseased | Systemic | 1 | 2 | [34] | PRJEB8392 |
| SS13 | 2005 | Vietnam | Human | Systemic | 1 | 2 | [34] | PRJEB2670 |
| SS14 | 2005 | Vietnam | Human | Systemic | 1 | 2 | [34] | PRJEB2670 |

| | | | | | | | | |
|---|---|---|---|---|---|---|---|---|
| SS15 | 2005 | Vietnam | Human | Systemic | 1 | 2 | [34] | PRJEB2670 |
| SS16 | 2005 | Vietnam | Human | Systemic | 1 | 2 | [34] | PRJEB2670 |
| SS17 | 2005 | Vietnam | Human | Systemic | 1 | 2 | [34] | PRJEB2670 |
| SS19 | 2005 | Vietnam | Human | Systemic | 1 | 2 | [34] | PRJEB2670 |
| SS20 | 2005 | Vietnam | Human | Systemic | 1 | 2 | [34] | PRJEB2670 |
| SS21 | 2005 | Vietnam | Human | Systemic | 1 | 2 | [34] | PRJEB2670 |
| SS22 | 2005 | Vietnam | Human | Systemic | 1 | 2 | [34] | PRJEB2670 |
| SS23 | 2005 | Vietnam | Human | Systemic | 1 | 2 | [34] | PRJEB2670 |
| SS24 | 2005 | Vietnam | Human | Systemic | 1 | 2 | [34] | PRJEB2670 |
| SS25 | 2005 | Vietnam | Human | Systemic | 1 | 2 | [34] | PRJEB2670 |
| SS26 | 2005 | Vietnam | Human | Systemic | 1 | 2 | [34] | PRJEB2670 |
| SS27 | 2006 | Vietnam | Human | Systemic | 1 | 2 | [34] | PRJEB2670 |
| SS28 | 2006 | Vietnam | Human | Systemic | 1 | 2 | [34] | PRJEB2670 |
| SS29 | 2006 | Vietnam | Human | Systemic | 1 | 2 | [34] | PRJEB2670 |
| SS30 | 2006 | Vietnam | Human | Systemic | 1 | 2 | [34] | PRJEB2670 |
| SS31 | 2006 | Vietnam | Human | Systemic | 1 | 2 | [34] | PRJEB2670 |
| SS32 | 2006 | Vietnam | Human | Systemic | 1 | 2 | [34] | PRJEB2670 |
| SS33 | 2006 | Vietnam | Human | Systemic | 1 | 2 | [34] | PRJEB2670 |
| SS35 | 2006 | Vietnam | Human | Systemic | 1 | 2 | [34] | PRJEB2670 |
| SS36 | 2006 | Vietnam | Human | Systemic | 1 | 2 | [34] | PRJEB2670 |
| SS37 | 2006 | Vietnam | Human | Systemic | 1 | 2 | [34] | PRJEB2670 |
| SS38 | 2006 | Vietnam | Human | Systemic | 1 | 2 | [34] | PRJEB2670 |
| SS40 | 2006 | Vietnam | Human | Systemic | 1 | 2 | [34] | PRJEB2670 |
| SS44 | 2009 | Vietnam | Human | Systemic | 1 | 2 | [34] | PRJEB2670 |
| SS46 | 2009 | Vietnam | Human | Systemic | 1 | 2 | [34] | PRJEB2670 |
| SS47 | 2009 | Vietnam | Human | Systemic | 1 | 2 | [34] | PRJEB2670 |

| | | | | | | | | |
|---|---|---|---|---|---|---|---|---|
| SS61 | 2010 | Vietnam | Human | Systemic | 1 | 2 | [34] | PRJEB2670 |
| SS65 | 2010 | Vietnam | Human | Systemic | 1 | 2 | [34] | PRJEB2670 |
| SS69 | 2010 | Vietnam | Human | Systemic | 1 | 2 | [34] | PRJEB2670 |
| SS73 | 2010 | Vietnam | Human | Systemic | 1 | 2 | [34] | PRJEB2670 |
| SS74 | 2010 | Vietnam | Human | Systemic | NA | 2 | [34] | PRJEB2670 |
| SS76 | 2010 | Vietnam | Human | Systemic | 1 | 2 | [34] | PRJEB2670 |
| SS78 | 2010 | Vietnam | Human | Systemic | 1 | 2 | [34] | PRJEB2670 |
| SS79 | 2010 | Vietnam | Human | Systemic | 1 | 2 | [34] | PRJEB2670 |
| SS81 | 2010 | Vietnam | Human | Systemic | 1 | 2 | [34] | PRJEB2670 |
| SS82 | 2010 | Vietnam | Human | Systemic | 1 | 2 | [34] | PRJEB2670 |
| SS83 | 2010 | Vietnam | Human | Systemic | 1 | 2 | [34] | PRJEB2670 |
| SS84 | 2010 | Vietnam | Human | Systemic | 1 | 2 | [34] | PRJEB2670 |
| SS85 | 2010 | Vietnam | Human | Systemic | 144 | 2 | [34] | PRJEB2670 |
| SS87 | 2010 | Vietnam | Human | Systemic | 1 | 2 | [34] | PRJEB2670 |
| SS967 | 2010 | UK | Pig Unknown | Unknown | 23 | 4 | [34] | PRJEB8392 |
| SS969 | 2010 | UK | Pig Diseased | Respiratory | 1 | 2 | [34] | PRJEB8392 |
| SS97 | 2010 | Vietnam | Human | Systemic | 144 | 2 | [34] | PRJEB2670 |
| SS970 | 2010 | UK | Pig Diseased | Systemic | 1 | 2 | [34] | PRJEB8392 |
| SS972 | 2010 | UK | Pig Diseased | Systemic | 1 | 2 | [34] | PRJEB8392 |
| SS973 | 2010 | UK | Pig Diseased | Systemic | 1 | 2 | [34] | PRJEB8392 |
| SS974 | 2010 | UK | Pig Unknown | Unknown | 28 | 2 | [34] | PRJEB8392 |
| SS975 | 2010 | UK | Pig Diseased | Respiratory | NA | NA | [34] | PRJEB8392 |
| SS976 | 2010 | UK | Pig Diseased | Systemic | 1 | 2 | [34] | PRJEB8392 |
| SS979 | 2010 | UK | Pig Unknown | Unknown | 1 | 14 | [34] | PRJEB8392 |
| SS981 | 2010 | UK | Pig Diseased | Systemic | NA | 9 | [34] | PRJEB8392 |
| SS982 | 2010 | UK | Pig Unknown | Unknown | 1 | 14 | [34] | PRJEB8392 |

| | | | | | | | | |
|---|---|---|---|---|---|---|---|---|
| SS983 | 2010 | UK | Pig Unknown | Unknown | 1 | 14 | [34] | PRJEB8392 |
| SS985 | 2010 | UK | Pig Unknown | Unknown | NA | NA | [34] | PRJEB8392 |
| SS986 | 2010 | UK | Pig Diseased | Respiratory | NA | 12 | [34] | PRJEB8392 |
| SS987 | 2010 | UK | Pig Unknown | Unknown | 1 | 14 | [34] | PRJEB8392 |
| SS989 | 2010 | UK | Pig Diseased | Respiratory | 28 | 2 | [34] | PRJEB8392 |
| SS99 | 2010 | Vietnam | Human | Systemic | 1 | 2 | [34] | PRJEB2670 |
| SS991 | 2010 | UK | Pig Diseased | Systemic | 1 | 2 | [34] | PRJEB8392 |
| SS992 | 2010 | UK | Pig Diseased | Respiratory | 1 | 2 | [34] | PRJEB8392 |
| SS993 | 2010 | UK | Pig Healthy | Non-Clinical | NA | NA | [34] | PRJEB8392 |
| SS994 | 2010 | UK | Pig Diseased | Systemic | 1 | 2 | [34] | PRJEB8392 |
| SS995 | 2010 | UK | Pig Unknown | Unknown | 1 | 14 | [34] | PRJEB8392 |
| SS996 | 2010 | UK | Pig Diseased | Respiratory | 28 | 2 | [34] | PRJEB8392 |
| SS997 | 2010 | UK | Pig Diseased | Respiratory | NA | 8 | [34] | PRJEB8392 |
| SS998 | 2010 | UK | Pig Diseased | Respiratory | 1 | 2 | [34] | PRJEB8392 |
| SS999 | 2010 | UK | Pig Diseased | Respiratory | 108 | 3 | [34] | PRJEB8392 |
| VS2903 | NA | Vietnam | Human | Systemic | 1 | 2 | [34] | PRJEB2670 |
| VS2923 | NA | Vietnam | Human | Systemic | 1 | 2 | [34] | PRJEB2670 |
| VS2979 | NA | Vietnam | Human | Systemic | 1 | 2 | [34] | PRJEB2670 |
| VS3121 | NA | Vietnam | Human | Systemic | 1 | 2 | [34] | PRJEB2670 |
| VS329 | NA | Vietnam | Human | Systemic | 1 | 2 | [34] | PRJEB2670 |
| VS3309 | NA | Vietnam | Human | Unknown | 1 | 2 | [34] | PRJEB2670 |
| VS3466 | NA | Vietnam | Human | Systemic | 1 | 2 | [34] | PRJEB2670 |
| VS3513 | 2009 | Vietnam | Human | Systemic | 1 | 2 | [34] | PRJEB2670 |
| VS3517 | NA | Vietnam | Human | Systemic | 1 | 2 | [34] | PRJEB2670 |
| VS3537 | NA | Vietnam | Human | Systemic | 1 | 2 | [34] | PRJEB2670 |
| VS3543 | NA | Vietnam | Human | Systemic | 1 | 2 | [34] | PRJEB2670 |

| | | | | | | | | |
|---|---|---|---|---|---|---|---|---|
| VS3569 | 2009 | Vietnam | Human | Systemic | 1 | 2 | [34] | PRJEB2670 |
| VS3575 | 2009 | Vietnam | Human | Systemic | 1 | 2 | [34] | PRJEB2670 |
| VS3587 | NA | Vietnam | Human | Systemic | 1 | 2 | [34] | PRJEB2670 |
| VS3640 | 2009 | Vietnam | Human | Systemic | 1 | 2 | [34] | PRJEB2670 |
| VS3699 | NA | Vietnam | Human | Systemic | 1 | 2 | [34] | PRJEB2670 |
| VS3709 | NA | Vietnam | Human | Systemic | 1 | 2 | [34] | PRJEB2670 |
| VS3756 | NA | Vietnam | Human | Systemic | 1 | 2 | [34] | PRJEB2670 |
| VS3804 | NA | Vietnam | Human | Systemic | 1 | 2 | [34] | PRJEB2670 |
| VS3861 | NA | Vietnam | Human | Systemic | 1 | 2 | [34] | PRJEB2670 |
| VS3865 | NA | Vietnam | Human | Systemic | 1 | 2 | [34] | PRJEB2670 |
| VS3878 | NA | Vietnam | Human | Systemic | 1 | 2 | [34] | PRJEB2670 |
| VS3881 | NA | Vietnam | Human | Unknown | 1 | 2 | [34] | PRJEB2670 |
| VS3886 | NA | Vietnam | Human | Systemic | 1 | 2 | [34] | PRJEB2670 |
| VS3888 | NA | Vietnam | Human | Systemic | 1 | 2 | [34] | PRJEB2670 |
| ERR1055550 | 1992 | Netherlands | Pig Diseased | Systemic | 1 | 14 | [35] | PRJEB11219 |
| ERR1055551 | 1992 | Netherlands | Pig Diseased | Systemic | 1 | 14 | [35] | PRJEB11219 |
| ERR1055552 | NA | Netherlands | Pig Diseased | Systemic | 13 | 14 | [35] | PRJEB11219 |
| ERR1055553 | 1996 | Netherlands | Pig Diseased | Systemic | 136 | 9 | [35] | PRJEB11219 |
| ERR1055554 | 1986 | Netherlands | Human | Systemic | 20 | 2 | [35] | PRJEB11219 |
| ERR1055555 | 1987 | Netherlands | Human | Systemic | 1 | 2 | [35] | PRJEB11219 |
| ERR1055556 | 1989 | Netherlands | Human | Systemic | 1 | 2 | [35] | PRJEB11219 |
| ERR1055557 | 1990 | Netherlands | Human | Systemic | 134 | 2 | [35] | PRJEB11219 |
| ERR1055558 | 1990 | Netherlands | Human | Systemic | 1 | 2 | [35] | PRJEB11219 |
| ERR1055559 | 1992 | Netherlands | Human | Systemic | 20 | 2 | [35] | PRJEB11219 |
| ERR1055560 | 1993 | Netherlands | Human | Systemic | 20 | 2 | [35] | PRJEB11219 |
| ERR1055561 | 1994 | Netherlands | Human | Systemic | 1 | 2 | [35] | PRJEB11219 |

| | | | | | | | | |
|---|---|---|---|---|---|---|---|---|
| ERR1055562 | 1994 | Netherlands | Human | Systemic | 20 | 2 | [35] | PRJEB11219 |
| ERR1055563 | 1994 | Netherlands | Human | Systemic | 1 | 2 | [35] | PRJEB11219 |
| ERR1055564 | 1994 | Netherlands | Human | Systemic | 20 | 2 | [35] | PRJEB11219 |
| ERR1055565 | 1995 | Netherlands | Human | Systemic | 20 | 2 | [35] | PRJEB11219 |
| ERR1055566 | 1997 | Netherlands | Human | Systemic | 1 | 2 | [35] | PRJEB11219 |
| ERR1055567 | 1997 | Netherlands | Human | Systemic | 1 | 2 | [35] | PRJEB11219 |
| ERR1055568 | 2000 | Netherlands | Human | Systemic | 1 | 2 | [35] | PRJEB11219 |
| ERR1055569 | 2000 | Netherlands | Human | Systemic | 20 | 2 | [35] | PRJEB11219 |
| ERR1055570 | 2000 | Netherlands | Human | Systemic | 1 | 2 | [35] | PRJEB11219 |
| ERR1055571 | 2001 | Netherlands | Human | Systemic | 1 | 2 | [35] | PRJEB11219 |
| ERR1055572 | 2001 | Netherlands | Human | Systemic | 20 | 2 | [35] | PRJEB11219 |
| ERR1055573 | 2003 | Netherlands | Human | Systemic | 20 | 2 | [35] | PRJEB11219 |
| ERR1055574 | 2005 | Netherlands | Human | Systemic | 20 | 2 | [35] | PRJEB11219 |
| ERR1055575 | 2006 | Netherlands | Human | Systemic | 146 | 2 | [35] | PRJEB11219 |
| ERR1055576 | 2006 | Netherlands | Human | Systemic | 1 | 2 | [35] | PRJEB11219 |
| ERR1055577 | 2007 | Netherlands | Human | Systemic | 1 | 2 | [35] | PRJEB11219 |
| ERR1055578 | 2006 | Netherlands | Pig Diseased | Systemic | 220 | 9 | [35] | PRJEB11219 |
| ERR1055579 | 2006 | Netherlands | Pig Diseased | Systemic | 13 | 14 | [35] | PRJEB11219 |
| ERR1055580 | 2006 | Netherlands | Pig Diseased | Systemic | 16 | 9 | [35] | PRJEB11219 |
| ERR1055581 | 2006 | Netherlands | Pig Diseased | Systemic | 16 | 9 | [35] | PRJEB11219 |
| ERR1055582 | 2006 | Netherlands | Pig Diseased | Systemic | 87 | 8 | [35] | PRJEB11219 |
| ERR1055583 | 2006 | Netherlands | Pig Diseased | Systemic | 29 | 7 | [35] | PRJEB11219 |
| ERR1055584 | 1991 | Netherlands | Pig Diseased | Systemic | 1 | 14 | [35] | PRJEB11219 |
| ERR1055585 | 1992 | Netherlands | Pig Diseased | Systemic | 132 | 14 | [35] | PRJEB11219 |
| ERR1055586 | 1996 | Netherlands | Pig Diseased | Systemic | 20 | 2 | [35] | PRJEB11219 |
| ERR1055587 | 1996 | Netherlands | Pig Diseased | Systemic | 16 | 9 | [35] | PRJEB11219 |

| | | | | | | | | |
|---|---|---|---|---|---|---|---|---|
| ERR1055588 | 1997 | Netherlands | Pig Diseased | Systemic | 16 | 9 | [35] | PRJEB11219 |
| ERR1055589 | 1998 | Netherlands | Pig Diseased | Systemic | 16 | 9 | [35] | PRJEB11219 |
| ERR1055590 | 1999 | Netherlands | Pig Diseased | Systemic | 16 | 9 | [35] | PRJEB11219 |
| ERR1055591 | 1999 | Netherlands | Pig Diseased | Systemic | 28 | 2 | [35] | PRJEB11219 |
| ERR1055592 | 1999 | Netherlands | Pig Diseased | Systemic | 1 | 2 | [35] | PRJEB11219 |
| ERR1055593 | 2007 | Netherlands | Pig Diseased | Systemic | 16 | 9 | [35] | PRJEB11219 |
| ERR1055594 | 2000 | Netherlands | Pig Diseased | Systemic | 1 | 2 | [35] | PRJEB11219 |
| ERR1055595 | 2000 | Netherlands | Pig Diseased | Systemic | 183 | 31 | [35] | PRJEB11219 |
| ERR1055596 | 2000 | Netherlands | Pig Diseased | Systemic | 16 | 9 | [35] | PRJEB11219 |
| ERR1055597 | 2001 | Netherlands | Pig Diseased | Systemic | 16 | 9 | [35] | PRJEB11219 |
| ERR1055598 | 2001 | Netherlands | Pig Diseased | Systemic | 1 | 14 | [35] | PRJEB11219 |
| ERR1055599 | 2001 | Netherlands | Pig Diseased | Systemic | 1 | 2 | [35] | PRJEB11219 |
| ERR1055600 | 2002 | Netherlands | Pig Diseased | Systemic | 218 | 16 | [35] | PRJEB11219 |
| ERR1055601 | 2002 | Netherlands | Pig Diseased | Systemic | 16 | 9 | [35] | PRJEB11219 |
| ERR1055602 | 2002 | Netherlands | Pig Diseased | Systemic | 29 | 7 | [35] | PRJEB11219 |
| ERR1055603 | 2002 | Netherlands | Pig Diseased | Systemic | 13 | 14 | [35] | PRJEB11219 |
| ERR1055604 | 2002 | Netherlands | Pig Diseased | Systemic | 16 | 9 | [35] | PRJEB11219 |
| ERR1055605 | 2002 | Netherlands | Pig Diseased | Systemic | 1 | 14 | [35] | PRJEB11219 |
| ERR1055606 | 2003 | Netherlands | Pig Diseased | Systemic | 1 | 2 | [35] | PRJEB11219 |
| ERR1055607 | 2003 | Netherlands | Pig Diseased | Systemic | 15 | 3 | [35] | PRJEB11219 |
| ERR1055608 | 2003 | Netherlands | Pig Diseased | Systemic | 136 | 9 | [35] | PRJEB11219 |
| ERR1055609 | 2003 | Netherlands | Pig Diseased | Systemic | 16 | 9 | [35] | PRJEB11219 |
| ERR1055610 | 2003 | Netherlands | Pig Diseased | Systemic | 13 | 14 | [35] | PRJEB11219 |
| ERR1055611 | 2004 | Netherlands | Pig Diseased | Systemic | 16 | 9 | [35] | PRJEB11219 |
| ERR1055612 | 2004 | Netherlands | Pig Diseased | Systemic | 17 | 4 | [35] | PRJEB11219 |
| ERR1055613 | 2004 | Netherlands | Pig Diseased | Systemic | NA | 7 | [35] | PRJEB11219 |

| | | | | | | | | |
|---|---|---|---|---|---|---|---|---|
| ERR1055614 | 2004 | Netherlands | Pig Diseased | Systemic | 15 | 3 | [35] | PRJEB11219 |
| ERR1055615 | 2004 | Netherlands | Pig Diseased | Systemic | 1 | 2 | [35] | PRJEB11219 |
| ERR1055616 | 2004 | Netherlands | Pig Diseased | Systemic | 29 | 7 | [35] | PRJEB11219 |
| ERR1055617 | 2005 | Netherlands | Pig Diseased | Systemic | 29 | 7 | [35] | PRJEB11219 |
| ERR1055618 | 2005 | Netherlands | Pig Diseased | Systemic | 155 | 9 | [35] | PRJEB11219 |
| ERR1055619 | 2005 | Netherlands | Pig Diseased | Systemic | 17 | 4 | [35] | PRJEB11219 |
| ERR1055620 | 2005 | Netherlands | Pig Diseased | Systemic | 16 | 9 | [35] | PRJEB11219 |
| ERR1055621 | 2005 | Netherlands | Pig Diseased | Systemic | 1 | 2 | [35] | PRJEB11219 |
| ERR1055622 | 2005 | Netherlands | Pig Diseased | Systemic | 151 | 9 | [35] | PRJEB11219 |
| ERR1055623 | 2005 | Netherlands | Pig Diseased | Systemic | 136 | 9 | [35] | PRJEB11219 |
| ERR1055624 | 2005 | Netherlands | Pig Diseased | Systemic | 16 | 9 | [35] | PRJEB11219 |
| ERR1055625 | 2006 | Netherlands | Pig Diseased | Systemic | 16 | 9 | [35] | PRJEB11219 |
| ERR1055626 | 2006 | Netherlands | Pig Diseased | Systemic | 136 | 9 | [35] | PRJEB11219 |
| ERR1055627 | 2006 | Netherlands | Pig Diseased | Systemic | 16 | 9 | [35] | PRJEB11219 |
| ERR1055628 | 2006 | Netherlands | Pig Diseased | Systemic | 1 | 2 | [35] | PRJEB11219 |
| ERR1055629 | 2006 | Netherlands | Pig Diseased | Systemic | 29 | 7 | [35] | PRJEB11219 |
| ERR1055630 | 2006 | Netherlands | Pig Diseased | Systemic | 20 | 2 | [35] | PRJEB11219 |
| ERR1055631 | 2006 | Netherlands | Pig Diseased | Systemic | 17 | 4 | [35] | PRJEB11219 |
| ERR1055632 | 2007 | Netherlands | Pig Diseased | Systemic | 16 | 9 | [35] | PRJEB11219 |
| ERR1055633 | 2007 | Netherlands | Pig Diseased | Systemic | 1 | 2 | [35] | PRJEB11219 |
| ERR1055634 | 2007 | Netherlands | Pig Diseased | Systemic | 184 | 17 | [35] | PRJEB11219 |
| ERR1055635 | 2007 | Netherlands | Pig Diseased | Systemic | 16 | 9 | [35] | PRJEB11219 |
| ERR1055636 | 2007 | Netherlands | Pig Diseased | Systemic | 1 | 2 | [35] | PRJEB11219 |
| ERR1055637 | 2007 | Netherlands | Pig Diseased | Systemic | 149 | 14 | [35] | PRJEB11219 |
| ERR1055638 | 2007 | Netherlands | Pig Diseased | Systemic | 136 | 9 | [35] | PRJEB11219 |
| ERR1055639 | 2007 | Netherlands | Pig Diseased | Systemic | 198 | 8 | [35] | PRJEB11219 |

| | | | | | | | | |
|---|---|---|---|---|---|---|---|---|
| ERR1055640 | 2008 | Netherlands | Pig Diseased | Systemic | 1 | 14 | [35] | PRJEB11219 |
| ERR1055641 | 2008 | Netherlands | Pig Diseased | Systemic | 1 | 2 | [35] | PRJEB11219 |
| ERR1055642 | 2008 | Netherlands | Pig Diseased | Systemic | 16 | 9 | [35] | PRJEB11219 |
| ERR1055643 | 2008 | Netherlands | Pig Diseased | Systemic | 20 | 2 | [35] | PRJEB11219 |
| ERR1055644 | 2008 | Netherlands | Pig Diseased | Systemic | 153 | 8 | [35] | PRJEB11219 |
| ERR1055645 | 2003 | Netherlands | Pig Diseased | Systemic | 13 | 14 | [35] | PRJEB11219 |
| ERR1055646 | 1982 | Netherlands | Pig Diseased | Systemic | 1 | 2 | [35] | PRJEB11219 |
| ERR1055647 | 1988 | Netherlands | Pig Diseased | Systemic | 1 | 2 | [35] | PRJEB11219 |
| 083_13A | 2013 | UK | Pig Healthy | Non-Clinical | NA | 6 | [36] | PRJEB6250 |
| 083_17A | 2013 | UK | Pig Healthy | Non-Clinical | NA | 25 | [36] | PRJEB6250 |
| 083_17B | 2013 | UK | Pig Healthy | Non-Clinical | NA | NA | [36] | PRJEB6250 |
| 083_17C | 2013 | UK | Pig Healthy | Non-Clinical | NA | NA | [36] | PRJEB6250 |
| 083_1C | 2013 | UK | Pig Healthy | Non-Clinical | NA | NA | [36] | PRJEB6250 |
| 083_22A | 2013 | UK | Pig Healthy | Non-Clinical | NA | 15 | [36] | PRJEB6250 |
| 083_24C | 2013 | UK | Pig Healthy | Non-Clinical | NA | 25 | [36] | PRJEB6250 |
| 083_25C | 2013 | UK | Pig Healthy | Non-Clinical | NA | 10 | [36] | PRJEB6250 |
| 083_2A | 2013 | UK | Pig Healthy | Non-Clinical | NA | 31 | [36] | PRJEB6250 |
| 083_2B | 2013 | UK | Pig Healthy | Non-Clinical | NA | 31 | [36] | PRJEB6250 |
| 083_3A | 2013 | UK | Pig Healthy | Non-Clinical | NA | 10 | [36] | PRJEB6250 |
| 083_3B | 2013 | UK | Pig Healthy | Non-Clinical | NA | 31 | [36] | PRJEB6250 |
| 125_1A | 2013 | UK | Pig Healthy | Non-Clinical | NA | 6 | [36] | PRJEB6250 |
| 270_16A | 2013 | UK | Pig Healthy | Non-Clinical | NA | NA | [36] | PRJEB6250 |
| 270_21B | 2013 | UK | Pig Healthy | Non-Clinical | NA | 30 | [36] | PRJEB6250 |
| 270_21C | 2013 | UK | Pig Healthy | Non-Clinical | NA | 30 | [36] | PRJEB6250 |
| 270_23C | 2013 | UK | Pig Healthy | Non-Clinical | NA | NA | [36] | PRJEB6250 |
| 270_6A | 2013 | UK | Pig Healthy | Non-Clinical | NA | 4 | [36] | PRJEB6250 |

| | | | | | | | | |
|---|---|---|---|---|---|---|---|---|
| 359_6B | 2013 | UK | Pig Healthy | Non-Clinical | NA | 10 | [36] | PRJEB6250 |
| 427_17B | 2013 | UK | Pig Healthy | Non-Clinical | NA | 15 | [36] | PRJEB6250 |
| 427_17C | 2013 | UK | Pig Healthy | Non-Clinical | NA | NA | [36] | PRJEB6250 |
| 427_5B | 2013 | UK | Pig Healthy | Non-Clinical | NA | 16 | [36] | PRJEB6250 |
| 427_7C | 2013 | UK | Pig Healthy | Non-Clinical | NA | 9 | [36] | PRJEB6250 |
| 632_4A | 2013 | UK | Pig Healthy | Non-Clinical | NA | 6 | [36] | PRJEB6250 |
| 632_6A | 2013 | UK | Pig Healthy | Non-Clinical | NA | 6 | [36] | PRJEB6250 |
| 632_6B | 2013 | UK | Pig Healthy | Non-Clinical | NA | 6 | [36] | PRJEB6250 |
| 632_6C | 2013 | UK | Pig Healthy | Non-Clinical | NA | 6 | [36] | PRJEB6250 |
| 632_7B | 2013 | UK | Pig Healthy | Non-Clinical | NA | 16 | [36] | PRJEB6250 |
| 632_7C | 2013 | UK | Pig Healthy | Non-Clinical | NA | 31 | [36] | PRJEB6250 |
| 632_9A | 2013 | UK | Pig Healthy | Non-Clinical | NA | 16 | [36] | PRJEB6250 |
| 632_9C | 2013 | UK | Pig Healthy | Non-Clinical | NA | 25 | [36] | PRJEB6250 |
| 635_10C | 2013 | UK | Pig Healthy | Non-Clinical | NA | 10 | [36] | PRJEB6250 |
| 635_11B | 2013 | UK | Pig Healthy | Non-Clinical | NA | 15 | [36] | PRJEB6250 |
| 635_3A | 2013 | UK | Pig Healthy | Non-Clinical | NA | 16 | [36] | PRJEB6250 |
| 635_6B | 2013 | UK | Pig Healthy | Non-Clinical | NA | 16 | [36] | PRJEB6250 |
| 635_6C | 2013 | UK | Pig Healthy | Non-Clinical | NA | 15 | [36] | PRJEB6250 |
| 635_8C | 2013 | UK | Pig Healthy | Non-Clinical | NA | 16 | [36] | PRJEB6250 |
| 635_9A | 2013 | UK | Pig Healthy | Non-Clinical | NA | NA | [36] | PRJEB6250 |
| 635_9C | 2013 | UK | Pig Healthy | Non-Clinical | NA | 15 | [36] | PRJEB6250 |
| 684_10A | 2013 | UK | Pig Healthy | Non-Clinical | NA | 5 | [36] | PRJEB6250 |
| 684_11B | 2013 | UK | Pig Healthy | Non-Clinical | NA | 28 | [36] | PRJEB6250 |
| 684_11C | 2013 | UK | Pig Healthy | Non-Clinical | NA | 25 | [36] | PRJEB6250 |
| 684_12A | 2013 | UK | Pig Healthy | Non-Clinical | NA | 25 | [36] | PRJEB6250 |
| 684_12B | 2013 | UK | Pig Healthy | Non-Clinical | NA | 16 | [36] | PRJEB6250 |

| | | | | | | | | |
|---|---|---|---|---|---|---|---|---|
| 684_12C | 2013 | UK | Pig Healthy | Non-Clinical | NA | NA | [36] | PRJEB6250 |
| 684_13A | 2013 | UK | Pig Healthy | Non-Clinical | NA | 16 | [36] | PRJEB6250 |
| 684_13C | 2013 | UK | Pig Healthy | Non-Clinical | NA | 16 | [36] | PRJEB6250 |
| 684_14B | 2013 | UK | Pig Healthy | Non-Clinical | NA | 9 | [36] | PRJEB6250 |
| 684_14C | 2013 | UK | Pig Healthy | Non-Clinical | NA | 9 | [36] | PRJEB6250 |
| 684_15B | 2013 | UK | Pig Healthy | Non-Clinical | NA | 21 | [36] | PRJEB6250 |
| 684_16A | 2013 | UK | Pig Healthy | Non-Clinical | NA | 9 | [36] | PRJEB6250 |
| 684_17A | 2013 | UK | Pig Healthy | Non-Clinical | NA | 15 | [36] | PRJEB6250 |
| 684_17C | 2013 | UK | Pig Healthy | Non-Clinical | NA | 15 | [36] | PRJEB6250 |
| 684_18B | 2013 | UK | Pig Healthy | Non-Clinical | NA | NA | [36] | PRJEB6250 |
| 684_18C | 2013 | UK | Pig Healthy | Non-Clinical | NA | NA | [36] | PRJEB6250 |
| 684_19B | 2013 | UK | Pig Healthy | Non-Clinical | NA | 30 | [36] | PRJEB6250 |
| 684_1B | 2013 | UK | Pig Healthy | Non-Clinical | NA | 8 | [36] | PRJEB6250 |
| 684_20A | 2013 | UK | Pig Healthy | Non-Clinical | NA | 16 | [36] | PRJEB6250 |
| 684_20C | 2013 | UK | Pig Healthy | Non-Clinical | NA | 9 | [36] | PRJEB6250 |
| 684_21A | 2013 | UK | Pig Healthy | Non-Clinical | NA | 16 | [36] | PRJEB6250 |
| 684_21B_ref | 2013 | UK | Pig Healthy | Non-Clinical | NA | NA | [36] | PRJEB6250 |
| 684_21C | 2013 | UK | Pig Healthy | Non-Clinical | NA | 16 | [36] | PRJEB6250 |
| 684_24C | 2013 | UK | Pig Healthy | Non-Clinical | NA | NA | [36] | PRJEB6250 |
| 684_25C | 2013 | UK | Pig Healthy | Non-Clinical | NA | 15 | [36] | PRJEB6250 |
| 684_2A | 2013 | UK | Pig Healthy | Non-Clinical | NA | 25 | [36] | PRJEB6250 |
| 684_3A | 2013 | UK | Pig Healthy | Non-Clinical | NA | 15 | [36] | PRJEB6250 |
| 684_3B | 2013 | UK | Pig Healthy | Non-Clinical | NA | NA | [36] | PRJEB6250 |
| 684_3C | 2013 | UK | Pig Healthy | Non-Clinical | NA | 28 | [36] | PRJEB6250 |
| 684_4A | 2013 | UK | Pig Healthy | Non-Clinical | NA | 16 | [36] | PRJEB6250 |
| 684_5A | 2013 | UK | Pig Healthy | Non-Clinical | NA | 16 | [36] | PRJEB6250 |

| | | | | | | | | |
|---|---|---|---|---|---|---|---|---|
| 684_5B | 2013 | UK | Pig Healthy | Non-Clinical | NA | 28 | [36] | PRJEB6250 |
| 684_6A | 2013 | UK | Pig Healthy | Non-Clinical | NA | 15 | [36] | PRJEB6250 |
| 684_6B | 2013 | UK | Pig Healthy | Non-Clinical | NA | 16 | [36] | PRJEB6250 |
| 684_6C | 2013 | UK | Pig Healthy | Non-Clinical | NA | 9 | [36] | PRJEB6250 |
| 684_7A | 2013 | UK | Pig Healthy | Non-Clinical | NA | 25 | [36] | PRJEB6250 |
| 684_7C | 2013 | UK | Pig Healthy | Non-Clinical | NA | 25 | [36] | PRJEB6250 |
| 684_8A | 2013 | UK | Pig Healthy | Non-Clinical | NA | 16 | [36] | PRJEB6250 |
| 684_8C | 2013 | UK | Pig Healthy | Non-Clinical | NA | 16 | [36] | PRJEB6250 |
| 684_9B | 2013 | UK | Pig Healthy | Non-Clinical | NA | 9 | [36] | PRJEB6250 |
| 714_14B | 2014 | UK | Pig Healthy | Non-Clinical | NA | 28 | [36] | PRJEB6250 |
| 714_15A | 2014 | UK | Pig Healthy | Non-Clinical | 29 | 7 | [36] | PRJEB6250 |
| 714_19B | 2014 | UK | Pig Healthy | Non-Clinical | NA | NA | [36] | PRJEB6250 |
| 714_3A | 2014 | UK | Pig Healthy | Non-Clinical | NA | NA | [36] | PRJEB6250 |
| 714_7B | 2014 | UK | Pig Healthy | Non-Clinical | NA | NA | [36] | PRJEB6250 |
| 784_10A | 2013 | UK | Pig Healthy | Non-Clinical | NA | NA | [36] | PRJEB6250 |
| 784_10B | 2013 | UK | Pig Healthy | Non-Clinical | NA | 6 | [36] | PRJEB6250 |
| 784_15B | 2013 | UK | Pig Healthy | Non-Clinical | NA | NA | [36] | PRJEB6250 |
| 784_16B | 2013 | UK | Pig Healthy | Non-Clinical | NA | NA | [36] | PRJEB6250 |
| 784_17C | 2013 | UK | Pig Healthy | Non-Clinical | NA | NA | [36] | PRJEB6250 |
| 784_18C | 2013 | UK | Pig Healthy | Non-Clinical | NA | 31 | [36] | PRJEB6250 |
| 784_20A | 2013 | UK | Pig Healthy | Non-Clinical | NA | 6 | [36] | PRJEB6250 |
| 784_21A | 2013 | UK | Pig Healthy | Non-Clinical | NA | NA | [36] | PRJEB6250 |
| 784_23B | 2013 | UK | Pig Healthy | Non-Clinical | NA | 21 | [36] | PRJEB6250 |
| 784_23C | 2013 | UK | Pig Healthy | Non-Clinical | NA | NA | [36] | PRJEB6250 |
| 784_24A | 2013 | UK | Pig Healthy | Non-Clinical | NA | NA | [36] | PRJEB6250 |
| 784_2C | 2013 | UK | Pig Healthy | Non-Clinical | NA | 6 | [36] | PRJEB6250 |

| | | | | | | | | |
|---|---|---|---|---|---|---|---|---|
| 784_3A | 2013 | UK | Pig Healthy | Non-Clinical | NA | NA | [36] | PRJEB6250 |
| 784_5C | 2013 | UK | Pig Healthy | Non-Clinical | NA | 16 | [36] | PRJEB6250 |
| 784_6A | 2013 | UK | Pig Healthy | Non-Clinical | NA | 31 | [36] | PRJEB6250 |
| 784_7A | 2013 | UK | Pig Healthy | Non-Clinical | NA | 16 | [36] | PRJEB6250 |
| 784_7B | 2013 | UK | Pig Healthy | Non-Clinical | NA | 9 | [36] | PRJEB6250 |
| 784_8B | 2013 | UK | Pig Healthy | Non-Clinical | NA | 16 | [36] | PRJEB6250 |
| 784_9A | 2013 | UK | Pig Healthy | Non-Clinical | NA | 9 | [36] | PRJEB6250 |
| 876_15B | 2013 | UK | Pig Healthy | Non-Clinical | NA | 9 | [36] | PRJEB6250 |
| 876_15C | 2013 | UK | Pig Healthy | Non-Clinical | NA | 9 | [36] | PRJEB6250 |
| 876_19A | 2013 | UK | Pig Healthy | Non-Clinical | NA | 8 | [36] | PRJEB6250 |
| 876_1A | 2013 | UK | Pig Healthy | Non-Clinical | NA | 15 | [36] | PRJEB6250 |
| 876_20A | 2013 | UK | Pig Healthy | Non-Clinical | NA | 9 | [36] | PRJEB6250 |
| 876_21B | 2013 | UK | Pig Healthy | Non-Clinical | NA | 30 | [36] | PRJEB6250 |
| 876_2A | 2013 | UK | Pig Healthy | Non-Clinical | NA | 9 | [36] | PRJEB6250 |
| 876_3B | 2013 | UK | Pig Healthy | Non-Clinical | NA | 29 | [36] | PRJEB6250 |
| 876_6C | 2013 | UK | Pig Healthy | Non-Clinical | NA | 10 | [36] | PRJEB6250 |
| TMW_SS001 | NA | UK | Pig Unknown | Unknown | 1 | 2 | [36] | PRJNA628943 |
| TMW_SS002 | NA | UK | Pig Unknown | Unknown | NA | 2 | [36] | PRJNA628943 |
| TMW_SS003 | NA | UK | Pig Unknown | Unknown | NA | NA | [36] | PRJNA628943 |
| TMW_SS004 | NA | UK | Pig Unknown | Unknown | 87 | 8 | [36] | PRJNA628943 |
| TMW_SS005 | 2013 | UK | Pig Diseased | Respiratory | NA | 2 | [36] | PRJNA628943 |
| TMW_SS006 | 2013 | UK | Pig Unknown | Unknown | 1 | 2 | [36] | PRJNA628943 |
| TMW_SS007 | 2013 | UK | Pig Diseased | Systemic | 1 | 2 | [36] | PRJNA628943 |
| TMW_SS008 | 2013 | UK | Pig Unknown | Unknown | NA | 9 | [36] | PRJNA628943 |
| TMW_SS009 | 2013 | UK | Pig Diseased | Respiratory | NA | 2 | [36] | PRJNA628943 |
| TMW_SS010 | 2013 | UK | Pig Diseased | Respiratory | 1 | 2 | [36] | PRJNA628943 |

| | | | | | | | | |
|---|---|---|---|---|---|---|---|---|
| TMW_SS011 | 2013 | UK | Pig Diseased | Respiratory | NA | 14 | [36] | PRJNA628943 |
| TMW_SS012 | 2013 | UK | Pig Unknown | Unknown | NA | 15 | [36] | PRJNA628943 |
| TMW_SS013 | 2013 | UK | Pig Unknown | Unknown | 29 | NA | [36] | PRJNA628943 |
| TMW_SS014 | 2013 | UK | Pig Unknown | Unknown | NA | NA | [36] | PRJNA628943 |
| TMW_SS015 | 2013 | UK | Pig Unknown | Unknown | NA | 2 | [36] | PRJNA628943 |
| TMW_SS016 | 2013 | UK | Pig Unknown | Unknown | 1 | 2 | [36] | PRJNA628943 |
| TMW_SS017 | 2013 | UK | Pig Unknown | Unknown | 1 | 14 | [36] | PRJNA628943 |
| TMW_SS018 | 2013 | UK | Pig Diseased | Respiratory | NA | 8 | [36] | PRJNA628943 |
| TMW_SS019 | 2013 | UK | Pig Unknown | Unknown | 1 | 14 | [36] | PRJNA628943 |
| TMW_SS020 | 2013 | UK | Pig Unknown | Unknown | NA | NA | [36] | PRJNA628943 |
| TMW_SS021 | 2013 | UK | Pig Unknown | Unknown | NA | 29 | [36] | PRJNA628943 |
| TMW_SS022 | 2013 | UK | Pig Diseased | Systemic | 1 | 2 | [36] | PRJNA628943 |
| TMW_SS023 | 2013 | UK | Pig Unknown | Unknown | 1 | 14 | [36] | PRJNA628943 |
| TMW_SS024 | 2013 | UK | Pig Unknown | Unknown | NA | NA | [36] | PRJNA628943 |
| TMW_SS025 | NA | UK | Pig Unknown | Unknown | 1 | 2 | [36] | PRJNA628943 |
| TMW_SS026 | 2013 | UK | Pig Unknown | Unknown | 27 | 3 | [36] | PRJNA628943 |
| TMW_SS027 | 2013 | UK | Pig Unknown | Unknown | NA | 8 | [36] | PRJNA628943 |
| TMW_SS028_ref | 2013 | UK | Pig Unknown | Unknown | NA | 29 | [36] | PRJNA1010137 |
| TMW_SS029 | 2013 | UK | Pig Unknown | Unknown | NA | 15 | [36] | PRJNA628943 |
| TMW_SS031 | 2013 | UK | Pig Unknown | Unknown | 1 | 14 | [36] | PRJNA628943 |
| TMW_SS032 | 2013 | UK | Pig Unknown | Unknown | 31 | 3 | [36] | PRJNA628943 |
| TMW_SS033 | 2013 | UK | Pig Unknown | Unknown | 28 | 2 | [36] | PRJNA628943 |
| TMW_SS034 | 2013 | UK | Pig Unknown | Unknown | NA | NA | [36] | PRJNA628943 |
| TMW_SS035 | 2013 | UK | Pig Unknown | Unknown | NA | NA | [36] | PRJNA628943 |
| TMW_SS036 | 2013 | UK | Pig Unknown | Unknown | NA | 2 | [36] | PRJNA628943 |
| TMW_SS037 | 2013 | UK | Pig Unknown | Unknown | NA | NA | [36] | PRJNA628943 |

| | | | | | | | | |
|---|---|---|---|---|---|---|---|---|
| TMW_SS038 | 2013 | UK | Pig Unknown | Unknown | 1 | 2 | [36] | PRJNA628943 |
| TMW_SS039 | 2013 | UK | Pig Unknown | Unknown | 1 | 2 | [36] | PRJNA628943 |
| TMW_SS040 | 2013 | UK | Pig Diseased | Systemic | 1 | 2 | [36] | PRJNA628943 |
| TMW_SS041 | 2013 | UK | Pig Unknown | Unknown | NA | 3 | [36] | PRJNA628943 |
| TMW_SS042 | 2013 | UK | Pig Unknown | Unknown | NA | 4 | [36] | PRJNA628943 |
| TMW_SS043 | 2014 | UK | Pig Diseased | Respiratory | NA | 2 | [36] | PRJNA628943 |
| TMW_SS044 | 2014 | UK | Pig Unknown | Unknown | NA | 2 | [36] | PRJNA628943 |
| TMW_SS045 | 2014 | UK | Pig Unknown | Unknown | NA | NA | [36] | PRJNA628943 |
| TMW_SS046 | 2014 | UK | Pig Unknown | Unknown | NA | 2 | [36] | PRJNA628943 |
| TMW_SS047 | NA | UK | Pig Unknown | Unknown | 28 | 2 | [36] | PRJNA628943 |
| TMW_SS048 | 2014 | UK | Pig Diseased | Respiratory | 28 | 2 | [36] | PRJNA628943 |
| TMW_SS049 | 2014 | UK | Pig Unknown | Unknown | NA | 2 | [36] | PRJNA628943 |
| TMW_SS051 | 2014 | UK | Pig Unknown | Unknown | 29 | NA | [36] | PRJNA628943 |
| TMW_SS052 | 2014 | UK | Pig Unknown | Unknown | 29 | NA | [36] | PRJNA628943 |
| TMW_SS053 | 2014 | UK | Pig Unknown | Unknown | 28 | 2 | [36] | PRJNA628943 |
| TMW_SS054 | 2014 | UK | Pig Unknown | Unknown | NA | 28 | [36] | PRJNA628943 |
| TMW_SS055 | 2014 | UK | Pig Unknown | Unknown | 2 | 2 | [36] | PRJNA628943 |
| TMW_SS056 | 2014 | UK | Pig Unknown | Unknown | 352 | 2 | [36] | PRJNA628943 |
| TMW_SS057 | 2014 | UK | Pig Unknown | Unknown | 28 | 2 | [36] | PRJNA628943 |
| TMW_SS058 | NA | UK | Pig Unknown | Unknown | 1 | 2 | [36] | PRJNA628943 |
| TMW_SS059 | 2014 | UK | Pig Diseased | Systemic | 1 | 14 | [36] | PRJNA628943 |
| TMW_SS060 | 2014 | UK | Pig Diseased | Systemic | 1 | 2 | [36] | PRJNA628943 |
| TMW_SS061 | 2014 | UK | Pig Unknown | Unknown | 1 | 2 | [36] | PRJNA628943 |
| TMW_SS062 | 2014 | UK | Pig Unknown | Unknown | 146 | 2 | [36] | PRJNA628943 |
| TMW_SS063 | 2014 | UK | Pig Unknown | Unknown | 87 | 8 | [36] | PRJNA628943 |
| TMW_SS064 | 2014 | UK | Pig Unknown | Unknown | 87 | 8 | [36] | PRJNA628943 |

| | | | | | | | | |
|---|---|---|---|---|---|---|---|---|
| TMW_SS065 | 2014 | UK | Pig Unknown | Unknown | 87 | NA | [36] | PRJNA628943 |
| TMW_SS066 | 2014 | UK | Pig Unknown | Unknown | NA | 18 | [36] | PRJNA628943 |
| TMW_SS067 | 2014 | UK | Pig Unknown | Unknown | NA | NA | [36] | PRJNA628943 |
| TMW_SS069 | 2014 | UK | Pig Unknown | Unknown | NA | 2 | [36] | PRJNA628943 |
| TMW_SS070 | 2014 | UK | Pig Diseased | Respiratory | 94 | 4 | [36] | PRJNA628943 |
| TMW_SS071 | 2014 | UK | Pig Diseased | Systemic | NA | 2 | [36] | PRJNA628943 |
| TMW_SS072 | 2014 | UK | Pig Unknown | Unknown | NA | 7 | [36] | PRJNA628943 |
| TMW_SS073 | 2014 | UK | Pig Diseased | Respiratory | 87 | 8 | [36] | PRJNA628943 |
| TMW_SS074 | 2014 | UK | Pig Diseased | Respiratory | 87 | NA | [36] | PRJNA628943 |
| TMW_SS075 | 2014 | UK | Pig Unknown | Unknown | NA | NA | [36] | PRJNA628943 |
| TMW_SS076 | 2014 | UK | Pig Diseased | Respiratory | NA | 8 | [36] | PRJNA628943 |
| TMW_SS077 | 2014 | UK | Pig Unknown | Unknown | NA | 14 | [36] | PRJNA628943 |
| TMW_SS078 | 2014 | UK | Pig Unknown | Unknown | NA | NA | [36] | PRJNA628943 |
| TMW_SS079 | 2014 | UK | Pig Diseased | Respiratory | NA | 2 | [36] | PRJNA628943 |
| TMW_SS080 | 2014 | UK | Pig Unknown | Unknown | 29 | NA | [36] | PRJNA628943 |
| TMW_SS081 | 2014 | UK | Pig Diseased | Systemic | 1 | 2 | [36] | PRJNA628943 |
| TMW_SS082 | 2014 | UK | Pig Diseased | Respiratory | 87 | 8 | [36] | PRJNA628943 |
| TMW_SS083 | 2014 | UK | Pig Diseased | Systemic | 1 | 2 | [36] | PRJNA628943 |
| TMW_SS084 | NA | UK | Pig Unknown | Unknown | 28 | 2 | [36] | PRJNA628943 |
| TMW_SS085 | 2014 | UK | Pig Unknown | Unknown | 28 | 2 | [36] | PRJNA628943 |
| TMW_SS086 | 2014 | UK | Pig Diseased | Systemic | 13 | 14 | [36] | PRJNA628943 |
| TMW_SS087 | 2014 | UK | Pig Unknown | Unknown | NA | NA | [36] | PRJNA628943 |
| TMW_SS089 | 2014 | UK | Pig Unknown | Unknown | 1 | 2 | [36] | PRJNA628943 |
| TMW_SS090 | 2014 | UK | Pig Diseased | Systemic | 1 | 14 | [36] | PRJNA628943 |
| TMW_SS091 | 2014 | UK | Pig Unknown | Unknown | NA | 29 | [36] | PRJNA628943 |
| TMW_SS092 | 2014 | UK | Pig Unknown | Unknown | 29 | NA | [36] | PRJNA628943 |

| | | | | | | | | |
|---|---|---|---|---|---|---|---|---|
| TMW_SS093 | 2014 | UK | Pig Unknown | Unknown | 1 | 2 | [36] | PRJNA628943 |
| TMW_SS094 | 2014 | UK | Pig Unknown | Unknown | 87 | 8 | [36] | PRJNA628943 |
| TMW_SS095 | 2014 | UK | Pig Unknown | Unknown | 28 | 2 | [36] | PRJNA628943 |
| TMW_SS096 | 2014 | UK | Pig Diseased | Systemic | 1 | 2 | [36] | PRJNA628943 |
| TMW_SS097 | 2014 | UK | Pig Unknown | Unknown | 1 | 14 | [36] | PRJNA628943 |
| TMW_SS098 | NA | UK | Pig Unknown | Unknown | 28 | 2 | [36] | PRJNA628943 |
| TMW_SS099 | 2014 | UK | Pig Unknown | Unknown | 1 | 2 | [36] | PRJNA628943 |
| TMW_SS100 | 2014 | UK | Pig Diseased | Respiratory | 28 | 2 | [36] | PRJNA628943 |
| TMW_SS101 | 2014 | UK | Pig Unknown | Unknown | 28 | 2 | [36] | PRJNA628943 |
| TMW_SS102 | 2014 | UK | Pig Unknown | Unknown | 1 | 2 | [36] | PRJNA628943 |
| TMW_SS103 | 2014 | UK | Pig Unknown | Unknown | 1 | 14 | [36] | PRJNA628943 |
| TMW_SS104 | 2014 | UK | Pig Diseased | Systemic | NA | 2 | [36] | PRJNA628943 |
| TMW_SS105 | 2014 | UK | Pig Unknown | Unknown | 28 | 2 | [36] | PRJNA628943 |
| TMW_SS106 | 2015 | UK | Pig Healthy | Non-Clinical | NA | 28 | [36] | PRJNA628943 |
| TMW_SS107 | 2015 | UK | Pig Healthy | Non-Clinical | NA | 28 | [36] | PRJNA628943 |
| TMW_SS108 | 2015 | UK | Pig Healthy | Non-Clinical | NA | 15 | [36] | PRJNA628943 |
| TMW_SS109 | 2015 | UK | Pig Healthy | Non-Clinical | NA | 15 | [36] | PRJNA628943 |
| TMW_SS110 | 2015 | UK | Pig Healthy | Non-Clinical | NA | 28 | [36] | PRJNA628943 |
| TMW_SS111 | 2015 | UK | Pig Healthy | Non-Clinical | NA | 28 | [36] | PRJNA628943 |
| TMW_SS112 | 2015 | UK | Pig Healthy | Non-Clinical | NA | 28 | [36] | PRJNA628943 |
| TMW_SS113 | 2015 | UK | Pig Healthy | Non-Clinical | NA | 28 | [36] | PRJNA628943 |
| TMW_SS114 | 2015 | UK | Pig Healthy | Non-Clinical | NA | 28 | [36] | PRJNA628943 |
| TMW_SS115 | 2015 | UK | Pig Healthy | Non-Clinical | NA | 28 | [36] | PRJNA628943 |
| TMW_SS116 | 2015 | UK | Pig Healthy | Non-Clinical | NA | 15 | [36] | PRJNA628943 |
| TMW_SS117 | 2015 | UK | Pig Healthy | Non-Clinical | NA | 15 | [36] | PRJNA628943 |
| TMW_SS118 | 2015 | UK | Pig Healthy | Non-Clinical | NA | 28 | [36] | PRJNA628943 |

| | | | | | | | | |
|---|---|---|---|---|---|---|---|---|
| TMW_SS119 | 2015 | UK | Pig Healthy | Non-Clinical | NA | 15 | [36] | PRJNA628943 |
| SS_0013 | 2013 | China | Pig Healthy | Non-Clinical | NA | 16 | [37] | PRJNA428542 |
| SS_0025 | 2013 | China | Pig Healthy | Non-Clinical | NA | 16 | [37] | PRJNA428542 |
| SS_0030 | 2013 | China | Pig Healthy | Non-Clinical | NA | 16 | [37] | PRJNA428542 |
| SS_0031 | 2013 | China | Pig Healthy | Non-Clinical | NA | 16 | [37] | PRJNA428542 |
| SS_0036 | 2013 | China | Pig Healthy | Non-Clinical | NA | 16 | [37] | PRJNA428542 |
| SS_0052 | 2013 | China | Pig Healthy | Non-Clinical | NA | 16 | [37] | PRJNA428542 |
| SS_0053 | 2013 | China | Pig Healthy | Non-Clinical | NA | 16 | [37] | PRJNA428542 |
| SS_0061 | 2013 | China | Pig Healthy | Non-Clinical | NA | 31 | [37] | PRJNA428542 |
| SS_0073 | 2013 | China | Pig Healthy | Non-Clinical | NA | 16 | [37] | PRJNA428542 |
| SS_0077 | 2013 | China | Pig Healthy | Non-Clinical | NA | 16 | [37] | PRJNA428542 |
| SS_0081 | 2013 | China | Pig Healthy | Non-Clinical | NA | 16 | [37] | PRJNA428542 |
| SS_0082 | 2013 | China | Pig Healthy | Non-Clinical | NA | 31 | [37] | PRJNA428542 |
| SS_0083 | 2013 | China | Pig Healthy | Non-Clinical | NA | 16 | [37] | PRJNA428542 |
| SS_0111 | 2013 | China | Pig Healthy | Non-Clinical | NA | 16 | [37] | PRJNA428542 |
| SS_0123 | 2013 | China | Pig Healthy | Non-Clinical | NA | 12 | [37] | PRJNA428542 |
| SS_0138 | 2013 | China | Pig Healthy | Non-Clinical | NA | 15 | [37] | PRJNA428542 |
| SS_0139 | 2013 | China | Pig Healthy | Non-Clinical | NA | 16 | [37] | PRJNA428542 |
| SS_0142 | 2013 | China | Pig Healthy | Non-Clinical | NA | 16 | [37] | PRJNA428542 |
| SS_0143 | 2013 | China | Pig Healthy | Non-Clinical | NA | 16 | [37] | PRJNA428542 |
| SS_0148 | 2013 | China | Pig Healthy | Non-Clinical | NA | 9 | [37] | PRJNA428542 |
| SS_0166 | 2013 | China | Pig Healthy | Non-Clinical | NA | NA | [37] | PRJNA428542 |
| SS_0179 | 2013 | China | Pig Healthy | Non-Clinical | 117 | 3 | [37] | PRJNA428542 |
| SS_0180 | 2013 | China | Pig Healthy | Non-Clinical | 499 | 5 | [37] | PRJNA428542 |
| SS_0183 | 2013 | China | Pig Healthy | Non-Clinical | 250 | 16 | [37] | PRJNA428542 |
| SS_0189 | 2013 | China | Pig Healthy | Non-Clinical | NA | NA | [37] | PRJNA428542 |

| | | | | | | | | |
|---|---|---|---|---|---|---|---|---|
| SS_0191 | 2013 | China | Pig Healthy | Non-Clinical | 117 | 3 | [37] | PRJNA428542 |
| SS_0192 | 2013 | China | Pig Healthy | Non-Clinical | 117 | 3 | [37] | PRJNA428542 |
| SS_0207 | 2013 | China | Pig Healthy | Non-Clinical | 117 | 3 | [37] | PRJNA428542 |
| SS_0217 | 2013 | China | Pig Healthy | Non-Clinical | 250 | 16 | [37] | PRJNA428542 |
| SS_0262 | 2013 | China | Pig Healthy | Non-Clinical | NA | 16 | [37] | PRJNA428542 |
| SS_0324 | 2013 | China | Pig Healthy | Non-Clinical | NA | 16 | [37] | PRJNA428542 |
| SS_0326 | 2013 | China | Pig Healthy | Non-Clinical | NA | 16 | [37] | PRJNA428542 |
| SS_0347 | 2013 | China | Pig Healthy | Non-Clinical | NA | 25 | [37] | PRJNA428542 |
| SS_0350 | 2013 | China | Pig Healthy | Non-Clinical | NA | 25 | [37] | PRJNA428542 |
| SS_0355 | 2013 | China | Pig Healthy | Non-Clinical | NA | 16 | [37] | PRJNA428542 |
| SS_0356 | 2013 | China | Pig Healthy | Non-Clinical | NA | 16 | [37] | PRJNA428542 |
| SS_0378 | 2013 | China | Pig Healthy | Non-Clinical | NA | 15 | [37] | PRJNA428542 |
| SS_0393 | 2013 | China | Pig Healthy | Non-Clinical | NA | 12 | [37] | PRJNA428542 |
| SS_0424-1 | 2013 | China | Pig Healthy | Non-Clinical | 477 | 16 | [37] | PRJNA428542 |
| SS_0615 | 2013 | China | Pig Healthy | Non-Clinical | NA | NA | [37] | PRJNA428542 |
| SS_0629 | 2013 | China | Pig Healthy | Non-Clinical | NA | NA | [37] | PRJNA428542 |
| SS_0635 | 2013 | China | Pig Healthy | Non-Clinical | NA | NA | [37] | PRJNA428542 |
| SS_0639 | 2013 | China | Pig Healthy | Non-Clinical | NA | NA | [37] | PRJNA428542 |
| SS_0649 | 2013 | China | Pig Healthy | Non-Clinical | NA | NA | [37] | PRJNA428542 |
| SS_0667 | 2013 | China | Pig Healthy | Non-Clinical | NA | 15 | [37] | PRJNA428542 |
| SS_0676 | 2013 | China | Pig Healthy | Non-Clinical | NA | NA | [37] | PRJNA428542 |
| SS_0681 | 2013 | China | Pig Healthy | Non-Clinical | NA | 30 | [37] | PRJNA428542 |
| SS_0707 | 2013 | China | Pig Healthy | Non-Clinical | NA | NA | [37] | PRJNA428542 |
| SS_0712 | 2013 | China | Pig Healthy | Non-Clinical | NA | NA | [37] | PRJNA428542 |
| SS_0768 | 2013 | China | Pig Healthy | Non-Clinical | 188 | 16 | [37] | PRJNA428542 |
| SS_0786 | 2013 | China | Pig Healthy | Non-Clinical | NA | 16 | [37] | PRJNA428542 |

| | | | | | | | | |
|---|---|---|---|---|---|---|---|---|
| SS_0789 | 2013 | China | Pig Healthy | Non-Clinical | NA | 15 | [37] | PRJNA428542 |
| SS_0796 | 2013 | China | Pig Healthy | Non-Clinical | NA | NA | [37] | PRJNA428542 |
| SS_0797 | 2013 | China | Pig Healthy | Non-Clinical | NA | NA | [37] | PRJNA428542 |
| SS_0801 | 2013 | China | Pig Healthy | Non-Clinical | 188 | 16 | [37] | PRJNA428542 |
| SS_0803 | 2013 | China | Pig Healthy | Non-Clinical | NA | 16 | [37] | PRJNA428542 |
| SS_0805 | 2013 | China | Pig Healthy | Non-Clinical | NA | 5 | [37] | PRJNA428542 |
| SS_0810 | 2013 | China | Pig Healthy | Non-Clinical | NA | 28 | [37] | PRJNA428542 |
| SS_0811 | 2013 | China | Pig Healthy | Non-Clinical | NA | 28 | [37] | PRJNA428542 |
| SS_0812 | 2013 | China | Pig Healthy | Non-Clinical | 11 | 14 | [37] | PRJNA428542 |
| SS_0832 | 2013 | China | Pig Healthy | Non-Clinical | 11 | 14 | [37] | PRJNA428542 |
| SS_0849 | 2013 | China | Pig Healthy | Non-Clinical | NA | 15 | [37] | PRJNA428542 |
| SS_0857 | 2013 | China | Pig Healthy | Non-Clinical | 198 | 8 | [37] | PRJNA428542 |
| SS_0858 | 2013 | China | Pig Healthy | Non-Clinical | 198 | 8 | [37] | PRJNA428542 |
| SS_0859 | 2013 | China | Pig Healthy | Non-Clinical | 198 | 8 | [37] | PRJNA428542 |
| SS_0862 | 2013 | China | Pig Healthy | Non-Clinical | 198 | 8 | [37] | PRJNA428542 |
| SS_0863 | 2013 | China | Pig Healthy | Non-Clinical | 198 | 8 | [37] | PRJNA428542 |
| SS_0869 | 2013 | China | Pig Healthy | Non-Clinical | 198 | 8 | [37] | PRJNA428542 |
| SS_0889 | 2013 | China | Pig Healthy | Non-Clinical | 198 | 8 | [37] | PRJNA428542 |
| SS_0890 | 2013 | China | Pig Healthy | Non-Clinical | 198 | 8 | [37] | PRJNA428542 |
| SS_0893 | 2013 | China | Pig Healthy | Non-Clinical | NA | 15 | [37] | PRJNA428542 |
| SS_0895 | 2013 | China | Pig Healthy | Non-Clinical | NA | 15 | [37] | PRJNA428542 |
| SS_0899 | 2013 | China | Pig Healthy | Non-Clinical | NA | 31 | [37] | PRJNA428542 |
| SS_0900 | 2013 | China | Pig Healthy | Non-Clinical | NA | 15 | [37] | PRJNA428542 |
| SS_0907 | 2013 | China | Pig Healthy | Non-Clinical | NA | 15 | [37] | PRJNA428542 |
| SS_0927 | 2013 | China | Pig Healthy | Non-Clinical | NA | 31 | [37] | PRJNA428542 |
| SS_0939 | 2013 | China | Pig Healthy | Non-Clinical | NA | 31 | [37] | PRJNA428542 |

| | | | | | | | | |
|---|---|---|---|---|---|---|---|---|
| SS_0942 | 2013 | China | Pig Healthy | Non-Clinical | 94 | 4 | [37] | PRJNA428542 |
| SS_0944 | 2013 | China | Pig Healthy | Non-Clinical | 94 | 4 | [37] | PRJNA428542 |
| SS_0949 | 2013 | China | Pig Healthy | Non-Clinical | NA | 29 | [37] | PRJNA428542 |
| SS_0955 | 2013 | China | Pig Healthy | Non-Clinical | NA | 29 | [37] | PRJNA428542 |
| SS_0957 | 2013 | China | Pig Healthy | Non-Clinical | 188 | 16 | [37] | PRJNA428542 |
| SS_0963 | 2013 | China | Pig Healthy | Non-Clinical | NA | NA | [37] | PRJNA428542 |
| SS_0964 | 2013 | China | Pig Healthy | Non-Clinical | NA | 29 | [37] | PRJNA428542 |
| SS_0965 | 2013 | China | Pig Healthy | Non-Clinical | 188 | 16 | [37] | PRJNA428542 |
| SS_0966 | 2013 | China | Pig Healthy | Non-Clinical | 188 | 16 | [37] | PRJNA428542 |
| SS_0969 | 2013 | China | Pig Healthy | Non-Clinical | NA | 29 | [37] | PRJNA428542 |
| SS_0970 | 2013 | China | Pig Healthy | Non-Clinical | NA | 30 | [37] | PRJNA428542 |
| SS_0985 | 2013 | China | Pig Healthy | Non-Clinical | NA | 29 | [37] | PRJNA428542 |
| SS_1008 | 2013 | China | Pig Healthy | Non-Clinical | NA | NA | [37] | PRJNA428542 |
| SS_1012 | 2013 | China | Pig Healthy | Non-Clinical | 188 | 16 | [37] | PRJNA428542 |
| SS_1024-1 | 2013 | China | Pig Healthy | Non-Clinical | 1 | 2 | [37] | PRJNA428542 |
| SS_1034 | 2013 | China | Pig Healthy | Non-Clinical | NA | 31 | [37] | PRJNA428542 |
| SS_1044 | 2013 | China | Pig Healthy | Non-Clinical | 94 | 4 | [37] | PRJNA428542 |
| SS_1047 | 2013 | China | Pig Healthy | Non-Clinical | 188 | 16 | [37] | PRJNA428542 |
| SS_1050 | 2013 | China | Pig Healthy | Non-Clinical | NA | 29 | [37] | PRJNA428542 |
| SS_1051 | 2013 | China | Pig Healthy | Non-Clinical | NA | 29 | [37] | PRJNA428542 |
| SS_1052 | 2013 | China | Pig Healthy | Non-Clinical | NA | 29 | [37] | PRJNA428542 |
| SS_1053 | 2013 | China | Pig Healthy | Non-Clinical | NA | 29 | [37] | PRJNA428542 |
| SS_1059 | 2013 | China | Pig Healthy | Non-Clinical | NA | 31 | [37] | PRJNA428542 |
| SS_1060 | 2013 | China | Pig Healthy | Non-Clinical | NA | 31 | [37] | PRJNA428542 |
| SS_1069 | 2013 | China | Pig Healthy | Non-Clinical | NA | 31 | [37] | PRJNA428542 |
| SS_1070 | 2013 | China | Pig Healthy | Non-Clinical | NA | 29 | [37] | PRJNA428542 |

| | | | | | | | | |
|---|---|---|---|---|---|---|---|---|
| SS_1076 | 2013 | China | Pig Healthy | Non-Clinical | NA | NA | [37] | PRJNA428542 |
| SS_1100 | 2013 | China | Pig Healthy | Non-Clinical | NA | NA | [37] | PRJNA428542 |
| SS_1140 | 2013 | China | Pig Healthy | Non-Clinical | NA | NA | [37] | PRJNA428542 |
| SS_1141-1 | 2013 | China | Pig Healthy | Non-Clinical | NA | NA | [37] | PRJNA428542 |
| SS_1164 | 2013 | China | Pig Healthy | Non-Clinical | NA | NA | [37] | PRJNA428542 |
| SS_1170 | 2013 | China | Pig Healthy | Non-Clinical | NA | NA | [37] | PRJNA428542 |
| SS_1171 | 2013 | China | Pig Healthy | Non-Clinical | NA | NA | [37] | PRJNA428542 |
| SS_1176 | 2013 | China | Pig Healthy | Non-Clinical | NA | 15 | [37] | PRJNA428542 |
| SS_1180 | 2013 | China | Pig Healthy | Non-Clinical | NA | 15 | [37] | PRJNA428542 |
| SS_1188 | 2013 | China | Pig Healthy | Non-Clinical | NA | NA | [37] | PRJNA428542 |
| SS_1194 | 2013 | China | Pig Healthy | Non-Clinical | NA | 15 | [37] | PRJNA428542 |
| SS_1200 | 2013 | China | Pig Healthy | Non-Clinical | NA | 16 | [37] | PRJNA428542 |
| SS_1214 | 2013 | China | Pig Healthy | Non-Clinical | NA | 16 | [37] | PRJNA428542 |
| SS_1215 | 2013 | China | Pig Healthy | Non-Clinical | NA | 16 | [37] | PRJNA428542 |
| SS_1219 | 2013 | China | Pig Healthy | Non-Clinical | NA | NA | [37] | PRJNA428542 |
| SS_1247 | 2013 | China | Pig Healthy | Non-Clinical | 477 | 16 | [37] | PRJNA428542 |
| SS_1251 | 2013 | China | Pig Healthy | Non-Clinical | 477 | 16 | [37] | PRJNA428542 |
| SS_1259 | 2013 | China | Pig Healthy | Non-Clinical | 477 | 16 | [37] | PRJNA428542 |
| SS_1261 | 2013 | China | Pig Healthy | Non-Clinical | NA | 11 | [37] | PRJNA428542 |
| SS_1262 | 2013 | China | Pig Healthy | Non-Clinical | 477 | 16 | [37] | PRJNA428542 |
| SS_1263 | 2013 | China | Pig Healthy | Non-Clinical | 477 | 16 | [37] | PRJNA428542 |
| SS_1264 | 2013 | China | Pig Healthy | Non-Clinical | 477 | 16 | [37] | PRJNA428542 |
| SS_1267 | 2013 | China | Pig Healthy | Non-Clinical | NA | 15 | [37] | PRJNA428542 |
| SS_1276 | 2013 | China | Pig Healthy | Non-Clinical | NA | 16 | [37] | PRJNA428542 |
| SS_1279 | 2013 | China | Pig Healthy | Non-Clinical | NA | 12 | [37] | PRJNA428542 |
| SS_1313 | 2013 | China | Pig Healthy | Non-Clinical | NA | 15 | [37] | PRJNA428542 |

| | | | | | | | | |
|---|---|---|---|---|---|---|---|---|
| SS_1314 | 2013 | China | Pig Healthy | Non-Clinical | 477 | 16 | [37] | PRJNA428542 |
| SS_1317 | 2013 | China | Pig Healthy | Non-Clinical | 477 | 16 | [37] | PRJNA428542 |
| SS_1318 | 2013 | China | Pig Healthy | Non-Clinical | 477 | 16 | [37] | PRJNA428542 |
| SS_1324 | 2013 | China | Pig Healthy | Non-Clinical | NA | 16 | [37] | PRJNA428542 |
| SS_1342 | 2013 | China | Pig Healthy | Non-Clinical | NA | 11 | [37] | PRJNA428542 |
| SS_1356 | 2013 | China | Pig Healthy | Non-Clinical | 423 | NA | [37] | PRJNA428542 |
| SS_1358 | 2013 | China | Pig Healthy | Non-Clinical | 7 | 2 | [37] | PRJNA428542 |
| SS_1360 | 2013 | China | Pig Healthy | Non-Clinical | 423 | 4 | [37] | PRJNA428542 |
| SS_1367 | 2013 | China | Pig Healthy | Non-Clinical | NA | 4 | [37] | PRJNA428542 |
| SS_1369 | 2013 | China | Pig Healthy | Non-Clinical | NA | 5 | [37] | PRJNA428542 |
| SS_1380 | 2013 | China | Pig Healthy | Non-Clinical | NA | 12 | [37] | PRJNA428542 |
| SS_1381 | 2013 | China | Pig Healthy | Non-Clinical | NA | 12 | [37] | PRJNA428542 |
| SS_1383 | 2013 | China | Pig Healthy | Non-Clinical | NA | 12 | [37] | PRJNA428542 |
| SS_1416 | 2013 | China | Pig Healthy | Non-Clinical | NA | NA | [37] | PRJNA428542 |
| SS_1496 | 2013 | China | Pig Healthy | Non-Clinical | NA | 21 | [37] | PRJNA428542 |
| SS_1721 | 2013 | China | Pig Healthy | Non-Clinical | NA | 31 | [37] | PRJNA428542 |
| SS_1722 | 2013 | China | Pig Healthy | Non-Clinical | NA | 31 | [37] | PRJNA428542 |
| SS_1781 | 2013 | China | Pig Healthy | Non-Clinical | 188 | 16 | [37] | PRJNA428542 |
| SS_1782 | 2013 | China | Pig Healthy | Non-Clinical | 188 | 16 | [37] | PRJNA428542 |
| SS_1783 | 2013 | China | Pig Healthy | Non-Clinical | 188 | 16 | [37] | PRJNA428542 |
| SS_1961 | 2013 | China | Pig Healthy | Non-Clinical | NA | 31 | [37] | PRJNA428542 |
| SS_1982 | 2013 | China | Pig Healthy | Non-Clinical | NA | 28 | [37] | PRJNA428542 |
| SS_2010 | 2013 | China | Pig Healthy | Non-Clinical | NA | 28 | [37] | PRJNA428542 |
| SS_2039 | 2013 | China | Pig Healthy | Non-Clinical | NA | 16 | [37] | PRJNA428542 |
| SS_2041 | 2013 | China | Pig Healthy | Non-Clinical | NA | 28 | [37] | PRJNA428542 |
| SS_2049-1 | 2013 | China | Pig Healthy | Non-Clinical | NA | 16 | [37] | PRJNA428542 |

| | | | | | | | | |
|---|---|---|---|---|---|---|---|---|
| SS_2052 | 2013 | China | Pig Healthy | Non-Clinical | NA | 30 | [37] | PRJNA428542 |
| SS_2070 | 2013 | China | Pig Healthy | Non-Clinical | NA | 28 | [37] | PRJNA428542 |
| SS_2080 | 2013 | China | Pig Healthy | Non-Clinical | NA | 29 | [37] | PRJNA428542 |
| SS_2089 | 2013 | China | Pig Healthy | Non-Clinical | NA | 15 | [37] | PRJNA428542 |
| SS_2100 | 2013 | China | Pig Healthy | Non-Clinical | NA | 29 | [37] | PRJNA428542 |
| SS_2101 | 2013 | China | Pig Healthy | Non-Clinical | NA | 21 | [37] | PRJNA428542 |
| SS_2151 | 2013 | China | Pig Healthy | Non-Clinical | NA | NA | [37] | PRJNA428542 |
| SS_2169 | 2013 | China | Pig Healthy | Non-Clinical | NA | 12 | [37] | PRJNA428542 |
| SS_2203-1 | 2013 | China | Pig Healthy | Non-Clinical | NA | 29 | [37] | PRJNA428542 |
| SS_2214 | 2013 | China | Pig Healthy | Non-Clinical | NA | 31 | [37] | PRJNA428542 |
| SS_2215 | 2013 | China | Pig Healthy | Non-Clinical | NA | 16 | [37] | PRJNA428542 |
| SS_2233 | 2013 | China | Pig Healthy | Non-Clinical | NA | NA | [37] | PRJNA428542 |
| SS_2254 | 2013 | China | Pig Healthy | Non-Clinical | NA | 30 | [37] | PRJNA428542 |
| SS_2261 | 2013 | China | Pig Healthy | Non-Clinical | NA | 15 | [37] | PRJNA428542 |
| SS_2283 | 2013 | China | Pig Healthy | Non-Clinical | 7 | 2 | [37] | PRJNA428542 |
| SS_2355 | 2013 | China | Pig Healthy | Non-Clinical | 7 | 5 | [37] | PRJNA428542 |
| SS_2440 | 2013 | China | Pig Healthy | Non-Clinical | NA | 16 | [37] | PRJNA428542 |
| SS_2464 | 2013 | China | Pig Healthy | Non-Clinical | 499 | 5 | [37] | PRJNA428542 |
| SS_2481 | 2013 | China | Pig Healthy | Non-Clinical | NA | 31 | [37] | PRJNA428542 |
| SS_2737 | 2014 | China | Pig Healthy | Non-Clinical | NA | 30 | [37] | PRJNA428542 |
| SS_2761 | 2014 | China | Pig Healthy | Non-Clinical | NA | NA | [37] | PRJNA428542 |
| SS_2764 | 2014 | China | Pig Healthy | Non-Clinical | NA | NA | [37] | PRJNA428542 |
| SS_2795 | 2014 | China | Pig Healthy | Non-Clinical | NA | NA | [37] | PRJNA428542 |
| SS_2797 | 2014 | China | Pig Healthy | Non-Clinical | NA | NA | [37] | PRJNA428542 |
| SS_2828 | 2014 | China | Pig Healthy | Non-Clinical | NA | 16 | [37] | PRJNA428542 |
| SS_2829 | 2014 | China | Pig Healthy | Non-Clinical | NA | NA | [37] | PRJNA428542 |

| | | | | | | | | |
|---|---|---|---|---|---|---|---|---|
| SS_2838 | 2014 | China | Pig Healthy | Non-Clinical | NA | NA | [37] | PRJNA428542 |
| SS_2873 | 2014 | China | Pig Healthy | Non-Clinical | NA | 15 | [37] | PRJNA428542 |
| SS_3096 | 2014 | China | Pig Healthy | Non-Clinical | NA | NA | [37] | PRJNA428542 |
| SS_3099 | 2014 | China | Pig Healthy | Non-Clinical | NA | NA | [37] | PRJNA428542 |
| 21928 | 2006 | Thailand | Human | Meningitis | 105 | 14 | [25] | NA |
| 25779 | 2007 | Thailand | Human | Sepsis | 105 | 14 | [25] | NA |
| 26390 | 2008 | Thailand | Human | Meningitis | 105 | 14 | [25] | NA |
| 27024 | 2008 | Thailand | Human | Meningitis | 127 | 14 | [25] | NA |
| 28075 | 2008 | Thailand | Human | Sepsis | 237 | 14 | [25] | NA |
| 30333 | 2009 | Thailand | Human | Spondylodiscitis | 105 | 14 | [25] | NA |
| 31998 | 2010 | Thailand | Human | Meningitis | 105 | 14 | [25] | NA |
| 32481 | 2010 | Thailand | Human | Sepsis | 105 | 14 | [25] | NA |
| 32516 | 2010 | Thailand | Human | Meningitis | 105 | 14 | [25] | NA |
| 35728 | 2011 | Thailand | Human | Sepsis | 105 | 14 | [25] | NA |
| 42841 | 2012 | Thailand | Human | Sepsis & Meningitis | 105 | 14 | [25] | NA |
| SAMEA111506260 | 2020 | Thailand | Human | NA | NA | 2 | [28] | PRJEB56918 |
| SAMN03652189 | 2002 | Thailand | Human | NA | 28 | 2 | [6] | PRJNA283528 |
| SAMN04231436 | 2000 | Thailand | Human | Meningitis | 25 | 2 | [5] | PRJNA301006 |
| SAMN04231437 | 2000 | Thailand | Human | Sepsis | 25 | 2 | [5] | PRJNA301006 |
| SAMN04231438 | 2001 | Thailand | Human | Meningitis | 25 | 2 | [5] | PRJNA301006 |
| SAMN04231439 | 2002 | Thailand | Human | Sepsis | 25 | 2 | [5] | PRJNA301006 |
| SAMN04231440 | 2002 | Thailand | Human | Sepsis | 25 | 2 | [5] | PRJNA301006 |
| SAMN04231441 | 2001 | Thailand | Human | Sepsis | 25 | 2 | [5] | PRJNA301006 |
| SAMN04231442 | 2002 | Thailand | Human | Sepsis | 25 | 2 | [5] | PRJNA301006 |
| SAMN04231474 | 2001 | Thailand | Human | Endocarditis | 25 | 2 | [5] | PRJNA301006 |
| SAMN05919918 | 2013 | Thailand | Human | Sepsis | 16 | 9 | [7] | PRJNA349071 |

| | | | | | | | | |
|---|---|---|---|---|---|---|---|---|
| SAMN12911088 | 2007 | Thailand | Human | NA | 1 | 2 | [2] | PRJNA575754 |
| SAMN12911089 | 2007 | Thailand | Pig Diseased | Sepsis | 1 | 2 | [2] | PRJNA575754 |
| SAMN12911090 | 2012 | Thailand | Pig Healthy | Healthy | NA | NT | [2] | PRJNA575754 |
| SAMN12911092 | 2007 | Thailand | Pig Healthy | Healthy | 1 | 2 | [2] | PRJNA575754 |
| SAMN12911093 | 2007 | Thailand | Pig Healthy | Healthy | 1 | 2 | [2] | PRJNA575754 |
| SAMN12911094 | 2007 | Thailand | Pig Diseased | Sepsis | 1 | 2 | [2] | PRJNA575754 |
| SAMN12911095 | 2007 | Thailand | Pig Diseased | Sepsis | NA | 2 | [2] | PRJNA575754 |
| SAMN12911096 | 2012 | Thailand | Pig Diseased | Sepsis | NA | 2 | [2] | PRJNA575754 |
| SAMN13172488 | 2007 | Thailand | Human | Sepsis | 104 | 2 | [2] | PRJNA575754 |
| SAMN17277757 | 2010 | Thailand | Human | Sepsis | 221 | 24 | [29] | PRJNA691075 |
| SAMN17277758 | 2010 | Thailand | Human | Sepsis | 234 | 24 | [29] | PRJNA691075 |
| SAMN19600291 | 2012 | Thailand | Human | Sepsis | 221 | 24 | [29] | PRJNA691075 |
| SAMN20090113 | 2020 | Thailand | Human | Sepsis | 221 | 5 | [29] | PRJNA691075 |
| SAMN17277759 | 2010 | Thailand | Human | Sepsis | 105 | 14 | [30] | PRJNA691075 |
| SAMN17277760 | 2009 | Thailand | Human | Meningitis | 25 | 2 | [30] | PRJNA691075 |
| SAMN17277761 | 2008 | Thailand | Human | Sepsis | 104 | 2 | [30] | PRJNA691075 |
| SAMN17277762 | 2012 | Thailand | Human | Sepsis | 233 | 2 | [30] | PRJNA691075 |
| SAMN17277763 | 2012 | Thailand | Human | Meningitis | 233 | 2 | [30] | PRJNA691075 |
| SAMN17277764 | 2012 | Thailand | Human | Sepsis | 379 | 2 | [30] | PRJNA691075 |
| SAMN17277765 | 2012 | Thailand | Human | Meningitis | 233 | 2 | [30] | PRJNA691075 |
| SAMN17277766 | 2011 | Thailand | Human | Sepsis | 233 | 2 | [30] | PRJNA691075 |
| SAMN18585446 | 2007 | Thailand | Human | Sepsis | 104 | 2 | [30] | PRJNA691075 |
| SAMN18585447 | 2007 | Thailand | Human | Meningitis | 1 | 2 | [30] | PRJNA691075 |
| SAMN29502508 | 2015 | Thailand | Pig Healthy | Healthy | 1923 | 16 | [3] | PRJNA418954 |
| SAMN29502509 | 2015 | Thailand | Pig Healthy | Healthy | 1923 | 16 | [3] | PRJNA418954 |
| SAMN29502515 | 2015 | Thailand | Pig Healthy | Healthy | 1924 | NT | [3] | PRJNA418954 |

| | | | | | | | | |
|---|---|---|---|---|---|---|---|---|
| SAMN29502516 | 2015 | Thailand | Pig Healthy | Healthy | 1924 | 4 | [3] | PRJNA418954 |
| SAMN29502517 | 2015 | Thailand | Pig Healthy | Healthy | 1924 | 4 | [3] | PRJNA418954 |
| SAMN29502527 | 2015 | Thailand | Pig Healthy | Healthy | 1925 | NA | [3] | PRJNA418954 |
| SAMN29502528 | 2015 | Thailand | Pig Healthy | Healthy | 1925 | 16 | [3] | PRJNA418954 |
| SAMN29502529 | 2015 | Thailand | Pig Healthy | Healthy | 1926 | NA | [3] | PRJNA418954 |
| SAMN29502530 | 2015 | Thailand | Pig Healthy | Healthy | 1926 | 11 | [3] | PRJNA418954 |
| SAMN29502531 | 2015 | Thailand | Pig Healthy | Healthy | 1927 | NA | [3] | PRJNA418954 |
| SAMN29502533 | 2015 | Thailand | Pig Healthy | Healthy | 1928 | NA | [3] | PRJNA418954 |
| SAMN29502534 | 2015 | Thailand | Pig Healthy | Healthy | 1929 | NA | [3] | PRJNA418954 |
| SAMN29502535 | 2015 | Thailand | Pig Healthy | Healthy | 1929 | NA | [3] | PRJNA418954 |
| SAMN29502537 | 2015 | Thailand | Pig Healthy | Healthy | 1929 | NA | [3] | PRJNA418954 |
| SAMN29502538 | 2015 | Thailand | Pig Healthy | Healthy | 1930 | NA | [3] | PRJNA418954 |
| SAMN29502541 | 2015 | Thailand | Pig Healthy | Healthy | 1931 | NA | [3] | PRJNA418954 |
| SAMN29502542 | 2015 | Thailand | Pig Healthy | Healthy | 1932 | NT | [3] | PRJNA418954 |
| SAMN29502550 | 2015 | Thailand | Pig Healthy | Healthy | 1933 | 16 | [3] | PRJNA418954 |
| SAMN29502560 | 2015 | Thailand | Pig Healthy | Healthy | 1934 | NT | [3] | PRJNA418954 |
| SAMN29502561 | 2015 | Thailand | Pig Healthy | Healthy | 1934 | NT | [3] | PRJNA418954 |
| SAMN29502567 | 2015 | Thailand | Pig Healthy | Healthy | 1930 | 29 | [3] | PRJNA418954 |
| SAMN29502568 | 2015 | Thailand | Pig Healthy | Healthy | 1935 | NA | [3] | PRJNA418954 |
| SAMN29502569 | 2015 | Thailand | Pig Healthy | Healthy | 1936 | 4 | [3] | PRJNA418954 |
| SAMN29502573 | 2015 | Thailand | Pig Healthy | Healthy | 1937 | 31 | [3] | PRJNA418954 |
| SAMN29502574 | 2015 | Thailand | Pig Healthy | Healthy | NA | NT | [3] | PRJNA418954 |
| SAMN29502575 | 2010 | Thailand | Human | Meningitis | 104 | 2 | [3] | PRJNA418954 |
| SAMN29502576 | 2010 | Thailand | Human | Meningitis | 105 | 2 | [3] | PRJNA418954 |
| SAMN29502577 | 2013 | Thailand | Pig Diseased | Sepsis | 1 | 2 | [3] | PRJNA418954 |
| SAMN29502578 | 2013 | Thailand | Pig Diseased | Sepsis | 1 | 2 | [3] | PRJNA418954 |

| | | | | | | | | |
|---|---|---|---|---|---|---|---|---|
| SAMN29502579 | 2013 | Thailand | Pig Healthy | Healthy | 1938 | 31 | [3] | PRJNA418954 |
| SAMN29502580 | 2013 | Thailand | Pig Healthy | Healthy | 28 | 2 | [3] | PRJNA418954 |
| SAMN29502581 | 2013 | Thailand | Pig Healthy | Healthy | 1939 | 2 | [3] | PRJNA418954 |
| SAMN29502582 | 2013 | Thailand | Pig Healthy | Healthy | 104 | 2 | [3] | PRJNA418954 |
| SAMN29502583 | 2013 | Thailand | Pig Healthy | Healthy | 104 | 2 | [3] | PRJNA418954 |
| SAMN29502584 | 2013 | Thailand | Pig Healthy | Healthy | 1940 | 28 | [3] | PRJNA418954 |
| SAMN29502585 | 2013 | Thailand | Pig Healthy | Healthy | NA | NT | [3] | PRJNA418954 |
| SAMN31277171 | 2011 | Thailand | Pig Healthy | Healthy | NA | 1 | [31] | PRJNA691075 |
| SAMN31277172 | 2011 | Thailand | Human | Sepsis | 105 | 14 | [31] | PRJNA691075 |
| SAMN31277173 | 2011 | Thailand | Pig Healthy | Healthy | NA | 4 | [32] | PRJNA691075 |
| SAMN31277174 | 2011 | Thailand | Human | Sepsis | 94 | 4 | [32] | PRJNA691075 |
| SAMN31277175 | 2010 | Thailand | Pig Diseased | Pneumonia | NA | 4 | [32] | PRJNA691075 |
| SAMN31277176 | 2011 | Thailand | Pig Healthy | Healthy | NA | 4 | [32] | PRJNA691075 |
| SAMN31277177 | 2011 | Thailand | Pig Healthy | Healthy | NA | 4 | [32] | PRJNA691075 |
| SAMN31277178 | 2011 | Thailand | Pig Healthy | Healthy | NA | 4 | [32] | PRJNA691075 |
| SAMN31277179 | 2011 | Thailand | Pig Healthy | Healthy | NA | 4 | [32] | PRJNA691075 |
| STC127 | 2021 | Thailand | Human | Sepsis | 1688 | 2 | [1] | PRJNA853715 |
| STC129 | 2021 | Thailand | Human | Sepsis & Meningitis | 1688 | 2 | [1] | PRJNA853715 |
| STC130 | 2021 | Thailand | Human | Sepsis | 379 | 2 | [1] | PRJNA853715 |
| STC131 | 2021 | Thailand | Human | Sepsis | 233 | 2 | [1] | PRJNA853715 |
| STC152 | 2021 | Thailand | Human | Meningitis | 104 | 2 | [1] | PRJNA853715 |
| STC153 | 2021 | Thailand | Human | NA | 233 | 2 | [1] | PRJNA853715 |
| STC154 | 2021 | Thailand | Human | Spondylodiscitis | 1688 | 2 | [1] | PRJNA853715 |
| STC78 | 2021 | Thailand | Human | Sepsis | 1656 | 2 | [1] | PRJNA853715 |
| STC407 | 2021 | Thailand | Human | Sepsis | NA | 2 | [27] | NA |
| SAMEA103980380 | 2015 | Netherlands | Human | NA | 1 | 2 | [17] | PRJEB20417 |

| | | | | | | | | |
|---|---|---|---|---|---|---|---|---|
| SAMD00009287 | NA | Netherlands | Human | NA | 6 | 14 | [17] | PRJDB1485 |
| SAMN05442334 | 1995 | Argentina | Human | NA | 1 | 2 | [17] | PRJNA335427 |
| SAMN05442335 | 2003 | Argentina | Human | NA | 1 | 2 | [17] | PRJNA335427 |
| SAMN05442336 | 2003 | Argentina | Human | NA | 1 | 2 | [17] | PRJNA335427 |
| SAMN05442338 | 2003 | Argentina | Human | NA | 1 | 2 | [17] | PRJNA335427 |
| SAMN05442339 | 2013 | Argentina | Human | NA | 1 | 2 | [17] | PRJNA335427 |
| SAMN05442340 | 2013 | Argentina | Human | NA | 1 | 2 | [17] | PRJNA335427 |
| SAMN05442341 | 2012 | Argentina | Human | NA | 1 | 2 | [17] | PRJNA335427 |
| SAMN05442342 | 2014 | Argentina | Human | NA | 1 | 2 | [17] | PRJNA335427 |
| SAMN05442343 | 2012 | Argentina | Human | NA | 1 | 2 | [17] | PRJNA335427 |
| SAMN05442344 | 2014 | China | Human | NA | 1 | 2 | [17] | PRJNA335427 |
| SAMN08435268 | 2016 | China | Human | NA | 1 | 2 | [17] | PRJNA432119 |
| SAMN08435269 | 2016 | China | Human | NA | 1 | 2 | [17] | PRJNA432119 |
| SAMN08435270 | 2016 | China | Human | NA | 1 | 2 | [17] | PRJNA432119 |
| SAMN04166386 | 1998 | China | Human | NA | 7 | 2 | [17] | PRJNA298752 |
| SAMN04166388 | 1999 | China | Human | NA | 7 | 2 | [17] | PRJNA298752 |
| SAMN08238443 | 2016 | China | Human | NA | 7 | 2 | [17] | PRJNA427455 |
| SAMN08238502 | 2011 | China | Human | NA | 7 | 2 | [17] | PRJNA427455 |
| SAMN08238506 | 2016 | China | Human | NA | 7 | 2 | [17] | PRJNA427455 |
| SAMN05231907 | 2014 | China | Human | NA | 658 | 2 | [17] | PRJNA325429 |
| SAMN08435267 | 2016 | China | Human | NA | 665 | 2 | [17] | PRJNA432119 |
| SAMN02470636 | NA | Germany | Human | NA | 1 | 2 | [17] | PRJNA171412 |
| SAMN08799542 | 2016 | Italy | Human | NA | 1 | 2 | [17] | PRJNA445803 |
| SAMD00009297 | NA | Japan | Human | NA | 1 | 2 | [17] | PRJDB1485 |
| SAMD00113017 | 2017 | Japan | Human | NA | 28 | 2 | [17] | PRJDB6789 |
| SAMD00066479 | 2016 | Japan | Human | NA | 752 | 5 | [17] | PRJDB5283 |

| | | | | | | | | |
|---|---|---|---|---|---|---|---|---|
| SAMEA4355200 | 2012 | Togo | Human | NA | 1 | 2 | [17] | PRJEB12952 |
| SAMEA4355202 | 2014 | Togo | Human | NA | 1 | 2 | [17] | PRJEB12952 |
| SAMEA4355203 | 2013 | Togo | Human | NA | 1 | 2 | [17] | PRJEB12952 |
| SAMEA4355204 | 2014 | Togo | Human | NA | 1 | 2 | [17] | PRJEB12952 |
| SAMEA4355190 | 2010 | Togo | Human | NA | 956 | 2 | [17] | PRJEB12952 |
| SAMEA4355191 | 2011 | Togo | Human | NA | 956 | 2 | [17] | PRJEB12952 |
| SAMEA4355192 | 2011 | Togo | Human | NA | 956 | 2 | [17] | PRJEB12952 |
| SAMEA4355193 | 2011 | Togo | Human | NA | 956 | 2 | [17] | PRJEB12952 |
| SAMEA4355194 | 2012 | Togo | Human | NA | 956 | 2 | [17] | PRJEB12952 |
| SAMEA4355195 | 2012 | Togo | Human | NA | 956 | 2 | [17] | PRJEB12952 |
| SAMEA4355196 | 2012 | Togo | Human | NA | 956 | 2 | [17] | PRJEB12952 |
| SAMEA4355197 | 2012 | Togo | Human | NA | 956 | 2 | [17] | PRJEB12952 |
| SAMEA4355198 | 2012 | Togo | Human | NA | 956 | 2 | [17] | PRJEB12952 |
| SAMEA4355199 | 2012 | Togo | Human | NA | 956 | 2 | [17] | PRJEB12952 |
| SAMEA4355201 | 2012 | Togo | Human | NA | 956 | 2 | [17] | PRJEB12952 |
| SAMN11854239 | 2015 | USA | Pig Diseased | NA | 225 | 7 | [17] | PRJNA544735 |
| SAMN04376555 | NA | Canada | Pig Diseased | NA | 225 | 7 | [17] | PRJNA307190 |
| SAMN11854154 | NA | USA | Pig Diseased | NA | 225 | 7 | [17] | PRJNA544735 |
| SAMN11854151 | 2014 | USA | Pig Diseased | NA | 225 | 7 | [17] | PRJNA544735 |
| SAMEA3233991 | 2010 | UK | Pig Diseased | NA | 907 | 7 | [17] | PRJEB8392 |
| SAMN04376560 | NA | Canada | Pig Diseased | NA | 32 | 7 | [17] | PRJNA307190 |
| SAMN02469560 | 2012 | China | Pig Healthy | NA | 32 | 7 | [17] | PRJNA171476 |
| CZ1 | NA | Czech Republic | Human | NA | 11 | 14 | [38] | PRJNA917482 |
| CZ2 | NA | Czech Republic | Human | NA | 1 | 2 | [38] | PRJNA917482 |
| CZ3 | NA | Czech Republic | Human | NA | 1 | 2 | [38] | PRJNA917482 |
| CZ4 | 2001 | Czech Republic | Human | | 25 | 2 | [38] | PRJNA917482 |

| | | | | | | | | |
|---|---|---|---|---|---|---|---|---|
| CZ5 | 2003 | Czech Republic | Human | Meningitis | 1 | 2 | [38] | PRJNA917482 |
| CZ6 | 2003 | Czech Republic | Human | Meningitis | 1 | 2 | [38] | PRJNA917482 |
| CZ7 | 2003 | Czech Republic | Human | Meningitis | 25 | 2 | [38] | PRJNA917482 |
| CZ8 | 2004 | Czech Republic | Human | NA | 25 | 2 | [38] | PRJNA917482 |
| CZ9 | 2005 | Czech Republic | Human | Meningitis | 1 | 2 | [38] | PRJNA917482 |
| CZ10 | 2011 | Czech Republic | Human | Meningitis | 1 | 2 | [38] | PRJNA917482 |
| CZ11 | 2012 | Czech Republic | Human | Meningitis and sepsis | 1 | 2 | [38] | PRJNA917482 |
| CZ12 | 2011 | Czech Republic | Human | Sepsis | 29 | 7 | [38] | PRJNA917482 |
| CZ13 | 2012 | Czech Republic | Human | Sepsis | 1 | 2 | [38] | PRJNA917482 |
| CZ14 | 2014 | Czech Republic | Human | Meningitis and sepsis | 1 | 2 | [38] | PRJNA917482 |
| CZ15 | 2015 | Czech Republic | Human | NA | NA | 2 | [38] | PRJNA917482 |
| CZ16 | 2016 | Czech Republic | Human | Sepsis | 1 | 2 | [38] | PRJNA917482 |
| CZ17 | 2005 | Czech Republic | Human | Meningitis | 1660 | 2 | [38] | PRJNA917482 |
| DE1 | 1998 | Germany | Human | NA | 1 | 2 | [38] | PRJNA917482 |
| DE2 | 2001 | Germany | Human | NA | 1 | 2 | [38] | PRJNA917482 |
| DE3 | 2003 | Germany | Human | NA | 1 | 2 | [38] | PRJNA917482 |
| DE4 | 2004 | Germany | Human | NA | 1 | 2 | [38] | PRJNA917482 |
| DE5 | 2005 | Germany | Human | NA | 1 | 2 | [38] | PRJNA917482 |
| DE6 | 2005 | Germany | Human | NA | 1 | 14 | [38] | PRJNA917482 |
| DE7 | 2004 | Germany | Human | NA | 1 | 14 | [38] | PRJNA917482 |
| DE8 | 2008 | Germany | Human | NA | 19 | 2 | [38] | PRJNA917482 |
| DE9 | 2008 | Germany | Human | NA | 19 | 2 | [38] | PRJNA917482 |
| DE10 | 2010 | Germany | Human | NA | 1 | 2 | [38] | PRJNA917482 |
| DE11 | 2010 | Germany | Human | NA | 1 | 2 | [38] | PRJNA917482 |
| DE12 | 1990 | Germany | Human | NA | 1661 | 2 | [38] | PRJNA917482 |
| DE13 | NA | Germany | Human | NA | 1 | 2 | [38] | PRJNA917482 |

| | | | | | | | | |
|---|---|---|---|---|---|---|---|---|
| DE14 | 1994 | Germany | Human | NA | 1 | 14 | [38] | PRJNA917482 |
| DE15 | 1994 | Germany | Human | NA | 1662 | 2 | [38] | PRJNA917482 |
| DE16 | 1999 | Germany | Human | NA | 1663 | 5 | [38] | PRJNA917482 |
| DE17 | 2000 | Germany | Human | NA | 1 | 2 | [38] | PRJNA917482 |
| DE18 | 2001 | Germany | Human | NA | 1 | 2 | [38] | PRJNA917482 |
| DE19 | 2002 | Germany | Human | NA | 1 | 2 | [38] | PRJNA917482 |
| DE20 | 2004 | Germany | Human | NA | 1 | 14 | [38] | PRJNA917482 |
| DE21 | 2018 | Germany | Human | NA | 1707 | 5 | [38] | PRJNA917482 |
| DE22 | 2002 | Germany | Human | NA | 19 | 2 | [38] | PRJNA917482 |
| DK1 | 2016 | Denmark | Human | NA | 1 | 2 | [38] | PRJNA917482 |
| SP1 | 2014 | Spain | Human | NA | 1 | 2 | [38] | PRJNA917482 |
| UK1 | 2017 | UK | Human | Meningitis | 1708 | 2 | [38] | PRJNA917482 |
| UK2 | 2014 | Ireland | Human | Meningitis | 1 | 14 | [38] | PRJNA917482 |
| UK3 | 2015 | Ireland | Human | Sepsis | 1 | 2 | [38] | PRJNA917482 |
| UK4 | 2014 | Ireland | Human | Meningitis | 1 | 14 | [38] | PRJNA917482 |
| UK5 | 2010 | UK | Human | Sepsis | 1 | 2 | [38] | PRJNA917482 |
| UK6 | 2014 | Ireland | Human | Meningitis | 1 | 14 | [38] | PRJNA917482 |
| SWI1 | 2018 | Switzerland | Human | Sepsis | 1 | 14 | [38] | PRJNA917482 |
| SAMN25220527 | 2013 | China | Human | NA | NA | NA | [39] | PRJNA799925 |
| SAMN25212000 | 2016 | China | Human | NA | NA | NA | [39] | PRJNA799908 |
| SAMN18029937 | 2016 | China | Human | NA | 373 | 7 | [39] | PRJNA702910 |
| SAMN21240299 | 2019 | China | Human | Meningitis | NA | 2 | [39] | PRJNA760999 |
| SAMN12784745 | 2014 | China | Human | NA | NA | 14 | [39] | PRJNA566245 |
| SAMN12784744 | 2016 | China | Human | NA | NA | 14 | [39] | PRJNA566245 |
| SAMN12784743 | 2016 | China | Human | NA | NA | 2 | [39] | PRJNA566245 |
| SAMN12784742 | 2016 | China | Human | NA | NA | 2 | [39] | PRJNA566245 |

| | | | | | | | | |
|---|---|---|---|---|---|---|---|---|
| SAMN12784741 | 2016 | China | Human | NA | NA | 2 | [39] | PRJNA566245 |
| SAMN12784740 | 2016 | China | Human | NA | NA | 14 | [39] | PRJNA566245 |
| SAMN12784739 | 2016 | China | Human | NA | NA | 2 | [39] | PRJNA566245 |
| SAMN12784738 | 2016 | China | Human | NA | NA | 2 | [39] | PRJNA566245 |
| SAMN12784737 | 2016 | China | Human | NA | NA | 2 | [39] | PRJNA566245 |
| SAMN12784736 | 2016 | China | Human | NA | NA | 2 | [39] | PRJNA566245 |
| SAMN12784735 | 2016 | China | Human | NA | NA | 2 | [39] | PRJNA566245 |
| SAMN12784734 | 2010 | China | Human | NA | NA | 2 | [39] | PRJNA566245 |
| SAMN12784733 | 2016 | China | Human | NA | NA | 2 | [39] | PRJNA566245 |
| SAMN12784732 | 2016 | China | Human | NA | NA | 2 | [39] | PRJNA566245 |
| SAMN12784731 | 2016 | China | Human | NA | NA | 2 | [39] | PRJNA566245 |
| SAMN12784730 | 2010 | China | Human | NA | NA | 2 | [39] | PRJNA566245 |
| SAMN12784729 | 2016 | China | Human | NA | NA | 2 | [39] | PRJNA566245 |
| SAMN12784728 | 2016 | China | Human | NA | NA | 2 | [39] | PRJNA566245 |
| SAMN12784727 | 2016 | China | Human | NA | NA | 2 | [39] | PRJNA566245 |
| SAMN12784726 | 2016 | China | Human | NA | NA | 2 | [39] | PRJNA566245 |
| SAMN12784725 | 2016 | China | Human | NA | NA | 2 | [39] | PRJNA566245 |
| SAMN12784724 | 2016 | China | Human | NA | NA | 2 | [39] | PRJNA566245 |
| SAMN12784723 | 2012 | China | Human | NA | NA | 2 | [39] | PRJNA566245 |
| SAMN12784722 | 2015 | China | Human | NA | NA | 2 | [39] | PRJNA566245 |
| SAMN12784721 | 2015 | China | Human | NA | NA | 2 | [39] | PRJNA566245 |
| SAMN12784720 | 2013 | China | Human | NA | NA | 2 | [39] | PRJNA566245 |
| SAMN12784719 | 2013 | China | Human | NA | NA | 2 | [39] | PRJNA566245 |
| SAMN12784718 | 2011 | China | Human | NA | NA | 2 | [39] | PRJNA566245 |
| SAMN12784717 | 2009 | China | Human | NA | NA | 2 | [39] | PRJNA566245 |
| SAMN12784716 | 2014 | China | Human | NA | NA | 2 | [39] | PRJNA566245 |

| | | | | | | | | |
|---|---|---|---|---|---|---|---|---|
| SAMN12784715 | 2005 | China | Human | NA | NA | 14 | [39] | PRJNA566245 |
| SAMN12784714 | 2005 | China | Human | NA | NA | 2 | [39] | PRJNA566245 |
| SAMN12784713 | 2007 | China | Human | NA | NA | 14 | [39] | PRJNA566245 |
| SAMN12784712 | 2007 | China | Human | NA | NA | 2 | [39] | PRJNA566245 |
| SAMN12784711 | 2006 | China | Human | NA | NA | 2 | [39] | PRJNA566245 |
| SAMN12784710 | 2005 | China | Human | NA | NA | 2 | [39] | PRJNA566245 |
| SAMN12784709 | 2005 | China | Human | NA | NA | 2 | [39] | PRJNA566245 |
| SAMN12784708 | 2007 | China | Human | NA | NA | 2 | [39] | PRJNA566245 |
| SAMN12784707 | 2007 | China | Human | NA | NA | 2 | [39] | PRJNA566245 |
| SAMN12784706 | 2014 | China | Human | NA | NA | 2 | [39] | PRJNA566245 |
| SAMN12784705 | 2016 | China | Human | NA | NA | 2 | [39] | PRJNA566245 |
| SAMN12784704 | 2015 | China | Human | NA | NA | 2 | [39] | PRJNA566245 |
| SAMN12784703 | 2015 | China | Human | NA | NA | 2 | [39] | PRJNA566245 |
| SAMN12784702 | 2016 | China | Human | NA | NA | 2 | [39] | PRJNA566245 |
| SAMN12784701 | 2016 | China | Human | NA | NA | 2 | [39] | PRJNA566245 |
| SAMN12784700 | 2012 | China | Human | NA | NA | 2 | [39] | PRJNA566245 |
| SAMN12784699 | 2012 | China | Human | NA | NA | 14 | [39] | PRJNA566245 |
| SAMN12784698 | 2013 | China | Human | NA | NA | 14 | [39] | PRJNA566245 |
| SAMN12784697 | 2016 | China | Human | NA | NA | 2 | [39] | PRJNA566245 |
| SAMN12784696 | 2015 | China | Human | NA | NA | 2 | [39] | PRJNA566245 |
| SAMN12784695 | 2015 | China | Human | NA | NA | 2 | [39] | PRJNA566245 |
| SAMN12784694 | 2015 | China | Human | NA | NA | 14 | [39] | PRJNA566245 |
| SAMN12784693 | 2015 | China | Human | NA | NA | 14 | [39] | PRJNA566245 |
| SAMN12784692 | 2014 | China | Human | NA | NA | 2 | [39] | PRJNA566245 |
| SAMN12784691 | 2014 | China | Human | NA | NA | 2 | [39] | PRJNA566245 |
| SAMN12784690 | 2014 | China | Human | NA | NA | 2 | [39] | PRJNA566245 |

| | | | | | | | | |
|---|---|---|---|---|---|---|---|---|
| SAMN12784689 | 2013 | China | Human | NA | NA | 2 | [39] | PRJNA566245 |
| SAMN12784688 | 2014 | China | Human | NA | NA | 2 | [39] | PRJNA566245 |
| SAMN12784687 | 2014 | China | Human | NA | NA | 2 | [39] | PRJNA566245 |
| SAMN12784686 | 2014 | China | Human | NA | NA | 14 | [39] | PRJNA566245 |
| SAMN12784685 | 2013 | China | Human | NA | NA | 2 | [39] | PRJNA566245 |
| SAMN12784684 | 2013 | China | Human | NA | NA | 2 | [39] | PRJNA566245 |
| SAMN12784683 | 2013 | China | Human | NA | NA | 2 | [39] | PRJNA566245 |
| SAMN12784682 | 2013 | China | Human | NA | NA | 14 | [39] | PRJNA566245 |
| SAMN12784681 | 2012 | China | Human | NA | NA | 2 | [39] | PRJNA566245 |
| SAMN12784680 | 2012 | China | Human | NA | NA | 14 | [39] | PRJNA566245 |
| SAMN12784679 | 2012 | China | Human | NA | NA | 2 | [39] | PRJNA566245 |
| SAMN12784678 | 2013 | China | Human | NA | NA | 2 | [39] | PRJNA566245 |
| SAMN12784677 | 2011 | China | Human | NA | NA | 14 | [39] | PRJNA566245 |
| SAMN12784676 | 2011 | China | Human | NA | NA | 2 | [39] | PRJNA566245 |
| SAMN12784675 | 2014 | China | Human | NA | NA | 2 | [39] | PRJNA566245 |
| SAMN12784674 | 2014 | China | Human | NA | NA | 2 | [39] | PRJNA566245 |
| SAMN12784673 | 2014 | China | Human | NA | NA | 2 | [39] | PRJNA566245 |
| SAMN12784672 | 2011 | China | Human | NA | NA | 2 | [39] | PRJNA566245 |
| SAMN12784671 | 2012 | China | Human | NA | NA | 2 | [39] | PRJNA566245 |
| SAMN12784670 | 2011 | China | Human | NA | NA | 2 | [39] | PRJNA566245 |
| SAMN12784669 | 2011 | China | Human | NA | NA | 2 | [39] | PRJNA566245 |
| SAMN12784668 | 2012 | China | Human | NA | NA | 2 | [39] | PRJNA566245 |
| SAMN12784667 | 2010 | China | Human | NA | NA | 14 | [39] | PRJNA566245 |
| SAMN12784666 | 2010 | China | Human | NA | NA | 2 | [39] | PRJNA566245 |
| SAMN12784665 | 2013 | China | Human | NA | NA | 2 | [39] | PRJNA566245 |
| SAMN12784664 | 2014 | China | Human | NA | NA | 2 | [39] | PRJNA566245 |

| | | | | | | | | |
|---|---|---|---|---|---|---|---|---|
| SAMN12784663 | 2014 | China | Human | NA | NA | 2 | [39] | PRJNA566245 |
| SAMN12784662 | 2012 | China | Human | NA | NA | 2 | [39] | PRJNA566245 |
| SAMN12784661 | 2012 | China | Human | NA | NA | 2 | [39] | PRJNA566245 |
| SAMN12784660 | 2012 | China | Human | NA | NA | 2 | [39] | PRJNA566245 |
| SAMN12784659 | 2013 | China | Human | NA | NA | 14 | [39] | PRJNA566245 |
| SAMN12784658 | 2013 | China | Human | NA | NA | 2 | [39] | PRJNA566245 |
| SAMN12784657 | 2016 | China | Human | NA | NA | 14 | [39] | PRJNA566245 |
| SAMN12784656 | 2016 | China | Human | NA | NA | 2 | [39] | PRJNA566245 |
| SAMN12784655 | 2016 | China | Human | NA | NA | 2 | [39] | PRJNA566245 |
| SAMN12784654 | 2016 | China | Human | NA | NA | 2 | [39] | PRJNA566245 |
| SAMN12784653 | 2016 | China | Human | NA | NA | 2 | [39] | PRJNA566245 |
| SAMN18839405 | 2007 | China | Human | NA | NA | 2 | [40] | PRJNA724034 |
| SAMN18839402 | 2005 | China | Human | NA | NA | 2 | [40] | PRJNA724034 |
| SAMN18839382 | 2005 | China | Human | NA | NA | 2 | [40] | PRJNA724034 |
| SAMN18839378 | 2007 | China | Human | NA | NA | 2 | [40] | PRJNA724034 |
| SAMN18839377 | 2007 | China | Human | NA | NA | 2 | [40] | PRJNA724034 |
| SAMN18839376 | 2007 | China | Human | NA | NA | 2 | [40] | PRJNA724034 |
| SAMN18839375 | 2019 | China | Human | NA | NA | 2 | [40] | PRJNA724034 |
| SAMN18839373 | 2019 | China | Human | NA | NA | 2 | [40] | PRJNA724034 |
| SS-194 | 2018 | Thailand | Pig Diseased | Respiratory | 104 | 7 | [26] | NA |
| SS-230 | 2018 | Thailand | Pig Diseased | Respiratory | 94 | 7 | [26] | NA |
| SS-236 | 2018 | Thailand | Pig Diseased | Respiratory | 94 | 7 | [26] | NA |
| SS-500 | 2020 | Thailand | Pig Diseased | Respiratory | 104 | 7 | [26] | NA |
| SS-560 | 2020 | Thailand | Pig Diseased | Respiratory | NA | 7 | [26] | NA |
| 6407_ref | NA | Denmark | Pig Diseased | Unknown | NA | 4 | NA | PRJNA254762 |
| 90_1330_ref | 1990 | Canada | Pig Healthy | Non-Clinical | NA | NA | NA | PRJNA295196 |

| | | | | | | | | |
|---|---|---|---|---|---|---|---|---|
| A7_ref | 2007 | China | Pig Diseased | Systemic | NA | 2 | NA | PRJNA63483 |
| BM407_ref | 2004 | Vietnam | Human | Systemic | NA | 2 | NA | PRJEA32237 |
| CS100322_ref | 2010 | China | Pig Unknown | Unknown | NA | 2 | NA | PRJNA414693 |
| D12_ref | NA | China | Pig Diseased | Systemic | NA | 9 | NA | PRJNA65469 |
| D9_ref | NA | NA | NA | Systemic | NA | 7 | NA | PRJNA65435 |
| DN13_ref | 2013 | China | Pig Unknown | Unknown | NA | 9 | NA | PRJNA319315 |
| GZ0565_ref | 2005 | China | Pig Diseased | Systemic | NA | 9 | NA | PRJNA337985 |
| GZ1_ref | 2005 | China | Human | Systemic | NA | 2 | NA | PRJNA18737 |
| HA0609_ref | 2006 | China | Pig Unknown | Unknown | NA | 2 | NA | PRJNA414694 |
| JS14_ref | NA | China | Pig Diseased | Systemic | NA | 14 | NA | PRJNA61797 |
| _ref | 2014 | China | Human | Systemic | NA | NA | NA | PRJNA325429 |
| NSUI002_ref | 2008 | Canada | Pig Unknown | Unknown | NA | 2 | NA | PRJNA283547 |
| NSUI060_ref | 2008 | Canada | Pig Diseased | Systemic | NA | 2 | NA | PRJNA298205 |
| P1_7_ref | 1984 | UK | Pig Diseased | Systemic | NA | 2 | NA | PRJNA352 |
| S10_ref | 1982 | Netherlands | Pig Diseased | Systemic | NA | 2 | NA | PRJEB35407 |
| S735_ref | NA | Netherlands | Pig Diseased | Respiratory | NA | 2 | NA | PRJNA163139 |
| SC070731_ref | 2007 | China | Pig Diseased | Systemic | NA | NA | NA | PRJNA178071 |
| SC19_ref | 2005 | China | Pig Diseased | Systemic | NA | 2 | NA | PRJNA383459 |
| SC84_ref | 2005 | China | Human | Systemic | NA | NA | NA | PRJEA32239 |
| SS12_ref | NA | China | Pig Diseased | Respiratory | NA | 1/2 | NA | PRJNA65433 |
| SS2_1_ref | 1998 | China | Pig Diseased | Unknown | NA | 2 | NA | PRJNA359165 |
| ST1_ref | NA | NA | NA | Systemic | NA | NA | NA | PRJNA65471 |
| ST3_ref | NA | China | Pig Diseased | Systemic | NA | 1 | NA | PRJNA65245 |
| T15_ref | NA | Netherlands | Pig Healthy | Non-Clinical | NA | NA | NA | PRJNA206218 |
| TL13_ref | NA | China | Pig Unknown | Unknown | NA | 16 | NA | PRJNA171404 |
| YB51_ref | NA | China | Pig Healthy | Non-Clinical | NA | 3 | NA | PRJNA213888 |

| | | | | | | | | |
|---|---|---|---|---|---|---|---|---|
| ZY05719_ref | 2005 | China | Pig Diseased | Systemic | NA | 2 | NA | PRJNA242223 |
| SAMD00039895 | 2000 | Thailand | Human | Endocarditis | NA | 2 | NA | PRJDB4183 |
| SAMD00039896 | 2000 | Thailand | Human | NA | NA | 2 | NA | PRJDB4183 |
| SAMD00039897 | 2000 | Thailand | Human | NA | NA | 2 | NA | PRJDB4183 |
| SAMD00039898 | 2000 | Thailand | Human | Meningitis | NA | 2 | NA | PRJDB4183 |
| SAMD00039899 | 2001 | Thailand | Human | NA | NA | 2 | NA | PRJDB4183 |
| SAMD00039900 | 2001 | Thailand | Human | NA | NA | 2 | NA | PRJDB4183 |
| SAMD00039901 | 2000 | Thailand | Human | Meningitis | NA | 2 | NA | PRJDB4183 |
| SAMD00039902 | 2002 | Thailand | Human | Meningitis | NA | 2 | NA | PRJDB4183 |
| SAMD00039903 | 2002 | Thailand | Human | NA | NA | 2 | NA | PRJDB4183 |
| SAMD00039904 | 2002 | Thailand | Human | NA | NA | 2 | NA | PRJDB4183 |
| SAMD00039905 | 2002 | Thailand | Human | NA | NA | 2 | NA | PRJDB4183 |
| SAMD00039906 | 2002 | Thailand | Human | Meningitis | NA | 2 | NA | PRJDB4183 |
| SAMD00039907 | 2001 | Thailand | Human | Meningitis | NA | 2 | NA | PRJDB4183 |
| SAMD00039908 | 2001 | Thailand | Human | Sepsis | NA | 2 | NA | PRJDB4183 |
| SAMD00039909 | 2000 | Thailand | Human | NA | NA | 2 | NA | PRJDB4183 |
| SAMD00039910 | 2002 | Thailand | Human | NA | NA | 2 | NA | PRJDB4183 |
| SAMD00039911 | 2002 | Thailand | Human | NA | NA | 2 | NA | PRJDB4183 |
| SAMN27578139 | 2005 | China | Human | NA | NA | NA | NA | PRJNA826693 |
| SAMN27553195 | 2006 | China | Human | Meningitis | NA | NA | NA | PRJNA826693 |
| SAMD00039914 | 2005 | Japan | Human | Meningitis | NA | 2 | NA | PRJDB4183 |
| SAMD00039913 | 2010 | Japan | Human | Sepsis | NA | 2 | NA | PRJDB4183 |
| SAMD00039912 | 1994 | Japan | Human | Sepsis | NA | 2 | NA | PRJDB4183 |
| SAMN26241748 | 2021 | China | Human | Meningitis | NA | NA | NA | PRJNA810294 |
| SAMN26241767 | 2017 | China | Human | NA | NA | NA | NA | PRJNA810299 |
| SAMN26241766 | 2016 | China | Human | NA | NA | NA | NA | PRJNA810299 |

| | | | | | | | | |
|---|---|---|---|---|---|---|---|---|
| SAMN26241765 | 2015 | China | Human | NA | NA | NA | NA | PRJNA810299 |
| SAMN26241764 | 2009 | China | Human | NA | NA | NA | NA | PRJNA810299 |
| SAMN26241763 | 2006 | China | Human | NA | NA | NA | NA | PRJNA810299 |
| SAMN13975591 | 2015 | USA | Pig Unknown | NA | NA | 7 | NA | PRJNA604583 |
| SAMN17982943 | 2016 | China | Pig Diseased | NA | 225 | 7 | NA | PRJNA702910 |
| SAMN17982944 | NA | China | Pig Diseased | NA | 225 | 7 | NA | PRJNA702910 |
| SAMN17982945 | NA | China | Pig Healthy | NA | 225 | 7 | NA | PRJNA702910 |
| SAMN18117670 | 2019 | Canada | Pig Diseased | Meningitis | 1614 | 7 | NA | PRJNA702910 |
| SAMN18117671 | 2019 | Canada | Pig Diseased | NA | 32 | 7 | NA | PRJNA702910 |
| SAMN18117673 | 2018 | Canada | Pig Diseased | Meningitis | 32 | 7 | NA | PRJNA702910 |
| SAMN20087851 | 2012 | China | Pig Healthy | Healthy | 32 | 7 | NA | PRJNA702910 |
| SAMD00009282 | 1998 | Thailand | Human | Meningitis | 104 | 2 | NA | PRJDB1485 |
| SAMD00009289 | 2002 | Thailand | Human | Pulmonary Edema | 104 | 2 | NA | PRJDB1485 |
| 41594 | 2012 | Thailand | Human | Sepsis | 236 | 2 | This_Project | PRJNA1210814 |
| 34584 | 2011 | Thailand | Human | Sepsis | 515 | 2 | This_Project | PRJNA1210814 |
| 36900 | 2011 | Thailand | Human | Sepsis | 516 | 2 | This_Project | PRJNA1210814 |
| STC19 | 2021 | Thailand | Human | NA | | 2 | This_Project | PRJNA1210814 |
| 23167 | 2006 | Thailand | Human | Meningitis | 1 | 2 | This_Project | PRJNA1210814 |
| 24526 | 2007 | Thailand | Human | Meningitis | 126 | 2 | This_Project | PRJNA1210814 |
| 26647 | 2008 | Thailand | Human | Meningitis | 1 | 2 | This_Project | PRJNA1210814 |
| 32599 | 2010 | Thailand | Human | Sepsis | 11 | 2 | This_Project | PRJNA1210814 |
| 36363 | 2011 | Thailand | Human | Sepsis | 144 | 2 | This_Project | PRJNA1210814 |
| 35736 | 2011 | Thailand | Human | Sepsis | 298 | 2 | This_Project | PRJNA1210814 |
| 40128 | 2012 | Thailand | Human | Sepsis | 337 | 2 | This_Project | PRJNA1210814 |
| STC217 | 2013 | Thailand | Human | NA | 1 | 2 | This_Project | PRJNA1210814 |
| STC224 | 2014 | Thailand | Human | NA | 1 | 2 | This_Project | PRJNA1210814 |

| | | | | | | | | |
|---|---|---|---|---|---|---|---|---|
| STC231 | 2015 | Thailand | Human | NA | 1 | 2 | This_Project | PRJNA1210814 |
| STC238 | 2016 | Thailand | Human | NA | 1 | 2 | This_Project | PRJNA1210814 |
| STC247 | 2017 | Thailand | Human | NA | 1 | 2 | This_Project | PRJNA1210814 |
| STC275 | 2019 | Thailand | Human | NA | 1 | 2 | This_Project | PRJNA1210814 |
| STC293 | 2020 | Thailand | Human | NA | 105 | 2 | This_Project | PRJNA1210814 |
| STC123 | 2021 | Thailand | Human | NA | 105 | 2 | This_Project | PRJNA1210814 |
| 35896 | 2011 | Thailand | Human | Meningitis | 105 | 2 | This_Project | PRJNA1210814 |
| 27964 | 2008 | Thailand | Human | Sepsis | 105 | 14 | This_Project | PRJNA1210814 |
| 28074 | 2008 | Thailand | Human | Sepsis | 105 | 14 | This_Project | PRJNA1210814 |
| 30198 | 2009 | Thailand | Human | Sepsis & Meningitis | 105 | 14 | This_Project | PRJNA1210814 |
| 32941 | 2010 | Thailand | Human | Sepsis | 105 | 14 | This_Project | PRJNA1210814 |
| 39957 | 2012 | Thailand | Human | Sepsis | 105 | 14 | This_Project | PRJNA1210814 |
| 38826 | 2011 | Thailand | Human | Sepsis | 105 | 14 | This_Project | PRJNA1210814 |
| 22696 | 2006 | Thailand | Human | Spondylodiscits | 104 | 2 | This_Project | PRJNA1210814 |
| 23163 | 2006 | Thailand | Human | Meningitis | 104 | 2 | This_Project | PRJNA1210814 |
| 25777 | 2007 | Thailand | Human | Sepsis | 104 | 2 | This_Project | PRJNA1210814 |
| 24529 | 2007 | Thailand | Human | Meningitis | 104 | 2 | This_Project | PRJNA1210814 |
| 29281 | 2008 | Thailand | Human | Sepsis | 104 | 2 | This_Project | PRJNA1210814 |
| 26127 | 2008 | Thailand | Human | Meningitis | 104 | 2 | This_Project | PRJNA1210814 |
| 31106 | 2009 | Thailand | Human | Sepsis | 104 | 2 | This_Project | PRJNA1210814 |
| 30341 | 2009 | Thailand | Human | Sepsis | 104 | 2 | This_Project | PRJNA1210814 |
| 33029 | 2010 | Thailand | Human | Sepsis | 104 | 2 | This_Project | PRJNA1210814 |
| 32861 | 2010 | Thailand | Human | Sepsis | 104 | 2 | This_Project | PRJNA1210814 |
| 33004 | 2010 | Thailand | Human | Sepsis | 104 | 2 | This_Project | PRJNA1210814 |
| 36921 | 2011 | Thailand | Human | Sepsis | 104 | 2 | This_Project | PRJNA1210814 |
| 36369 | 2011 | Thailand | Human | Sepsis | 512 | 2 | This_Project | PRJNA1210814 |

| | | | | | | | | |
|---|---|---|---|---|---|---|---|---|
| 40414 | 2012 | Thailand | Human | Sepsis | 514 | 2 | This_Project | PRJNA1210814 |
| 40621 | 2012 | Thailand | Human | Sepsis | 392 | 2 | This_Project | PRJNA1210814 |
| 39971 | 2012 | Thailand | Human | Sepsis | 512 | 2 | This_Project | PRJNA1210814 |
| 40133 | 2012 | Thailand | Human | Sepsis | 513 | 2 | This_Project | PRJNA1210814 |
| 40302 | 2012 | Thailand | Human | Sepsis | 393 | 2 | This_Project | PRJNA1210814 |
| STC242 | 2016 | Thailand | Human | NA | 104 | 2 | This_Project | PRJNA1210814 |
| STC246 | 2017 | Thailand | Human | NA | 104 | 2 | This_Project | PRJNA1210814 |
| STC244 | 2017 | Thailand | Human | NA | 104 | 2 | This_Project | PRJNA1210814 |
| STC256 | 2019 | Thailand | Human | NA | 104 | 2 | This_Project | PRJNA1210814 |
| STC253 | 2019 | Thailand | Human | NA | 104 | 2 | This_Project | PRJNA1210814 |
| STC290 | 2020 | Thailand | Human | NA | 104 | 2 | This_Project | PRJNA1210814 |
| STC291 | 2020 | Thailand | Human | NA | 104 | 2 | This_Project | PRJNA1210814 |
| STC292 | 2020 | Thailand | Human | NA | 104 | 2 | This_Project | PRJNA1210814 |
| STC282 | 2020 | Thailand | Human | NA | 104 | 2 | This_Project | PRJNA1210814 |
| STC283 | 2020 | Thailand | Human | NA | 104 | 2 | This_Project | PRJNA1210814 |
| STC72 | 2021 | Thailand | Human | NA | 104 | 2 | This_Project | PRJNA1210814 |
| STC269 | 2021 | Thailand | Human | NA | 104 | 2 | This_Project | PRJNA1210814 |
| STC266 | 2021 | Thailand | Human | NA | 104 | 2 | This_Project | PRJNA1210814 |
| STC268 | 2021 | Thailand | Human | NA | 104 | 2 | This_Project | PRJNA1210814 |
| STC267 | 2021 | Thailand | Human | NA | 104 | 2 | This_Project | PRJNA1210814 |
| STC262 | 2021 | Thailand | Human | NA | 104 | 2 | This_Project | PRJNA1210814 |
| STC270 | 2021 | Thailand | Human | NA | 104 | 2 | This_Project | PRJNA1210814 |
| 30389 | 2009 | Thailand | Human | Sepsis | 233 | 2 | This_Project | PRJNA1210814 |
| 27968 | 2008 | Thailand | Human | Sepsis | 233 | 2 | This_Project | PRJNA1210814 |
| 30823 | 2009 | Thailand | Human | Sepsis | 233 | 2 | This_Project | PRJNA1210814 |
| 33394 | 2010 | Thailand | Human | Sepsis | 233 | 2 | This_Project | PRJNA1210814 |

| | | | | | | | | |
|---|---|---|---|---|---|---|---|---|
| 35940 | 2011 | Thailand | Human | Sepsis | 233 | 2 | This_Project | PRJNA1210814 |
| 37880 | 2011 | Thailand | Human | Sepsis | 379 | 2 | This_Project | PRJNA1210814 |
| 35949 | 2011 | Thailand | Human | Sepsis | 233 | 2 | This_Project | PRJNA1210814 |
| 35688 | 2011 | Thailand | Human | Sepsis | 233 | 2 | This_Project | PRJNA1210814 |
| 37554 | 2011 | Thailand | Human | Sepsis | 233 | 2 | This_Project | PRJNA1210814 |
| 39236 | 2012 | Thailand | Human | Sepsis | 233 | 2 | This_Project | PRJNA1210814 |
| 42527 | 2012 | Thailand | Human | Sepsis | 233 | 2 | This_Project | PRJNA1210814 |
| 40750 | 2012 | Thailand | Human | Sepsis | 233 | 2 | This_Project | PRJNA1210814 |
| 40804 | 2012 | Thailand | Human | Sepsis | 233 | 2 | This_Project | PRJNA1210814 |
| 41477 | 2012 | Thailand | Human | Sepsis | 233 | 2 | This_Project | PRJNA1210814 |
| 41219 | 2012 | Thailand | Human | Sepsis | 233 | 2 | This_Project | PRJNA1210814 |
| STC223 | 2014 | Thailand | Human | NA | 233 | 2 | This_Project | PRJNA1210814 |
| STC233 | 2015 | Thailand | Human | NA | 233 | 2 | This_Project | PRJNA1210814 |
| STC235 | 2015 | Thailand | Human | NA | 233 | 2 | This_Project | PRJNA1210814 |
| STC239 | 2016 | Thailand | Human | NA | 233 | 2 | This_Project | PRJNA1210814 |
| STC243 | 2016 | Thailand | Human | NA | 233 | 2 | This_Project | PRJNA1210814 |
| STC250 | 2017 | Thailand | Human | NA | 233 | 2 | This_Project | PRJNA1210814 |
| STC257 | 2019 | Thailand | Human | NA | 379 | 2 | This_Project | PRJNA1210814 |
| STC274 | 2019 | Thailand | Human | NA | 379 | 2 | This_Project | PRJNA1210814 |
| STC279 | 2020 | Thailand | Human | NA | 233 | 2 | This_Project | PRJNA1210814 |
| STC286 | 2020 | Thailand | Human | NA | 233 | 2 | This_Project | PRJNA1210814 |
| STC280 | 2020 | Thailand | Human | NA | 379 | 2 | This_Project | PRJNA1210814 |
| STC141 | 2021 | Thailand | Human | NA | 233 | 2 | This_Project | PRJNA1210814 |
| STC135 | 2021 | Thailand | Human | NA | 379 | 2 | This_Project | PRJNA1210814 |
| STC31 | 2021 | Thailand | Human | NA | New | 2 | This_Project | PRJNA1210814 |
| STC212 | 2021 | Thailand | Human | NA | 233 | 2 | This_Project | PRJNA1210814 |

| | | | | | | | | |
|---|---|---|---|---|---|---|---|---|
| STC173 | 2021 | Thailand | Human | NA | 233 | 2 | This_Project | PRJNA1210814 |
| STC260 | 2021 | Thailand | Human | NA | 1656 | 2 | This_Project | PRJNA1210814 |
| 22714 | 2006 | Thailand | Human | Sepsis | 25 | 2 | This_Project | PRJNA1210814 |
| 25781 | 2007 | Thailand | Human | Meningitis | 103 | 2 | This_Project | PRJNA1210814 |
| 24205 | 2007 | Thailand | Human | Sepsis | 25 | 2 | This_Project | PRJNA1210814 |
| 27103 | 2008 | Thailand | Human | Sepsis | 25 | 2 | This_Project | PRJNA1210814 |
| 31095 | 2009 | Thailand | Human | Sepsis | 103 | 2 | This_Project | PRJNA1210814 |
| 33243 | 2010 | Thailand | Human | Sepsis | 381 | 2 | This_Project | PRJNA1210814 |
| 34758 | 2011 | Thailand | Human | Sepsis | 380 | 2 | This_Project | PRJNA1210814 |
| 36375 | 2011 | Thailand | Human | Sepsis | 395 | 2 | This_Project | PRJNA1210814 |
| 40417 | 2012 | Thailand | Human | Sepsis | 25 | 2 | This_Project | PRJNA1210814 |
| 23164 | 2006 | Thailand | Human | Meningitis | 28 | 2 | This_Project | PRJNA1210814 |
| 28739 | 2008 | Thailand | Human | Sepsis | 28 | 2 | This_Project | PRJNA1210814 |
| 36763 | 2011 | Thailand | Human | Sepsis | 28 | NT | This_Project | PRJNA1210814 |
| 42319 | 2012 | Thailand | Human | Sepsis | 382 | 2 | This_Project | PRJNA1210814 |
| 34536 | 2010 | Thailand | Pig Healthy | Healthy | 299 | 2 | This_Project | PRJNA1210814 |
| 34537 | 2010 | Thailand | Pig Healthy | Healthy | 104 | 2 | This_Project | PRJNA1210814 |
| 34384 | 2010 | Thailand | Pig Healthy | Healthy | 2 | 2 | This_Project | PRJNA1210814 |
| 35125 | 2010 | Thailand | Pig Healthy | Healthy | 28 | 2 | This_Project | PRJNA1210814 |
| 35150 | 2010 | Thailand | Pig Healthy | Healthy | 1 | 2 | This_Project | PRJNA1210814 |
| 35415 | 2010 | Thailand | Pig Healthy | Healthy | 28 | 2 | This_Project | PRJNA1210814 |
| 35781 | 2010 | Thailand | Pig Healthy | Healthy | 1 | 14 | This_Project | PRJNA1210814 |
| 34677 | 2010 | Thailand | Pig Healthy | Healthy | NA | 7 | This_Project | PRJNA1210814 |
| 34703 | 2010 | Thailand | Pig Healthy | Healthy | NA | 7 | This_Project | PRJNA1210814 |
| 34555 | 2010 | Thailand | Pig Healthy | Healthy | NA | 16 | This_Project | PRJNA1210814 |
| 34538 | 2010 | Thailand | Pig Healthy | Healthy | NA | 16 | This_Project | PRJNA1210814 |

| | | | | | | | | |
|---|---|---|---|---|---|---|---|---|
| 34560 | 2010 | Thailand | Pig Healthy | Healthy | NA | 24 | This_Project | PRJNA1210814 |
| 34408 | 2010 | Thailand | Pig Healthy | Healthy | NA | 15 | This_Project | PRJNA1210814 |
| 34679 | 2010 | Thailand | Pig Healthy | Healthy | NA | 29 | This_Project | PRJNA1210814 |
| 41445 | 2012 | Thailand | Human | Sepsis | 105 | 2 | This_Project | PRJNA1210814 |
| 30346 | 2009 | Thailand | Human | Endocarditis | 233 | 2 | This_Project | PRJNA1210814 |
| 30827 | 2009 | Thailand | Human | ARDS | 1 | 2 | This_Project | PRJNA1210814 |
| 34391 | 2010 | Thailand | Pig Healthy | Healthy | NA | 31 | This_Project | PRJNA1210814 |
| 34428 | 2010 | Thailand | Pig Healthy | Healthy | 117 | 3 | This_Project | PRJNA1210814 |
| 34429 | 2010 | Thailand | Pig Healthy | Healthy | NA | 9 | This_Project | PRJNA1210814 |
| 34533 | 2010 | Thailand | Pig Healthy | Healthy | NA | 31 | This_Project | PRJNA1210814 |
| 34552 | 2010 | Thailand | Pig Healthy | Healthy | NA | 19 | This_Project | PRJNA1210814 |
| 34688 | 2010 | Thailand | Pig Healthy | Healthy | NA | 7 | This_Project | PRJNA1210814 |
| 36602 | 2011 | Thailand | Human | NA | 379 | 2 | This_Project | PRJNA1210814 |
| 36610 | 2011 | Thailand | Human | Sepsis | 391 | 2 | This_Project | PRJNA1210814 |
| 38727 | 2011 | Thailand | Human | Sepsis | 233 | 2 | This_Project | PRJNA1210814 |
| 39825 | 2012 | Thailand | Human | Meningitis | 105 | 14 | This_Project | PRJNA1210814 |
| 40301 | 2012 | Thailand | Human | Sepsis | 233 | 2 | This_Project | PRJNA1210814 |
| 40614 | 2012 | Thailand | Human | Sepsis | 1 | 2 | This_Project | PRJNA1210814 |
| STC7 | 2019 | Thailand | Human | NA | 11 | 14 | This_Project | PRJNA1210814 |
| STC147 | 2021 | Thailand | Human | NA | 25 | 2 | This_Project | PRJNA1210814 |
| STC149 | 2021 | Thailand | Human | NA | New | 14 | This_Project | PRJNA1210814 |
| STC228 | 2014 | Thailand | Human | NA | 104 | 2 | This_Project | PRJNA1210814 |
| STC240 | 2016 | Thailand | Human | NA | 379 | 2 | This_Project | PRJNA1210814 |
| STC251 | 2019 | Thailand | Human | NA | 104 | 2 | This_Project | PRJNA1210814 |

**Supplementary Table 6**. Isolates selected for MIC testing with penicillin and ceftriaxone. NA stands for not available. CC stands for clonal complex.

| Isolate | Allele | CC | BioSample |
|---|---|---|---|
| 1584695 | 14 | 16 | SAMN05919918 |
| 33329 | 21 | 221/234 | SAMN17277757 |
| 32098 | 21 | 221/234 | SAMN17277758 |
| 27715 | 31 | 104 | SAMN17277761 |
| 40085 | 26 | 233 | SAMN17277764 |
| 24525 | 10 | 104 | SAMN18585446 |
| 26154 | 1 | 1 | SAMN18585447 |
| 39565 | 22 | 221/234 | SAMN19600291 |
| 41570 | 21 | 221/234 | SAMN20090113 |
| 35541 | 35 | NA | SAMN31277171 |
| 34572 | 19 | NA | SAMN31277173 |
| 36054 | 51 | 94 | SAMN31277174 |
| TRG6 | 18 | NA | SAMN31277175 |
| 34545 | 19 | NA | SAMN31277176 |
| 34553 | 19 | NA | SAMN31277177 |
| 34693 | 18 | NA | SAMN31277178 |
| 34704 | 18 | NA | SAMN31277179 |
| 23163 | 3 | 104 | SAMN46258699 |
| 24205 | 15 | 25 | SAMN46258702 |
| 26127 | 30 | 104 | SAMN46258707 |
| 26647 | 3 | 104 | SAMN46258708 |
| 27103 | 16 | 25 | SAMN46258709 |
| 28074 | 23 | 1 | SAMN46258712 |
| 28739 | 12 | 28 | SAMN46258713 |

| | | | |
|---|---|---|---|
| 29281 | 8 | 104 | SAMN46258714 |
| 30341 | 9 | 104 | SAMN46258716 |
| 31095 | 15 | 25 | SAMN46258721 |
| 32599 | 2 | 1 | SAMN46258723 |
| 32861 | 9 | 104 | SAMN46258724 |
| 33004 | 32 | 104 | SAMN46258726 |
| 34391 | 49 | NA | SAMN46258731 |
| 34408 | 48 | NA | SAMN46258732 |
| 34428 | 13 | 28 | SAMN46258733 |
| 34429 | 66 | NA | SAMN46258734 |
| 34533 | 37 | NA | SAMN46258735 |
| 34536 | 12 | 28 | SAMN46258736 |
| 34538 | 62 | NA | SAMN46258738 |
| 34552 | 34 | NA | SAMN46258739 |
| 34555 | 40 | NA | SAMN46258740 |
| 34584 | 15 | 29 | SAMN46258742 |
| 34677 | 52 | NA | SAMN46258743 |
| 34679 | 36 | NA | SAMN46258744 |
| 34688 | 52 | NA | SAMN46258745 |
| 34703 | 52 | NA | SAMN46258746 |
| 34758 | 16 | 25 | SAMN46258747 |
| 36369 | 7 | 104 | SAMN46258758 |
| 36375 | 9 | 25 | SAMN46258759 |
| 36763 | 12 | 28 | SAMN46258762 |
| 36900 | 16 | 29 | SAMN46258763 |
| 40301 | 28 | 233 | SAMN46258775 |

| | | | |
|---|---|---|---|
| 40414 | 30 | 104 | SAMN46258777 |
| 41594 | 24 | NA | SAMN46258786 |
| 42527 | 28 | 233 | SAMN46258788 |
| STC127 | 43 | 1688 | SAMN29401333 |
| STC131 | 29 | 233 | SAMN29401336 |
| STC135 | 44 | 233 | SAMN46258790 |
| STC141 | 46 | 233 | SAMN46258791 |
| STC152 | 7 | 104 | SAMN29401337 |
| STC154 | 43 | 1688 | SAMN29401339 |
| STC19 | 43 | 1688 | SAMN46258795 |
| STC212 | 28 | 233 | SAMN46258796 |
| STC231 | 1 | 1 | SAMN46258801 |
| STC233 | 26 | 233 | SAMN46258802 |
| STC239 | 26 | 233 | SAMN46258805 |
| STC242 | 3 | 104 | SAMN46258807 |
| STC246 | 30 | 104 | SAMN46258810 |
| STC260 | 46 | 233 | SAMN46258817 |
| STC266 | 33 | 104 | SAMN46258819 |
| STC269 | 33 | 104 | SAMN46258822 |
| STC270 | 33 | 104 | SAMN46258823 |
| STC274 | 44 | 233 | SAMN46258824 |
| STC280 | 44 | 233 | SAMN46258827 |
| STC292 | 7 | 104 | SAMN46258833 |
| STC293 | 1 | 1 | SAMN46258834 |
| STC31 | 47 | 233 | SAMN46258835 |
| STC407 | 61 | 233 | NA |

| | | | |
|---|---|---|---|
| STC7 | 38 | 1 | SAMN46258836 |
| STC72 | 45 | 104 | This study |
| STC78 | 46 | 233 | This study |

**Supplementary Table 7**. Metadata of isolates selected for BEAST analysis. NA stands for not available. UK stands for United Kingdom. USA stands for United States of America. ST stands for ST.

| Isolate name | Year | Country | Host | Disease association | ST | Serotype | Reference | Bioproject |
|---|---|---|---|---|---|---|---|---|
| 33164_Y2125 | 1996 | UK | Pig Diseased | Respiratory | 34 | 7 | [16] | PRJNA1009400 |
| 1506978 | 2013 | Canada | Pig Diseased | Systemic | 32 | 7 | [20] | PRJNA628943 |
| 1615460 | 2014 | Canada | Pig Unknown | Unknown | 225 | 7 | [20] | PRJNA628943 |
| 87-4647 | 1987 | Canada | Pig Unknown | Unknown | 225 | 7 | [20] | PRJNA628943 |
| 89-5189 | 1989 | Canada | Pig Unknown | Unknown | 225 | 7 | [20] | PRJNA628943 |
| SS1022 | 2010 | UK | Pig Diseased | Respiratory | 32 | 7 | [34] | PRJEB8392 |
| 0435-10_S10_L001 | 2019 | USA | Pig Diseased | Unknown | 225 | 7 | [16] | PRJNA972671 |
| 0435-9_S9_L001 | 2019 | USA | Pig Diseased | Unknown | 225 | 7 | [16] | PRJNA972671 |
| 108-24D_S23_L001 | 2018 | USA | Pig Diseased | Unknown | 225 | 7 | [16] | PRJNA972671 |
| 2957-4_S4_L001 | 2018 | USA | Pig Diseased | Unknown | 225 | 7 | [16] | PRJNA972671 |
| A-08-1-TT-01-SS-C4 | NA | Myanmar | Pig Unknown | Unknown | 225 | 7 | [16] | PRJNA1012585 |
| 26647 | 2008 | Thailand | Human | Meningitis | 1 | 2 | This_Project | PRJNA1210814 |
| STC238 | 2016 | Thailand | Human | NA | 1 | 2 | This_Project | PRJNA1210814 |
| 22696 | 2006 | Thailand | Human | Spondylodiscits | 104 | 2 | This_Project | PRJNA1210814 |
| 23163 | 2006 | Thailand | Human | Meningitis | 104 | 2 | This_Project | PRJNA1210814 |
| 25777 | 2007 | Thailand | Human | Sepsis | 104 | 2 | This_Project | PRJNA1210814 |
| 24529 | 2007 | Thailand | Human | Meningitis | 104 | 2 | This_Project | PRJNA1210814 |
| 29281 | 2008 | Thailand | Human | Sepsis | 104 | 2 | This_Project | PRJNA1210814 |
| 26127 | 2008 | Thailand | Human | Meningitis | 104 | 2 | This_Project | PRJNA1210814 |
| 31106 | 2009 | Thailand | Human | Sepsis | 104 | 2 | This_Project | PRJNA1210814 |
| 30341 | 2009 | Thailand | Human | Sepsis | 104 | 2 | This_Project | PRJNA1210814 |
| 33029 | 2010 | Thailand | Human | Sepsis | 104 | 2 | This_Project | PRJNA1210814 |
| 32861 | 2010 | Thailand | Human | Sepsis | 104 | 2 | This_Project | PRJNA1210814 |

| | | | | | | | | |
|---|---|---|---|---|---|---|---|---|
| 33004 | 2010 | Thailand | Human | Sepsis | 104 | 2 | This_Project | PRJNA1210814 |
| 36921 | 2011 | Thailand | Human | Sepsis | 104 | 2 | This_Project | PRJNA1210814 |
| 36369 | 2011 | Thailand | Human | Sepsis | 512 | 2 | This_Project | PRJNA1210814 |
| 40414 | 2012 | Thailand | Human | Sepsis | 514 | 2 | This_Project | PRJNA1210814 |
| 40621 | 2012 | Thailand | Human | Sepsis | 392 | 2 | This_Project | PRJNA1210814 |
| 39971 | 2012 | Thailand | Human | Sepsis | 512 | 2 | This_Project | PRJNA1210814 |
| 40133 | 2012 | Thailand | Human | Sepsis | 513 | 2 | This_Project | PRJNA1210814 |
| 40302 | 2012 | Thailand | Human | Sepsis | 393 | 2 | This_Project | PRJNA1210814 |
| STC242 | 2016 | Thailand | Human | NA | 104 | 2 | This_Project | PRJNA1210814 |
| STC246 | 2017 | Thailand | Human | NA | 104 | 2 | This_Project | PRJNA1210814 |
| STC244 | 2017 | Thailand | Human | NA | 104 | 2 | This_Project | PRJNA1210814 |
| STC256 | 2019 | Thailand | Human | NA | 104 | 2 | This_Project | PRJNA1210814 |
| STC253 | 2019 | Thailand | Human | NA | 104 | 2 | This_Project | PRJNA1210814 |
| STC290 | 2020 | Thailand | Human | NA | 104 | 2 | This_Project | PRJNA1210814 |
| STC291 | 2020 | Thailand | Human | NA | 104 | 2 | This_Project | PRJNA1210814 |
| STC292 | 2020 | Thailand | Human | NA | 104 | 2 | This_Project | PRJNA1210814 |
| STC282 | 2020 | Thailand | Human | NA | 104 | 2 | This_Project | PRJNA1210814 |
| STC283 | 2020 | Thailand | Human | NA | 104 | 2 | This_Project | PRJNA1210814 |
| STC72 | 2021 | Thailand | Human | NA | 104 | 2 | This_Project | PRJNA1210814 |
| STC269 | 2021 | Thailand | Human | NA | 104 | 2 | This_Project | PRJNA1210814 |
| STC266 | 2021 | Thailand | Human | NA | 104 | 2 | This_Project | PRJNA1210814 |
| STC268 | 2021 | Thailand | Human | NA | 104 | 2 | This_Project | PRJNA1210814 |
| STC267 | 2021 | Thailand | Human | NA | 104 | 2 | This_Project | PRJNA1210814 |
| STC262 | 2021 | Thailand | Human | NA | 104 | 2 | This_Project | PRJNA1210814 |
| STC270 | 2021 | Thailand | Human | NA | 104 | 2 | This_Project | PRJNA1210814 |
| 30389 | 2009 | Thailand | Human | Sepsis | 233 | 2 | This_Project | PRJNA1210814 |

| | | | | | | | | |
|---|---|---|---|---|---|---|---|---|
| 27968 | 2008 | Thailand | Human | Sepsis | 233 | 2 | This_Project | PRJNA1210814 |
| 30823 | 2009 | Thailand | Human | Sepsis | 233 | 2 | This_Project | PRJNA1210814 |
| 33394 | 2010 | Thailand | Human | Sepsis | 233 | 2 | This_Project | PRJNA1210814 |
| 35940 | 2011 | Thailand | Human | Sepsis | 233 | 2 | This_Project | PRJNA1210814 |
| 37880 | 2011 | Thailand | Human | Sepsis | 379 | 2 | This_Project | PRJNA1210814 |
| 35949 | 2011 | Thailand | Human | Sepsis | 233 | 2 | This_Project | PRJNA1210814 |
| 35688 | 2011 | Thailand | Human | Sepsis | 233 | 2 | This_Project | PRJNA1210814 |
| 37554 | 2011 | Thailand | Human | Sepsis | 233 | 2 | This_Project | PRJNA1210814 |
| 39236 | 2012 | Thailand | Human | Sepsis | 233 | 2 | This_Project | PRJNA1210814 |
| 42527 | 2012 | Thailand | Human | Sepsis | 233 | 2 | This_Project | PRJNA1210814 |
| 40750 | 2012 | Thailand | Human | Sepsis | 233 | 2 | This_Project | PRJNA1210814 |
| 40804 | 2012 | Thailand | Human | Sepsis | 233 | 2 | This_Project | PRJNA1210814 |
| 41477 | 2012 | Thailand | Human | Sepsis | 233 | 2 | This_Project | PRJNA1210814 |
| 41219 | 2012 | Thailand | Human | Sepsis | 233 | 2 | This_Project | PRJNA1210814 |
| STC223 | 2014 | Thailand | Human | NA | 233 | 2 | This_Project | PRJNA1210814 |
| STC233 | 2015 | Thailand | Human | NA | 233 | 2 | This_Project | PRJNA1210814 |
| STC235 | 2015 | Thailand | Human | NA | 233 | 2 | This_Project | PRJNA1210814 |
| STC239 | 2016 | Thailand | Human | NA | 233 | 2 | This_Project | PRJNA1210814 |
| STC243 | 2016 | Thailand | Human | NA | 233 | 2 | This_Project | PRJNA1210814 |
| STC250 | 2017 | Thailand | Human | NA | 233 | 2 | This_Project | PRJNA1210814 |
| STC257 | 2019 | Thailand | Human | NA | 379 | 2 | This_Project | PRJNA1210814 |
| STC274 | 2019 | Thailand | Human | NA | 379 | 2 | This_Project | PRJNA1210814 |
| STC279 | 2020 | Thailand | Human | NA | 233 | 2 | This_Project | PRJNA1210814 |
| STC286 | 2020 | Thailand | Human | NA | 233 | 2 | This_Project | PRJNA1210814 |
| STC280 | 2020 | Thailand | Human | NA | 379 | 2 | This_Project | PRJNA1210814 |
| STC141 | 2021 | Thailand | Human | NA | 233 | 2 | This_Project | PRJNA1210814 |

| | | | | | | | | |
|---|---|---|---|---|---|---|---|---|
| STC135 | 2021 | Thailand | Human | NA | 379 | 2 | This_Project | PRJNA1210814 |
| STC31 | 2021 | Thailand | Human | NA | | 2 | This_Project | PRJNA1210814 |
| STC212 | 2021 | Thailand | Human | NA | 233 | 2 | This_Project | PRJNA1210814 |
| STC173 | 2021 | Thailand | Human | NA | 233 | 2 | This_Project | PRJNA1210814 |
| STC260 | 2021 | Thailand | Human | NA | 1656 | 2 | This_Project | PRJNA1210814 |
| 36375 | 2011 | Thailand | Human | Sepsis | 395 | 2 | This_Project | PRJNA1210814 |
| 34537 | 2010 | Thailand | Pig Healthy | Healthy | 104 | 2 | This_Project | PRJNA1210814 |
| 30346 | 2009 | Thailand | Human | Endocarditis | 233 | 2 | This_Project | PRJNA1210814 |
| 36602 | 2011 | Thailand | Human | NA | 379 | 2 | This_Project | PRJNA1210814 |
| 36610 | 2011 | Thailand | Human | Sepsis | 391 | 2 | This_Project | PRJNA1210814 |
| 38727 | 2011 | Thailand | Human | Sepsis | 233 | 2 | This_Project | PRJNA1210814 |
| 40301 | 2012 | Thailand | Human | Sepsis | 233 | 2 | This_Project | PRJNA1210814 |
| STC149 | 2021 | Thailand | Human | NA | New | 14 | This_Project | PRJNA1210814 |
| STC228 | 2014 | Thailand | Human | NA | 104 | 2 | This_Project | PRJNA1210814 |
| STC240 | 2016 | Thailand | Human | NA | 379 | 2 | This_Project | PRJNA1210814 |
| STC251 | 2019 | Thailand | Human | NA | 104 | 2 | This_Project | PRJNA1210814 |
| SAMN13172488 | 2007 | Thailand | Human | Sepsis | 104 | 2 | [2] | PRJNA575754 |
| SAMN12911095 | 2007 | Thailand | Pig Diseased | Sepsis | NA | 2 | [2] | PRJNA575754 |
| SAMN29502575 | 2010 | Thailand | Human | Meningitis | 104 | 2 | [3] | PRJNA418954 |
| SAMN29502582 | 2013 | Thailand | Pig Healthy | Healthy | 104 | 2 | [3] | PRJNA418954 |
| SAMN29502583 | 2013 | Thailand | Pig Healthy | Healthy | 104 | 2 | [3] | PRJNA418954 |
| SAMD00039909 | 2000 | Thailand | Human | NA | NA | 2 | NA | PRJDB4183 |
| SAMN17277761 | 2008 | Thailand | Human | Sepsis | 104 | 2 | [30] | PRJNA691075 |
| SAMN17277762 | 2012 | Thailand | Human | Sepsis | 233 | 2 | [30] | PRJNA691075 |
| SAMN17277763 | 2012 | Thailand | Human | Meningitis | 233 | 2 | [30] | PRJNA691075 |
| SAMN17277764 | 2012 | Thailand | Human | Sepsis | 379 | 2 | [30] | PRJNA691075 |

| | | | | | | | | |
|---|---|---|---|---|---|---|---|---|
| SAMN17277765 | 2012 | Thailand | Human | Meningitis | 233 | 2 | [30] | PRJNA691075 |
| SAMN17277766 | 2011 | Thailand | Human | Sepsis | 233 | 2 | [30] | PRJNA691075 |
| SAMN18585446 | 2007 | Thailand | Human | Sepsis | 104 | 2 | [30] | PRJNA691075 |
| SAMEA111506260 | 2020 | Thailand | Human | NA | NA | 2 | [30] | PRJNA691075 |
| STC78 | 2021 | Thailand | Human | Sepsis | 1656 | 2 | [1] | PRJNA853715 |
| STC130 | 2021 | Thailand | Human | Sepsis | 379 | 2 | [1] | PRJNA853715 |
| STC131 | 2021 | Thailand | Human | Sepsis | 233 | 2 | [1] | PRJNA853715 |
| STC152 | 2021 | Thailand | Human | Meningitis | 104 | 2 | [1] | PRJNA853715 |
| STC153 | 2021 | Thailand | Human | NA | 233 | 2 | [1] | PRJNA853715 |
| STC407 | 2021 | Thailand | Human | Sepsis | NA | 2 | [27] | NA |
| SAMN11854239 | 2015 | USA | Pig Diseased | NA | 225 | 7 | [17] | PRJNA544735 |
| SAMN04376555 | NA | Canada | Pig Diseased | NA | 225 | 7 | [17] | PRJNA307190 |
| SAMN11854154 | NA | USA | Pig Diseased | NA | 225 | 7 | [17] | PRJNA544735 |
| SAMN11854151 | 2014 | USA | Pig Diseased | NA | 225 | 7 | [17] | PRJNA544735 |
| SAMEA3233991 | 2010 | UK | Pig Diseased | NA | 907 | 7 | [17] | PRJEB8392 |
| SAMN04376560 | NA | Canada | Pig Diseased | NA | 32 | 7 | [17] | PRJNA307190 |
| SAMN02469560 | 2012 | China | Pig Healthy | NA | 32 | 7 | [17] | PRJNA171476 |
| SAMN13975591 | 2015 | USA | Pig Unknown | NA | NA | 7 | NA | PRJNA604583 |
| SAMN17982943 | 2016 | China | Pig Diseased | NA | 225 | 7 | NA | PRJNA702910 |
| SAMN17982944 | NA | China | Pig Diseased | NA | 225 | 7 | NA | PRJNA702910 |
| SAMN17982945 | NA | China | Pig Healthy | NA | 225 | 7 | NA | PRJNA702910 |
| SAMN18117670 | 2019 | Canada | Pig Diseased | Meningitis | 1614 | 7 | NA | PRJNA702910 |
| SAMN18117671 | 2019 | Canada | Pig Diseased | NA | 32 | 7 | NA | PRJNA702910 |
| SAMN18117673 | 2018 | Canada | Pig Diseased | Meningitis | 32 | 7 | NA | PRJNA702910 |
| SAMN20087851 | 2012 | China | Pig Healthy | Healthy | 32 | 7 | NA | PRJNA702910 |
| SAMD00009282 | 1998 | Thailand | Human | Meningitis | 104 | 2 | NA | PRJDB1485 |

| | | | | | | | | |
|---|---|---|---|---|---|---|---|---|
| SAMD00009289 | 2002 | Thailand | Human | Pulmonary Edema | 104 | 2 | NA | PRJDB1485 |
| SS-194 | 2018 | Thailand | Pig Diseased | Respiratory | 104 | 7 | [26] | NA |
| SS-500 | 2020 | Thailand | Pig Diseased | Respiratory | 104 | 7 | [26] | NA |

**Supplementary Table 8**. Isolates selected for serotype 2 capsule locus analysis. NA stands for not Available. CC stands for Clonal complex. UK stands for United Kingdom. USA stands for United States of America.

| Isolate name | Year | Country | Host | CC | Serotype | Reference | Bioproject |
|---|---|---|---|---|---|---|---|
| 22696 | 2006 | Thailand | Human | 104 | 2 | This_project | PRJNA1210814 |
| 22714 | 2006 | Thailand | Human | 25 | 2 | This_project | PRJNA1210814 |
| 23163 | 2006 | Thailand | Human | 104 | 2 | This_project | PRJNA1210814 |
| 23164 | 2006 | Thailand | Human | 28 | 2 | This_project | PRJNA1210814 |
| 24526 | 2007 | Thailand | Human | 1 | 2 | This_project | PRJNA1210814 |
| 24529 | 2007 | Thailand | Human | 104 | 2 | This_project | PRJNA1210814 |
| 25777 | 2007 | Thailand | Human | 104 | 2 | This_project | PRJNA1210814 |
| 26127 | 2008 | Thailand | Human | 104 | 2 | This_project | PRJNA1210814 |
| 26647 | 2008 | Thailand | Human | 104 | 2 | This_project | PRJNA1210814 |
| 27103 | 2008 | Thailand | Human | 25 | 2 | This_project | PRJNA1210814 |
| 27968 | 2008 | Thailand | Human | 233 | 2 | This_project | PRJNA1210814 |
| 29281 | 2008 | Thailand | Human | 104 | 2 | This_project | PRJNA1210814 |
| 30341 | 2009 | Thailand | Human | 104 | 2 | This_project | PRJNA1210814 |
| 30346 | 2009 | Thailand | Human | 233 | 2 | This_project | PRJNA1210814 |
| 30389 | 2009 | Thailand | Human | 233 | 2 | This_project | PRJNA1210814 |
| 30823 | 2009 | Thailand | Human | 233 | 2 | This_project | PRJNA1210814 |
| 31095 | 2009 | Thailand | Human | 25 | 2 | This_project | PRJNA1210814 |
| 31106 | 2009 | Thailand | Human | 104 | 2 | This_project | PRJNA1210814 |
| 32861 | 2010 | Thailand | Human | 104 | 2 | This_project | PRJNA1210814 |
| 33029 | 2010 | Thailand | Human | 104 | 2 | This_project | PRJNA1210814 |
| 33243 | 2010 | Thailand | Human | 25 | 2 | This_project | PRJNA1210814 |
| 33394 | 2010 | Thailand | Human | 233 | 2 | This_project | PRJNA1210814 |
| 34384 | 2010 | Thailand | Pig Healthy | 1 | 2 | This_project | PRJNA1210814 |

| | | | | | | | |
|---|---|---|---|---|---|---|---|
| 34537 | 2010 | Thailand | Pig Healthy | 104 | 2 | This_project | PRJNA1210814 |
| 34584 | 2011 | Thailand | Human | 29 | 2 | This_project | PRJNA1210814 |
| 34758 | 2011 | Thailand | Human | 25 | 2 | This_project | PRJNA1210814 |
| 35688 | 2011 | Thailand | Human | 233 | 2 | This_project | PRJNA1210814 |
| 35940 | 2011 | Thailand | Human | 233 | 2 | This_project | PRJNA1210814 |
| 35949 | 2011 | Thailand | Human | 233 | 2 | This_project | PRJNA1210814 |
| 36363 | 2011 | Thailand | Human | 1 | 2 | This_project | PRJNA1210814 |
| 36369 | 2011 | Thailand | Human | 104 | 2 | This_project | PRJNA1210814 |
| 36375 | 2011 | Thailand | Human | 25 | 2 | This_project | PRJNA1210814 |
| 36602 | 2011 | Thailand | Human | 233 | 2 | This_project | PRJNA1210814 |
| 36610 | 2011 | Thailand | Human | 104 | 2 | This_project | PRJNA1210814 |
| 36900 | 2011 | Thailand | Human | 29 | 2 | This_project | PRJNA1210814 |
| 36921 | 2011 | Thailand | Human | 104 | 2 | This_project | PRJNA1210814 |
| 37554 | 2011 | Thailand | Human | 233 | 2 | This_project | PRJNA1210814 |
| 37880 | 2011 | Thailand | Human | 233 | 2 | This_project | PRJNA1210814 |
| 38727 | 2011 | Thailand | Human | 233 | 2 | This_project | PRJNA1210814 |
| 39236 | 2012 | Thailand | Human | 233 | 2 | This_project | PRJNA1210814 |
| 39971 | 2012 | Thailand | Human | 104 | 2 | This_project | PRJNA1210814 |
| 40128 | 2012 | Thailand | Human | 1 | 2 | This_project | PRJNA1210814 |
| 40301 | 2012 | Thailand | Human | 233 | 2 | This_project | PRJNA1210814 |
| 40302 | 2012 | Thailand | Human | 233 | 2 | This_project | PRJNA1210814 |
| 40414 | 2012 | Thailand | Human | 104 | 2 | This_project | PRJNA1210814 |
| 40417 | 2012 | Thailand | Human | 25 | 2 | This_project | PRJNA1210814 |
| 40750 | 2012 | Thailand | Human | 233 | 2 | This_project | PRJNA1210814 |
| 40804 | 2012 | Thailand | Human | 233 | 2 | This_project | PRJNA1210814 |
| 41219 | 2012 | Thailand | Human | 233 | 2 | This_project | PRJNA1210814 |

| | | | | | | | |
|---|---|---|---|---|---|---|---|
| 41477 | 2012 | Thailand | Human | 233 | 2 | This_project | PRJNA1210814 |
| 42319 | 2012 | Thailand | Human | 28 | 2 | This_project | PRJNA1210814 |
| 42527 | 2012 | Thailand | Human | 233 | 2 | This_project | PRJNA1210814 |
| 1654537 | 2014 | Canada | Pig Diseased | 28 | 2 | [20] | PRJNA628943 |
| 1666725 | 2014 | Canada | Pig Unknown | 25 | 2 | [20] | PRJNA628943 |
| 108-54D_S51_L001 | 2018 | USA | Pig Diseased | 28 | 2 | [16] | PRJNA972671 |
| 19632_PH2016-048 | 2016 | Germany | Pig Diseased | 1 | 2 | [16] | PRJNA1009400 |
| 19635_PH2016-052 | 2016 | Germany | Pig Diseased | 28 | 2 | [16] | PRJNA1009400 |
| 19659_PH2016-077 | 2016 | Germany | Pig Diseased | 28 | 2 | [16] | PRJNA1009400 |
| 19718_PH2016-139 | 2016 | Germany | Pig Healthy | 28 | 2 | [16] | PRJNA1009400 |
| 19806_M105244_S28 | 2017 | Spain | Pig Diseased | 1 | 2 | [16] | PRJNA1009400 |
| 19807_M105244_S29 | 2017 | Spain | Pig Diseased | 1 | 2 | [16] | PRJNA1009400 |
| 19859_M105435_R31 | 2017 | Spain | Pig Diseased | 28 | 2 | [16] | PRJNA1009400 |
| 19860_M105455_R32 | 2017 | Spain | Pig Diseased | 28 | 2 | [16] | PRJNA1009400 |
| 19893_SS15055_N2_C15 | 2015 | Spain | Wild Boar Healthy | 1 | 2 | [16] | PRJNA1009400 |
| 21147_DNR14 | 2018 | Denmark | Pig Diseased | 28 | 2 | [16] | PRJNA1009400 |
| 21166_DNR33 | 2018 | Denmark | Pig Diseased | 28 | 2 | [16] | PRJNA1009400 |
| 21210_DNS27 | 2018 | Denmark | Pig Diseased | 1 | 2 | [16] | PRJNA1009400 |
| 21213_DNS30 | 2018 | Denmark | Pig Diseased | 28 | 2 | [16] | PRJNA1009400 |
| 21302_NLR18 | 2017 | Netherlands | Pig Diseased | 1 | 2 | [16] | PRJNA1009400 |
| 21321_NLR37 | 2017 | Netherlands | Pig Diseased | 28 | 2 | [16] | PRJNA1009400 |
| 21331_NLR47 | 2017 | Netherlands | Pig Diseased | 28 | 2 | [16] | PRJNA1009400 |
| 21413_NLC29 | 2017 | Netherlands | Pig Healthy | 28 | 2 | [16] | PRJNA1009400 |
| 28284_HSPI233 | 1997 | Spain | Pig Unknown | 28 | 2 | [16] | PRJNA1009400 |
| 28313_HNL2852 | 1983 | Netherlands | Pig Diseased | 20 | 2 | [16] | PRJNA1009400 |
| 28330_201988791 | 2004 | Denmark | Pig Diseased | 1 | 2 | [16] | PRJNA1009400 |

| | | | | | | | |
|---|---|---|---|---|---|---|---|
| 28349_2019887920 | 2005 | Denmark | Pig Diseased | 1 | 2 | [16] | PRJNA1009400 |
| 33148_Y0515 | 1989 | UK | Pig Diseased | 1 | 2 | [16] | PRJNA1009400 |
| 33151_Y0642 | 1990 | UK | Pig Diseased | 25 | 2 | [16] | PRJNA1009400 |
| 4057-28_S28_L001 | 2019 | USA | Pig Diseased | 25 | 2 | [16] | PRJNA972671 |
| 4057-46_S46_L001 | 2019 | USA | Pig Diseased | 25 | 2 | [16] | PRJNA972671 |
| 4057-48_S48_L001 | 2019 | USA | Pig Diseased | 25 | 2 | [16] | PRJNA972671 |
| 89-1591 | 1989 | Canada | Pig Unknown | 25 | 2 | [20] | PRJNA628943 |
| 89-6891-2 | 1989 | Canada | Pig Diseased | 25 | 2 | [20] | PRJNA628943 |
| BM358a | 2003 | Vietnam | Human | 1 | 2 | [34] | PRJEB2670 |
| CZ17 | 2005 | Czech Republic | Human | 87 | 2 | [38] | PRJNA917482 |
| CZ4 | 2001 | Czech Republic | Human | 25 | 2 | [38] | PRJNA917482 |
| CZ7 | 2003 | Czech Republic | Human | 25 | 2 | [38] | PRJNA917482 |
| CZ8 | 2004 | Czech Republic | Human | 25 | 2 | [38] | PRJNA917482 |
| D15-037104 | NA | USA | Pig Unknown | 28 | 2 | [38] | PRJNA628943 |
| D15-049887 | NA | USA | Pig Unknown | 28 | 2 | [38] | PRJNA628943 |
| DE12 | 1990 | Germany | Human | 94 | 2 | [38] | PRJNA917482 |
| DE13 | NA | Germany | Human | 1 | 2 | [38] | PRJNA917482 |
| DE19 | 2002 | Germany | Human | 1 | 2 | [38] | PRJNA917482 |
| DE8 | 2008 | Germany | Human | 87 | 2 | [38] | PRJNA917482 |
| DE9 | 2008 | Germany | Human | 87 | 2 | [38] | PRJNA917482 |
| ERR1055554 | 1986 | Netherlands | Human | 20 | 2 | [35] | PRJEB11219 |
| ERR1055566 | 1997 | Netherlands | Human | 1 | 2 | [35] | PRJEB11219 |
| ERR1055586 | 1996 | Netherlands | Pig Diseased | 20 | 2 | [35] | PRJEB11219 |
| ERR1055628 | 2006 | Netherlands | Pig Diseased | 1 | 2 | [3536] | PRJEB11219 |
| LSS25 | 2011 | UK | Pig Healthy | 28 | 2 | [38] | PRJEB8392 |
| MA4T3-3A | 2016 | Canada | Pig Healthy | 28 | 2 | [20] | PRJNA628943 |

| | | | | | | | |
|---|---|---|---|---|---|---|---|
| MY1C3-1B | 2016 | Canada | Pig Healthy | 28 | 2 | [20] | PRJNA628943 |
| SAMD00009297 | NA | Japan | Human | 1 | 2 | [17] | PRJDB1485 |
| SAMD00039896 | 2000 | Thailand | Human | 25 | 2 | NA | PRJDB4183 |
| SAMD00039897 | 2000 | Thailand | Human | 25 | 2 | NA | PRJDB4183 |
| SAMD00039900 | 2001 | Thailand | Human | 25 | 2 | NA | PRJDB4183 |
| SAMD00039902 | 2002 | Thailand | Human | 25 | 2 | NA | PRJDB4183 |
| SAMD00039903 | 2002 | Thailand | Human | 25 | 2 | NA | PRJDB4183 |
| SAMD00039904 | 2002 | Thailand | Human | NA | 2 | NA | PRJDB4183 |
| SAMD00039906 | 2002 | Thailand | Human | 25 | 2 | NA | PRJDB4183 |
| SAMD00039907 | 2001 | Thailand | Human | 25 | 2 | NA | PRJDB4183 |
| SAMD00039908 | 2001 | Thailand | Human | 25 | 2 | NA | PRJDB4183 |
| SAMD00039909 | 2000 | Thailand | Human | 104 | 2 | NA | PRJDB4183 |
| SAMD00039911 | 2002 | Thailand | Human | 28 | 2 | NA | PRJDB4183 |
| SAMEA4355195 | 2012 | Togo | Human | 1 | 2 | [17] | PRJEB12952 |
| SAMEA4355196 | 2012 | Togo | Human | 1 | 2 | [17] | PRJEB12952 |
| SAMN03652189 | 2002 | Thailand | Human | 28 | 2 | [6] | PRJNA283528 |
| SAMN04231436 | 2000 | Thailand | Human | 25 | 2 | [5] | PRJNA301006 |
| SAMN04231437 | 2000 | Thailand | Human | 25 | 2 | [5] | PRJNA301006 |
| SAMN04231442 | 2002 | Thailand | Human | 25 | 2 | [5] | PRJNA301006 |
| SAMN05442340 | 2013 | Argentina | Human | 1 | 2 | [17] | PRJNA335427 |
| SAMN05442342 | 2014 | Argentina | Human | 1 | 2 | [17] | PRJNA335427 |
| SAMN05442343 | 2012 | Argentina | Human | 1 | 2 | [17] | PRJNA335427 |
| SAMN05442344 | 2014 | China | Human | 1 | 2 | [17] | PRJNA335427 |
| SAMN08238506 | 2016 | China | Human | 1 | 2 | [17] | PRJNA427455 |
| SAMN08435268 | 2016 | China | Human | 1 | 2 | [17] | PRJNA432119 |
| SAMN12911095 | 2007 | Thailand | Pig Diseased | 104 | 2 | [2] | PRJNA575754 |

| | | | | | | | |
|---|---|---|---|---|---|---|---|
| SAMN12911096 | 2012 | Thailand | Pig Diseased | 25 | 2 | [2] | PRJNA575754 |
| SAMN13172488 | 2007 | Thailand | Human | 104 | 2 | [2] | PRJNA575754 |
| SAMN17277760 | 2009 | Thailand | Human | 25 | 2 | [30] | PRJNA691075 |
| SAMN17277761 | 2008 | Thailand | Human | 104 | 2 | [30] | PRJNA691075 |
| SAMN17277762 | 2012 | Thailand | Human | 233 | 2 | [30] | PRJNA691075 |
| SAMN17277763 | 2012 | Thailand | Human | 233 | 2 | [30] | PRJNA691075 |
| SAMN17277764 | 2012 | Thailand | Human | 233 | 2 | [30] | PRJNA691075 |
| SAMN17277766 | 2011 | Thailand | Human | 233 | 2 | [30] | PRJNA691075 |
| SRR7369587 | 2010 | Australia | Pig Unknown | 1 | 2 | [33] | PRJNA476804 |
| SRR7369589 | NA | Australia | Pig Unknown | 1 | 2 | [33] | PRJNA476804 |
| SRR7369624 | 2016 | Australia | Pig Unknown | 28 | 2 | [33] | PRJNA476804 |
| SRR7369629 | 2016 | Australia | Pig Unknown | 25 | 2 | [33] | PRJNA476804 |
| SRR7369704 | 2014 | Australia | Pig Unknown | 1 | 2 | [33] | PRJNA476804 |
| SRR7369706 | 2014 | Australia | Pig Unknown | 25 | 2 | [33] | PRJNA476804 |
| SRR7369718 | 2016 | Australia | Pig Unknown | 28 | 2 | [33] | PRJNA476804 |
| SS1012 | 2010 | UK | Pig Diseased | 1 | 2 | [34] | PRJEB8392 |
| SS1036 | 2010 | UK | Pig Unknown | 1 | 2 | [34] | PRJEB8392 |
| SS1053 | 2010 | UK | Pig Diseased | 25 | 2 | [34] | PRJEB8392 |
| SS1064 | 2010 | UK | Pig Diseased | 25 | 2 | [34] | PRJEB8392 |
| SS15 | 2005 | Vietnam | Human | 1 | 2 | [34] | PRJEB2670 |
| SS19 | 2005 | Vietnam | Human | 1 | 2 | [34] | PRJEB2670 |
| STC123 | 2021 | Thailand | Human | 1 | 2 | This_project | PRJNA1210814 |
| STC135 | 2021 | Thailand | Human | 233 | 2 | This_project | PRJNA1210814 |
| STC141 | 2021 | Thailand | Human | 233 | 2 | This_project | PRJNA1210814 |
| STC147 | 2021 | Thailand | Human | 25 | 2 | This_project | PRJNA1210814 |
| STC152 | 2021 | Thailand | Human | 104 | 2 | This_project | PRJNA1210814 |

| | | | | | | | |
|---|---|---|---|---|---|---|---|
| STC173 | 2021 | Thailand | Human | 233 | 2 | This_project | PRJNA1210814 |
| STC212 | 2021 | Thailand | Human | 233 | 2 | This_project | PRJNA1210814 |
| STC223 | 2014 | Thailand | Human | 233 | 2 | This_project | PRJNA1210814 |
| STC228 | 2014 | Thailand | Human | 104 | 2 | This_project | PRJNA1210814 |
| STC231 | 2015 | Thailand | Human | 1 | 2 | This_project | PRJNA1210814 |
| STC233 | 2015 | Thailand | Human | 233 | 2 | This_project | PRJNA1210814 |
| STC235 | 2015 | Thailand | Human | 233 | 2 | This_project | PRJNA1210814 |
| STC238 | 2016 | Thailand | Human | 233 | 2 | This_project | PRJNA1210814 |
| STC239 | 2016 | Thailand | Human | 233 | 2 | This_project | PRJNA1210814 |
| STC240 | 2016 | Thailand | Human | 233 | 2 | This_project | PRJNA1210814 |
| STC242 | 2016 | Thailand | Human | 104 | 2 | This_project | PRJNA1210814 |
| STC243 | 2016 | Thailand | Human | 233 | 2 | This_project | PRJNA1210814 |
| STC246 | 2017 | Thailand | Human | 104 | 2 | This_project | PRJNA1210814 |
| STC250 | 2017 | Thailand | Human | 233 | 2 | This_project | PRJNA1210814 |
| STC251 | 2019 | Thailand | Human | 104 | 2 | This_project | PRJNA1210814 |
| STC253 | 2019 | Thailand | Human | 104 | 2 | This_project | PRJNA1210814 |
| STC256 | 2019 | Thailand | Human | 104 | 2 | This_project | PRJNA1210814 |
| STC257 | 2019 | Thailand | Human | 233 | 2 | This_project | PRJNA1210814 |
| STC260 | 2021 | Thailand | Human | 233 | 2 | This_project | PRJNA1210814 |
| STC262 | 2021 | Thailand | Human | 104 | 2 | This_project | PRJNA1210814 |
| STC266 | 2021 | Thailand | Human | 104 | 2 | This_project | PRJNA1210814 |
| STC267 | 2021 | Thailand | Human | 104 | 2 | This_project | PRJNA1210814 |
| STC268 | 2021 | Thailand | Human | 104 | 2 | This_project | PRJNA1210814 |
| STC269 | 2021 | Thailand | Human | 104 | 2 | This_project | PRJNA1210814 |
| STC270 | 2021 | Thailand | Human | 104 | 2 | This_project | PRJNA1210814 |
| STC274 | 2019 | Thailand | Human | 233 | 2 | This_project | PRJNA1210814 |

| | | | | | | | |
|---|---|---|---|---|---|---|---|
| STC275 | 2019 | Thailand | Human | 1 | 2 | This_project | PRJNA1210814 |
| STC279 | 2020 | Thailand | Human | 233 | 2 | This_project | PRJNA1210814 |
| STC280 | 2020 | Thailand | Human | 233 | 2 | This_project | PRJNA1210814 |
| STC286 | 2020 | Thailand | Human | 233 | 2 | This_project | PRJNA1210814 |
| STC290 | 2020 | Thailand | Human | 104 | 2 | This_project | PRJNA1210814 |
| STC291 | 2020 | Thailand | Human | 104 | 2 | This_project | PRJNA1210814 |
| STC292 | 2020 | Thailand | Human | 104 | 2 | This_project | PRJNA1210814 |
| STC293 | 2020 | Thailand | Human | 1 | 2 | This_project | PRJNA1210814 |
| STC31 | 2021 | Thailand | Human | 233 | 2 | This_project | PRJNA1210814 |
| STC407 | 2021 | Thailand | Human | 233 | 2 | This_project | PRJNA1210814 |
| STC72 | 2021 | Thailand | Human | 104 | 2 | This_project | PRJNA1210814 |
| TMW_SS053 | 2014 | UK | Pig Unknown | 28 | 2 | [36] | PRJNA628943 |
| TMW_SS105 | 2014 | UK | Pig Unknown | 28 | 2 | [36] | PRJNA628943 |
| UK3 | 2015 | Ireland | Human | 1 | 2 | [38] | PRJNA917482 |

### References

1 Brizuela J, Kajeekul R, Roodsant TJ, *et al.* *Streptococcus suis* outbreak caused by an emerging zoonotic strain with acquired multi-drug resistance in Thailand. *Microb Genomics* 2023; **9**. DOI:10.1099/mgen.0.000952.

2 Yongkiettrakul S, Wongsurawat T, Jenjaroenpun P, *et al.* Genome sequences of antibiotic-resistant *Streptococcus suis* strains isolated from human patients and diseased and asymptomatic pigs in Thailand. *Infect Genet Evol* 2021; **87**: 104674.

3 Kittiwan N, Calland JK, Mourkas E, *et al.* Genetic diversity and variation in antimicrobial-resistance determinants of non-serotype 2 *Streptococcus suis* isolates from healthy pigs. *Microb Genomics* 2022; **8**: 000882.

4 Fittipaldi N, Xu J, Lacouture S, *et al.* Lineage and Virulence of *Streptococcus suis* Serotype 2 Isolates from North America. *Emerg Infect Dis* 2011; **17**: 2239.

5 Athey TBT, Teatero S, Takamatsu D, *et al.* Population Structure and Antimicrobial Resistance Profiles of *Streptococcus suis* Serotype 2 Sequence Type 25 Strains. *PLOS ONE* 2016; **11**: e0150908.

6 Athey TBT, Auger J-P, Teatero S, *et al.* Complex Population Structure and Virulence Differences among Serotype 2 *Streptococcus suis* Strains Belonging to Sequence Type 28. *PLOS ONE* 2015; **10**: e0137760.

7 Zheng H, Du P, Qiu X, *et al.* Genomic comparisons of *Streptococcus suis* serotype 9 strains recovered from diseased pigs in Spain and Canada. *Vet Res* 2018; **49**: 1.

8 Prjibelski A, Antipov D, Meleshko D, Lapidus A, Korobeynikov A. Using SPAdes De Novo Assembler. *Curr Protoc Bioinforma* 2020; **70**: e120.

9 Bolger AM, Lohse M, Usadel B. Trimmomatic: a flexible trimmer for Illumina sequence data. *Bioinformatics* 2014; **30**: 2114–20.

10 Gurevich A, Saveliev V, Vyahhi N, Tesler G. QUAST: quality assessment tool for genome assemblies. *Bioinforma Oxf Engl* 2013; **29**: 1072–5.

11 Seemann T. Prokka: rapid prokaryotic genome annotation. *Bioinformatics* 2014; **30**: 2068–9.

12 Wick RR, Judd LM, Gorrie CL, Holt KE. Unicycler: Resolving bacterial genome assemblies from short and long sequencing reads. *PLOS Comput Biol* 2017; **13**: e1005595.

13 Tonkin-Hill G, MacAlasdair N, Ruis C, *et al.* Producing polished prokaryotic pangenomes with the Panaroo pipeline. *Genome Biol* 2020; **21**: 180.

14 Page AJ, Taylor B, Delaney AJ, *et al.* SNP-sites: rapid efficient extraction of SNPs from multi- FASTA alignments. *Microb Genomics* 2016; **2**: pp: e000056.

15 Nguyen L-T, Schmidt HA, von Haeseler A, Minh BQ. IQ-TREE: A Fast and Effective Stochastic Algorithm for Estimating Maximum-Likelihood Phylogenies. *Mol Biol Evol* 2015; **32**: 268–74.

16 Murray GGR, Hossain ASMdM, Miller EL, *et al.* The emergence and diversification of a zoonotic pathogen from within the microbiota of intensively farmed pigs. *Proc Natl Acad Sci* 2023; **120**: e2307773120.

17 Roodsant TJ, Van Der Putten BCL, Tamminga SM, Schultsz C, Van Der Ark KCH. Identification of *Streptococcus suis* putative zoonotic virulence factors: A systematic review and genomic meta-analysis. *Virulence* 2021; **12**: 2787–97.

18 Price MN, Dehal PS, Arkin AP. FastTree 2 – Approximately Maximum-Likelihood Trees for Large Alignments. *PLoS ONE* 2010; **5**: e9490.

19 Paradis E, Schliep K. ape 5.0: an environment for modern phylogenetics and evolutionary analyses in R. *Bioinformatics* 2019; **35**: 526–8.

20 Hadjirin NF, Miller EL, Murray GGR, *et al.* Large-scale genomic analysis of antimicrobial resistance in the zoonotic pathogen *Streptococcus suis*. *BMC Biol* 2021; **19**: 191.

21 Liu M, Li X, Xie Y, *et al.* ICEberg 2.0: an updated database of bacterial integrative and conjugative elements. *Nucleic Acids Res* 2019; **47**: D660–5.

22 Wishart DS, Han S, Saha S, *et al.* PHASTEST: faster than PHASTER, better than PHAST. *Nucleic Acids Res* 2023; **51**: W443–50.

23 Bertelli C, Laird MR, Williams KP, *et al.* IslandViewer 4: expanded prediction of genomic islands for larger-scale datasets. *Nucleic Acids Res* 2017; **45**: W30–5.

24 Ross K, Varani AM, Snesrud E, *et al.* TnCentral: a Prokaryotic Transposable Element Database and Web Portal for Transposon Analysis. *mBio* 2021; **12**: 10.1128/mbio.02060-21.

25 Boueroy P, Brizuela J, Roodsant TJ, *et al.* Genomic analysis and virulence of human *Streptococcus suis* serotype 14. *Eur J Clin Microbiol Infect Dis Off Publ Eur Soc Clin Microbiol* 2024; published online Dec 28. DOI:10.1007/s10096-024-05029-w.

26 Lunha K, Chumpol W, Jiemsup S, Samngamnim S, Assavacheep P, Yongkiettrakul S. Relationship between Penicillin-Binding Proteins Alterations and β-Lactams Non-Susceptibility of Diseased Pig-Isolated *Streptococcus suis*. *Antibiotics* 2023; **12**: 158.

27 Isolate information: id-2917 (STC407) - *Streptococcus suis* isolates. https://pubmlst.org/bigsdb?page=info&db=pubmlst_ssuis_isolates&id=2917 (accessed Jan 15, 2025).

28 Nag S, Larsen G, Szarvas J, *et al.* Whole genomes from bacteria collected at diagnostic units around the world 2020. *Sci Data* 2023; **10**: 628.

29 Kerdsin A, Hatrongjit R, Wongsurawat T, *et al.* Genomic Characterization of *Streptococcus suis* Serotype 24 Clonal Complex 221/234 From Human Patients. *Front Microbiol* 2021; **12**: 812436.

30 Bamphensin N, Chopjitt P, Hatrongjit R, *et al.* Non-Penicillin-Susceptible *Streptococcus suis* Isolated from Humans. *Pathog Basel Switz* 2021; **10**: 1178.

31 Hatrongjit R, Fittipaldi N, Jenjaroenpun P, *et al.* Genomic comparison of two *Streptococcus suis* serotype 1 strains recovered from porcine and human disease cases. *Sci Rep* 2023; **13**: 5380.

32 Zhu J, Wang J, Kang W, *et al. Streptococcus suis* serotype 4: a population with the potential pathogenicity in humans and pigs. *Emerg Microbes Infect*; **13**: 2352435.

33 O'Dea MA, Laird T, Abraham R, *et al.* Examination of Australian *Streptococcus suis* isolates from clinically affected pigs in a global context and the genomic characterisation of ST1 as a predictor of virulence. *Vet Microbiol* 2018; **226**: 31–40.

34 Weinert LA, Chaudhuri RR, Wang J, *et al.* Genomic signatures of human and animal disease in the zoonotic pathogen *Streptococcus suis*. *Nat Commun* 2015; **6**: 6740.

35 Willemse N, Howell KJ, Weinert LA, *et al.* An emerging zoonotic clone in the Netherlands provides clues to virulence and zoonotic potential of *Streptococcus suis*. *Sci Rep* 2016; **6**: 28984.

36 Wileman TM, Weinert LA, Howell KJ, *et al.* Pathotyping the Zoonotic Pathogen *Streptococcus suis*: Novel Genetic Markers To Differentiate Invasive Disease-Associated Isolates from Non-Disease-Associated Isolates from England and Wales. *J Clin Microbiol* 2019; **57**: e01712-18.

37 Zou G, Zhou J, Xiao R, *et al.* Effects of Environmental and Management-Associated Factors on Prevalence and Diversity of *Streptococcus suis* in Clinically Healthy Pig Herds in China and the United Kingdom. *Appl Environ Microbiol* 2018; **84**: e02590-17.

38 Brizuela J, Roodsant TJ, Hasnoe Q, *et al.* Molecular Epidemiology of Underreported Emerging Zoonotic Pathogen *Streptococcus suis* in Europe. *Emerg Infect Dis* 2024; **30**: 413–22.

39 Dong X, Chao Y, Zhou Y, *et al.* The global emergence of a novel *Streptococcus suis* clade associated with human infections. *EMBO Mol Med* 2021; **13**: e13810.

40 Guo G, Kong X, Du D, Wei D, Yu Y, Zhang W. Genome-wide association study identifies the virulence-associated marker in *Streptococcus suis* serotype 2. *Infect Genet Evol* 2021; **92**: 104894.
